# Refining the transcriptional landscapes for distinct clades of virulent phages infecting *pseudomonas aeruginosa*

**DOI:** 10.1101/2023.10.13.562163

**Authors:** Leena Putzeys, Laura Wicke, Maarten Boon, Vera van Noort, Jörg Vogel, Rob Lavigne

## Abstract

The introduction of high-throughput sequencing has resulted in a surge of available bacteriophage genomes, unveiling their tremendous genomic diversity. However, our current understanding of the complex transcriptional mechanisms that dictate their gene expression during infection is limited to a handful of model phages. Here, we applied ONT-cappable-seq to reveal the transcriptional architecture of six different clades of virulent phages infecting *Pseudomonas aeruginosa*. This long-read microbial transcriptomics approach is tailored to globally map transcription start and termination sites, transcription units and putative RNA-based regulators on dense phage genomes. Specifically, the full-length transcriptomes of LUZ19, LUZ24, 14-1, YuA, PAK_P3 and giant phage phiKZ during early, middle and late infection were collectively charted. Beyond pinpointing traditional promoter and terminator elements and transcription units, these transcriptional profiles provide insights in transcriptional attenuation and splicing events and allow straightforward validation of Group I intron activity. In addition, ONT-cappable-seq data can guide genome-wide discovery of novel regulatory element candidates, including non-coding RNAs and riboswitches. This work substantially expands the number of annotated phage-encoded transcriptional elements identified to date, shedding light on the intricate and diverse gene expression regulation mechanisms in *Pseudomonas* phages, which can ultimately be sourced as tools for biotechnological applications in phage and bacterial engineering.

## Introduction

Bacteriophages, viruses that infect bacteria, are the most abundant biological entities on our planet. The introduction of high-throughput sequencing technologies, has unveiled their ubiquitous nature and exceptional genomic diversity, which in turn has produced a growing catalogue of phage genomic sequences (Dion, Oechslin and Moineau 2020). According to the National Center for Biotechnology Information (NCBI), as of May 2023, more than 900 *Pseudomonas* phage genomes have been sequenced. The majority of these phages belong to the *Caudoviricetes* class of tailed phages and infect the human opportunistic pathogen *Pseudomonas aeruginosa*.

Based on the 2022 International Committee on Taxonomy of Viruses (ICTV) report, *Pseudomonas aeruginosa* phages now span over 20 different genera, further reflecting the widespread and diverse nature of their bacterial host (De Smet *et al*. 2017; Turner *et al*. 2023). *Pbunavirus, Pakpunavirus*, and *Phikmvvirus* currently represent the three lytic genera with the most members (Figure 1). Despite the large number of available phage genomes, in-depth knowledge on their transcriptional landscapes and gene regulation mechanisms remain scarce beyond a limited number of model phages (Yang *et al*. 2014). Yet, charting the transcriptome architectures of phages is key to fully understand the different layers of gene regulation during the infection process (Salmond and Fineran 2015; Hör, Gorski and Vogel 2018; Ofir and Sorek 2018).

**Figure 1:**
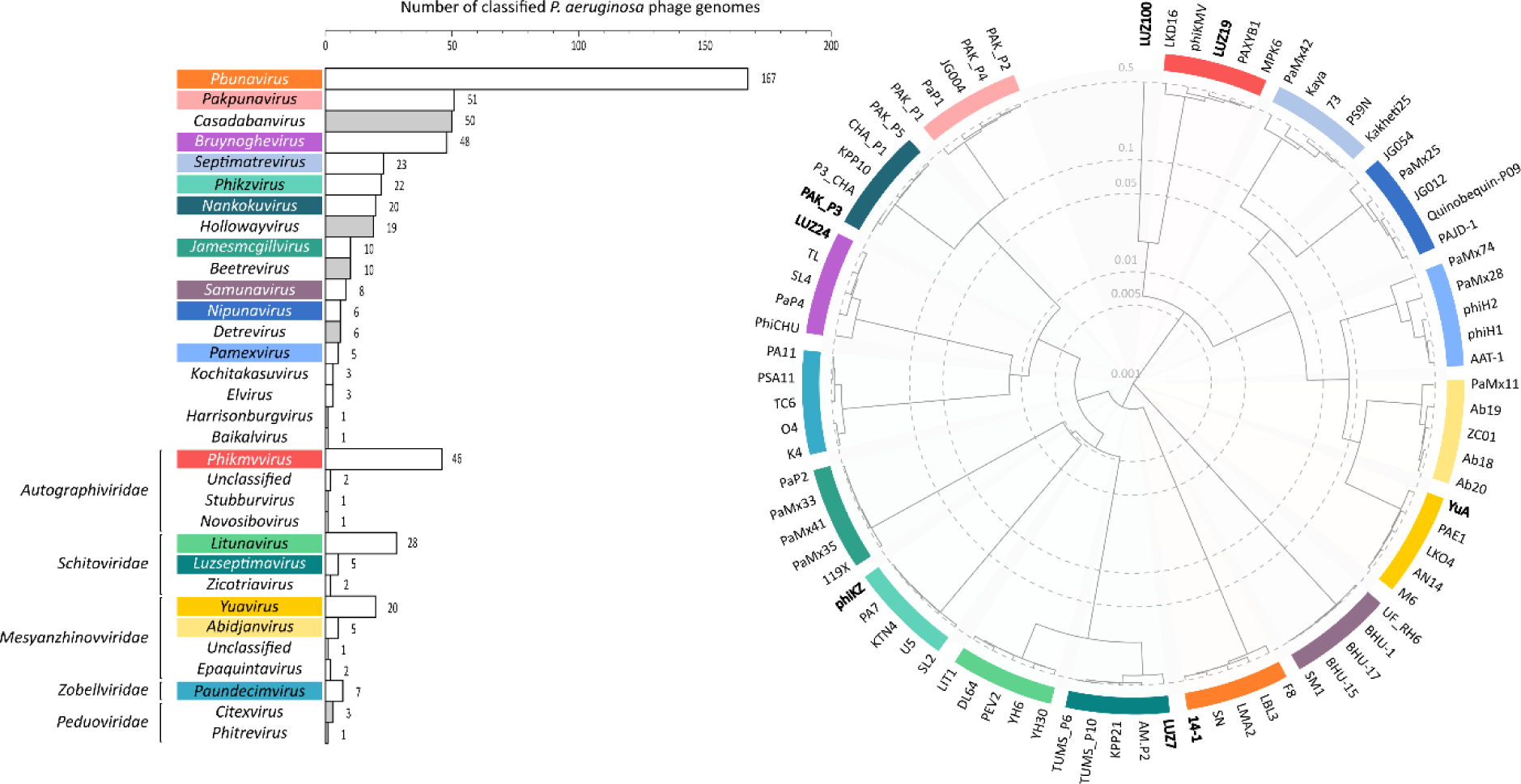
Classification and phylogenetic tree of lytic *Pseudomonas aeruginosa* phages. As of May 2023, the genomes of *P. aeruginosa*-infecting phages are classified in over 20 different genera according to the current ICTV taxonomy release (2022) (Turner *et al*. 2023). The bar plot (left panel) shows the number of classified *P. aeruginosa* phages in each genus with their respective family-level taxa, excluding phage families with less than two members. Genera associated with temperate phages are indicated in grey and are not shown in the phylogenetic tree. For the lytic phage genera with at least five members, five representative were selected and their genomes were used to construct a protein-level phylogenetic tree using VipTree (Nishimura *et al*. 2017) (right panel). Colours in the tree represent the corresponding genus in the bar plot. Branch lengths are logarithmically scaled and represent the genomic similarity scores (S_G_) of the phage genomes (normalized scores by TBLASTX). Representative phages used in this study, or sequenced previously using ONT-cappable-seq (LUZ7 and LUZ100), are indicated in bold.

In the last decade, phage transcriptomics research has largely been limited to classical short-read RNA-sequencing (RNA-seq) experiments, providing valuable insights in phage temporal gene expression patterns and host responses (Ceyssens *et al*. 2014; Blasdel *et al*. 2017; Yang *et al*. 2019; Kornienko *et al*. 2020; Li *et al*. 2020; Lood *et al*. 2020; Brandão *et al*. 2021). However, RNA-seq generally lacks the capacity to distinguish between primary and processed transcripts, which obscures the discovery of key transcriptional initiation and termination events at the original 5’ and 3’ boundaries of primary transcripts. To this end, specialised transcriptomic approaches have been developed that allow targeted sequencing of either 5’ or 3’ transcript ends (Sharma *et al*. 2010; Boutard *et al*. 2016; Dar *et al*. 2016; Ettwiller *et al*. 2016), or enable the profiling of primary prokaryotic transcripts in full-length (Yan *et al*. 2018; Putzeys *et al*. 2022). Of these, ONT-cappable-seq is a recent, long-read, nanopore-based cDNA sequencing method that permits end-to-end sequencing of primary prokaryotic transcripts, concurrently delineating both 5’ and 3’ RNA extremities, and revealing operon structures. Recently, ONT-cappable-seq was successfully introduced to study the transcriptomes and RNA biology of *Pseudomonas aeruginosa* phages LUZ7 (*Luzseptimavirus*) and LUZ100 (unclassified *Autographiviridae*) (Putzeys *et al*. 2022, 2023) and *Thermus thermophilus* phage P23-45 (*Oshimavirus*) (Chaban *et al*. 2022), yielding high-resolution genome-wide maps of their transcription start sites (TSS), transcription terminator sites (TTS) and transcription unit (TU) architectures.

In this work, ONT-cappable-seq is applied to profile the full-length transcriptomes of virulent *P. aeruginosa* phage representatives of major taxonomic clades (Figure 1), including LUZ19 (*Phikmvvirus*), LUZ24 (*Bruynoghevirus*), 14-1 (*Pbunavirus*), YuA (*Yuavirus*), PAK_P3 (*Nankokuvirus*) and giant phage phiKZ (*Phikzvirus*) (Ceyssens *et al*. 2008a, 2008b, 2009, 2014; Lavigne *et al*. 2013; Chevallereau *et al*. 2016). The transcriptional strategies of these phages are highly diverse and show various levels of dependency on the host transcriptional apparatus. While the majority of phages used in this study rely almost exclusively on the machinery of their host to initiate gene transcription, LUZ19 and phiKZ are equipped with their own RNA polymerase(s) and phage-specific promoter sequences. Using ONT-cappable-seq, we refined their distinct transcriptional architectures and discovered novel phage-encoded regulatory features, shedding light on the diversified and intricate transcriptional regulation mechanisms that *Pseudomonas* phages use to orchestrate their gene expression.

## Materials & methods

### Bacterial strains, growth conditions and bacteriophage propagation

*P. aeruginosa* strains PAO1 (DSM 22644) (Oberhardt *et al*. 2008), PAK (Takeya and Amako 1966) and Li010 (Pirnay *et al*. 2002) were cultured in Lysogeny Broth (LB) medium at 37°C. Six different lytic *P. aeruginosa* phages from different genera were selected to represent a diverse set of characterised phages (Table 1). PAO1 was used to amplify phages LUZ19, YuA, 14-1 and phiKZ. Alternatively, phages PAK_P3 and LUZ24 were amplified on host strains PAK and Li010, respectively. For phage amplification, the bacterial host was grown to early exponential phase (optical density of OD_600_ = 0.3) and infected with a high-titer lysate of the appropriate phage. Following overnight incubation at 37°C, the phage lysate was purified and concentrated using polyethylene glycol 8000 (PEG8000) precipitation. The resulting phage stocks were stored in phage buffer (10 mM NaCl, 10 mM MgSO_4_, 10 mM Tris-HCl, pH 7.5) at 4°C.

**Table 1:** Overview of the lytic *P. aeruginosa* phages used in this work. For each phage, infection conditions and RNA sampling timepoints used for ONT-cappable-seq are indicated. MOI: multiplicity of infection.

| Phage | Genus | host strain | MOI | Sampling timepoints post-infection (min) | Reference |
| --- | --- | --- | --- | --- | --- |
| **LUZ19** | *Phikmvvirus* | PAO1 | 75 | 5, 10, 15 | (Lavigne *et al.* 2013) |
| **YuA** | *Yuavirus* | PAO1 | 75 | 5, 15, 25, 45, 65 | (Ceyssens *et al.* 2008b) |
| **phiKZ** | *Phikzvirus* | PAO1 | 15 | 5, 10, 15, 20, 30, 40, 50, 60, 70, 80 | (Ceyssens *et al.* 2014) |
| **14-1** | *Pbunavirus* | PAO1 | 25 | 3, 6, 9, 12 | (Ceyssens *et al.* 2009) |
| **LUZ24** | *Bruynoghevirus* | Li010 | 50 | 5, 15, 25, 35 | (Ceyssens *et al.* 2008a) |
| **PAK_P3** | *Nankokuvirus* | PAK | 40 | 3.5, 6.5, 10, 13, 16.5 | (Chevallereau *et al.* 2016) |

### P. aeruginosa *Li010 genome extraction, sequencing and hybrid assembly*

The full genome of *P. aeruginosa* strain Li010 was sequenced using a combination of Illumina short-read sequencing and Nanopore sequencing technology. For this, High-molecular weight genomic DNA (gDNA) of Li010 was extracted using the DNeasy UltraClean Microbial Kit (Qiagen) according to the manufacturer’s guidelines. Afterwards, the DNA sample was prepared using the Illumina DNA prep kit (Illumina, USA) and sequenced on an Illumina MiniSeq device. Raw read quality of the Illumina data was assessed using FastQC (v0.11.8) (bioinformatics.babraham.ac.uk), after which adapters and poor-quality bases were removed using Trimmomatic (v0.39) (Bolger, Lohse and Usadel 2014). The Rapid Barcoding Sequencing Kit (SQK-RBK004) was used to prepare the DNA for nanopore sequencing, which was subsequently loaded on a MinION flow cell (FLO-MIN106, R9.4.1) and sequenced for 24–48h. The nanopore data was basecalled in high-accuracy mode using Guppy (v6.3.8) and processed using Porechop (v0.2.3). Next, both short-read and long-read sequencing datasets were integrated to perform *de novo* hybrid assembly of the Li010 genome using Unicycler (v0.4.8) with default settings (Wick *et al*. 2017). Finally, the resolved genome of Li010 was deposited in NCBI GenBank (CP124600) and used as the host reference for ONT-cappable-seq data analysis of phage LUZ24.

### Phage infection conditions and RNA extraction

Bacterial cultures were grown to an OD_600_ of 0.3 and infected at high multiplicity of infection (MOI, see Table 1) of the appropriate phages to ensure a high infection rate of the bacterial cells, before incubation at 37°C (Table 1) (Chevallereau *et al*. 2016; De Smet *et al*. 2016; Brandão *et al*. 2021; Wicke *et al*. 2021). Phage-infected culture samples were collected at multiple timepoints during the infection cycle of each phage, as indicated in Table 1. The collected samples were treated with stop mix solution (95% ethanol, 5% phenol, saturated, pH 4.5) and immediately snap-frozen in liquid nitrogen. Next, samples were thawed on ice and centrifuged for 20 minutes at 4°C at 4,000 *g*. The pellets were resuspended in 0.5 mg/mL lysozyme in Tris-EDTA solution (pH 8) to lyse the cells, after which RNA was isolated using hot phenol extraction. The crude RNA samples were subsequently purified using ethanol precipitation and subjected to DNAse I treatment, followed by another round of ethanol precipitation and spin-column purification. Successful removal of genomic DNA was verified by PCR using a specific primer pair that targets *P. aeruginosa* (Supplementary Table S1). Finally, the RNA sample integrity was assessed on an Agilent 2100 Bioanalyzer using the RNA 6000 Pico kit. Samples with an RNA integrity number (RIN) greater than 9 were used for downstream processing and sequencing.

### ONT-cappable-seq library preparation

For each phage, individual RNA samples were pooled together in equal amounts to a final concentration of 5 µg prior to library preparation. The resulting samples were supplemented with 1 ng of a control RNA spike-in, which was transcribed *in vitro* using the HiScribe T7 high-yield RNA synthesis kit (New England Biolabs). Afterwards, ONT-cappable-seq library preparation was carried out for all six phage samples as described in prior work (Putzeys *et al*. 2022), resulting for each phage in a cDNA sample that was enriched for primary transcripts and a control cDNA sample that did not undergo discrimination between primary and processed transcripts (Putzeys *et al*. 2022). Equimolar amounts of the twelve cDNA samples (LUZ19_enriched_, LUZ19_control_, YuA_enriched_, YuA_control_, phiKZ_enriched_, phiKZ_control_, 14-1_enriched_, 14-1_control_, LUZ24_enriched_, LUZ24_control_, PAK_P3_enriched_, PAK_P3_control_) were pooled to a total of 100 fmol in a 23 µL sample volume. Finally, nanopore sequencing adapters were added to the cDNA library, which was subsequently loaded on a PromethION flow cell (R9.4.1). The flow cell was run on the PromethION 24 platform with live base-calling and demultiplexing enabled. After ∼24h, the flow cell was reloaded and refuelled, after which sequencing carried on until all pores were exhausted (>48h).

### ONT-cappable-seq data analysis

After base-calling, only raw reads which passed the default Phred-like quality score threshold (≥7) were retained for downstream analysis. The passed raw sequencing data was evaluated in terms of sequencing yield, quality and read length using NanoComp (v1.11.2) (De Coster *et al*. 2018). Next, reads from each sample were mapped to their reference phage genomes, LUZ19 (NC_010326.1); YuA (NC_010116.1); phiKZ (NC_004629.1); 14-1 (NC_011703.1); LUZ24 (NC_010325.1); PAK_P3 (NC_022970.1)) and host (PAO1 (NC_002516.2); PAK (LR657304.1); Li010 (CP124600) as described previously (Putzeys *et al*. 2022). The genomic alignments were visually inspected using Integrated Genomics Viewer (IGV) (Thorvaldsdóttir, Robinson and Mesirov 2013). For each sample, a summary of the sequencing output, read lengths and mapping data is provided in Supplementary Table S2.

Afterwards, identification of viral TSSs and TTSs was carried out using our ONT-cappable-seq data analysis workflow (https://github.com/LoGT-KULeuven/ONT-cappable-seq) (Putzeys *et al*. 2022), tailored to the phages in this dataset. Briefly, TSSs were identified by finding genomic positions with a local maxima of 5’ read ends using a peak calling algorithm. Afterwards, for each peak position, the enrichment ratio was calculated by dividing the read count per million mapped reads (RPM) in the enriched sample by the corresponding RPM in the control sample. Peak positions that surpassed the enrichment ratio threshold (T_TSS_) and had at least 30 reads starting at that position in the enriched sample, followed by manual curation, were annotated as a TSS. The enrichment ratio thresholds varied for the individual phages (T_Tss_ LUZ19 = 212.7; T_TSS_ YuA = 86.6; T_TSS_ phiKZ = 30; T_TSS_ 14-1 = 5.1; T_TSS_ LUZ24 = 5; T_TSS_ PAK_P3 = 29.8), depending on the enrichment ratio observed for the TSS of the T7 promoter in the RNA spike-in in each sample (Putzeys *et al*. 2022). Next, regions upstream of the annotated TSSs were uploaded in MEME (-50 to +1) and SAPPHIRE.CNN (-100 to +1) to identify motifs and *Pseudomonas* σ70 promoter sequences, respectively (Bailey *et al*. 2015; Coppens, Wicke and Lavigne 2022). Similarly, phage TTSs were annotated by determining genomic positions with a local accumulation of 3’ read ends that showed an average read reduction of at least 20% across the termination site, as described by Putzeys et al. (2022). For each TTS identified by ONT-cappable-seq, the -60 to +40 terminator region was analysed with ARNold to predict intrinsic, factor-independent transcriptional terminators (Naville *et al*. 2011). RNAfold (v2.4.13) was used to predict and calculate the secondary structure and minimum free energy of the annotated terminator regions (Lorenz *et al*. 2011). Finally, transcription units (TUs) for each phage were delineated by adjacent TSS and TTS annotated on the same strand, upon validating that at least one ONT-cappable-seq read spans the candidate TU. Where no clear TSS-TTS pair could be defined, the longest read was used for TU boundary determination.

### In vivo *promoter activity assay*

A subset of the phage-encoded host-specific promoters was experimentally validated in *vivo* using the SEVAtile-based expression system, as described previously (Lammens *et al*. 2021; Putzeys *et al*. 2023). In short, promoters were cloned in a pBGDes vector backbone upstream of a standardised ribosomal binding site (*BCD2*) and an *msfgfp* reporter gene (Mutalik *et al*. 2013; Lammens *et al*. 2021). A construct without a promoter sequence (pBGDes *BCD2-msfGFP*) and a construct with a constitutive promoter (pBGDes *Pem7-BCD2-msfGFP*) were included as controls. The resulting vectors were introduced to *Escherichia coli* PIR2 cells using heat-shock transformation and selectively plated on LB supplemented with kanamycin (50 µg/µL) (Hanahan 1983). In addition, the genetic constructs were transformed to *P. aeruginosa* PAO1 host cells by co-electroporation with a pTNS2 helper plasmid and subsequent plating on LB supplemented with gentamicin (30 µg/mL) (Choi, Kumar and Schweizer 2006). All primers and genetic parts used in this work are listed in Supplementary Table S1. Next, four biological replicates of each sample were inoculated in M9 minimal medium (1× M9 salts (BD Biosciences), 2 mM MgSO_4_, 0.1 mM CaCl_2_ (Sigma-Aldrich), 0.5% casein amino acids (LabM, Neogen), and 0.2% citrate (Sigma-Aldrich)), complemented with the appropriate antibiotic, and incubated at 37°C. The following day, samples were diluted 1:10 in fresh M9 medium in a 96-well black polystyrene COSTAR plate (Corning) with a clear flat bottom and transferred to a CLARIOstar® *Plus* Microplate Reader (BMG Labtech). OD_600_ and msfGFP fluorescence intensity levels (485 nm (ex)/528 nm (em)) were measured every 15 min for 12 h, while incubating at 37 °C. The relative msfGFP measurements were normalised for their respective OD600 values and subsequently converted to absolute units of the calibrant 5(6)-carboxyfluorescein (5(6)-FAM)) (Sigma-Aldrich). Finally, the data was visualised and analysed using the statistical software JMP 16 Pro (SAS Institute Inc.).

### PCR-based splicing validation

Purified RNA samples of LUZ24 collected 5 min, 15 min, 25 min, and 35 min post-infection were used to confirm the presence of a second putative intron in the LUZ24 genome. For this, 200 ng of total RNA was mixed with 100 pmol of a sequence-specific primer (P_RT,intron2_). Reverse transcription was carried out with 100 units of Maxima H Minus Reverse Transcriptase (Thermo Fisher), after incubation for 10 min at 25°C, 30 min at 50°C, and heat inactivation for 5 min at 85°C. Next, the resulting cDNA, LUZ24 genomic DNA, and LUZ24 phage lysate was used for PCR amplification (primers P_F,PCR_,_intron2_ and P_R,PCR,intron2_; Supplementary Table S1) and visually compared on a 1.5% agarose gel. As a control, the same experiment was carried out for the previously identified group I intron using a different set of primers (P_F_,_PCR,intron1_, P_R,PCR,intron1_ and P_RT,intron1_) (Supplementary Figure S1) (Ceyssens *et al*. 2008a).

### Northern blotting

To visualize specific RNA transcripts of interest, 5 µg of total RNA of each sample was electrophoretically resolved on a 6% polyacrylamide gel containing 7M urea. After blotting on an Amersham Hybond-XL (GE Healthcare, Chicago, IL, USA) membrane at 50 V, 4 °C for 1 h, transcripts were detected by gene-specific ^32^P-labeled oligonucleotides (Supplementary Table S1) in hybridization buffer (G-Biosciences, Saint Louis, MI, USA) and exposed to a phosphor screen overnight. Images were visualized using a Typhoon 9400 (Variable Mode Imager, Amersham Biosciences, Amersham, United Kingdom). The pUC8 ladder reference was size matched and cut from their corresponding membrane after the first read out. This was due to gradual fading of the signal after multiple exposures.

## Results & discussion

### *The global transcriptional landscapes of* P. aeruginosa phages

ONT-cappable-seq enables end-to-end sequencing of primary prokaryotic transcripts, allowing the simultaneous delineation of 5’ and 3’ transcript boundaries (Putzeys *et al*. 2022). During the ONT-cappable-seq library preparation, primary transcripts are enriched by capping their hallmarking triphosphorylated 5’ RNA ends with a desthiobiotin tag that can be captured by streptavidin beads. In parallel, a non-enriched control sample is prepared where the streptavidin enrichment step is omitted, retaining all processed RNA species. As such, comparative transcriptomics between the enriched and control samples allows discrimination between transcription start sites (TSSs) and processed 5’ ends. In addition, full-length transcriptional profiling by ONT-cappable-seq can discover transcription termination sites (TTS), as primary transcripts are more likely to have their original 3’ end intact (Yan *et al*. 2018; Putzeys *et al*. 2022). For each individual phage, cellular transcripts from multiple infection stages were pooled prior to library preparation, after which the resulting enriched and control samples (n = 12) were multiplexed and sequenced on a PromethION device, generating 0.7-22 million reads per sample (Supplementary Table S2).

Using this approach, we obtained comprehensive transcriptional maps of taxonomically distinct phages 14-1 (66.2 kb), LUZ19 (43.5 kb), LUZ24 (45.6 kb), PAK_P3 (88.1 kb), YuA (58.7 kb) and phiKZ (280.3 kb), shedding light on their distinct transcriptional patterns and architectures (Figure 2). In addition, we identified transcription initiation and terminator sites across the genome for each phage, allowing the discovery of novel regulatory elements and refined phage genome annotations (Supplementary Tables S3-S5).

**Figure 2:**
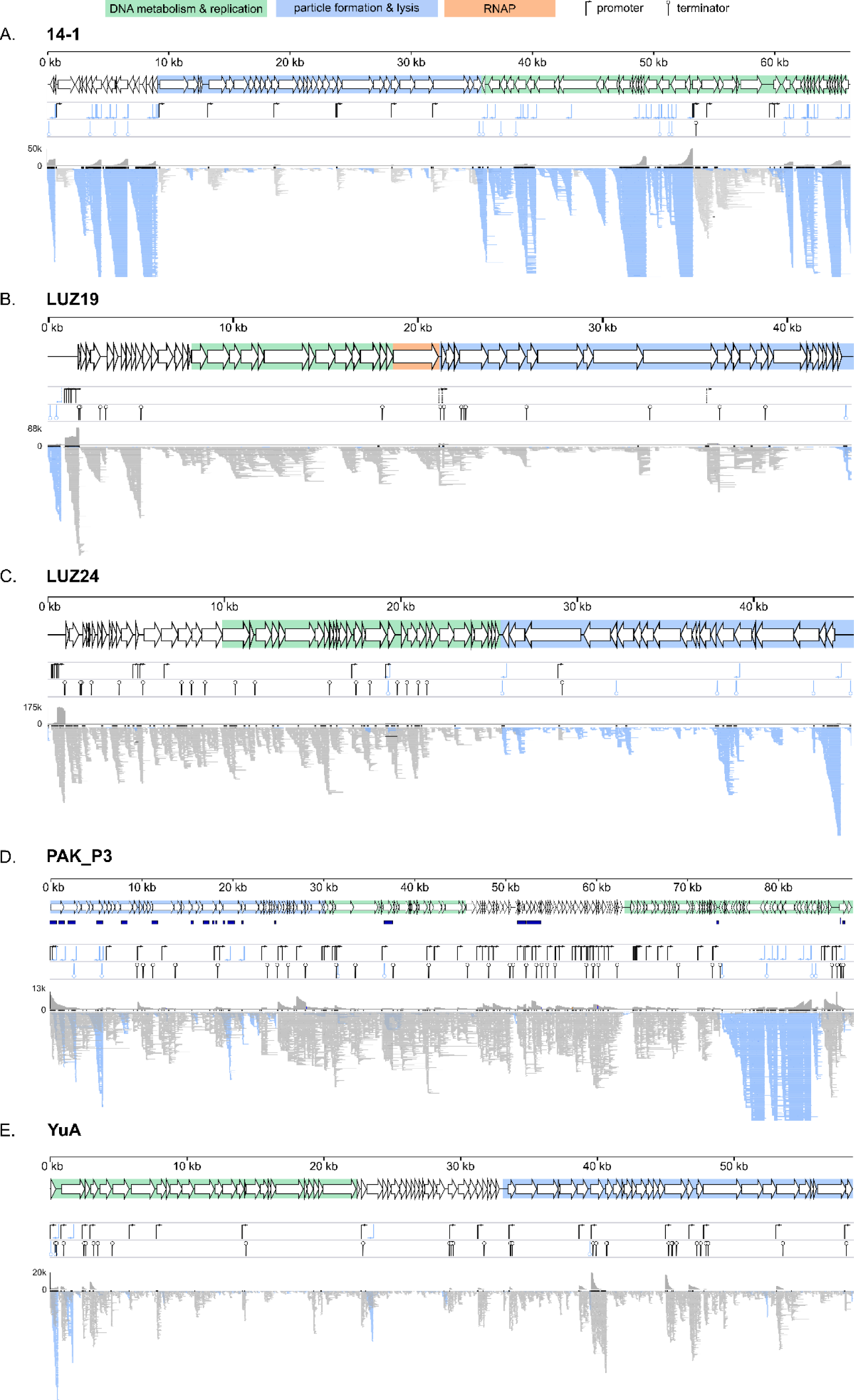
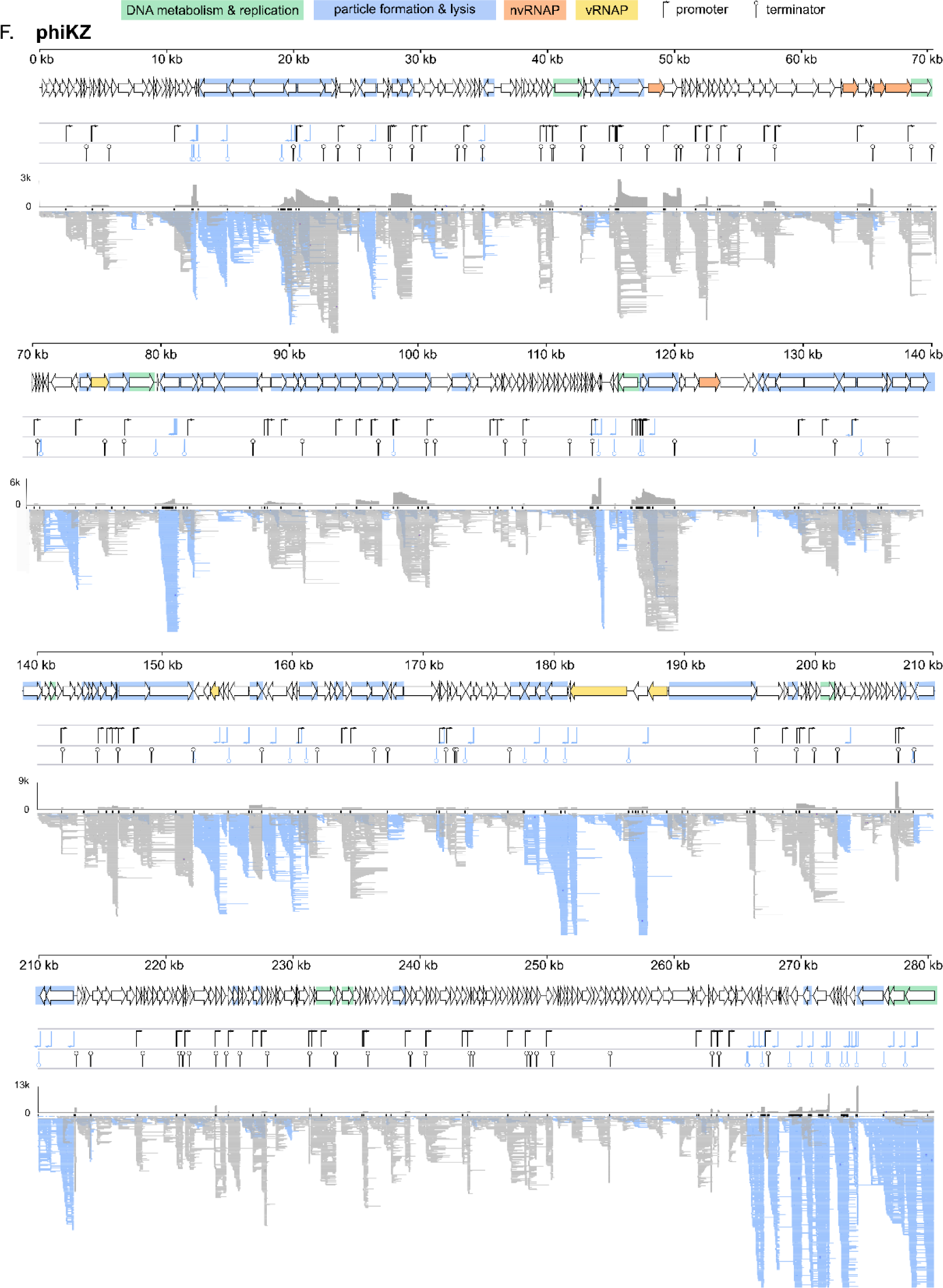
Genomic overview and transcriptional landscape of *Pseudomonas* phages 14-1 (A.), LUZ19 (B.), LUZ24 (C.), PAK_P3 (D.), YuA (E.) and phiKZ (F.), as obtained by ONT-cappable-seq. For each phage, the upper panel shows the annotated coding sequences of the phage genome. The genomic regions with genes involved in phage DNA metabolism (green), and virion morphogenesis and lysis (blue) are highlighted. The phage-encoded RNAPs of LUZ19 and phiKZ (nvRNAP) are indicated in orange. The subunits of the virion RNAP of phiKZ are depicted in yellow. Previously annotated anti-sense RNA species in PAK_P3 are indicated in dark blue underneath. The panel underneath displays the position and orientation of the promoters (arrows) and terminators (line with circle) identified in this work (Supplementary Tables S3-S5). For phage LUZ19, phage-specific promoters are indicated with a dotted arrow. The lower panel displays the ONT-cappable-seq data track, as visualised by IGV (down sampling with a 50 base window size, 50 reads per window). Reads aligning on the Watson and the Crick strand are indicated in grey and blue, respectively.

### Delineation of phage regulatory elements

#### Identification of phage TSS and promoter sequences

The phages in this work have distinct strategies to regulate their gene expression and display various degrees of dependency on the host transcriptional machinery. For example, phages 14-1, LUZ24, YuA and PAK_P3 rely fully on their bacterial host for progeny production, as they do not encode their own RNAP (Ceyssens *et al*. 2008a, 2008b, 2009; Chevallereau *et al*. 2016). As a consequence, the phages harbour strong promoter sequences and/or encode additional factors to hijack the host RNAP and alter its specificity. By contrast, phage phiKZ is equipped with two non-canonical multi-subunit RNAPs that recognise distinct promoters, supporting a phage transcriptional program that is completely independent of the host (Ceyssens *et al*. 2014). Conversely, LUZ19 encodes its own single-subunit RNAP, but relies on the host RNAP to carry out transcription of early and middle phage genes, resembling the characteristic transcriptional program of T7 (Ceyssens *et al*. 2006; Lavigne *et al*. 2013).

To uncover the dependency of the different phage genes on specific RNAPs, we mapped the TSSs of each phage individually (as specified in the M&M section), together with their associated promoter sequences. For this, phage genomes were screened for enriched positions with a local accumulation of 5’ read ends. Collectively, we identified a total of 320 TSS spread across the genomes of 14-1 (50), LUZ24 (16), LUZ19 (9), PAK_P3 (75), YuA (21), and phiKZ (149) (Supplementary Table S3). To assess whether the promoters encoded on phages that rely exclusively on the host transcription apparatus are similar to canonical σ70 promoters, the -50 to +1 regions upstream the annotated TSSs were analysed using the *Pseudomonas* promoter prediction tool SAPPHIRE.CNN (Coppens and Lavigne 2020; Coppens, Wicke and Lavigne 2022). Indeed, the majority of promoters from LUZ24 (62.5%), PAK_P3 (78.7%), and 14-1 (52%) show significant similarity to the σ70 promoter consensus of *Pseudomonas* (Figure 3A-C). Previous *in vivo* promoter trap experiments of LUZ24 revealed the presence of seven bacterial promoters (Ceyssens *et al*. 2008a). In this experiment, the phage genome was randomly sheared and the resulting fragments (200-400 bp) were cloned upstream a *lacZ* reporter gene to screen for promoter activity. Using ONT-cappable-seq, we confirm and refine the annotation of four of these reported promoters (LUZ24 P004-P006, P016), and identified six additional promoter sequences with a *P. aeruginosa* σ70 consensus motif. To make sure that no sequence motifs were overlooked, the remaining promoter elements from each phage were subjected to an additional motif search using MEME (Bailey *et al*. 2015). In the case of phage 14-1, this revealed the presence of a second motif in eleven viral promoter elements (E-value = 1.9e-007), which is characterised by a 5’-CTGGG-3’ core region located ∼5 bp from the TSS (Figure 3C).

**Figure 3:**
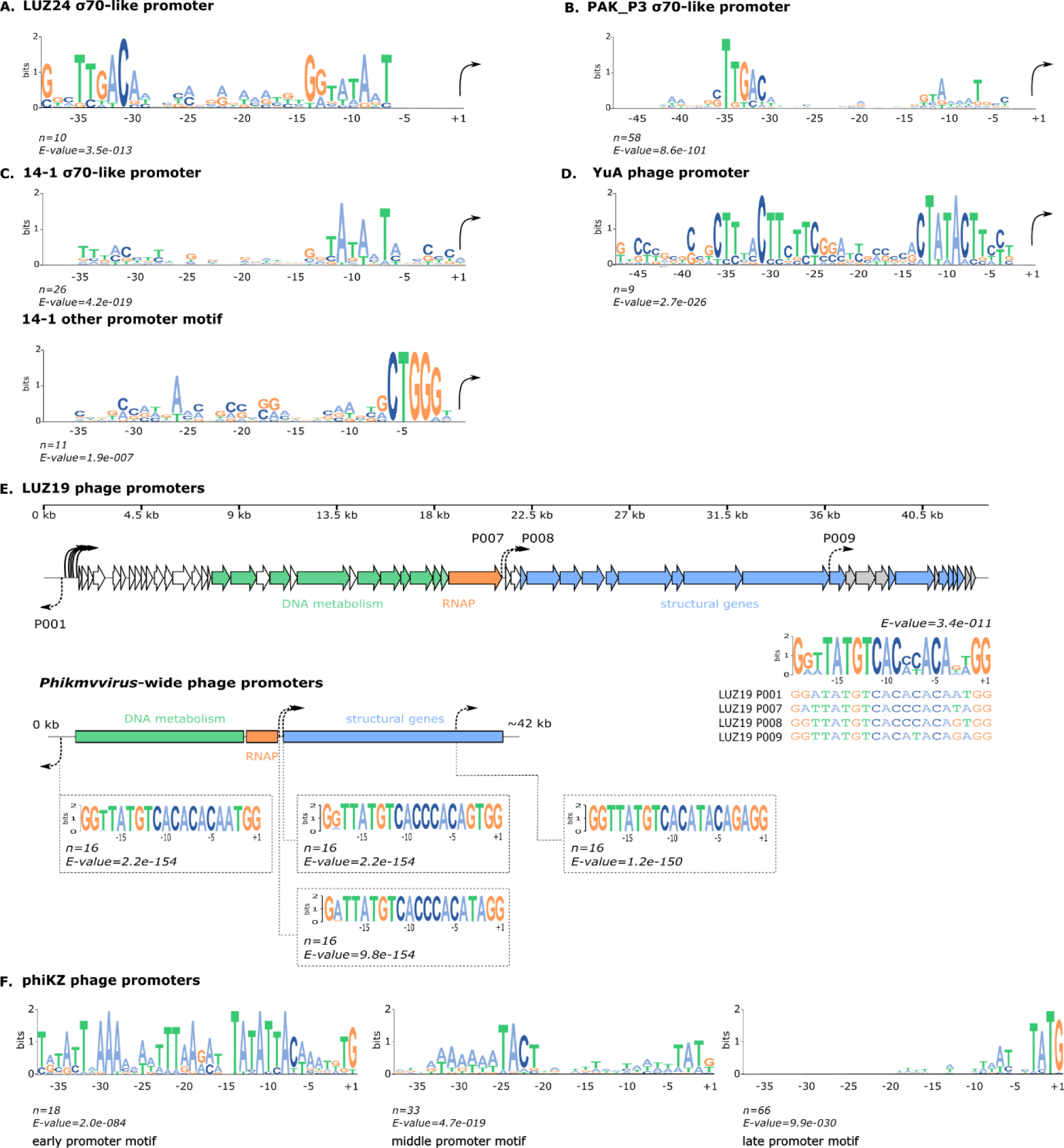
Phage promoter identification and motif analyses. Consensus sequences of σ70-like promoters encoded on the genomes of phages LUZ24 (**A.**), PAK_P3 (**B.)** and 14-1 (**C.** upper motif), as derived from 10, 58, and 26 sequences, respectively. Different promoter motifs were found for 11 and 9 TSS of phage 14-1 (**C**. lower motif) and YuA (**D.**), that do not resemble bacterial promoters. (**E.)** Phage promoters across the genome of LUZ19 (upper panel) and phiKMV-like relatives (lower panel). The phage RNAP is indicated in orange and genes involved in DNA metabolism and virion structure are depicted in green and blue, respectively. LUZ19 phage-specific promoters are indicated with dotted arrows (P001, P007-P009) and share a 20bp motif. The panel below represents the schematic genome organisation of *Phikmvviruse*s. Using the LUZ19 phage-specific promoter motif, we identified highly similar 20bp sequences in fifteen other *Phikmvvirus* species at genomic locations that match the LUZ19 promoter distribution. (**F.**) ONT-cappable-seq revealed respectively 18, 33 and 66 TSS that resemble the early, middle and late promoter motif of phage phiKZ. Motif analyses were carried out using MEME with the -50 to +1 region respective to the TSS.

YuA is dependent on both σ70 and σ54 binding factors to initiate transcription (Ceyssens *et al*. 2008b). Here, we identified two YuA promoters that can be associated with σ70-like (YuA P014) and σ54-like promoters regions (YuA P015), consistent with a previously performed fragment library promoter trap assay (Ceyssens *et al*. 2008b). In addition, nine promoter regions of YuA display a highly conserved sequence motif (MEME, E-value = 2.7e-026) (Figure 3D). Four of these phage promoters were reported previously, albeit no *in vivo* promoter activity could be observed, suggesting the need for additional (phage-encoded) factors to initiate transcription from these promoter regions (Ceyssens *et al*. 2008b). All but three of the remaining YuA promoters identified by ONT-cappable-seq were predicted to be σ70-like promoter sequences (Supplementary Table S3).

LUZ19 transcription is driven by successive actions of both the host-encoded and phage-encoded RNAP from their cognate promoter sequences. Based on ONT-cappable-seq data, we identified five strong promoters (LUZ19 P002-P006) near the leftmost region of the Watson strand of the LUZ19 genome, located upstream of gene *gp0.1,* consistent with the T7-archetypal transcriptional scheme. All but one of these promoters show significant sequence similarity to the characteristic bacterial promoter consensus, confirming the host-dependent transcription program of LUZ19 at the early infection stage (Lavigne *et al*. 2013). In addition, we identified four promoter sequences that share a highly conserved 20bp motif 5’-GgtTATGTCACacACAnnGG-3’ (lowercase letters represent lower level of conservation) (E-value = 3.4e-011), which is likely specifically recognised by the LUZ19 RNAP (Figure 3E). Two of the phage-specific LUZ19 promoters (P007-P008) are positioned directly downstream of the RNAP gene, while another one is located in the structural region (P009), in agreement with the promoter locations reported in phiKMV-like phage KP34 (Lu *et al*. 2019). However, interestingly, one of the phage-specific LUZ19 promoters (P001) is located on the opposite orientation of the coding sequences of the phage, which are generally found exclusively on the Watson strand. The antisense transcripts that originate from this promoter extend all the way towards the rightmost end of the genome, overlapping with structural genes *gp47-gp49*. One may hypothesize that this antisense transcription from phage-specific promoter P001 in the middle/late infection stage could play a role in preventing transcription elongation to early genes on the circularised phage genome.

Based on the promoter motif found in LUZ19, we assessed whether we could pinpoint the phage-specific promoters of other *Phikmvviru*s phage relatives infecting *Pseudomonas*, which have proven challenging to identify in the past (Ceyssens *et al*. 2006; Lavigne *et al*. 2013). For this, we screened the genomes of the 15 additional *Phikmvvirus* species according to the 2022 ICTV classification (Supplementary Table S6). Indeed, all phages contained almost identical 20 bp motifs at genomic positions that match the corresponding phage-specific promoters observed in LUZ19, including an antisense promoter in the early genomic region. This finding highlights a near perfect conservation of the sequence and genomic distribution of single-subunit RNAP promoters among the *Phikmvviru*s (Figure 3E). Notably, compared to T7-like phages, these viruses seemingly harbour a relatively small number of phage-specific promoters that are spread across the genome, suggesting that phiKMV-like phage transcription might be sustained by long-range processivity of RNAPs.

Finally, the transcriptional program of the giant *Pseudomonas* phage phiKZ is host-independent and relies exclusively on two distinct phage-encoded RNAPs. Upon infection, the phage co-injects its DNA along with a virion RNAP, which initiates transcription from 32 early phage promoters with a AT-rich consensus element, as revealed by primer extensions and (differential) RNA-seq profiling of PAO1 cells infected with phiKZ [23], [26]. Among the 149 phiKZ TSS and promoters identified by ONT-cappable-seq data, we recovered 16 AT-rich early promoters that were described previously, and find two additional promoters that share the phiKZ early promoter motif (Figure 3F, Supplementary Table S3). PhiKZ middle and late genes are transcribed by the non-virion-associated RNAP (nvRNAP), encoded among the phiKZ early genes. Here, we identified 33 and 66 viral TSS with upstream sequences that resemble the middle and late promoter motifs of phiKZ, respectively, of which 31 have been reported previously [23], [26] (Figure 3F, Supplementary Table S3). No significant motif could be associated with the 32 remaining upstream regions of the phiKZ TSS delineated by ONT-cappable-seq.

#### *In vivo* validation of host-specific phage promoters

Previous promoter trap studies demonstrated the presence of bacterial promoters encoded on the genomes of phage YuA and LUZ24 (Ceyssens *et al*. 2008b, 2008a). Using ONT-cappable-seq, we identified numerous promoters of 14-1, PAK_P3, and LUZ19 predicted to be recognised by the host transcriptional apparatus. As an experimental validation, the activity of a subset of promoters from 14-1, PAK_P3 and LUZ19 was evaluated *in vivo* using a fluorescence expression assay. For this, phage promoters were cloned in front of a standardised ribosome binding site (RBS) and a green fluorescent reporter protein (monomeric superfolder green fluorescent protein, msfGFP) gene, after which the genetic constructs were transformed to both *E. coli* and *P. aeruginosa* (Figure 4A). In addition, we created a construct without a promoter element (pEmpty) and a construct with the strong Pem7 constitutive promoter to serve as a negative and positive control, respectively. Interestingly, all σ70-like phage-encoded promoters (14-1 P001, P018, P036, P038; PAK_P3 P033, P034, P064, P067, P075; LUZ19 P002-P005), predicted by SAPPHIRE.CNN, are able to drive significant expression of the msfGFP reporter gene in *E. coli*, compared the negative control (Wilcoxon test, p<0.05), even though the *in vivo* activity of PAK_P3 promoters P067 and P075 is limited (Figure 4B). These data further validate the accuracy and efficacy of the SAPPHIRE.CNN tool. In addition, in *E. coli*, the majority of the host-specific phage promoters outperform the Pem7 control promoter, which is routinely used in microbial synthetic biology applications (Zobel *et al*. 2015). Unlike the phage-encoded promoters that resemble the canonical bacterial promoter consensus sequence, 14-1 promoters P011 and P032, and LUZ19 promoter P006 do not seem to drive transcription in *E. coli*, as no significant fluorescent signal could be observed. Instead, P011 and P032 contain the alternative promoter motif found in 14-1 (Figure 3C, lower motif) and no motif could be associated with LUZ19 P006. Our results suggest that additional phage-encoded transcriptional factors might be required for gene expression regulation from these promoters. Given that gp12 of 14-1 is thought to redirect the bacterial RNAP towards phage-specific promoters by interacting with the RNAP α-subunit (Bossche *et al*. 2014), P011 and P032 might rely on gp12 for transcription initiation, although this requires further investigation (unpublished results). Alternatively, other host-encoded factors, which are only expressed at specific phage-induced conditions, might also be needed for the activity of 14-1 P011, 14-1 P032 and LUZ19 P006. Finally, the majority of the promoters included in this fluorescence assay behave similarly in *P. aeruginosa* and in *E. coli*, displaying significant msfGFP expression exclusively for the σ70-like promoters (14-1 P001, P018, P036, P038; PAK_P3 P033, P064, P067; LUZ19 P005-P006) (Wilcoxon test, p<0.05), although no *in vivo* data could be gathered for LUZ19 P002-P004, PAK_P3 P034 and P075 in *P. aeruginosa* (Figure 4C). Collectively, LUZ19 P005, PAK_P3 P033 and 14-1 P001 display potent activity in both bacterial hosts.

**Figure 4:**
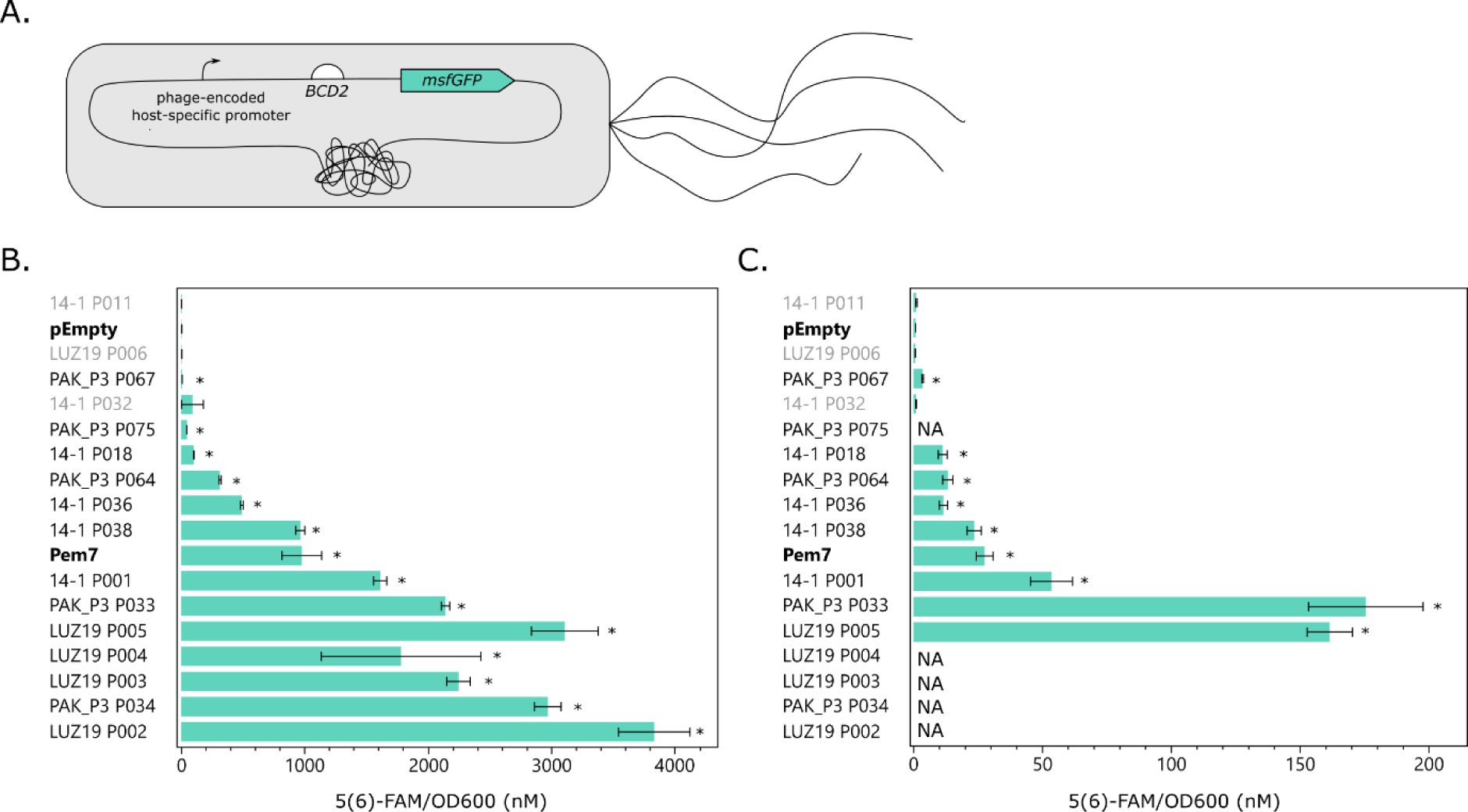
*In vivo* validation of the phage-encoded host-specific promoter activity of a subset promoters from 14-1, LUZ19 and PAK_P3. **(A.)** Schematic representation of the promoter trap construct used for *in vivo* promoter activity measurements. Phage-encoded host-specific promoters (arrow) are cloned upstream of a ribosomal binding site (BCD2) and *msfGFP* reporter gene. Bar plots showing the *in vivo* promoter activity in *E. coli*. **(B.)** and in *P. aeruginosa* (**C.**). The *in vivo* activity of the phage-encoded host-specific promoters was determined by measuring the msfGFP expression levels. The fluorescence intensity was normalised for OD and converted to an equivalent concentration of 5(6)-FAM (nM). Control msfGFP reporter constructs without promoter sequence (pEmpty) and with a constitutive promoter (Pem7) are indicated in bold. Phage promoters that could not be associated with σ70 bacterial promoters (14-1 P011, P032 and LUZ19 P005), as predicted by SAPPHIRE.CNN, are indicated in grey. Bars and error bars display the mean expression value (5(6)-FAM/OD_600_) and standard error of four biological replicates after 12h growth, respectively. Significant differences in promoter activity in comparison to the empty control construct are indicated with an ‘*’ (Wilcoxon test, p < 0.05).

#### Identification of phage transcription terminators

In addition to mapping transcription initiation sites, the ONT-cappable-seq data was leveraged to detect the 3’ boundaries of phage transcripts and annotate key transcription termination events in a global manner (Putzeys *et al*. 2022). This way, we located a total of 268 distinct transcription termination regions across the six phage genomes (14-1 (14), LUZ19 (18), LUZ24 (26), YuA (34), PAK_P3 (43), phiKZ (133)), validating 54 terminators that were predicted previously (Supplementary Table S4). Analysis of the -60 to +40 regions flanking the TTSs revealed that most of the phage terminators are prone to form energetically stable secondary structures upon transcription (Figure 5A). In general, there are two main prokaryotic transcription termination mechanisms: intrinsic termination and factor-dependent termination (Roberts 2019). Among the identified phage terminators, 84 were predicted by ARNold to be intrinsic, factor-independent transcription terminators (Naville *et al*. 2011), characterised by a canonical GC-rich hairpin structure followed by a polyuridine stretch. Interestingly, the vast majority of predicted intrinsic terminator sequences are encoded by phiKZ (>70%) and display a conserved nucleotide motif with a distinct stem-loop structure followed by a 5’-TTAT-3’ region (5’- B<u>CCYCCCC</u>HWWD<u>GGGRRGG</u>BYTTATKYYGT-3’, hairpin underlined, E-value = 1.7e-212), suggesting a shared mode of action at the RNA level (Figure 5B). Most of the termination sites either reside in intergenic regions (63%) or in a neighbouring gene sequence downsteam (28%). The remaining phage terminators (9%) are located in antisense orientation relative to the genes encoded on the phage genome. We find that the distances between the transcription termination siginals and the stop codon of the preceding gene varies extensively among all phages, although the length of most 3’ untranslated regions (3’UTRs) does not exceed 100 nt (58.2%) (Supplementary Table S4). In addition, seven overlapping bidirectional transcriptional terminator regions were discovered in the genomes of phiKZ (6) and PAK_P3 (1). Each of these bidirectional termination regions reside between a pair of convergent genes and contain two oppositely oriented TTSs located within 23-183 nt from each other.

**Figure 5:**
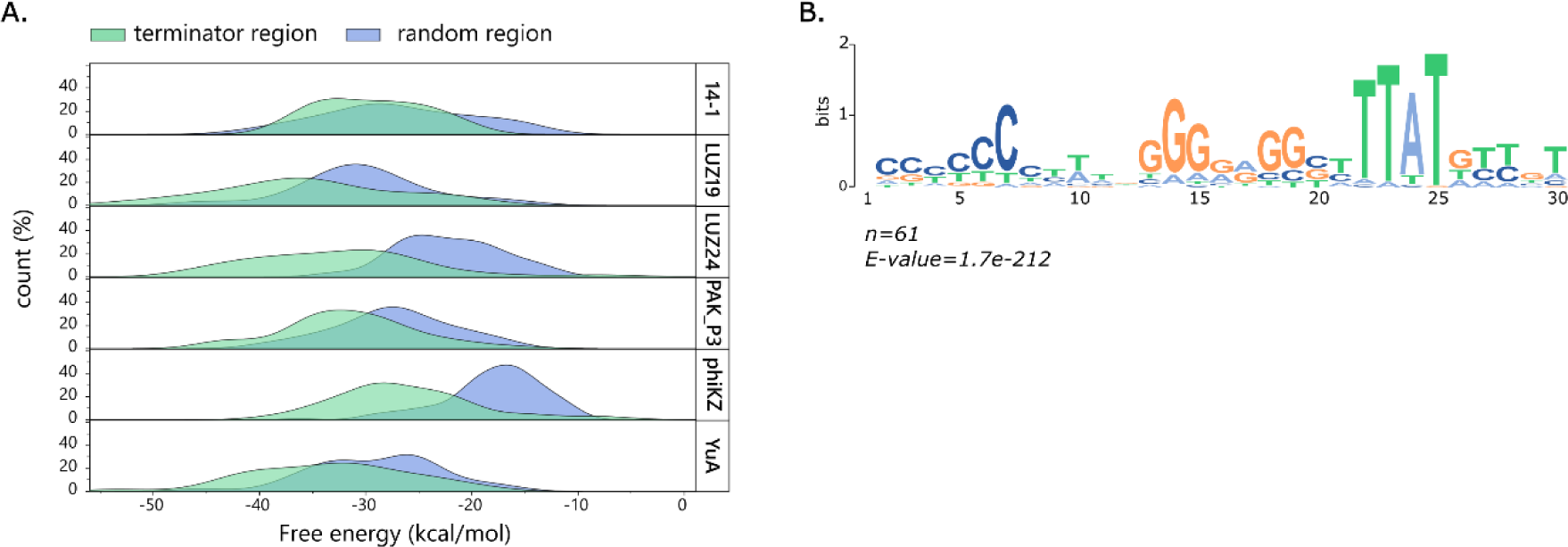
Phage terminator identification. (**A.)** Distribution of the minimum free energy (kcal/mol) of the -60 to +40 region surrounding the identified TTS of each phage (green) compared to an equal amount of randomised sequences of the same length selected across the corresponding phage genomes (blue). (**B.)** Conserved terminator motif identified for 61 terminator regions on the genome of phiKZ, which were predicted to act as intrinsic, factor-independent transcription terminators at the RNA level, as predicted by ARNold.

#### Delineation of phage transcription units

In addition to the identification of transcriptional signals that mark the start and end point of transcription, the long-read ONT-cappable-seq data can be leveraged to elucidate the transcription unit architectures of the individual phages. Transcription units (TUs) were annotated based on neighbouring TSSs and TTSs delineated in this work, resulting in a total of 520 TUs encoded in the genomes of 14-1 (55), LUZ19 (25), LUZ24 (22), PAK_P3 (130), phiKZ (247) and YuA (40) (Supplementary Table S5). Depending on the number of genes the TU encompasses, 134 (25.8%) and 257 (49.5%) TUs were classified as monocistronic or polycistronic, respectively. The remaining TUs (24.7%) did not fully span any genomic feature annotated on the phage genomes. On average, the TUs of LUZ19 and 14-1 are 0.8-1.2 kb and cover between one and two genes (Supplementary Figure S2). The TUs of YuA and LUZ24 are seemingly shorter (average TU length 483-722 bp), and do not encode more than two or three genes, respectively. By contrast, the TUs of phiKZ and PAK_P3 have an average length of 1.6-2 kb and can comprise up to 10-16 annotated genes.

In general, genes that are co-transcribed in the same TU are likely to be functionally related. This also seems to be the case for the *Pseudomonas* phages in this study. For example, whereas 14-1 TU015, LUZ19 TU025, phiKZ TU137, and PAK_P3 TU12 contain genes that are involved in virion structure and morphogenesis, the genes encompassed by LUZ24 TU016 and PAK_P3 TU040 play key roles in viral genome replication. In addition, the gene content of the phage TUs shows significant overlap, sharing at least one or more genes between individual TUs. In the case of 14-1 (52%), phiKZ (67%) and PAK_P3 (77%), the majority of genes in the defined TUs are transcribed in at least two different transcriptional contexts, which represents the number of unique gene combinations encoded by the TUs (Yan *et al*. 2018). Similar observations were made for *P. aeruginosa* phages LUZ7 and LUZ100 (Putzeys *et al*. 2022, 2023). Taken together, the widespread identification of overlapping TUs in diverse *P. aeruginosa* phages suggests that the alternative use of TU variants might be a common regulatory strategy to adjust gene expression of individual genes during phage infection.

The genomes of phages, including the *Pseudomonas* phages in this study, are endowed with a multitude of genes that lack functional annotation, often referred to as the ‘viral dark matter’ (Hatfull 2008, 2015). As the functional elucidation of these hypothetical proteins is a time-consuming and challenging endeavour (Roucourt and Lavigne 2009; Wan *et al*. 2021), ONT-cappable-seq data could be a first step to infer their role during phage infection. Knowledge of the transcriptional context of TUs encompassing such hypothetical genes could be leveraged to obtain clues on their function. For example, the longest version of PAK_P3 TU046 spans genes *gp65-gp69*, which are involved in thymidylate synthesis (*gp65*, *gp66*) or code for a putative ribonucleotide-diphosphate reductase (*gp67*, *gp69*) (Chevallereau *et al*. 2016). By contrast, PAK_P3 *gp68* lacks any amino acid sequence similarity to other proteins in the databases, and hence, its function remains elusive. However, the transcriptional context of PAK_P3 *gp68*, as derived from ONT-cappable-seq data, suggests that it is likely involved in nucleotide biosynthesis as well. In addition, the nuclear shell protein of phiKZ, *PHIKZ054* (Chaikeeratisak, Birkholz and Pogliano 2021), appears to be transcribed in many different TUs and transcriptional contexts (Supplementary Table S5), including phiKZ TU038. In phiKZ TU038, the phage shell protein is co-transcribed with a gene of unknown function, *PHIKZ053,* hinting towards a related role in the formation of the phage nucleus-like structure upon infection.

In contrast to genes encoded on polycistronic transcripts, phage morons are independently acquired autonomous genetic modules that generally contain their own promoter and terminator elements (Juhala *et al*. 2000). These nonessential moron genes are mainly associated with prophages and confer a fitness advantage for the bacterial host, including *P. aeruginosa* (Tsao *et al*. 2018; Taylor *et al*. 2019). Given that roughly 25% of the phage TUs identified by ONT-cappable-seq encompass a single gene flanked by a TSS and TTS, investigation of these TUs could point to similar portable transcription units in virulent phages. For example, phiKZ TU118, TU129, and TU123 respectively harbour hypothetical protein-coding genes *PHIKZ153, PHIKZ_p52*, and *PHIKZ169*, which are all embedded within another gene cluster encoded on the opposite strand, suggesting that they were acquired individually. In addition, phage 14-1 TU021 solely contains gene of unknown function *gp48* and has a slightly different G+C base composition (51%) compared to the 14-1 genome (56% G+C) and up- and downstream flanking genes with 60% and 57% G+C, respectively. Although the origin and function of the phage genes remains to be elucidated, these examples illustrate the potential of TU analysis to pinpoint transcriptionally independent loci, which could ultimately unveil moron-like sequences in (pro)phage genomes.

### ONT-cappable-seq guides the discovery of putative regulatory RNAs

Interestingly, TU analysis revealed that ∼25% of the phage TUs do not contain any previously predicted genomic features. Among these, a considerable number of TUs are the apparent result of premature transcription termination events, as their cognate TTSs were found immediately downstream of promoter elements, which could be reflective of regulatory events (Adams *et al*. 2021). Collectively, we identified 42 phage TTS that reside within 5’UTR regions, as defined by ONT-cappable-seq. Based on the global phage transcriptional profiles, 34 of these terminators (81%) appear to have two strict modes of transcriptional read-through, which either result in a long transcripts that span one or multiple genes, or a predominant short transcript version of less than 250 nt with the same 5’ extremity. For example, phiKZ T102 is located within the 5’UTR of *PHIKZ221*, downstream of the TSS associated with phiKZ P112. While some transcripts that start at this position fully cover genes *PHIKZ221* and *PHIKZ222*, the majority (>90%) is aborted prematurely by T102, resulting in high levels of a short RNA species of ∼71 nt (Figure 6A). Similarly, phiKZ TU237 gives rise to a small intergenic transcript of 141 nt in length, delineated at its 3’ end by terminator T131, upstream of gene *PHIKZ302* (Figure 6B). In addition, the abundance and length of these two short RNA species was confirmed by northern blot probing, which also captured the full-length transcript version, consistent with the associated ONT-cappable-seq cDNA reads (Figure 6). Given their abundance and size, these short structured fragments, which appear tightly controlled by conditional termination, might have a regulatory function (Dar *et al*. 2016; Adams *et al*. 2021; Felden and Augagneur 2021).

**Figure 6:**
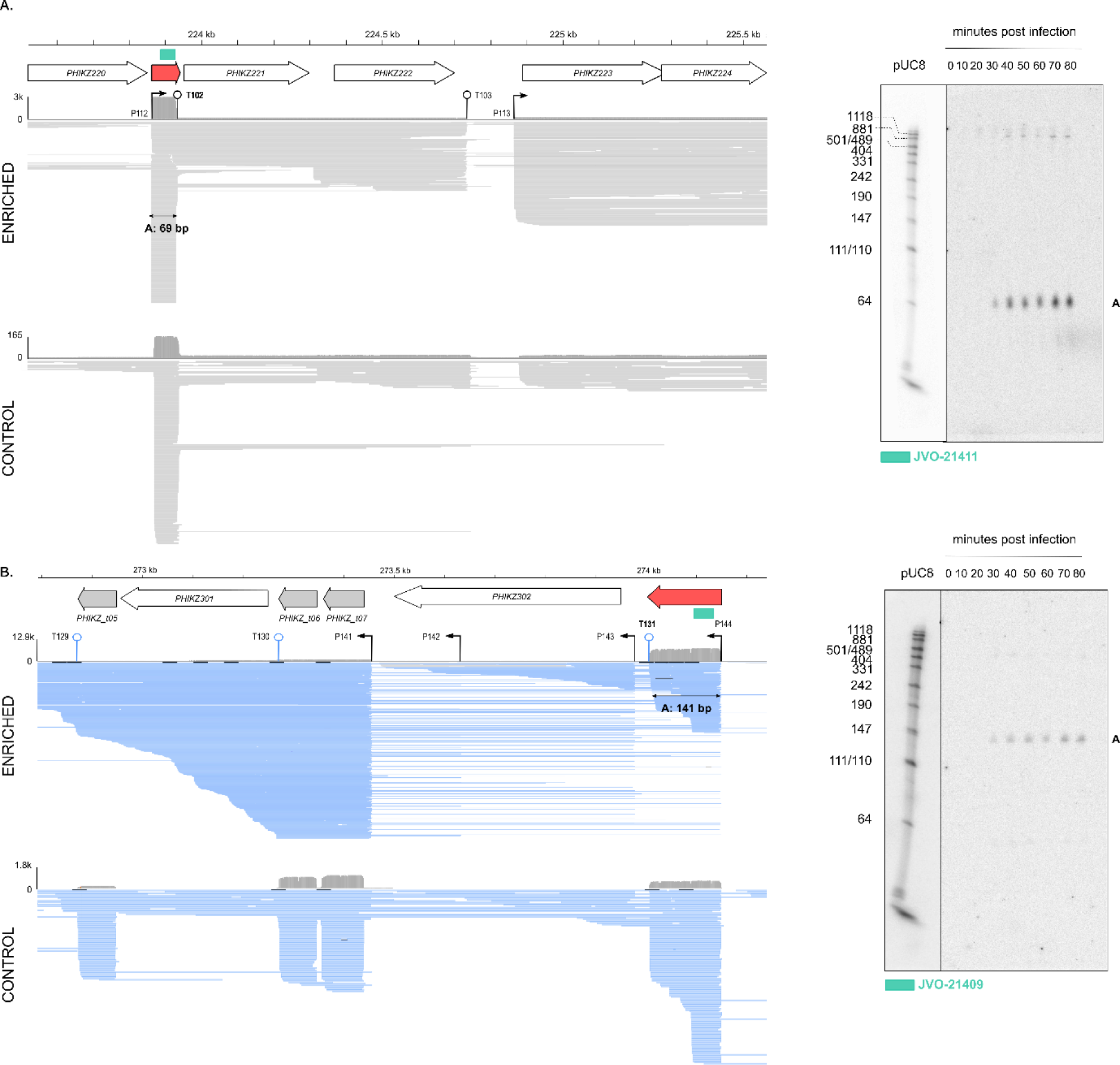
Example of transcription patterns in 5’UTR regions in phiKZ that reveal putative small non-coding RNA candidates. IGV data tracks of ONT-cappable-seq enriched and control datasets indicating the abundancy of short RNA species in the 5’UTR region of *PHIKZ221* (**A.**) and *PHIKZ302 (*(**B.**), suggesting that they play a regulatory role. The production of the small transcripts is seemingly controlled by premature transcription termination. Reads mapped on the Watson and Crick strand are indicated in grey and blue, respectively. The alignment view in panel B was down sampled in IGV for visualisation (window size = 25, number of reads per window = 50). Genes, tRNAs and ncRNA candidates are displayed as white arrows, grey arrows, and red filled arrows, respectively. The position and orientation of the promoters (arrows) and terminators (line with circle) are indicated. Northern blot probing (probe position in genome indicated with green rectangle) confirms the length and abundance of the ncRNA candidates, in agreement with the transcriptional profile.

Among the small 5’UTR-residing RNA species discovered by ONT-cappable-seq, two were previously reported as putative non-coding RNAs (ncRNA), and serve as a first validation of the ONT-cappable-seq approach for ncRNA discovery. Indeed, grad-seq data of *P. aeruginosa* cells infected with phiKZ revealed two phage-encoded ncRNA candidates associated with the 5’UTR and 3’UTR region of *p40* and *PHIKZ298*, respecitively (Gerovac *et al*. 2021). These putative ncRNAs were predicted to be 89 nt (3’UTR_PHIKZ298) and 196 nt (5’UTR_gp40) in length, respectively. However, closer inspection of the ONT-cappable-seq reads associated with 3’UTR_PHIKZ298, together with northern blot probing, justifies re-annotation of the transcript boundaries (270595-270485(-)). In addition, comparison of the 3’UTR_PHIKZ298 transcriptional profiles in the enriched and control datasets, together with the identified TSS associated with this transcript, suggests that it is likely generated by 5’UTR processing of the *PHIKZ297* mRNA as well (Supplementary Figure S3). After manual screening, the small RNA species for potential open reading frames (≥ 10 AA) preceded by a canonical Shine-Dalgarno sequence, ONT-cappable-seq data hints towards the presence of 29 5’UTR-derived ncRNA candidates in the genomes of 14-1 (1), LUZ19 (2), LUZ24 (2), YuA (3), PAK_P3 (4), and phiKZ (17) (Supplementary Table S7), significantly expanding the number of putative regulatory ncRNAs produced by virulent phages (Bloch *et al*. 2021). In addition, at the sequence level, these ncRNA candidates appear to be conserved across *P. aeruginosa* phage relatives within the same genus. For example, the sequence of the short RNA species identified upstream the PAK_P3 DNA primase/helicase gene (*gp49*) (fragment 31147-31308(+)), is also found upstream of the corresponding gene in other *Nankokuvirus* members, including KPP10 (NC_015272.2) and P3_CHA (KC862296.1) (BLASTN >90% identity, E-value < 2e-65) (Supplementary Figure S4). Similarly, a ncRNA candidate of YuA (39396-39519(+)) is located in the 5’UTR of the phage major head protein gene (*gp56*) and shows sequence conservation to the upstream region of the equivalent gene in *Yuavirus* relatives, such as M6 and LKO4 (BLASTN >97%, E-value < 5e-57). In conclusion, our results indicate that this technique provides the means to identify novel ncRNA candidates at a genome-wide scale, as well as offering clues towards their biogenesis.

Previous standard RNA-sequencing experiments of PAK_P3 predicted two putative ncRNAs, named ncRNA1 (89 nt) and ncRNA2 (132 nt), which were highly expressed during the late infection stages (Chevallereau *et al*. 2016). Using ONT-cappable-seq, we recovered both RNA species and refined their transcriptional boundaries with single nucleotide precision (ncRNA1: 85,248-85,458(+); ncRNA2: 86,307-86,426(+)). In addition, both RNA species lack an associated TSS, suggesting they are derived from extensive processing of the 3’UTR of the upstream gene. This also becomes apparent when comparing the PAK_P3 ONT-cappable-seq data tracks of the enriched and control datasets. These indicate that the small RNA species are predominantly present in the control dataset, which mainly contains processed transcripts (Supplementary Figures S5-S6). These results are in agreement with northern blot probing of ncRNA1 and ncRNA2, displaying several processed intermediates and a highly abundant RNA species of ∼210 nt and ∼120 nt in length, respectively, of which the signal becomes stronger over the course of infection (Supplementary Figures S5-S6). In addition to pointing out putative 5’UTR- and 3’UTR-derived ncRNAs, ONT-cappable-seq reveals twenty phage-encoded antisense RNA species of varying lengths with a defined TSS (LUZ19 (2), LUZ24 (2), YuA (4), PAK_P3 (8), phiKZ (4) (Supplementary Table S5). Among these, seven can be associated with anti-sense transcripts discovered in PAK_P3 by previous RNA-seq analysis (Chevallereau *et al*. 2016). Collectively, these results illustrate the potential of ONT-cappable-seq to finetune transcript annotation and discern novel RNA species that were previously overlooked by classic RNA-seq experiments. We envision that further functional studies on this extensive set of antisense transcripts and putative ncRNAs can offer valuable insights in their regulatory role and importance during phage infection (Bloch *et al*. 2021).

### Full-length transcriptional profiling enables straightforward detection of splicing activity

Group I introns, intervening sequences capable of self-splicing, are widespread in the genomes of bacteria and phages, including *Pseudomonas* phage LUZ24 (Ceyssens *et al*. 2008a; Lavigne and Vandersteegen 2013; Hausner, Hafez and Edgell 2014). The LUZ24 intron interrupts the coding sequence of the phage DNA polymerase, and restores the reading frame upon excision from the corresponding primary transcripts. The presence of bacteriophage group I introns is generally predicted from genomic surveys, and can be subsequently confirmed after PCR amplification of the cDNA. Given that ONT-cappable-seq allows full-length cDNA sequencing, we evaluated whether this intron can also be detected in our global transcriptome data of LUZ24. Indeed, visual inspection of the phage transcriptome data showed a considerable number of transcripts that lack the 669-bp fragment (19,143-19,811) which matches the Group I intron embedded in the DNA polymerase of LUZ24, confirming its *in vivo* splicing activity (Supplementary Figure S7). In addition, the relative number of spliced cDNA read alignments in this region is significantly lower in the sample enriched for primary transcripts (0.03%) compared to the control sample which was predominantly composed of processed transcripts (0.9%), as expected. Previous studies demonstrated that environmental cues can impact group I intron splicing efficiency (Sandegren and Sjöberg 2007; Belfort 2017). The observed occurrence of both spliced and unspliced cDNA is consistent with the experimental design in which multiple timepoints have been pooled, also hinting at condition-dependent splicing events. In addition, the full-length 669-bp fragment is relatively more abundant in the enriched dataset, representing 46.5% of the mapped reads in this region, compared to the control sample (1%). This suggests that its 5’ end holds a triphosphate group, which is enriched for during library preparation. Indeed, group I intron splicing is initiated through hydrophilic attack of the 5’ splice site by a free guanosine-5’-triphosphate, the latter of which is subsequently linked to the 5’ end of the intron (Cech 1990; Hausner, Hafez and Edgell 2014). Based on these data, it should be noted that the 5’ and 3’ ends associated with this fragment most likely do not represent genuine TSS (LUZ24 P011) and TTS (LUZ4 T016), respectively (highlighted in red in Supplementary Tables S3-S5). Previous genomic analysis of phiKZ also predicted the presence of Group I intron-containing endonucleases in the genes encoding gp56, gp72 and gp179 (Mesyanzhinov *et al*. 2002). However, careful surveillance of the phiKZ full-length transcriptional landscape did not provide evidence to support this hypothesized self-splicing activity under the growth conditions tested here.

In contrast, ONT-cappable-seq data revealed a considerable number of spliced cDNA reads in another LUZ24 gene, *gp2*. LUZ24 gp2 is a conserved hypothetical protein that was shown to be non-inhibitory in terms of host growth and biofilm formation (Wagemans *et al*. 2015). More specifically, a 34-bp fragment (1,659-1,692) appears to be removed near the 3’ end of 3.9% and 4.3% of the cDNA reads aligned to *gp2* in the enriched and control transcriptomic samples of LUZ24, respectively (Figure 7). Interestingly, we identified four ‘CAAGG’ repeats, paired two by two, at the boundaries of the spliced sequence. The pairs reside 16-bp from each other and the repeats in each pair lie seven base pairs apart. Moreover, the five nucleotides immediately downstream the intron sequence (AACTG) are identical to the 5’ sequence of the excised fragment. Manual inspection of individual reads indicated high mapping accuracy, which supports the hypothesis that the spliced cDNA reads are biologically relevant, and not the result of alignment errors due to error-prone nanopore reads. In addition, PCR of first-strand cDNA derived from *P. aeruginosa* cells infected with LUZ24, followed by gel electrophoresis, reveals the presence of a smaller, less intense fragment, suggesting that a short, intervening sequence is removed from a portion of the transcripts throughout infection (Supplementary Figure S1A). Although the molecular underpinnings of this short intervening sequence remain to be uncovered, these findings demonstrate that ONT-cappable-seq can be a valuable strategy to confirm, identify and elucidate splicing events in bacteriophages, and could readily be extended to their bacterial hosts.

**Figure 7:**
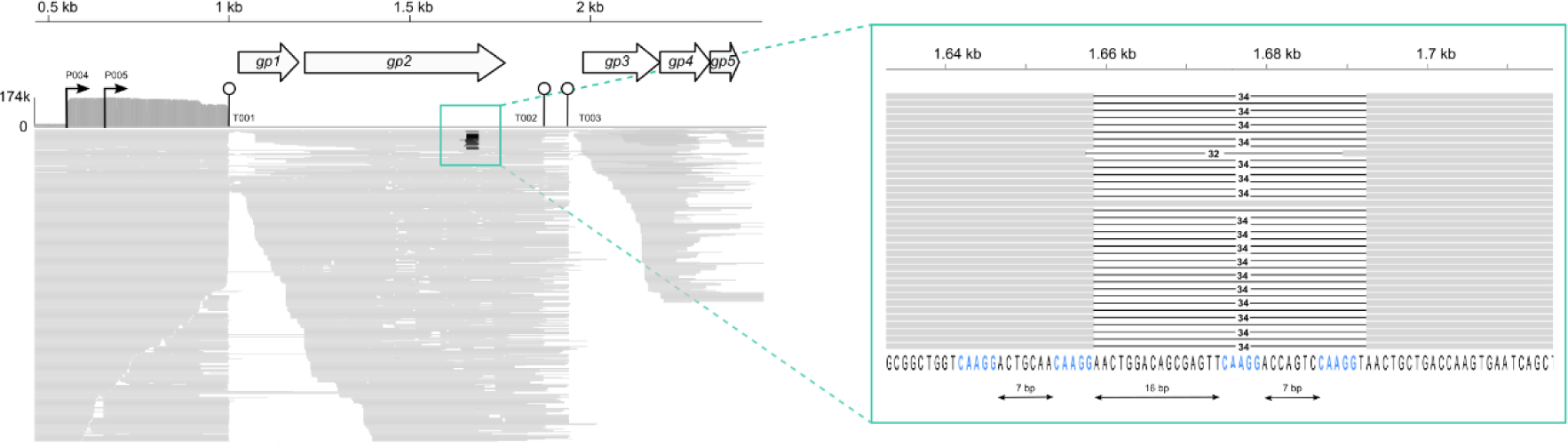
ONT-cappable-seq data suggests splicing activity in LUZ24 *gp2* transcripts. IGV visual representation of ONT-cappable-seq datatrack of a region of the LUZ24 genome spanning *gp1-gp5*. Read alignments show splicing of a 34-bp fragment in cDNA reads that map to *gp2*. Closer inspection of the boundaries of the putatively spliced fragment reveals four regularly interspaced ‘CAAGG’ repeats, paired two by two. The position and orientation of the promoters (arrows) and terminators (line with circle) are indicated. Reads mapping on the Watson and Crick strand are indicated in grey and blue, respectively.

## Conclusions & perspectives

In the last decade, classic RNA sequencing has been the main method to profile the transcriptional landscape of phage-infected bacteria, providing insights in temporal gene expression levels and host responses. However, these short-read methods are generally not suited to gain in-depth knowledge of the transcriptional regulatory mechanisms involved in phage infection, as they lack the ability to differentiate between primary and processed transcripts and information on transcript continuity is lost. By contrast, differential RNA-seq (dRNA-seq) performs differential treatment of the RNA sample with 5’P-dependent terminator exonuclease (TEX) to enrich primary transcripts prior to short-read sequencing, and is considered the golden standard for global prokaryotic TSS mapping (Sharma *et al*. 2010; Sharma and Vogel 2014). Alternatively, Cappable-seq is based on targeted enrichment for the 5’-PPP end of primary transcripts, followed by Illumina sequencing (Ettwiller *et al*. 2016). Here, we applied ONT-cappable-seq to profile the full-length primary transcriptomes of a diverse set of lytic phages infecting *P. aeruginosa*. Using this method, we pinpointed key regulatory elements that mark the start and end of transcription and delineated transcription units across the genomes of LUZ19, LUZ24, YuA, 14-1, PAK_P3 and phiKZ, significantly refining their transcriptional maps and highlighting the extensive diversity and complexity of transcriptional strategies across *Pseudomonas* phages. Compared to dRNA-seq, which has been applied to map early TSSs of phiKZ at single-nucleotide resolution (Wicke *et al*. 2021), we demonstrated extensive overlap with the phiKZ TSSs defined by ONT-cappable-seq, with a positional difference limited to 2 nt. Furthermore, we find that individual phage TUs are highly interconnected, suggesting that alternative TU usage might be a widespread regulatory strategy in *Pseudomonas* phages to balance and finetune gene expression levels in their densely coded genomes in response to different stimuli (Lee *et al*. 2019; Putzeys *et al*. 2022, 2023). In bacteria, 5’UTR conditional premature transcription is an important regulatory mechanism to modulate gene expression levels under different environmental stressors and conditions (Merino and Yanofsky 2005; Adams *et al*. 2021; Konikkat *et al*. 2021). ONT-cappable-seq data of the different phages also revealed numerous 5’UTR premature transcription termination events, pointing to 29 potential phage-encoded 5’UTR-derived ncRNA candidates, two of which were previously detected by phiKZ dRNA-seq and grad-seq experiments (Gerovac *et al*. 2021; Wicke *et al*. 2021). To our knowledge, this is the first study to source putative phage-encoded ncRNA at this scale. Finally, we find that ONT-cappable-seq offers a straightforward approach to study splicing events, as illustrated by the confirmed 669-bp Group I intron and the observed splicing activity in *gp2* in the LUZ24 genome. Collectively, this work highlights the wealth of information that can be gained from global ONT-cappable-seq experiments, uncovering transcript boundaries and transcriptome architectures, as well as introns and ncRNA candidates, all of which have remained largely understudied in phages to date.

However, while ONT-cappable-seq is a powerful standalone method to obtain a birds-eye view of phage transcriptomes and their regulatory features; classic, temporally-resolved RNA-seq experiments remain imperative for quantitative evaluations of gene expression in phage-infected cells. Notably, temporal gene expression information combined with global ONT-cappable-seq data could help infer preferential TSS and TTS usage of individual genes in specific infection stages, as well as reveal discrepancies that might point to interesting RNA processing events. Indeed, these methods collectively provide the means to globally examine gene expression and transcript boundaries, yet information on transcript modification, structure and interaction cannot be inferred and requires the adoption of other specialized transcriptomic methods (Hör, Gorski and Vogel 2018). We envision that the routine application of state-of-the-art transcriptomics approaches in the phage field, including but not limited to ONT-cappable-seq, will shed light on the RNA biology of non-model phages, to ultimately help bridge the existing knowledge gaps in the complex molecular mechanisms at play during phage infection.

## Data availability

The resolved genome of *P. aeruginosa* strain Li010 was deposited in NCBI GenBank (accession number CP124600). Raw and processed RNA sequencing files were made available under GEO accession number GSE231702. Any additional information is accessible from the authors upon request.

