## Supplementary Figures S1-S7, Supplementary Tables S1-S7 for "Refining the transcriptional landscapes for distinct clades of virulent phages infecting *pseudomonas aeruginosa*"

### Supplementary data

#### Supplementary Figures

A

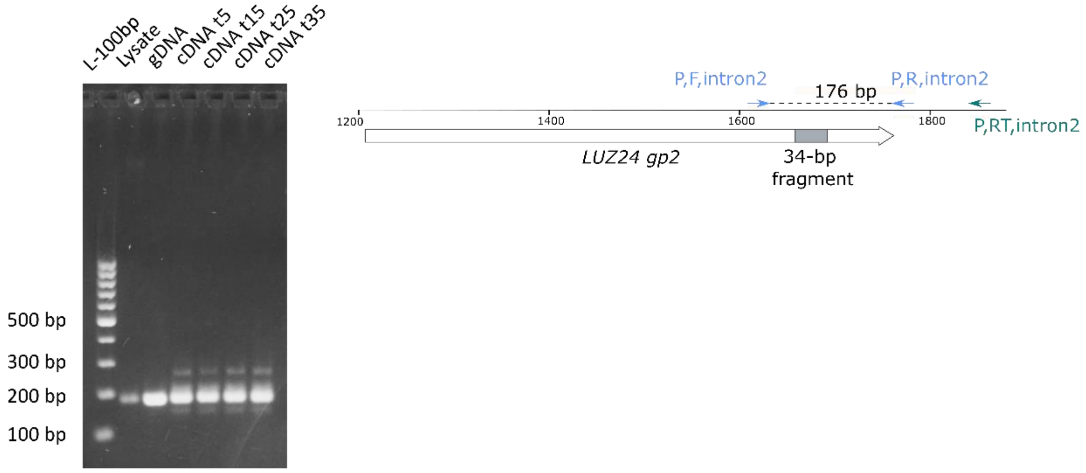


B.

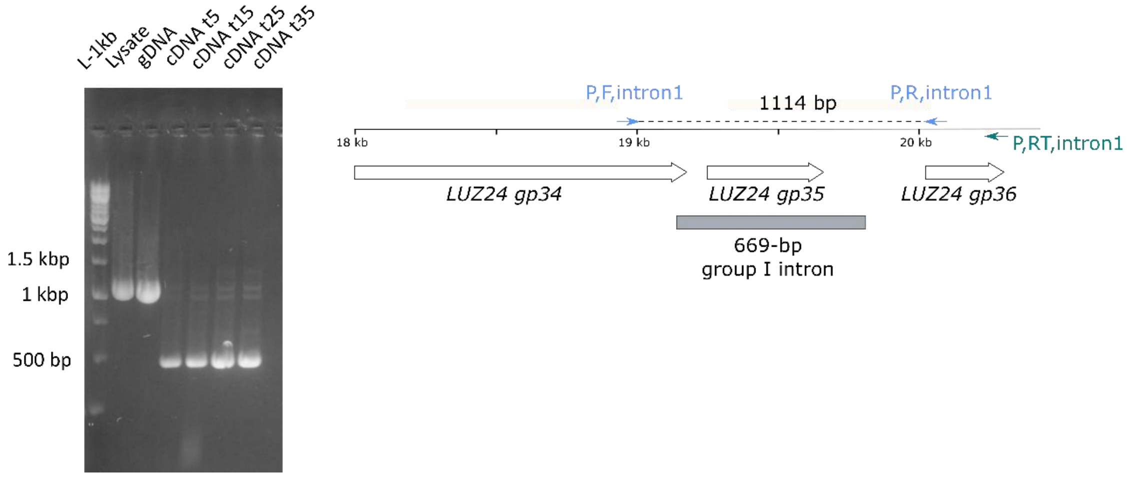


**Supplementary Figure S1: Demonstration of LUZ24 splicing activity of a 34-bp fragment in *gp2* transcripts (A) and the previously identified 669-bp group I intron (B).** Visualization of PCR products on a 1.5% agarose gel. PCR was performed on phage lysate and gDNA of LUZ24 using primer pairs that span the intron region of interest. Lanes labeled cDNA t5, t15, t25 and t35 represent PCR products derived from a cDNA template, which was generated on RNA samples from different infection stages of LUZ24 (5min, 15min, 25min, 35min). The lanes of the cDNA-derived PCR products show a band below the band observed in the lanes of the lysate and genomic DNA. Primers used for cDNA conversion and PCR amplification are indicated in green and blue, respectively.

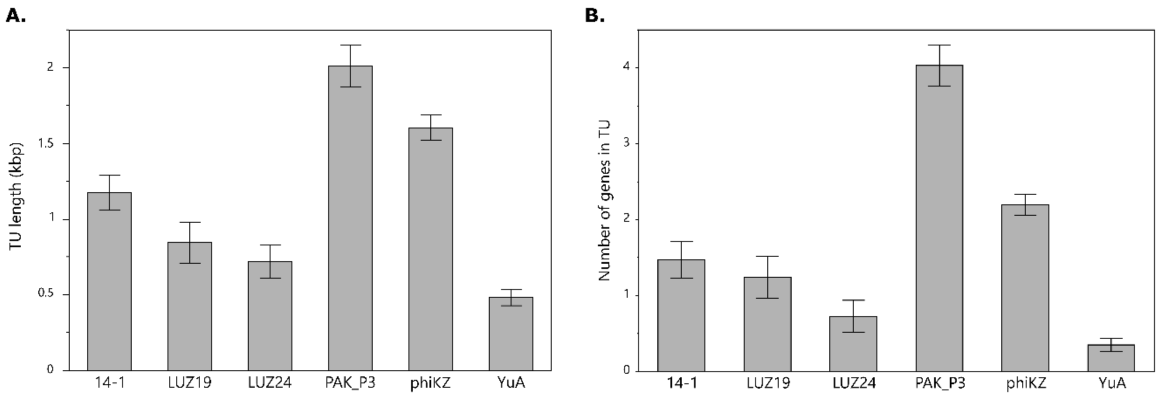


**Supplementary Figure S2: Characteristics of the phage transcription units**. Bar plots showing for each phage, the average length of the transcription units defined by ONT-cappable-seq (A)., and the average number of genes encoded within (B.). Error bars represent the standard error.

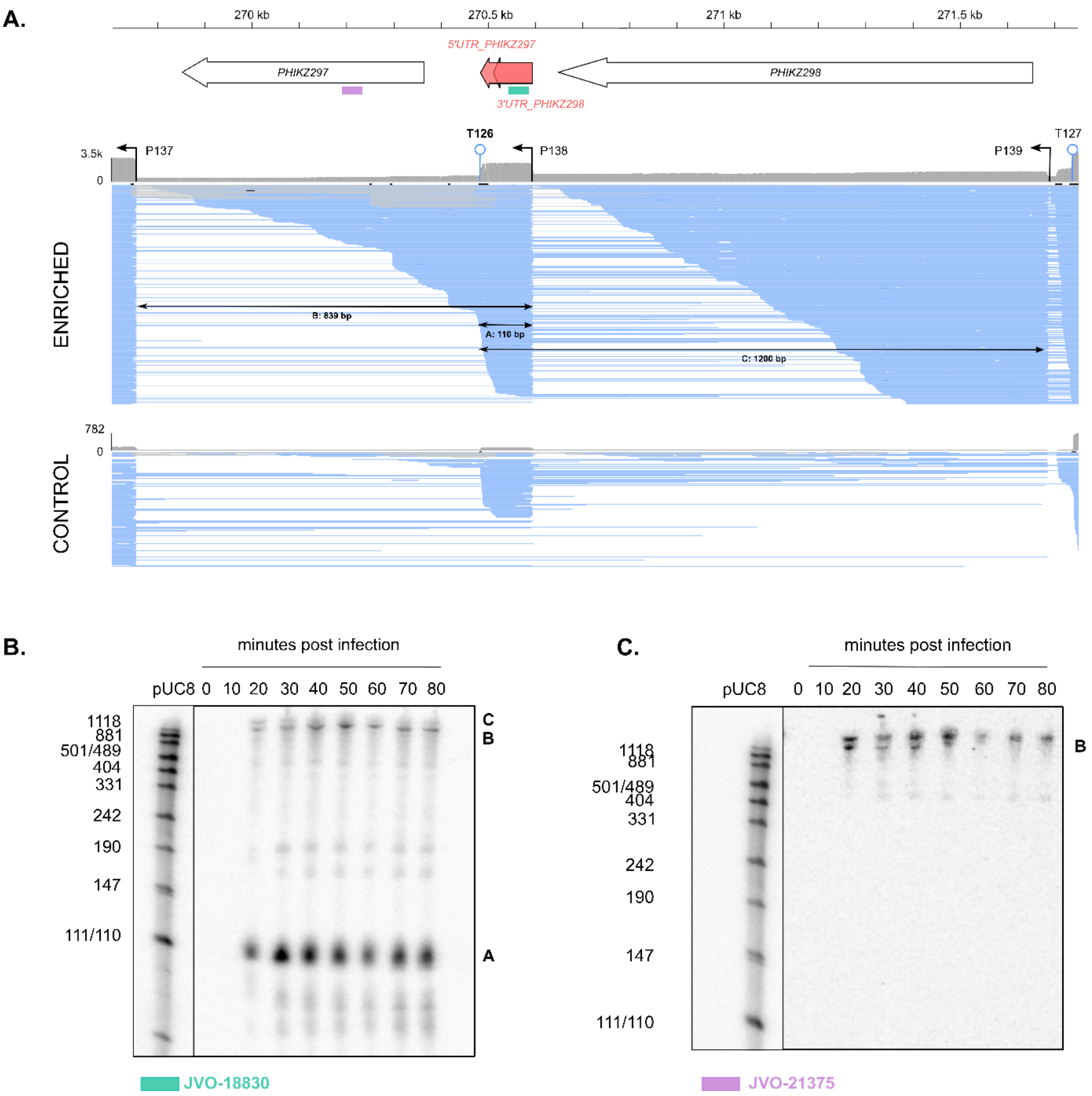


**Supplementary Figure S3: ONT-cappable-seq refines the boundaries of the phiKZ 3'UTR_PHIKZ298 ncRNA candidate and demonstrates that it can also be derived from the 5'UTR of the *PHIKZ297* mRNA. (A.)** IGV ONT-cappable-seq tracks of the enriched and the control phiKZ samples of the intergenic region between genes *PHIKZ297* and *PHIKZ298.* Previous Grad-seq experiments revealed the presence of the 3'UTR-derived ncRNA candidate, 3'UTR_PHIKZ298. Based on the TSS and TTS identified in this work, we re-annotated the transcript boundaries and find that it can be derived from the 5'UTR of PHIKZ297 as well (5'UTR_PHIKZ297). Genes and ncRNA candidates (old and new annotation) are displayed with black outlined arrows and red filled arrows, respectively. The position and orientation of the promoters (arrows) and terminators (line with circle) are indicated. Reads and terminators mapping on the Watson and Crick strand are indicated in grey and blue, respectively. The alignment view was downsampled in IGV for visualisation (window size = 5, number of reads per window = 25). **(B.)** Northern blot probing of the ncRNA candidate (green rectangle in part A) confirms abundance and size of the transcript (A: 110 nt) and shows longer intermediates (B: 830 nt & C: 1100 nt) that can be found in the transcriptional landscape. (**C.**) Northern blot probing of *PHIKZ297* (purple rectangle) suggests that the ncRNA candidate can originate from processing of the 839 nt transcript of PHIKZ297 (fragment B).

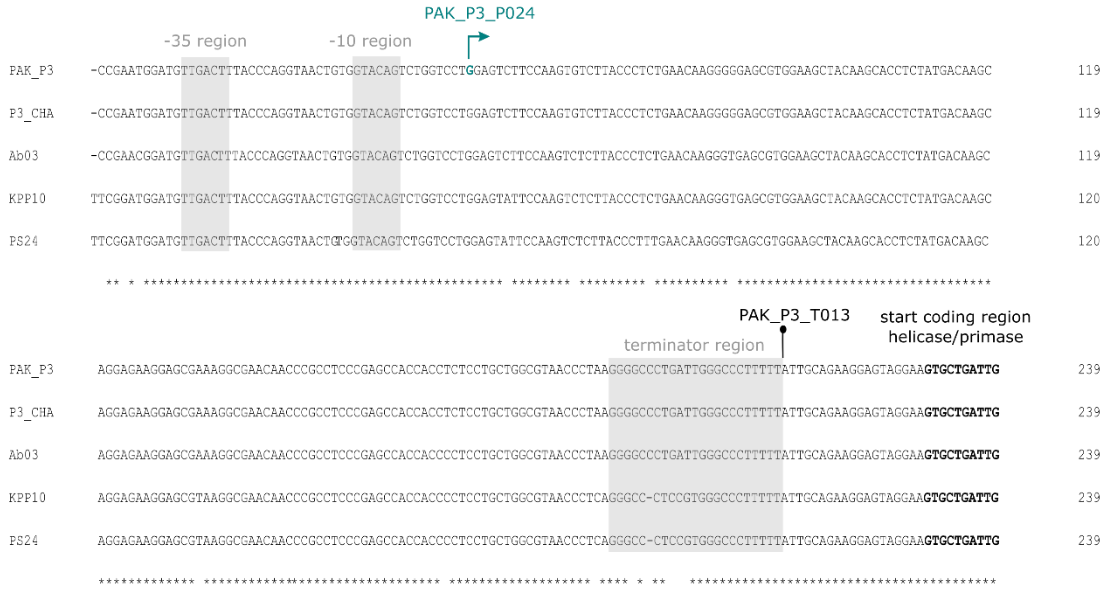


**Supplementary Figure S4: Multiple sequence alignment (Clustalω) of the 5'UTR region of the PAK_P3 helicase/primase gene (*gp49*) with the corresponding regions of *Nankokuvirus* phage relatives shows strong sequence conservation.** PAK_P3 transcription start- and termination sites defined by ONT-cappable-seq, together with their associated promoter and terminator regions are projected on the sequences of the other phages. The start of the coding region of the annotated helicase/primase gene in each phage is indicated in bold.

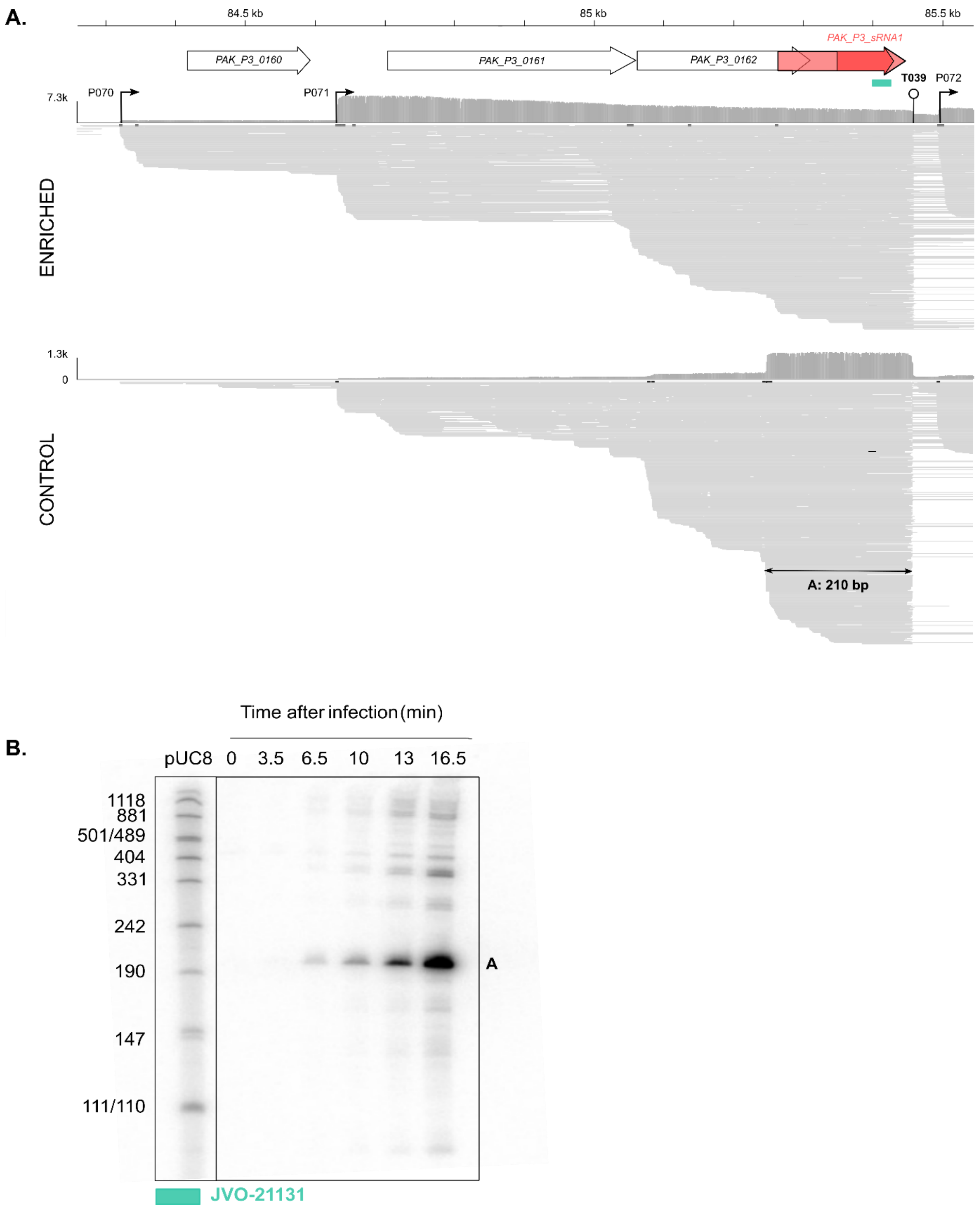


**Supplementary Figure S5: ONT-cappable-seq (A) and northern blot probing (B) of PAK_P3 ncRNA candidate sRNA1 warrants reannotation of the transcript boundaries and indicates it is derived from the 3'UTR of *PAK_P30162*. (A.)** IGV ONT-cappable-seq tracks of the enriched and the control PAK_P3 samples. Previous RNA-seq experiments revealed the presence of an 89-nt ncRNA candidate, sRNA1. Based on the TSS and TTS identified in this work, we re-annotated the transcript boundaries (now 210-nt long) and find that it is likely derived from the 3'UTR of *PAK_P3_162*. Genes and ncRNA candidates (old and new annotation) are displayed with black outlined arrows and red filled arrows, respectively. The position and orientation of the promoters (arrows) and terminators (line with circle) are indicated. Reads and terminators mapping on the Watson and Crick strand are indicated in grey and blue, respectively. The alignment view was downsampled in IGV for visualisation (window size = 5, number of reads per window = 25). (**B.**) Northern blot probing of the ncRNA candidate (green rectangle in part A) confirms abundance and size of the transcript (A: 210 nt) in the late infection stage and shows extensive processing, in agreement with the transcriptional landscape.

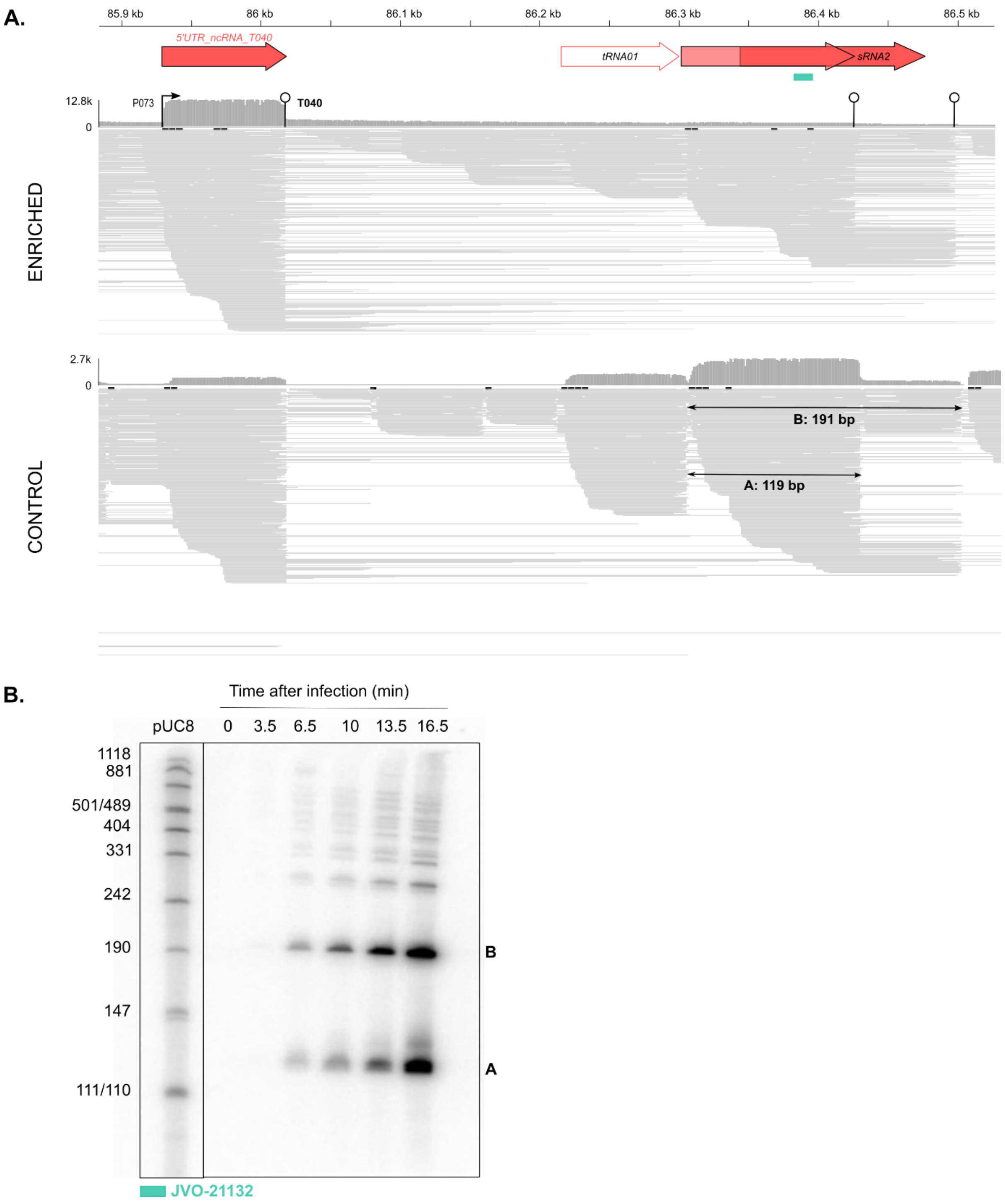


**Supplementary Figure S6: ONT-cappable-seq (A) and northern blot probing (B) of PAK_P3 ncRNA candidate sRNA2 shows the need to refines its annotation and indicates it is 3'UTR-derived. (A.)** IGV ONT-cappable-seq tracks of the enriched and the control PAK_P3 samples. Previous RNA-seq experiments revealed the presence of an 132-nt ncRNA candidate, sRNA2. Based on the TSS and TTS identified in this work, we re-annotated the transcript boundaries (now 119-nt long) and find that it is likely 3'UTR derived. tRNAs and ncRNA candidates (old and new annotation) are displayed with a red outlined white arrow, and red filled arrows, respectively. The position and orientation of the promoters (arrows) and terminators (line with circle) are indicated. Reads and terminators mapping on the Watson and Crick strand are indicated in grey and blue, respectively. The alignment view was downsampled in IGV for visualisation (window size = 5, number of reads per window = 25). (**B.**) Northern blot probing of the ncRNA candidate sRNA2 (green rectangle in part A) confirms abundance and size of the transcript (A: 119 nt) in the late infection stage and shows extensive processing, in agreement with the transcriptional landscape.

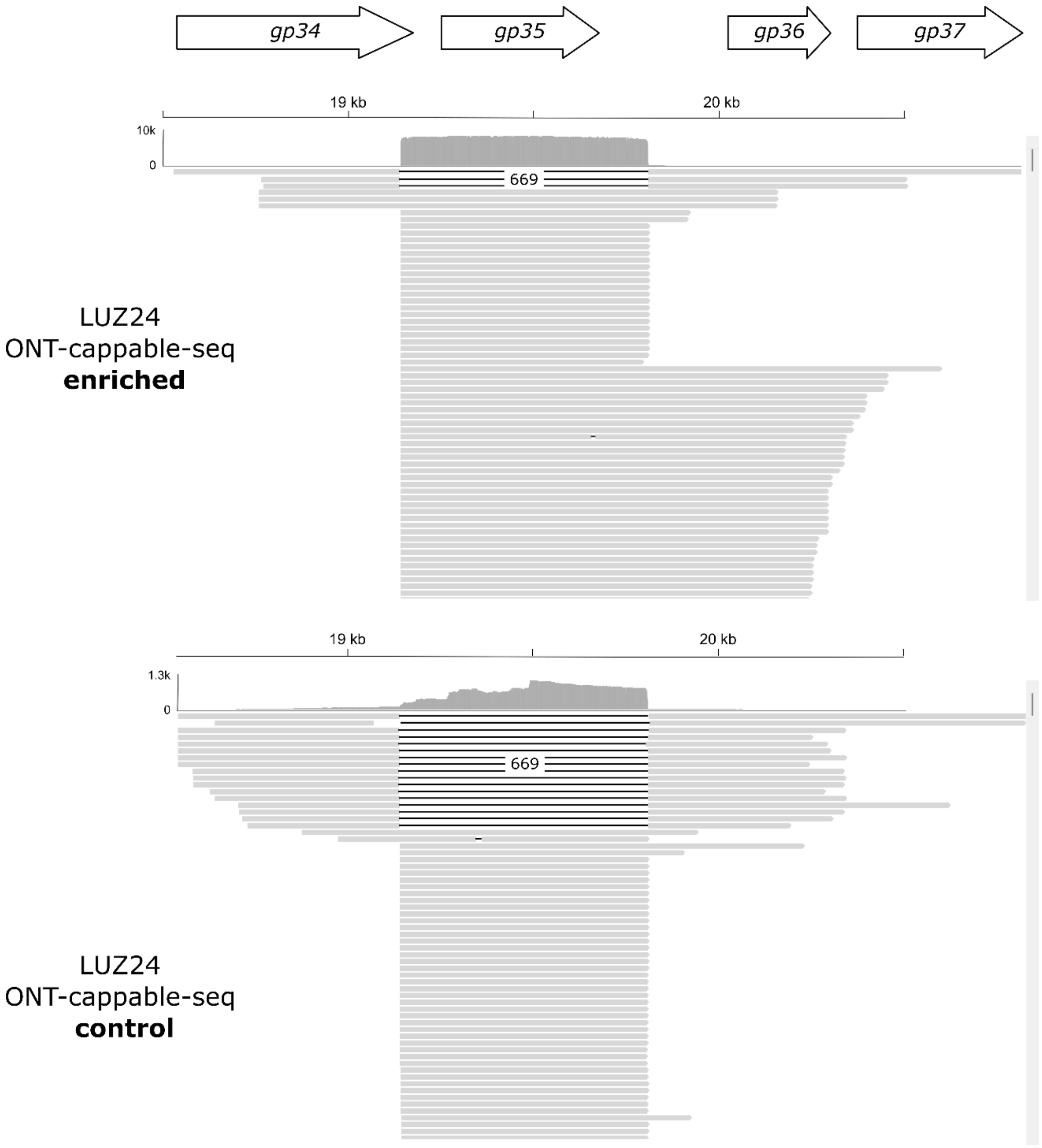


**Supplementary Figure S7: IGV ONT-cappable-seq data track window of the LUZ24 genomic region that comprises genes *gp34-gp37* and the group I intron.** Comparison between the LUZ24 transcriptional landscapes obtained by ONT-cappable-seq of the sample enriched for primary transcripts (upper panel) and control sample (lower panel). This alignment view only displays a small window, as indicated by the scroll bars on the right. Read alignment shows splicing of a 669-bp intron interrupting one of the DNA polymerase genes of LUZ24 (*gp34*). The proportion of spliced cDNA reads is higher in the control sample, which predominantly contains processed transcripts.

### Supplementary Tables

**Supplementary Table S1: Overview of primers and inserts used in this study.**

| Feature | Name | Sequence (5'-3') | Use |
|---|---|---|---|
| Primer | GlmS_Up_Pa | gtgcgactgctggagctgaa | gDNA removal test |
| | GlmS_down_Pa | gctctcgccgatcctctaca | |
| | PF,PCR,intron2 | cgatcgatcttcatgcacggctc | intron splicing assay |
| | PR,PCR,intron2 | aagcgccacccactactcgcatc | |
| | PRT,intron2 | tgcccacggtttaccatggac | |
| | PF,PCR,intron1 | gggtaagtggaagactgtaggaaag | |
| | PR,PCR,intron1 | caatgccttgtggatcatcacgtcgcc | |
| | PRT,intron1 | gccaaaggacagttaaccccgaggagttc | |
| | pBG_BsaI_F | gacggtctctaagagaattcgagctcggtac | promotor trap assay |
| | pBG_BsaI_R | gtcggtctcctagattaattaagacgtcttgac | |
| | ST_Pem7prom_F | tcggctcttccagaggtctcttctattgttgacaattaatcatcggc | |
| | ST_Pem7prom_R | atagctcttcacttggtctcgctgcggtttagttcctcaccttgtcg | |
| | ST_BCD2_F | ggcgctcttccagaggtctctgcaggcccaagttcacttaaaaagg | |
| | ST_BCD2_R | gccgctcttctcttggtctcgcattacctccttagcatgattaag | |
| | ST_msfGFP_F | gacgctcttccagaggtctcgaatgatcatgggaattcataaaggtg | |
| | ST_msfGFP_R | taggctcttctcttggtctcagttatttgtagagttcatccatgccg | |
| | 14-1_P001_F | tctacttcgatcagacactttacttccagtctgactgtcggcataatctcctca | |
| | 14-1_P001_R | ctgctgaggagattatgccgacagtcagactggaagtaaagtgtctgatcgaag | |
| | 14-1_P011_F | tctacggcaagattgttccgcgacatgaaaggcggcgtgctggattcctgggt | |
| | 14-1_P011_R | ctgcacccaggaatccagcacgccgcctttcatgtcgcggaacaatcttgccg | |
| | 14-1_P018_F | tctacaaaaagggataattcctaaccgggcctgggaaggctatactagacct | |
| | 14-1_P018_R | ctgcaggtctagtatagccttcccaggcccggttaggaattatcccttttg | |
| | 14-1_P032_F | tctagagcctcggttcgcagctatctgagcaagctggaagtcatgggctgggtc | |
| | 14-1_P032_R | ctgcgacccagcccatgacttccagcttgctcagatagctgcgaaccgaggctc | |
| | 14-1_P036_F | tctagattaaattcgctcgctttggcgagtcagctgatagtatactcttcct | |
| | 14-1_P036_R | ctgcaggaagagtatactatcagctgactcgccaaagcgagcgaatttaatc | |
| | 14-1_P038_F | tctaatttaatccaactttacttcggcaggaaagtggccgatactagagcc | |
| | 14-1_P038_R | ctgcggctctagtatcggccactttcctgccgaagtaaagttggattaaat | |
| | LUZ19_P002_F | tctacctggcgcgggtattgacacgtgggtagtaggtctgtagagttcgccct | |
| | LUZ19_P002_R | ctgcagggcgaactctacagacctactacccacgtgtcaatacccgcgccagg | |
| | LUZ19_P003_F | tctactggagacagggttgacacactgccagggcatcggtagagtacgcagcca | |
| | LUZ19_P003_R | ctgctggctgcgtactctaccgatgccctggcagtgtgtcaaccctgtctccag | |
| | LUZ19_P004_F | tctatcgctgaggggattgacacggtgcaggacacgcggtacagttcgcacca | |
| | LUZ19_P004_R | ctgctggtgcgaactgtaccgcgtgtcctgcaccgtgtcaatcccctcagcga | |
| | LUZ19_P005_F | tctaccccgacagaggattgacaagacaagcgcaagtctctaacatgcgcagca | |
| | LUZ19_P005_R | ctgctgctgcgcatgttagagacttgcgcttgtcttgtcaatcctctgtcgggg | |
| | LUZ19_P006_F | tctagatcgagaactagaaacctcggttaatcgcgactgacagtatgctgggta | |
| | LUZ19_P006_R | ctgctacccagcatactgtcagtcgcgattaaccgaggtttctagttctcgatc | |
| | PAKP3_P033_F | tctacggccaaagctgttgacaggttgtatggggccgggtaaactggccccta | |
| | PAKP3_P033_R | ctgctagggccagtttacccggccccatacaacctgtcaacagctttggccg | |
| | PAKP3_P034_F | tctatcgtctgatgaggttgctttctgacgtggagaatagtaaccttgtctcc | |

| | | | |
|---|---|---|---|
| | PAKP3_P034_R | ctgcggagacaaggttactattctccacgtcagaaagcaacctcatcagacga | |
| | PAKP3_P064_F | tctatcctgggtatattcttgacatcattgtcaagctggggtactacctttcg | |
| | PAKP3_P064_R | ctgccgaaaggtagtaccccagcttgacaatgatgtcaagaatatacccagga | |
| | PAKP3_P067_F | tctatgctctgtgggtgttcgtgcaaaggttggtgcagtgattgtccagggaa | |
| | PAKP3_P067_R | ctgcttccctggacaatcactgcaccaacctttgcacgaacacccacagagca | |
| | PAKP3_P075_F | tctagcatgctaggggcttgacaaagccaagcccctagcatatactgaccata | |
| | PAKP3_P075_R | ctgctatggtcagtatatgctaggggcttggctttgtcaagcccctagcatgc | |
| | JVO-18831 | cccgttatcactcgttgataattgatttg | Northern blot probing |
| | JVO-21131 | acctccggggcgtagggggttcg | |
| | JVO-21132 | tgctcagcggtgggagccctct | |
| | JVO-21411 | tataccccacaacataagtcat | |
| | JVO-21410 | aaccaatcggctagtatcttac | |
| | JVO-21409 | gaaacaagggggaaggatta | |
| | JVO-18830 | gttctaccaccgagtgtgtttagtgg | |
| | JVO-21375 | taatccaacagtcctgtattcac | |
| **Insert** | $P_{EM7}$ | ttgttgacaattaatcatcggcatagtatatcggcatagtataatacgacaaggtgag<br>gaactaaacc | promotor trap assay |
| | *BCD2* | gcccaagttcacttaaaaaggagatcaacaatgaaagcaattttcgtactgaaacat<br>cttaatcatgctaaggaggttttctaatg | |
| | *msfGFP* | atgatcatgggaattcataaaggtgaagaactgttcaccggtgttgttccgatcctggt<br>tgaactggatggtgatgttaacggccacaaattctctgttcgtggtgaaggtgaaggt<br>gatgcaaccaacggtaaactgaccctgaaattcatctgcactaccggtaaactgccg<br>gttccatggccgactctggtgactaccctgacctatggtgttcagtgtttttctcgttacc<br>cggatcacatgaagcagcatgatttcttcaaatctgcaatgccggaaggttatgtaca<br>ggagcgcaccatttctttcaaagacgatggcacctacaaaacccgtgcagaggttaa<br>atttgaaggtgatactctggtgaaccgtattgaactgaaaggcattgatttcaaagag<br>gacggcaacatcctgggccacaaactggaatataacttcaactcccataacgtttaca<br>tcaccgcagacaaacagaagaacggtatcaaagctaacttcaaaattcgccataacg<br>ttgaagacggtagcgtacagctggcggaccactaccagcagaacactccgatcggtg<br>atggtccggttctgctgccggataaccactacctgtccacccagtctaaactgtccaaa<br>gacccgaacgaaaagcgcgaccacatggtgctgctggagttcgttactgcagcaggt<br>atcacgcacggcatggatgaactctacaaataa | |

**Supplementary Table S2: Overview of ONT-cappable-seq sequencing yields, read lengths and mapping metrics for each sample.**

| Sample | Condition | total number of reads | Mean read length | Read length N50 | Total mapped reads | Total reads mapped to phage genome |
|---|---|---|---|---|---|---|
| LUZ19 + PAO1 | enriched | 789,320 | 541.3 | 570 | 689,504 | 118,275 |
| | control | 6,310,482 | 631.2 | 1688 | 5,380,550 | 16,543 |
| phiKZ + PAO1 | enriched | 983,507 | 643.7 | 737 | 860,294 | 129,998 |
| | control | 8,068,998 | 506.9 | 648 | 6,670,935 | 43,411 |
| 14-1 + PAO1 | enriched | 3,526,361 | 504.4 | 522 | 3,278,166 | 232,128 |
| | control | 6,076,471 | 1009.7 | 1729 | 5,692,604 | 11,901 |
| PAK_P3 + PAK | enriched | 1,490,051 | 597.5 | 556 | 1,383,016 | 143,136 |
| | control | 3,552,016 | 909.6 | 1730 | 3,266,261 | 20,599 |
| YuA + PAO1 | enriched | 6,369,687 | 342.8 | 333 | 5,062,107 | 94,273 |
| | control | 20,298,786 | 302.5 | 289 | 16,879,211 | 16,052 |
| LUZ24 + Li010 | enriched | 5,649,112 | 337.1 | 313 | 5,062,409 | 214,226 |
| | control | 22,083,138 | 353.1 | 385 | 18,206,325 | 186,255 |

**Supplementary Table S3: Overview of phage transcription start sites (TSS) and associated promoter sequences, as identified by ONT-cappable-seq data analysis.**

| ϕ | ID | TSS position | +/- | 50bp upstream region | predicted type | Experimental validation | Note |
|---|---|---|---|---|---|---|---|
| 14-1 | 14-1 P001 | 707 | - | CTTCGATCAGACACTTTACTTCCAGTCTGACTGTCGGCATAATCTCCTCA | sigma70 | this work | |
| | 14-1 P002 | 822 | + | ACGAACGGTTGTGCTCGACCGTCTGCTGCGGTGTCGATATACTCGGCCTA | sigma70 | - | |
| | 14-1 P003 | 3708 | - | CTCATGCGCCTGGACGATAGCGACGGTCGGCGCGCCTTCAATGCCGGGAC | - | - | |
| | 14-1 P004 | 3974 | - | TGAACATGTTCAACTCCCTTGTCGAAGCGGTTGATCGTCTGTGCCGGGAG | - | - | |
| | 14-1 P005 | 4070 | - | CCAAGGCTTCTTGGCACGCCGCAAACGGCGGGACTACGAAGCGCTGGGGT | other motif | - | |
| | 14-1 P006 | 4479 | - | GTGGAAGAAAATGCTTTACTTCCATTCTGAATGAAGGCAAAATAGCTCCA | sigma70 | - | |
| | 14-1 P007 | 5200 | - | CAGAGGCAAAGATTTGTTGTTGACGCCGGGCAGCAATATTCTATGTCTGC | sigma70 | - | |
| | 14-1 P008 | 5686 | - | GACCAACCACGCATCCATCGAAATGGGCATTGGACCGATCTGTGCCGGAA | - | - | |
| | 14-1 P009 | 6637 | - | GGTAGTGCGAAAGTGCTACCGTTTTCCATTTCTGGCGTATAGTCAACCCA | sigma70 | - | |
| | 14-1 P010 | 8684 | - | CGACGAAAAGCGCTTTACTCCTGGCGAGCAGATGGAATAGACTCTGCTCA | - | - | |
| | 14-1 P011 | 8970 | - | CGGCAAGATTGTTCCGCGACATGAAAGGCGGCGTGCTGGATTCCTGGGTG | other motif | this work, no significant activity in vivo | |
| | 14-1 P012 | 9116 | - | CTGAGAAATAATTCTTTACGGGAACTTTTCGGCAGATTAGACTCTAATTA | sigma70 | - | |
| | 14-1 P013 | 9247 | + | TGTGTTTGAATCTCTTTTGAACGTTTGATGTTTCCCCTATAATAAGCGCA | sigma70 | - | |
| | 14-1 P014 | 13257 | + | TATTGTTCCGCTTATTTACCCTTAGATGTACTGCGTATATAATACAGCCA | sigma70 | - | |
| | 14-1 P015 | 18699 | + | TACGGCGGAACTATCGACCATCCGGGCGGGACCAGGTATATTAGGGACGC | sigma70 | - | |
| | 14-1 P016 | 23797 | + | GGCGTGCAGATTCAGTCCCTTTCCGATGCAAATTTGCTAGACTTGGGAGC | sigma70 | - | |
| | 14-1 P017 | 23847 | + | CACCGGCGATTCGATATCTTCGGCCGCAAAATCGCTATATAATCGCGTGA | sigma70 | - | |
| | 14-1 P018 | 28318 | + | CAAAAAGGGATAATTCCTAACCGGGCCTGGGAAGGCTATACTAGACCTGC | sigma70 | this work | |
| | 14-1 P019 | 31714 | + | GCATACATCGGCCTACCCACTAGTGGGGCATTCAAGTTAGAATACAGAGT | sigma70 | - | |
| | 14-1 P020 | 36158 | - | CGTTAAACAGTTAAGCGCGAGCATTTTCGACGGGTTATAATAGCGCCCGA | sigma70 | - | |
| | 14-1 P021 | 36804 | - | TTCCATCGAACAGTTCAGCGAAGTCGAGCGCCGCCACACGATGCTGGGAT | other motif | - | |
| | 14-1 P022 | 38929 | - | AGACGGTTCGGTCAGCGACAAGAAGTCTGACTACTTCGCCGACACCCGTG | - | - | |
| | 14-1 P023 | 39095 | - | GCCTGCTGATCATTCCGGCCGCGTACCGCCGCGAGTTCATCCGCTGGGCC | other motif | - | |

| ϕ | ID | TSS position | +/- | 50bp upstream region | predicted type | Experimental validation | Note |
|---|---|---|---|---|---|---|---|
| | 14-1 P024 | 39481 | - | TAATATGTAATTTGCTTTACTTAGTTGCAACTTCCGTTATTATAAACCCA | sigma70 | - | |
| | 14-1 P025 | 40195 | - | AATATGTAAATCAGTAGCCGTATCAAGGAAGATGCGGCATAATATCTACT | sigma70 | - | |
| | 14-1 P026 | 43097 | - | ACTGTGTCCATACTTCGCGAATTGGCCAAGGAAGGCGTAGAATATACGGC | sigma70 | - | |
| | 14-1 P027 | 48622 | - | TTTCTCTTTCTCCGCTTCCACCACACTGGTCGCGGCCATCGACTGGGTGA | other motif | - | |
| | 14-1 P028 | 49234 | - | ATTTACAAAAGTGCTTTACTTCTGGATCAGGTGGGCGTAATATTCCCTCA | sigma70 | - | |
| | 14-1 P029 | 49599 | - | ACCCTAATCGTCGAATCCATTACTAAGGGAGTGGTGGAGACTGCTGCGGG | - | - | |
| | 14-1 P030 | 50236 | - | CATGATTCCTGATCAGAAAGTCATCATCGTAGGTGCTGGACTCGCCGGAC | - | - | |
| | 14-1 P031 | 50440 | - | ACGAAGCTGGCCAGGGCGATGACGAGAAGGCCGGTGATGAAGGCCAGGAG | - | - | |
| | 14-1 P032 | 50518 | - | GAGCCTCGGTTCGCAGCTATCTGAGCAAGCTGGAAGTCATGGGCTGGGTC | other motif | this work, no significant activity in vivo | |
| | 14-1 P033 | 51005 | - | AAGGAAAAGGCCGCCGCCAAGGAGGCCGAGCGTGCTGAAAAGGCCAAGGC | - | - | |
| | 14-1 P034 | 51317 | - | TCAGGCGAAAGGTATTCCCTTCTTCAATTAGTTTAGAGATAATCCTCTCA | sigma70 | - | |
| | 14-1 P035 | 51666 | - | TGCTTTTTGCCAAAGTTCGCATTATCGGCACGAACTATGAGCGCTGGGTG | other motif | - | |
| | 14-1 P036 | 53073 | - | GATTAAATTCGCTCGCTTTGGCGAGTCAGCTGATAGTATACTCTTCCTGC | sigma70 | this work | |
| | 14-1 P037 | 53097 | + | ACGTACAGTTATGCTTTACCTCTGCGCAGGAAGAGTATACTATCAGCTGA | sigma70 | - | |
| | 14-1 P038 | 53163 | + | AATTTAATCCAACTTTACTTCGGCAGGAAAGTGGCCGATACTAGAGCCGC | sigma70 | this work | |
| | 14-1 P039 | 54251 | + | TGAGCGCGTCGAAGTCGGGTTTCTGGGGAAGTGCGCCGCTTGCAAGGGAA | - | - | |
| | 14-1 P040 | 59399 | + | ATTAATAAAATTAATAGTAATTTGGTAATTTGGTATACTTTAGTATTTGA | sigma70 | - | |
| | 14-1 P041 | 59807 | + | CACAAGCAGCCCATAGACGCGATCCCTGGCCCCATAGTACAATCGCGCCA | sigma70 | - | |
| | 14-1 P042 | 60949 | - | AAGAGAATCTGAAGCCGGAAGAAGAAATCACCGAAGAGGAAGCGCTGGAA | other motif | - | |
| | 14-1 P043 | 61300 | - | GATGAAATAGATGTAGACACGCAGCATCGACCTAGGCATAATCTCTTTCA | sigma70 | - | |
| | 14-1 P044 | 62417 | - | TTGGAGGGCAGCTTATGTCCGATAAAGAACAAAGTGGCCATTACTGGGTG | other motif | - | |
| | 14-1 P045 | 62463 | - | GCGACTAACAACAACCAGCCTCTGCGACTCGGCAGAGGCTTTCTCATTGG | - | - | |
| | 14-1 P046 | 62562 | - | GCGACGAAATCACCCTGGAGCTTACCTGCCCGGCCGGCGACTACTGGGAA | other motif | - | |
| | 14-1 P047 | 63024 | - | GACGCAGTGTCCATCCTTCACAAACGTTAATCCATCGGCGCGGCTGGGAT | other motif | - | |
| | 14-1 P048 | 63303 | - | ATGCTTTACTCGGCGGTGAGAATCAGGGCATAATCTCTTCACCGGCTGAG | - | - | |
| | 14-1 P049 | 65240 | - | TTCCTCAGAAGTGCTTTACTCTATGAGATGACTGAGGTATAGTTACCTCA | sigma70 | - | |
| | 14-1 P050 | 65678 | - | CGAAACGAATAGCCAGCTCGCCCTTCGAATCCGCGAGCTAGAGGCCGAGC | - | - | |

| ϕ | ID | TSS position | +/- | 50bp upstream region | predicted type | Experimental validation | Note |
|---|---|---|---|---|---|---|---|
| LUZ19 | LUZ19 P001 | 732 | - | ACCCTGGAACCCGCGTGGCACTAAGGCTGCGGATATGTCACACACAATGG | phage promoter | - | |
| | LUZ19 P002 | 950 | + | CCTGGCGCGGGTATTGACACGTGGGTAGTAGGTCTGTAGAGTTCGCCCTG | sigma70 | this work | |
| | LUZ19 P003 | 1029 | + | CTGGAGACAGGGTTGACACACTGCCAGGGCATCGGTAGAGTACGCAGCCA | sigma70 | this work | |
| | LUZ19 P004 | 1182 | + | ATCGCTGAGGGGATTGACACGGTGCAGGACACGCGGTACAGTTCGCACCA | sigma70 | this work | |
| | LUZ19 P005 | 1284 | + | CCCCGACAGAGGATTGACAAGACAAGCGCAAGTCTCTAACATGCGCAGCA | sigma70 | this work | |
| | LUZ19 P006 | 1561 | + | GATCGAGAACTAGAAACCTCGGTTAATCGCGACTGACAGTATGCTGGGTA | - | this work | |
| | LUZ19 P007 | 21134 | + | GGGACTCCTGGGCGTTCTTCTGCTGAGGCGGATTATGTCACCCACATAGG | phage promoter | - | |
| | LUZ19 P008 | 21331 | + | GATGGCTACTATGAAGACCCACCGCCCTACGGTTATGTCACCCACAGTGG | phage promoter | - | |
| | LUZ19 P009 | 35609 | + | GACTGGACGCCAGTCCTGGCGCGGGTTCGCGGTTATGTCACATACAGAGG | phage promoter | - | |
| LUZ24 | LUZ24 P001 | 238 | + | CTTGATCCCCTATTGACATCCCACGATACCCTATGTTAATCAAAGACTGT | sigma70 | - | |
| | LUZ24 P002 | 331 | + | TATGAAAAACCAGTAGACAACCCCGTAAAGAATATGGCGTAATGGCGCCG | - | - | |
| | LUZ24 P003 | 410 | + | AAACCCCAGGGGCTTGACAAACCCCGCGGAACCGGGCCTAATGGCCCCCA | sigma70 | - | |
| | LUZ24 P004 | 554 | + | AGGTATCGGGGGTTGACAAGGGATCGGTGAAGCGGTATAGTTCGCCCCGG | sigma70 | Ceyssens et al. (2008a) | |
| | LUZ24 P005 | 657 | + | TGGCCGGCACCGCTTGACACTGTAAGGCTCAGTCGGTATGATGGGCACCA | sigma70 | Ceyssens et al. (2008a) | |
| | LUZ24 P006 | 4869 | + | GTAACATCTAGGCCTTGACATTCTGCTGGAAGTGTGGTATAATAACCTTA | sigma70 | Ceyssens et al. (2008a) | |
| | LUZ24 P007 | 5143 | + | CTGTAAAGAAGGAGGTGCATCGTGGCCACGGTAACTAGAATCAGTAAGGG | sigma70 | - | |
| | LUZ24 P008 | 5249 | + | CGGAATCCCGGAAGGACCTCACCATCCCGATGGTAAACATCTGCAATGGG | - | - | |
| | LUZ24 P009 | 6630 | + | GCCACCTGTTGGTATCTACCCACGAGAATACCTCTGGTAAACTGGTGGAG | sigma70 | - | |
| | LUZ24 P010 | 17248 | + | TTCCACAAGAAGGCTTGACAAATCCAGAAAAGTGTGGTATAATAACCTTA | sigma70 | - | |
| | LUZ24 P011 | 19143 | + | CAGATAGCCGCAGGGCTTCCCACACGGGACGATGCTAAGACCTTCATCTG | - | - | 5' boundary of group I intron |
| | LUZ24 P012 | 19353 | - | TGATCCTACCCCACTCCTTGCACTCGTCTGAGCAGTAGTGGTGGCTAGGG | - | - | |
| | LUZ24 P013 | 25961 | - | ACTTTGAGCCAGTGAAATGGGAGGATATTGAGGGGAAGCCTGAGCTTCTG | - | - | |
| | LUZ24 P014 | 28915 | + | TAGCTTCTGGGGTATTGACACCTGCGGATGTATCTGAGATAATCCTGTCG | sigma70 | - | |
| | LUZ24 P015 | 39133 | - | ATTGTGTCTTATGGCAAATCGATTGGATACTCCGAGGATGAGATCCGAGG | - | - | |
| | LUZ24 P016 | 44883 | - | AAAGAAGAAAAAAGACTTGACAAAATAGAAAAAGTGTGATATAATAGTAT | sigma70 | Ceyssens et al. (2008a) | |

| ϕ | ID | TSS position | +/ - | 50bp upstream region | predicted type | Experimental validation | Note |
|---|---|---|---|---|---|---|---|
| PAK_P3 | PAK_P3 P001 | 100 | + | CAACTCCACTGCGAACATCACCGTGTTCGGTGGAGCTGCCGGGGCTGGTA | - | - | |
| | PAK_P3 P002 | 300 | + | TCAAATGGAAGGACAAGGACGGGAAATTCGTCTTCTCCTCGGGGGCTGAG | - | - | |
| | PAK_P3 P003 | 768 | - | TGAAGTAGCCGCTTGCTTCTTCTCGGGCAAACCAGCTACCATGGAGCAGT | sigma70 | - | |
| | PAK_P3 P004 | 1638 | - | CTCTGTACCGCCCTTGACACTCCATTCTACACGGCTTACCATCAATTCGA | sigma70 | - | |
| | PAK_P3 P005 | 2898 | - | TTGGGAGTCGGCTGGTTTTCTTCAGCCATTTCAGTTATCCTGGTTTGCGG | sigma70 | - | |
| | PAK_P3 P006 | 5470 | - | CGCCGGAGCGAAGAAGGTCTTGAACAGGTCTTCGGTGCCGCTGCCGGGGA | - | - | |
| | PAK_P3 P007 | 5814 | - | GCGGATGTTGCCATTGTACATGTCGATAAAATATTGGTAATCTTCGTCCG | sigma70 | - | |
| | PAK_P3 P008 | 6167 | + | TCGCGATTTGCGGCTTTACGAGCAGCAGAACGTGGGGTACAATGACATGC | sigma70 | - | |
| | PAK_P3 P009 | 9603 | + | AATATGAACACCTTGACTTACTCTCCAAGAGAGGTTACGATCCTGATCGC | sigma70 | - | |
| | PAK_P3 P010 | 12231 | + | AAAGATGTTGTTGACAGCATCTTCCGTTCCATGACAGAGTATGGTACCGT | - | - | |
| | PAK_P3 P011 | 17986 | + | ACCAAGATGGGCTGGACAAAATTGGTAAAGAGGTGGTATACTCTAGACTA | sigma70 | - | |
| | PAK_P3 P012 | 18530 | + | CAGTCGGGGCCCCTTGGTGGGGTTTGGGGCAAGCTACTGAGATTTCCCGG | - | - | |
| | PAK_P3 P013 | 19745 | - | CCACTATACCTGCGTAATCGGCTGTCAGAGGTGGGAGCGTGATATCCTGG | - | - | |
| | PAK_P3 P014 | 21185 | - | TGGCCATTTCTACTTGACGATCTGCATCTCGTAGTTCATTATCTCTCCAG | sigma70 | - | |
| | PAK_P3 P015 | 23122 | + | AGCTGGTGGGTATGGAAAATACAGCGAGCAACACGCTAAACTGGCCCGTG | sigma70 | - | |
| | PAK_P3 P016 | 24923 | + | CCGGAGAAAAGTAGTTGACATCCCCCACAACACCCTATACAGTGGCCACA | sigma70 | - | |
| | PAK_P3 P017 | 25397 | + | ATCACTGGAACCGTTGACACAGGAATATAAAGGAGGAACAATGAAGAACA | - | - | |
| | PAK_P3 P018 | 26977 | + | ATCCCACAACTGTTGACACAGGGGAGAAGGAGGAGTATCGTGCATACCGT | sigma70 | - | |
| | PAK_P3 P019 | 29755 | + | TTATGCCGACATCCTTGCCATGGCTGAAGGTAAGAGCTATAATCCTGCCA | sigma70 | - | |
| | PAK_P3 P020 | 29962 | + | TCGCCAGATTCGAGACGATGAAGAGTTCGCCCGACTGGTCGAGCTGGGCA | - | - | |
| | PAK_P3 P021 | 30016 | + | ACAGGCCGACATCATGAATGAATCGTTCTACTGGGTTGACATGCAGGGGC | - | - | |
| | PAK_P3 P022 | 30038 | + | TCGTTCTACTGGGTTGACATGCAGGGGCGTTTCAGGCAGTATACTCCTCA | sigma70 | - | |
| | PAK_P3 P023 | 30858 | + | GTATGTGAATGGGGTAGACGGCAATGACAAGAAAGCCCAGTATGACACCG | sigma70 | - | |
| | PAK_P3 P024 | 31149 | + | CGAATGGATGTTGACTTTACCCAGGTAACTGTGGTACAGTCTGGTCCTGG | sigma70 | - | |
| | PAK_P3 P025 | 31313 | + | TCTCCTGCTGGCGTAACCCTAAGGGGCCCTGATTGGGCCCTTTTTATTGC | - | - | |
| | PAK_P3 P026 | 31341 | + | CTGATTGGGCCCTTTTTATTGCAGAAGGAGTAGGAAGTGCTGATTGGGAA | - | - | |
| | PAK_P3 P027 | 36310 | + | TTTGATAAAAGTCCTTGACTTTCAATAGCTTATATGCTTCAATAGCGTTC | sigma70 | - | |
| | PAK_P3 P028 | 41196 | + | CCTGGAAGAGGTTTACTTGAAGCTGCGTGCGTGGTATAATACCCACCCGT | sigma70 | - | |

| ϕ | ID | TSS position | +/ - | 50bp upstream region | predicted type | Experimental validation | Note |
|---|---|---|---|---|---|---|---|
| | PAK_P3 P029 | 42029 | + | AAGGGCAACAACGCTGTCTATGACGCCGAGAAGATGGTATTCTTCTCTCA | sigma70 | - | |
| | PAK_P3 P030 | 44517 | + | GGAACCATGGGCTTCCATACTCTGCTACAACAGCGTATGATCCCGTTCGA | sigma70 | - | |
| | PAK_P3 P031 | 46660 | + | GTACAATTTCATGTTGACTAGCCGAGCCGGTTGACCTATCATAGCCCCCA | sigma70 | - | |
| | PAK_P3 P032 | 47278 | + | TCACATTTTCCTTGAACTGGTGGTCAGACGGGAGTACAATCTGGTATCGT | sigma70 | - | |
| | PAK_P3 P033 | 47939 | + | TACGGCCAAAGCTGTTGACAGGTTGTATGGGGCCGGGTAAACTGGCCCTA | sigma70 | this work | |
| | PAK_P3 P034 | 48439 | + | TCGTCTGATGAGGTTGCTTTCTGACGTGGAGAATAGTAACCTTGTCTCCG | sigma70 | this work | |
| | PAK_P3 P035 | 51049 | + | TTTCTTAAAAAGGGGTTGACTTATGCCATGGCTGCCATAGAATGACCCCA | sigma70 | - | |
| | PAK_P3 P036 | 51771 | + | GTGAAATAAACGCTTGACAAACTGGCTAACCGCCCGCACAATAGCACTCA | sigma70 | - | |
| | PAK_P3 P037 | 51944 | + | CCGCTGGGGAGTCAAGCTGAATATCTCCCCCAGCGTCCGAGTGCTGGGCT | - | - | |
| | PAK_P3 P038 | 52664 | + | CCTGAAATAAAGTGTTGACACGGCATCTCGTGTCTGTAGAATAGGCATCA | sigma70 | - | |
| | PAK_P3 P039 | 53142 | + | AGGAAGAATAAGGGTTGACAAGGGGCCAGGAAACTCATAAACTGGCCCCA | sigma70 | - | |
| | PAK_P3 P040 | 54361 | + | GATCAAAAAGGGCTTGACGTGCGCAACGGATGGCCCTAGTATCTCTCTCA | sigma70 | - | |
| | PAK_P3 P041 | 55259 | + | CCTGAAATAAAGTGTTGACACTATAGGCCCTTCCTGTAGAATGGCCGCCA | sigma70 | - | |
| | PAK_P3 P042 | 55652 | + | CGAATCCCCGGGGCTTGACACCAGCGATAGTCCCGATAGAATAGCCATCA | sigma70 | - | |
| | PAK_P3 P043 | 57078 | + | TGACCGTATAAAGTTGACAAGACCCTTTTGGGTTTGTAAGATGCGTGTCA | sigma70 | - | |
| | PAK_P3 P044 | 57408 | + | CCCTGAATAAAAGCTTGACTTCCCTCCCGCTTCCCGTAGACTAGGCTTCA | sigma70 | - | |
| | PAK_P3 P045 | 58020 | + | TCTTCTAAAATCGCTTGACAGCTTAGGCCATTCGCGTAGAATGGCCCTCA | sigma70 | - | |
| | PAK_P3 P046 | 58656 | + | TTTTCCTACAGGGCTTGACCTGGCCCCCTCTTCCTGTAGAATGCGCATCA | sigma70 | - | |
| | PAK_P3 P047 | 58850 | + | AGTGAAAAGTGGATTGCATTCATGGTGGAGCGTGGGTATAGTGCCTCTGA | sigma70 | - | |
| | PAK_P3 P048 | 58993 | + | TGACCGTTTGGACCGAAGAATTTGAAAAAGCGGCAGCGGCTGAGGGCTGG | - | - | |
| | PAK_P3 P049 | 59369 | + | CTGAAAAACATCTTGACGGGGGTCCCAGGCTTCGGCATAATGACCCCCGT | sigma70 | - | |
| | PAK_P3 P050 | 59950 | + | TGGGGATTTAATGCTTTACGGGGCCCGGCAAGGGCCCCATAATGCACCCA | sigma70 | - | |
| | PAK_P3 P051 | 60536 | + | CGCAAATAAGGCTGTTGACAGCCTAGGCCATTCCTGTAGAATGGCCATCA | sigma70 | - | |
| | PAK_P3 P052 | 61068 | + | TTGACAGAAAAAGCTTGACTCCTCTTCTCCCTGGGGGCATTATGTCTCCA | sigma70 | - | |
| | PAK_P3 P053 | 63712 | + | TGAATTGAAAAATGTTGACAGGCCCCTGCCGGGGCTTCACAATGGCCCCA | sigma70 | - | |
| | PAK_P3 P054 | 63855 | + | AGACTCCCAAGGACATCTGTAGTGCCCTGTGTGAGCTGGCTAGCTGGGCT | - | - | |
| | PAK_P3 P055 | 64001 | + | CTGGCGATGGTGAACTGGAAGCTGGAATACCTAATGTCCTGGGCTGGGTT | - | - | |
| | PAK_P3 P056 | 64117 | + | GAACCCCAACAGGTTGCGTTTCCTGGACATGGTGACTATGATGTCCAAGA | sigma70 | - | |

| ϕ | ID | TSS position | +/- | 50bp upstream region | predicted type | Experimental validation | Note |
|---|---|---|---|---|---|---|---|
| | PAK_P3 P057 | 65172 | + | AGAATGAATACCTTGACTGGGAGTACGAACTGTACGATAATCCCAAGCGA | sigma70 | - | |
| | PAK_P3 P058 | 66422 | + | AGCAGCATGGACCTTGCCCGTGCCCTGGAAGTGCTGTATGATGCAGGCTA | sigma70 | - | |
| | PAK_P3 P059 | 67450 | + | GTGAGGAAGGGTTTGACACAGAGGGAGAGATTTGGAAGAATGATCGCCGA | sigma70 | - | |
| | PAK_P3 P060 | 70777 | + | GGTTTGCCCCTATTGCAAGAACCTAAGGCCTGAGTTATAGTCGGGCCTGA | sigma70 | - | |
| | PAK_P3 P061 | 72404 | + | TCCAGCCCCAGGACTTGACAGTCCTGGGGCTTTGTCGTATCTTCATCTCT | sigma70 | - | |
| | PAK_P3 P062 | 72475 | + | TGAAAAAATTCTCTGAGACTTACTACTCCTGGTCTGGGGCTGCCAGCCGG | - | - | |
| | PAK_P3 P063 | 78100 | - | TACCTTCCTCCCCTTGATTTCCACAGAGGAAGTCCGTAAACTGCTCCTCA | sigma70 | - | |
| | PAK_P3 P064 | 78951 | - | TCCTGGGTATATTCTTGACATCATTGTCAAGCTGGGGTACTACCTTTCGG | sigma70 | - | |
| | PAK_P3 P065 | 79497 | - | TTCCAGGGGTGGTGTACTTCCGTTGCCAGTTCTCGTACACTTGTGATGGA | sigma70 | - | |
| | PAK_P3 P066 | 80430 | - | CCGTAGACTAAGGTTGACAACCCCAAGGAGGCCAGGCATACTGGCCTCCA | sigma70 | - | |
| | PAK_P3 P067 | 82342 | - | ATGCTCTGTGGGTGTTCGTGCAAAGGTTGGTGCAGTGATTGTCCAGGGAA | sigma70 | - | |
| | PAK_P3 P068 | 83179 | - | CGCCTGAAATAAGTTGACAGAGCAGGAGAGTCTGTGTAGGATGCCCCTGC | sigma70 | - | |
| | PAK_P3 P069 | 84247 | - | TAGCTTGTCAAGCCTTGACTTCTGCTTCAGATGGTGGTATAGTGTACTCT | sigma70 | - | |
| | PAK_P3 P070 | 84321 | + | GCAGAAGTCAAGGCTTGACAAGCTAAAGAATTTATAGTAATATTATTAAA | sigma70 | - | |
| | PAK_P3 P071 | 84633 | + | ATGTGACTAGGGATTGACAAACACACCTACAGGATGCTATAGTATCTATA | sigma70 | - | |
| | PAK_P3 P072 | 85495 | + | TCCACCGGCACTCTTGACACGGCGTACTGAACACGGTTTAATACCCTCCA | sigma70 | - | |
| | PAK_P3 P073 | 85931 | + | GCCCGCCAAAATCTCTTGACTTTATCTGAAAATGTGGTAAAGTGGATTTA | sigma70 | - | |
| | PAK_P3 P074 | 86865 | - | ATCCACTATCTGTTGCGCCAGGCCTCCAGGGCCCGTAAGATATGGAATGG | sigma70 | - | |
| | PAK_P3 P075 | 88067 | + | AGCATGCTAGGGGCTTGACAAAGCCAAGCCCCTAGCATATACTGACCATA | sigma70 | this work | |
| phiKZ | phiKZ P001 | 2176 | + | GCTTTATGTCATGTATTCAAACATATTTAAACAGTATATTACAAAGGTGA | early promoter | Ceyssens et al. (2014); Wicke et al. (2021) | |
| | phiKZ P002 | 4180 | + | GAAATGAAATAATACATTCAAACCATTTTAACCAGTATATCACAACAGTG | early promoter | Ceyssens et al. (2014); Wicke et al. (2021) | |
| | phiKZ P003 | 10695 | + | acaggaaattctaatggaagacttcaaagaaatcgagatttacagtatga | late promoter | - | |
| | phiKZ P004 | 12388 | - | TACAAGCTGACCATTGCCAATGTAACCTAGGAAAGTTTTATTATCATATA | - | - | |
| | phiKZ P005 | 12482 | - | GTTAAGCAACTCTAGCGAACAAAAGATTACTGGAGTTAGAATTAATATGA | middle promoter | Wicke et al. (2021) | |
| | phiKZ P006 | 14812 | - | GGTGGTTCCTAGGGGGCTTTATAACGTCTTTTAACACATTATCCATATGA | late promoter | - | |
| | phiKZ P007 | 19891 | - | GTTGATGCTCGTCTATATAAAAAGATTGAACAACGTACTGGTATTCCTAG | - | - | |

| ϕ | ID | TSS position | +/- | 50bp upstream region | predicted type | Experimental validation | Note |
|---|---|---|---|---|---|---|---|
| | phiKZ P008 | 19894 | + | TGCCATCCGAATATATTGGAACTGCCATTCGTTCTGCTGAATAATCCTAG | late promoter | - | |
| | phiKZ P009 | 20123 | - | AACTCTATACAATCCCACCGGCGGTGGAAAATTATATCCTAAACAATATG | late promoter | Ceyssens et al. (2014) | |
| | phiKZ P010 | 20273 | + | ATGGATAGTAAAAAATAACTAGTATGGTCAACTATGATTAATATCTATGA | late promoter | - | |
| | phiKZ P011 | 21311 | - | AGGAAGTCAATTTCACCAGGTGCAGTAGTATGCGCAGCAGCGGCTGGGTT | - | - | |
| | phiKZ P012 | 23599 | + | GTTGTTAAAAACTATGCTAAAAAATAATTAAGATATATATTACATGTTTG | early promoter | Ceyssens et al. (2014); Wicke et al. (2021) | |
| | phiKZ P013 | 26501 | - | gtaattacatggaataaataacataagtacttttatgacttatgatttga | middle promoter | - | |
| | phiKZ P014 | 26543 | + | TGGAGAATCAAATCATAAGTCATAAAAGTACTTATGTTATTTATTCCATG | middle promoter | Ceyssens et al. (2014); Wicke et al. (2021) | |
| | phiKZ P015 | 27554 | + | TTGCCATCTACTTTTAAGATAACTGAATCTACTTTTGTTGCTGCTGATTT | - | - | |
| | phiKZ P016 | 27710 | + | CCTCTCCTTATGGAGAGGGCTTATGATGTATTTTCTCTTTTATCATCTGT | late promoter | - | |
| | phiKZ P017 | 29413 | + | AGGCTTTATGCTGAGTTAAAAAAAGTTTAAGATCTATATTACTAATTTGC | early promoter | Ceyssens et al. (2014); Wicke et al. (2021) | |
| | phiKZ P018 | 30150 | + | AAGGTGTAACCAGAATTAAAACAAGTGATCGAGATGCAGCTGATATGCTG | - | - | |
| | phiKZ P019 | 33472 | + | AAATCAGAATTTTGCAGATATTAAATAACTAGCGATAATTAATAATCTGT | middle promoter | - | |
| | phiKZ P020 | 35112 | - | ATGAGTTATTTGTATCTATGGGTGAAGCACAGTTATCAGGTGGTGCTGAG | late promoter | - | |
| | phiKZ P021 | 39463 | + | GTTATATTCTAGAAGGTGAAGACTTTGGTATGGACTGTTACAATGATGTG | late promoter | - | |
| | phiKZ P022 | 39935 | + | AAAAGGTCTGTTTGGTGTAGTTGTATTACTAGATGAGTAATACCCTATGT | late promoter | - | |
| | phiKZ P023 | 40441 | + | TATGGCTCCTTTTATGTTGTTTTATTTTTACTTACTTACATCATCTAATG | middle promoter | Ceyssens et al. (2014) | |
| | phiKZ P024 | 42692 | + | TTATTACGCGTGTTGCTGACGTTAGAACTATCATTGGAAATACTATGGGT | late promoter | - | |
| | phiKZ P025 | 44937 | + | ATGGTTATTATCCGATTAAAAATTAATTGAGATCTATATCATAGTTTGGT | early promoter | Ceyssens et al. (2014); Wicke et al. (2021) | |
| | phiKZ P026 | 45377 | + | AGTAAATGATAACTGCATTTTATAAAAATACTGATAGTTATTATTTAATA | middle promoter | Wicke et al. (2021) | |
| | phiKZ P027 | 45490 | + | ATGGAAATTGTAAAATGAAAAATAAATAATACTCAGATTGTAATAATATG | middle promoter | - | |
| | phiKZ P028 | 45583 | + | AGCGTCTATGTTGTAATGAAAAAAATTTGAGATTTATATTACTAAATTGA | early promoter | Ceyssens et al. (2014); Wicke et al. (2021) | |

| ϕ | ID | TSS position | +/- | 50bp upstream region | predicted type | Experimental validation | Note |
|---|---|---|---|---|---|---|---|
| | phiKZ P029 | 49168 | + | TAATGATAATTCTATCAGCATGTCAGTTTTATGCCTTAAAGCGCCTAATG | middle promoter | Wicke et al. (2021) | |
| | phiKZ P030 | 51729 | + | GCGCCGTGAAGTTATACTGAATTCAATTTCAGATTTATATTACTAAGGTG | early promoter | Wicke et al. (2021) | |
| | phiKZ P031 | 52559 | + | AGTCTGCACGAGAAAGTTGACATTATCCGTATGTGGAATGCTATTGATGG | late promoter | - | |
| | phiKZ P032 | 53699 | + | CCAAGATATTATTAATAGGGAAAGAAGTACTTCATTACCTTATCTGTTGA | middle promoter | Wicke et al. (2021) | |
| | phiKZ P033 | 57100 | + | AGCATACAAGAAATTCAAATAGAAGAAATACTTGTTGCGATAATATTACG | middle promoter | Wicke et al. (2021) | |
| | phiKZ P034 | 57998 | + | ACTAAATGCCGTTATACTAAAAAAAATTTGAGATCTATATTACTAACATG | early promoter | Wicke et al. (2021) | |
| | phiKZ P035 | 64454 | + | TCAGAAGATGCCAGCTGGTGCTTTCAAGATGACTGATTTTCATGGTGGTA | - | - | |
| | phiKZ P036 | 68441 | + | AAAGATTAGCCTTAATATTTCCCATTGTGAGATCTTGATATACGCTATGA | late promoter | - | |
| | phiKZ P037 | 70913 | + | TTGTGGTTATTTAATATGATAATCATGCATATTAAATAGACATGCTATGT | late promoter | - | |
| | phiKZ P038 | 74086 | - | ATAGTTATTCTTTAATATAGTGTAAAATTACCAAATGTGTTAAATATATG | middle promoter | Wicke et al. (2021) | |
| | phiKZ P039 | 74150 | + | TTTGGTAATTTTACACTATATTAAAGAATAACTATCTGTCCAACCCTATG | middle promoter | Wicke et al. (2021) | |
| | phiKZ P040 | 77926 | + | TGCTAACCATCGTCTAACTGGTGTAGTTAATGATATTGCCGATATTGCTG | middle promoter | Wicke et al. (2021) | |
| | phiKZ P041 | 81776 | - | GGGAGGCTTTATGCTGATTATTACTTTTACTAGGTACTAATATAGTACGT | - | - | |
| | phiKZ P042 | 81893 | - | ACGCCTATATATTCCGATCCTTATCCAGGTCCTAAAACACTGATAGCTGG | late promoter | Ceyssens et al. (2014) | |
| | phiKZ P043 | 81932 | + | AATTTATAATCATAAAGATAAGGCTGCGTTGTATAGAATTCAATCTTATG | late promoter | Ceyssens et al. (2014) | |
| | phiKZ P044 | 81969 | - | ATACGTAGTGCATAAGATTGAATTCTATACAACGCAGCCTTATCTTTATG | late promoter | Ceyssens et al. (2014) | |
| | phiKZ P045 | 82828 | + | TCACGCTTAACATTCACATCTAACACTTCATTAGAGATTTCTATCGTATG | late promoter | - | |
| | phiKZ P046 | 88779 | + | TGCTTGCAAATCAAATGTGTTAAATTGATACGGGACATCATCTACATATG | - | - | |
| | phiKZ P047 | 89053 | + | ATCATTTAATAGCATTGTGAGCTTTCTATCGATCTTACAATTATCGTATG | late promoter | - | |
| | phiKZ P048 | 90115 | + | TTGATGAAATTAAAATGGCTGAAAAGAATAAAGGTCCGGATATCACTGAG | - | - | |
| | phiKZ P049 | 94330 | + | GAAGTAACAAACATAATGAGGAACCCTTCGGGGTTCCTCTTATGCTATGT | late promoter | - | |
| | phiKZ P050 | 95917 | + | CCAATAAATATATCTAAGTGTTCATTTTTATCTAAGTACCTAATTCTATG | late promoter | - | |
| | phiKZ P051 | 97086 | + | AAGACTATGCCACCTACCTAACGAATATTGGTACTGTGCTTAAAGCTTGG | - | - | |

| ϕ | ID | TSS position | +/- | 50bp upstream region | predicted type | Experimental validation | Note |
|---|---|---|---|---|---|---|---|
| | phiKZ P052 | 98764 | + | ACCTATTTAAATTAACATGTCGTAATTTGACTGCCATACTCAATGTATGC | late promoter | - | |
| | phiKZ P053 | 98803 | + | TCAATGTATGCGATGAAAATCTAAAGTAAGATTCCAGCGCAATATTTTGA | late promoter | - | |
| | phiKZ P054 | 101390 | + | ACGGGGAGCCTTCATGGCTCCCTTTTATGCTGTTAGACCATGATCTTATG | late promoter | - | |
| | phiKZ P055 | 106304 | + | aacagttgacaacaatgttgttaaaacctactacggtgtattattaggtg | - | - | |
| | phiKZ P056 | 106889 | + | TAATTGGAAAGGTGTATTTGAAAGATCTTAAACAGTATATTACACATATG | early promoter | Wicke et al. (2021) | |
| | phiKZ P057 | 108843 | + | TGTTCGTCAATGCTGCAATGACTGAGACAATGGATAGTGCTGTTGCTGAG | late promoter | - | |
| | phiKZ P058 | 114163 | + | TGTCTAGGGTATGATGAGTACGACCCCAGCGATTACAGTAATTATTTATG | late promoter | - | |
| | phiKZ P059 | 114456 | - | CATTGGATAACATTATCGGCTTGCGTTTTCGCAACCGACATGCTATGGGT | late promoter | - | |
| | phiKZ P060 | 114922 | - | AGTATTACTCAATGATACTGGTAATAATATACTAGATTATTAACAATATG | middle promoter | Wicke et al. (2021) | |
| | phiKZ P061 | 116062 | - | ATTTTGTTATGCCGTTTGATCCAGACCGTATTGGTGGTATTAGATGGGAT | - | - | |
| | phiKZ P062 | 117303 | + | TCTTCACGACAACATACTTGTAGTGATTGTAATACCTGTTGGGTAGCTGG | - | - | |
| | phiKZ P063 | 117616 | + | CATTTAGATAGCTAGTAATTTTAGTGAATGTATTTGCTATATTGCTATGT | late promoter | Ceyssens et al. (2014) | |
| | phiKZ P064 | 117754 | + | ATAAAACAGCAGGTGTACAATTTAACAAATTGTTAGACACCAACTATATA | - | - | |
| | phiKZ P065 | 117934 | + | AAATAGACGGTCCTGAACAATTTAAGATGAAGGATTTACACAATCGTTTA | - | - | |
| | phiKZ P066 | 118012 | + | GGGATGCAGGTAACCGTAATTTAGTGTACGATTATTACCTATATCCTGTA | - | - | |
| | phiKZ P067 | 118114 | + | ATCCTAGACGTGTCTCAATTCAACTTAACCATACTTTCTTTGATAGTATG | late promoter | - | |
| | phiKZ P068 | 118184 | + | CCGCCCACGTGAATCCGTGGCGTCAACACAAATTTTTACCTATTATTTGG | - | - | |
| | phiKZ P069 | 119075 | - | ATAGTTAGATTCTGCTCGTTCTTGATGGTGTACGGAGGAACAAGAGCCGG | - | - | |
| | phiKZ P070 | 130258 | + | TCGATAAAGCTCTGTATTTATTTGATCGGCAATATCATCAAATTTTATGA | late promoter | - | |
| | phiKZ P071 | 132129 | + | AACCCTGAACTCCAAGATCGTTTAAGGAGAATGCACATTTCATCTATGGG | late promoter | - | |
| | phiKZ P072 | 134420 | + | ACTAGTGTGTACATCTATTAGTCTCCATGGAATGATTGGACAATCCTATG | late promoter | - | |
| | phiKZ P073 | 134428 | - | GCCCTCCTTTAATAAAACGCTATTGCGCAATTTGTTAGCGTAATGATGTG | - | - | |
| | phiKZ P074 | 142840 | + | TTAATTGGAAAAGGTGAAGGATTAGTATTCCATCTATCCCAATTACATGA | - | - | |
| | phiKZ P075 | 145659 | + | AAAACTATAATAGCCTTCCTCTTCAAGAGGAAGGCTTTATAATGCTATGT | late promoter | - | |
| | phiKZ P076 | 146355 | + | GGCGATAGGGGTGATGAGGTATGTCAACTCCAGACACTCTTAAATTTATG | late promoter | - | |
| | phiKZ P077 | 146766 | + | ATTGAAAATTATGGCATGAAGTATGGCGTACTTACTGATCCAACTGGGGC | - | - | |
| | phiKZ P078 | 147221 | + | TCTTATGTTGTTAAAATATTACTCATTAGCAATAGTACATTACTGTATGA | late promoter | - | |

| ϕ | ID | TSS position | +/ - | 50bp upstream region | predicted type | Experimental validation | Note |
|---|---|---|---|---|---|---|---|
| | phiKZ P079 | 148383 | + | CGATATTGATCTTGGTGCACCCACTACTGAAGGTTATGCTGATATTGTTG | - | - | |
| | phiKZ P080 | 154974 | - | GAATTCAATACCTGTCTATCAAGGGGTATCTCGCTCAAGCTAATGGTATG | late promoter | - | |
| | phiKZ P081 | 155550 | - | ttaaatcaggtgcgtatcaactcgcacctacttggttaatggttatggga | - | - | |
| | phiKZ P082 | 157220 | - | AGGATTGTTGAATATGTAGACATAAAATTACTCTCCAAATCAATTAATTG | middle promoter | Ceyssens et al. (2014) | |
| | phiKZ P083 | 157249 | + | CATTTTTTAAGTTAAACAGTCAATTAATTGATTTGGAGAGTAATTTTATG | late promoter | Ceyssens et al. (2014) | |
| | phiKZ P084 | 159280 | - | TATATAAGGCAAATGAGCCAGATGAGTATCTCCAGTTACTAATTAATATG | late promoter | - | |
| | phiKZ P085 | 161035 | + | ATAATACCAGACTTTTTATTTTAACACAACTAGAACAGACTGATTTTATA | late promoter | - | |
| | phiKZ P086 | 161236 | - | ACTGATACTCAGCTGTAATGTCAACAGGAATTTCGTCCCTTACGCTATGA | late promoter | - | |
| | phiKZ P087 | 164370 | + | GGATCCGCTATTATTTCGTCTTAAAGTAACTATAGTGCTCAATTGAATGA | late promoter | - | |
| | phiKZ P088 | 165041 | + | GCAGAGAATCTTCTCATTGAGAATGGTATTATCTCCTAACAACCTTATGA | late promoter | - | |
| | phiKZ P089 | 171868 | + | AGTTTTCACAACTACACTAAAAAATAATTAAGATATATATTACTACTATG | early promoter | Wicke et al. (2021) | |
| | phiKZ P090 | 172165 | - | ATTGCGTATCTATATTTGCAGATACTCCAGTAATACGATTGATAATATGG | late promoter | - | |
| | phiKZ P091 | 172385 | + | AATTTTTTTATTACACTAAAAAAAGTTTAAGATCTATATTACAGACATGT | early promoter | Wicke et al. (2021) | |
| | phiKZ P092 | 174427 | - | TCCTTTTGGAAGCCCCGGAAATATTGAATAAGTTTCTTAAACATGATATA | - | - | |
| | phiKZ P093 | 179469 | - | GGAAACTTCCCTTGGTATATTAACTATTATGACAAAGCCAAATTATTTGA | late promoter | - | |
| | phiKZ P094 | 181662 | - | CAGCTCCCCATCTCTGAATGGGGAGTTTTTAACTGAGGCTGATATTGCAG | - | - | |
| | phiKZ P095 | 182372 | - | GAGATATGGTGAAAGTTGGGGATCATGTCGACTTAAATTCAATTCTATGT | late promoter | - | |
| | phiKZ P096 | 187788 | - | CGACTCAGTATCCATATCGCCTTCTATTCTCGCGCCATTGGGGGCGGATT | - | - | |
| | phiKZ P097 | 195984 | + | AAGCGGCTAATCCATCTGCACAAACTGGACAAGAGCAAGCTAATACTATA | late promoter | - | |
| | phiKZ P098 | 198121 | + | TTGCTAACACCACTGGTCGCTTCTAAATACTTGGAATAGTTATTTATTGT | middle promoter | Wicke et al. (2021) | |
| | phiKZ P099 | 199232 | + | AGGCTTTATTTCGTATTAAAAAAAATTTGAGATTTATATTACTAGTGTGT | early promoter | Wicke et al. (2021) | |
| | phiKZ P100 | 199463 | + | AGTAGTTGTATCTATACACTCACTTCAAATCATGAGTTAATCGCTGGGGA | - | - | |
| | phiKZ P101 | 200175 | + | ATCGAAACCCACAACCTAAATGAAAAAGTAAGCATGTCTGATGCCGGGGT | - | - | |
| | phiKZ P102 | 203327 | - | TAACCACAGCCACACAGGTTAAATAATTTGCGTGTAATGGTATTGTATGT | late promoter | - | |
| | phiKZ P103 | 206786 | + | AATTAGCCGAGAAGTATATTCGTGCTTGTAGTGCAATAAAACGATGCTGG | - | - | |

| ϕ | ID | TSS position | +/- | 50bp upstream region | predicted type | Experimental validation | Note |
|---|---|---|---|---|---|---|---|
| | phiKZ P104 | 207035 | + | GAGAACCATGATGGTTCTCTCTTTTTGACGTAAGTGCATTATTGTTATGA | late promoter | - | |
| | phiKZ P105 | 210184 | - | TATTATATTCGAGTTGACTCACCTATTAAGACGGCTGATATTCCCGTGTA | - | - | |
| | phiKZ P106 | 211034 | - | CATATGCTTCCCTTTCAACAGCGGTCCGTATGACGATGGCTGATACTATG | late promoter | - | |
| | phiKZ P107 | 212808 | - | GCTCTATATCATTGCACTGTGCGTATTTTTATTCGTTAGCTAATCTTATG | late promoter | Ceyssens et al. (2014) | |
| | phiKZ P108 | 217730 | + | ATATCAGTGTTTGCTCTTTGGAGGTAATTACTCTGGAGCATCTAATATGT | late promoter | - | |
| | phiKZ P109 | 220796 | + | ATAGTGGTTTATACATTTAAAGCATTTTAAACAGTATATCACAAATGTGA | early promoter | Wicke et al. (2021) | |
| | phiKZ P110 | 220879 | + | GTCACGCAATACTATTTCTTCCTGGAGTGAACTTGGTTCAATGCTGGGTG | - | - | |
| | phiKZ P111 | 221507 | + | TAAGACAAAATTAACTCGTGCGGAAATTAACCGATTACAAATACAATATG | middle promoter | Wicke et al. (2021) | |
| | phiKZ P112 | 223862 | + | AGCTGAGAATAACAAAGTATCTGTTATTAATGAGTAAATACAATACTATG | late promoter | | |
| | phiKZ P113 | 224867 | + | ATCTGTAATGAAATAAAATTCAAACGGTACTAAACATACTAACCCTATGA | late promoter | | |
| | phiKZ P114 | 226820 | + | CCATTATGATGAGTATTATGGTTTAGATGTACTGGCATGTGCATCTTATG | middle promoter | Wicke et al. (2021) | |
| | phiKZ P115 | 227508 | + | GTTAAGGTTGTAGAATGATTTAAGGCTATTTTAGTAGCTTTATATTCTGA | late promoter | - | |
| | phiKZ P116 | 231207 | + | TGACTCGTATTGGTCTGTAATAAAAATACTTCTTATAGACTAATAATATA | middle promoter | - | |
| | phiKZ P117 | 231384 | + | gtgcttttgtaattagaccgtttgaaaataccattacgcataatgatacg | late promoter | - | |
| | phiKZ P118 | 232161 | + | AGTAACTGCCCTCATTGGTGCCCCATTCGTAATGTACTTACATTGTATGA | late promoter | - | |
| | phiKZ P119 | 235361 | + | TCTGCACAACTTAATCGTCATCACAATACTCGTATAGTTAATTATTATGT | late promoter | - | |
| | phiKZ P120 | 235469 | + | TGGCGTAAACATAATCCGTTTAACTGCGGTAGGTCTCGTTGTATGCTATG | late promoter | - | |
| | phiKZ P121 | 238722 | + | TGGGCTCCTATTAAAAATTTAATAGAATATTAATTAAAAATTATCTTATG | early promoter | - | |
| | phiKZ P122 | 240373 | + | TTATCTTTTTTTGTATTTAAACTATTTAAAAAGCTATATTACAAATATGA | early promoter | Wicke et al. (2021) | |
| | phiKZ P123 | 243202 | + | TCAAAAGGAAAATAGGATGAAATACGTCTTCTCGCTCATTATCACTATGG | late promoter | - | |
| | phiKZ P124 | 243620 | + | ATGGGTAATCTCTCTTATGGGAATGATTACGTATGTTCTTTATGATACGT | middle promoter | - | |
| | phiKZ P125 | 246721 | + | ATTTTATTTTTTTGTATTTAAAGCATTTTAAACAGTATATTACAAATGTG | early promoter | Wicke et al. (2021) | |
| | phiKZ P126 | 248091 | + | TGGTATCCGACTCAAAGACAAAAATATTATCCAGCTACGTTCTATTATGA | late promoter | - | |

| ϕ | ID | TSS position | +/- | 50bp upstream region | predicted type | Experimental validation | Note |
|---|---|---|---|---|---|---|---|
| | phiKZ P127 | 249790 | + | AAGAAAATTAGTGTAGAATGCTATAAAGGTTTATAATGGTCAATTTTATA | late promoter | - | |
| | phiKZ P128 | 261514 | + | AAATACATGTATTTAATTTTGAAAACATACTGGAGTTATTGATGTATTGA | middle promoter | - | |
| | phiKZ P129 | 262671 | + | ATCTAAAGAGGTTATGTAAAAAATAGTAAAAGAAATACACTATTCTATGT | early promoter | - | |
| | phiKZ P130 | 263172 | + | CAATGGTGATTATACTGCGGGGAACAAATACTTGCAGTAATACCATATGT | middle promoter | - | |
| | phiKZ P131 | 264056 | + | GATCTTACCTGCATTTTCAATGACCGCGAATATGCCAATAACATATGGGG | late promoter | - | |
| | phiKZ P132 | 265938 | - | GTTATCCAGATATTGTAACGGATTAAAATACTTACAGCTATCTAGTATGT | middle promoter | Wicke et al. (2021) | |
| | phiKZ P133 | 266394 | - | CAGTTGGTTATCGTATTACAAAGAAATAATACTCCAGATTATTTAATATG | middle promoter | Wicke et al. (2021) | |
| | phiKZ P134 | 266808 | - | CTTTATTTATTATAATAAATTAAAATAATACTTTCATATACACTATATGT | middle promoter | Wicke et al. (2021) | |
| | phiKZ P135 | 266897 | + | ATAAATAAAGTTATATTCAAACCTATTTAAAAGGTATATTACAAATGTGA | early promoter | Wicke et al. (2021) | |
| | phiKZ P136 | 267851 | - | AGAGTGGTCTATTTATAAGTAAAATTATACATAGAGTTAATATTATTTGT | middle promoter | - | |
| | phiKZ P137 | 269756 | - | TACATAATGTCATAACAAGAAATAAATAATACTCAGATTGTAATAATATG | middle promoter | Wicke et al. (2021) | |
| | phiKZ P138 | 270595 | - | ATAATATTTATAACTATTTAGATAATTATACTGGCACGATGATATTATGT | middle promoter | - | |
| | phiKZ P139 | 271686 | - | TAAAGAGTAGCTTCGGCTACTCTTTTATGTTGCCATGGGTTATCTTATGA | late promoter | - | |
| | phiKZ P140 | 271922 | - | ACTGGACTGATCGCCCGTAATAAAATATTACTGGTTATGTAATACTATGT | middle promoter | Wicke et al. (2021) | |
| | phiKZ P141 | 273454 | - | GTTGAATTTAAAATAAGTTGTAATAAAATACCTAGCATTACATGTTATGT | middle promoter | Wicke et al. (2021) | |
| | phiKZ P142 | 273633 | - | TCCTAAACTCATAAAAGAGAAATTAACAATTGATCTAATGAATATTTATG | late promoter | - | |
| | phiKZ P143 | 273977 | - | CTTCGGGGAGGCTTTATATTTTGTTTTTACGTATGTATATTAAAATATGT | late promoter | Wicke et al. (2021) | |
| | phiKZ P144 | 274148 | - | AGGTTAGCAATGGGTAATGAAAACTACCAGTAATGTATCTAATATTATGT | late promoter | - | |
| | phiKZ P145 | 276905 | - | tgggaaattgtaaagactgaagttacctggattaattatctattctctga | late promoter | - | |
| | phiKZ P146 | 277797 | - | AATTGGGAGGGTGTTATTCCCAATTTATACTAACCTCTATTATTTATTGT | middle promoter | Wicke et al. (2021) | |

| ϕ | ID | TSS position | +/- | 50bp upstream region | predicted type | Experimental validation | Note |
|---|---|---|---|---|---|---|---|
| | phiKZ P147 | 278754 | - | CACGGGCGCGGTGAGATTGCTACTTGTTCACTGGCTGCTATTGCAGTGGA | - | - | |
| | phiKZ P148 | 280201 | - | AATTCATTTAAAATGGGTCGTAAAAATACTTAGTTGATCTAAAGATATGT | middle promoter | Wicke et al. (2021) | |
| | phiKZ P149 | 280271 | - | TAAGGCTACAGAGGTTTTTAAGATAAAGATATCCAATGTACCATAATATG | middle promoter | - | |
| YuA | YuA P001 | 40 | + | AATTTGCCAAGTTGTCTTGGGTTCCGTAAAATACGGCCCA | sigma70 | - | |
| | YuA P002 | 612 | - | TTCTTCTTCTTCTTAACTTACTTAACTTACTTACCTTAAACTCCCGGACG | phage promoter | - | |
| | YuA P003 | 814 | + | CATGATTCTCCGCTCGCCTTCGCCGCCCCGCTGGGCTATACTTCCGGGCA | phage promoter | - | |
| | YuA P004 | 1715 | - | GCGTCCCGCTCAATCCCGCCCGTCTCCAGCCACACGATATACTCATCACG | phage promoter | - | |
| | YuA P005 | 2349 | + | GACGGGTCTGACCTTGACCGCAGCCAAGACTGTCATTTATTATTCCAACA | sigma70 | - | |
| | YuA P006 | 2914 | + | TGCCTCGGGCCTCTTTACTTCTTCCGATTGACCGGCTATACTTCTTCAGC | phage promoter | - | |
| | YuA P007 | 5809 | + | ACCTCGTTATGATGTAGCCCAAGAACAGGAGCAACGATATGATAGAAAGC | sigma70 | - | |
| | YuA P008 | 7758 | + | GTTGCCCGCCCCACTTTACTTCTTCGGATTGTGGCACTATACTTCTTTCA | phage promoter | Ceyssens et al. (2008b), no activity detected | |
| | YuA P009 | 14010 | + | CTGGAAGAAGGCCCAGGACGATCTGGACGCGGAGAAGTCCAAGGCGAAGT | - | - | |
| | YuA P010 | 22676 | + | TGACCTGAACCGCTTTACTTCTTCGGATTGACCCACTATACTTCTTCACA | phage promoter | Ceyssens et al. (2008b), no activity detected | |
| | YuA P011 | 23521 | - | AGGCCCAGACATCCAGGAACTTTTCGCCCCGGACCACAATCACAGTTTCG | - | | |
| | YuA P012 | 29094 | + | TTGTATGAGGGGGCTTTACTTCTTCGGAGGCCCCCACTATACTTTCTTCA | phage promoter | Ceyssens et al. (2008b), no activity detected | |
| | YuA P013 | 31134 | + | ACGCCCGGCGGGGCTTTACTTCTTCGGGTCCCGCCACTATACTTTCTTCA | phage promoter | Ceyssens et al. (2008b), no activity detected | |
| | YuA P014 | 33417 | + | CGCGACCCTCGCCCTTGGCTAATGTACAACCTCACGCTATACTCGCGGCA | sigma70 | Ceyssens et al. (2008b) | |
| | YuA P015 | 38492 | + | GCAAGGCTGGCCTTGACCCTGAAAAGTTCTGAGGCTACAATATGAACCGT | sigma54 | Ceyssens et al. (2008b) | |
| | YuA P016 | 39396 | + | TTTGGCCAGGGGGCTTCCCTTTTTCGCAACTTGGTACTATACTGACGTCA | phage promoter | - | |
| | YuA P017 | 44790 | + | CGGGGGACAAGGCTTGTCTCTTGCTGAAAAATGCGCGATAATCGCCCGGG | sigma70 | - | |
| | YuA P018 | 46497 | + | CAATGAATTCCCTTACCTTCCGAACGCGGCGGACGTATAATCAACGGAGC | sigma70 | - | |

| ϕ | ID | TSS position | +/- | 50bp upstream region | predicted type | Experimental validation | Note |
|---|---|---|---|---|---|---|---|
| | YuA P019 | 47574 | + | GCCGCAATTGACCAAGCTGAGTCGGTCATTCGCGGGTACAATTATGGGGA | sigma70 | - | |
| | YuA P020 | 52854 | + | ATGTTCCAGGCTTGCCAAGGTGGTGGGGTTCACGATAAAGTATTGCGGGA | sigma70 | - | |
| | YuA P021 | 57837 | + | CTACGCTCCCGGGAACTTCCGGGTGAACAGTCTCCAGTATCCTTCGGCTA | - | - | |

**Supplementary Table S4: Overview of phage transcription termination sites and associated terminator sequences, as identified by ONT-cappable-seq data analysis.**

| ϕ | ID | TTS position | +/- | -60 to +40 sequence region | predicted type | ARNold hairpin energy (kcal/mol) |
|---|---|---|---|---|---|---|
| 14-1 | 14-1 T001 | 178 | - | GGCTGAGCACGGACGAAGGCGAAGAGTAATCATCAGGCGCGGCCGGTTGGTCGCGCCATTCACAGGAGAAT<br>TGTCGTGTCCAAGCTAATCCCTTTCAACA | - | - |
| | 14-1 T002 | 3581 | - | AAAGTTATGCTGCGAATCGTCCAATCCATCCTCGCGATCCTCATCGGGATCGCTTGGGCCTGCGCCGACTA<br>TGGAGTCGTGGCCGCCGAACTTCCTCCAT | - | - |
| | 14-1 T003 | 5619 | - | TCGGATGATCTACATAACCATCAAAGCCATGGCCTGGTTCGCCCTCCTATGGGCGACCGGCTTGGCCATCG<br>TAACTCTCACCATACATTTATGCTACTGA | - | - |
| | 14-1 T004 | 6661 | - | ATCCGTGTAGCCTGACCCTCTGCGATCCGACGAACGGTAGTGCGAAAGTGCTACCGTTTTCCATTTCTGGC<br>GTATAGTCAACCCATCGAAACGAACTTTC | intrinsic | -9.3 |
| | 14-1 T005 | 35525 | - | ATGCCGGAGTCTTCGGCGAACAGGGAATAGTTCGACTTCCCGGTCAGCTGGATCAAGCCGCGCCCGCGATA<br>CTTCCAACCATCGCCATCCTGCTCGCATC | - | - |
| | 14-1 T006 | 35820 | - | AAGATGGTTGAAGAGGCAAAGCCAGAAAGCAAAAAGGCCGCTGAATAAGCGGCCTTTTCTTTGGGTGGAAT<br>GTCAGGAGCCTTGATTGATCGCGGCGATG | intrinsic | -16 |
| | 14-1 T007 | 37289 | - | ACGCCAACGAAACCATCTGAGCCGGCTTACCGCCATTTAGCCCGGTTCGCCGGGCTTCTTTTCTGGAGAAC<br>TGTATGAAACCCATGCTCGCATCGAATTT | intrinsic | -12.2 |
| | 14-1 T008 | 38499 | - | CCGAAGCCGAAGAGTTCTAAATCACCGGCCGGAGAAAGGCGCTGTAATGGCGCCTTTCTTTTGAGGAGTCG<br>AAATGGACTGGAAAGACATAGGTAGTAAG | intrinsic | -11.1 |
| | 14-1 T009 | 50323 | - | CGCTCTTCTGAGACTGAATCGTCTAGCTCTTCAAGGGACTCATGAAAATGGGTCCCTTTTTTATCCTCTCA<br>AGTTTTGTACACTCAGAAGGAACACATCA | intrinsic | -12.6 |
| | 14-1 T010 | 51089 | - | CCAAGGCCAAGGAAGCCGAACAGAAGAAAGCCGAGCGCGAAGAGAAGCGCAAGGCCGAGCGTGAGAAGAAG<br>GAAGCAGAGCGCGCCGAGAAGGCGAAGGA | - | - |
| | 14-1 T011 | 51354 | - | AGTTCCCTGGAGTCTGGACTTTCCCGAAAGAAAAATATTTAGAGAAGTTCAGGCGAAAGGTATTCCCTTCT<br>TCAATTAGTTTAGAGATAATCCTCTCACA | - | - |
| | 14-1 T012 | 53335 | + | CGAACCTTAAGCCCGGTTTCCTGCGTCGCAGCCGCGAGCTGGACCCGGCGCTGGCCGTTCGCATCCGTCGC<br>GAGCTGATCCACGCCGAAGCATCCGACTT | - | - |
| | 14-1 T013 | 60583 | - | TGGAGAATAAGAAACCCAAAAGAAAGCCCCGGATTCTGAACTAGAGTCCGGGGCTTTGTCGCTTCTGATCG<br>CTATTGGTCGGCTTCCTAGAACAGCCCTA | intrinsic | -15.9 |
| | 14-1 T014 | 62469 | - | AGTCGTTCTGTTGAAGCGCGACTAACAACAACCAGCCTCTGCGACTCGGCAGAGGCTTTCTCATTGGAGGG<br>CAGCTTATGTCCGATAAAGAACAAAGTGG | - | - |
| LUZ19 | LUZ19 T001 | 141 | - | AGGGGTTGACACTCCCGACAGGATGCGTAGAATGGGCGGCGACAGGGCAGGGAAGCCCACCAAGCCGACAG<br>GCGAAGCACCGCACCACCGGCGAAGCCGG | - | - |
| | LUZ19 T002 | 457 | - | GGGCCAGGAAGGCACAAGAGGTAAGGCTGAGCATAGAGGCAAGGCTAGCCACTCCCGCACCACCACCCGAC<br>AGGACCAGCAGAGGCCACAGGAGGCACAG | - | - |

| ϕ | ID | TTS position | +/- | -60 to +40 sequence region | predicted type | ARNold hairpin energy (kcal/mol) |
|---|---|---|---|---|---|---|
| | LUZ19 T003 | 1704 | + | GCAGGACAAGCCAGAACCGCTGATACTACAGGGACTATGCCAATGCCAAGGTTTGTCCCTTGAGGCCCTTC<br>CACCGAGGGGATTCAAGAGACAGACCTAG | - | - |
| | LUZ19 T004 | 1768 | + | GCCCTTCCACCGAGGGGATTCAAGAGACAGACCTAGTGAGGATTGCCGATGGCACACTTCAAGGCTAAGGC<br>TCCCAAGTCGCCCTTTGCTGCTCAGGTAG | - | - |
| | LUZ19 T005 | 2852 | + | AGAGCAAGCGTGCTGCCGCCCATTGACTCAAGTCGAACGCCTGCTAAGCGGGCGTTCCGCTGGAATCAATG<br>ACAACTGGAGAAACACGATGAGCTTCAAA | - | - |
| | LUZ19 T006 | 3156 | + | TGAAGCGGTGACTCAAGTCATGGCCCTGGCGGGCGACCCTCGCCTACTCCGGGGCCATCGCTGGACTCATC<br>ACAAGCGAGAACTAAACCATGCAAGCTTT | - | - |
| | LUZ19 T007 | 5048 | + | CACCCAAGTGACACTCACCGCCGAGTGACCAAGGCGAAGGCTGGTGCGCCAGCCTTCCACCGTGGCCATTC<br>CTTGCCGCGAACCAACTCAACCGAGGAGC | - | - |
| | LUZ19 T008 | 18088 | + | CAAGCTTCCCATGCACCTGGTAGTCGAGGGGTTCCCGCTGCTGAAGCGGGAGCTTGCTCGAATGATGCAAT<br>GGGCTGCCGAGGTCAAGGGGTATCTGCCG | - | - |
| | LUZ19 T009 | 21207 | + | GTCCAAGGCCCTCGTAGAGGGAGCGGGGGAGAGGAGAGGTCAGGGAAGACCTGGTAGAGGAGAGGTGAAGA<br>TGAGAATGGATGACTACGAAGGATTCTAG | - | - |
| | LUZ19 T010 | 21415 | + | CGGCCCGTCCTGTCACGTTCACCTCTCAGCAGATCGAGTGGTTAGAACAGACCTTCCCCGAACATCAGATC<br>GGTCCTGGAACCACGATGGAAGACATCCA | - | - |
| | LUZ19 T011 | 22355 | + | CGGGACGGGAGCGTGGAGAGTCGAGCCATCGAGTTCGCCAAGACCACGCTGCCCTATCTGATGGTCGATCC<br>CATGTCCGGCAGCCGGGGAGTCGTAGAGC | - | - |
| | LUZ19 T012 | 22491 | + | ACCTCGCCGCCAAGCTGGCGAGATCGCTGTTCCCCACGGGGATTCCGTTCTTCCGATCCGAACTCACTGAT<br>GCGATCCGCCGCGAGGCCGACAGCCGGGA | - | - |
| | LUZ19 T013 | 22582 | + | CAGCCGGGACACAGACATTACCGAAGTGACCGCTGCCTTGGCTCGGGTGGATCGCAAAGCAACACAGCGCC<br>TGTTCCAGAACGCCTCCCTGGCGGTCCTG | - | - |
| | LUZ19 T014 | 25861 | + | GCGCCTTCGACATCACCGCGTGATGCCACGAAACCCCGCACTTCGGTGTGGGGTTTCTTCAAAGCCTAACG<br>ACCCGCGCAGATTCCCTGCGTGGGTTTTT | intrinsic | -13 |
| | LUZ19 T015 | 32528 | + | GGGCGATGCGCTCCAGCTCGACGCAGACCTTGCGGCGAACAGTACCGGCGAGGTGGACCCTAACTTCGACG<br>CTGGGACCTATGGGGTCCAGGCGCTCCAG | - | - |
| | LUZ19 T016 | 36296 | + | CCGAGACCATCCACGTTGCAGATGGGGTCGAGGCTGTCTTCAGTCTCGACTTCCCGTTCCTGCGGCGTGAG<br>GACGTATTCGTCCAGGTCGATAAGATACT | - | - |
| | LUZ19 T017 | 38768 | + | AGGTTGACTATGCCAAGGTGGCCCTGGACGACAAACGTCCGGGGGCTACCGCCTGGGCGCAGGTTATCGTG<br>CCCTACACACGCAACGGGAACCTCTACGT | - | - |
| | LUZ19 T018 | 43099 | - | CAGGCAACATCCAGGCAAGGCAAGGATGGCCCACCAGGCGGAGGGCCTACGGGGGAAGGGTGGCTGATCA<br>GAGTCGGGAGGGTTCTCCCACGACGATAT | - | - |

| ϕ | ID | TTS position | +/- | -60 to +40 sequence region | predicted type | ARNold hairpin energy (kcal/mol) |
|---|---|---|---|---|---|---|
| LUZ24 | LUZ24 T001 | 998 | + | CCCTACGATAGCCATCCGGCTCCCTCTTACAAGTAACCTCGGAGTTAGGTAGGATATATTACAAAGGAATA ACCCTTTGCAATGTATCTTCTAGAGGGAC | - | - |
| | LUZ24 T002 | 1872 | + | CCTCACCAGCAGCGTATCCATCGATATCTGTCCATGGTAAACCGTGGGCAGGTATCGAGGGCGAAGCGATA CGCAAATAGGCATTGCCTGTAATGGCCAG | - | - |
| | LUZ24 T003 | 1941 | + | TACGCAAATAGGCATTGCCTGTAATGGCCAGGGGCGTACTATGACGCGCCCCTTGGTTCCTCCCTAGAGTA CATGGTAGGTCCGTGTATTGTATGGATGA | - | - |
| | LUZ24 T004 | 2480 | + | AGGCCATGGTGATTCCTGGCCGAAACCCCCACCGGACCTATGGTTGCAGGCTGGGGCGTCTTGGGAAATCA ACTAAGGAAACCATCCCGTGAAACGCAAT | - | - |
| | LUZ24 T005 | 4067 | + | TTTCACCTTGGCAGCCGTAGGTGTGGAGTCGAGCCGATCGGCTTACCTGCGGCACGCCAGGGAGGCTATGA TCCAATCCGGGGAGGCTTGCCCCCATTGT | - | - |
| | LUZ24 T006 | 5415 | + | GTCGCGGCCCCTTCTAAGAGGGCGTTAGTAATGCCCCTAGGAACTCCCTGGGGGTATTGCTGGCTAACTCA ACCAAGGAGAAAACAAATGGCCCGTATCA | - | - |
| | LUZ24 T007 | 7605 | + | CGGCCAGCACCAAGGAAGAGTTCGTGAAGCGTATCGTTTCCGTTCGACTCCTGGGTGCCCAGATGCTCCAC CCCTTCGGCATGAACATCATCGAGAACCT | - | - |
| | LUZ24 T008 | 8164 | + | GCGAATCATCAACGATAACGACGGCGCTGCGGCGGTAGCTGCACTTCAAGCGCTGGGGGTAAAGTATGAAT GACCTGAATAATCGCCATCGGTTGGCCGG | - | - |
| | LUZ24 T009 | 8922 | + | TGAAGGGTGACGGCTCTCTGCGGGATGGTAAGGAGTATGTCTTCTCTGGTCCCGCCTCTGGCCAGAAGGCA ATCACTCGGGTTGAATCCTTTGTGAGTGC | - | - |
| | LUZ24 T010 | 10631 | + | CGTTCACGGAGGCCGGAGTGCGAAAGGGTGCAGAACCCCAGACCACTCCGGTCACGTTTCTGGATCTCCCG ATTCCCGAAAGCGAAAGGCAAGCAGCGGC | - | - |
| | LUZ24 T011 | 11755 | + | CCCATGCCGGTTCCTAAGATGCGTACATGGGATCTAACTTAGAATTCCAGGGGCTATTGCTGGCTTCACTA CCCTCAACAGAAACAGGAGATTTGCCATG | - | - |
| | LUZ24 T012 | 15963 | + | AAGGCCAAAGGCATTCGCAAGAAAAAGTAAATGAAGAGGGGCCTTCGGGCCCCCGAGGACTCACTATGTTC AATCGAAAGCTTAGCATCAGTAACATCCT | - | - |
| | LUZ24 T013 | 17476 | + | CTCTCCGGGAGAAGCGTCGGGTATTCCTGAGCCTCCCCGCTGGATTCCTGGGGGTGAGCTACTTCTCAGGC TCCCACCTGAATCTGAGTTCCTACTCGCC | - | - |
| | LUZ24 T014 | 18243 | + | GGCCTTCAGCAAGCCATTCCTAAAGAGTGGTAAGCCGAACCAAAGGCTCCAGTCCCTGTGGCAACGTCTTG GGCACTTCGAGGTATCTGGTCCCTTCTCT | - | - |
| | LUZ24 T015 | 19270 | - | GCACTTTCTACAGTTCTTAATCTTAAACCATCCATTTGGGTACTTCATAAACACCTCCCATAAAATATCAG TATAACACATTCTACGGGAAATGTCAAGT | - | - |
| | LUZ24 T016 | 19811 | + | GGGGAAACCTTGGGGCAGCCGAGAGGCGGGCCAGTAGTAGCGACACTGGTTGAACTTTTGATGCGTTCATC TACGGTGCTGGGGATGCCAAGATTGGGAC | - | - |
| | LUZ24 T017 | 20338 | + | GGTCGCTCGTGGTATGAAACTCATTGATACATCTGGGGGTTGACTTTCAGCCCCCTCTGTGGTATAATACC TTCTTCCCTACGAGAGGTTTAAGATATGT | - | - |

| ϕ | ID | TTS position | + /- | -60 to +40 sequence region | predicted type | ARNold hairpin energy (kcal/mol) |
|---|---|---|---|---|---|---|
| | LUZ24 T018 | 20978 | + | CTCCTATCTAAGGGGTTGGAGGATCACGAGGCGGGGGTATGCCCACGGGGCTGCCCCTACTGCTTAATCGA<br>ATTCGAAAGAGTGTGGGGTGTAAGGGTGG | - | - |
| | LUZ24 T019 | 21477 | + | CCTAGTAACTACTCTGTAATGGGCATCGATGTAGACCTGGGGCTTCCCCCAGGCTACAGCCTAACGGAGGA<br>AGCTATGGACAAGGCCAAGCGTCAAGAAA | - | - |
| | LUZ24 T020 | 25726 | - | TGAACGCTCTACGGAGCTAAAGATAAGGCCCCTTGGGATGATACCCTTGGGGCCTTTTTTTTGGTTACTCG<br>ATGGATACCTACAGTGAACCTGCCCAAGA | intrinsic | NA |
| | LUZ24 T021 | 29129 | + | AAGCCTTGGCGAGTGGCTCTAGGAGCGACGTAGCTTCGTTTCCTAGGGCATCCCAGGCTTTCTTGACCGTA<br>CCCCTAGGCTCAAGCTCAGAAGCCTTAGG | - | - |
| | LUZ24 T022 | 32153 | - | AGGTGGTGGGCAGGAGCGGGTTCAGTTGAGTCGTCAAGGCTCCTACCGCCCCACCCACCTTAGGACAATGC<br>AACCGGGAGAGTCTAGTGGACAATCAAAC | - | - |
| | LUZ24 T023 | 37885 | - | TCTGATCCGTGACGTCGTAGCCCAGGTCTAACCCAACGGGGCCCTTCGGGGCCCTTGTTTCCCATTAGGAG<br>GTACATCATGAGTATCCAATCGACTTATG | - | - |
| | LUZ24 T024 | 38968 | - | ACCTGCCGCAGTTAAACCTGCTGCTCGTAGGGCGGAAGCCTCAGAGGGTTCTAAGCGGGTTAAAGCTGCCC<br>GCGCTAATCTGAGGAAAGACCAGTCGGTT | - | - |
| | LUZ24 T025 | 43329 | - | TCCGAACTCCTAGGAGACTGGAAGAACCCTGAGGCTTTTGGAACGGGGATGATCCCCAAGGAAGATATCGT<br>AGAGACGATTCGTAGGGAAGGTAAACCCG | - | - |
| | LUZ24 T026 | 45443 | - | GGCTAGGGGCTATACCCCTGGGGAGAGGGCCACCCCCACTTTATCTGGGGGACTCCTGGGGGGATATTCCC<br>CAGATACACCGGAGGATAAAATAGACTTT | - | - |
| PAK_P3 | PAK_P3 T001 | 2627 | - | CCTTGGAGTAAACATCCAAGTCAATCTCGTCGAGGTCGCCGAACTCGAAGGTGGGGAACTCTTCATCGGGT<br>CGATATCCGTTTAACCGGAACAGTTGAGG | - | - |
| | PAK_P3 T002 | 5673 | - | TAGGGCATATCCTACTCCTTTTCTCGGGACCAAAAAAGGGGGCCCGAAGGCCCCCGTTTTCTTAGGCCGCT<br>TTGCCTTTCACGACCAGTTGCGGGTACAT | intrinsic | -16.7 |
| | PAK_P3 T003 | 9540 | + | AACCTCGACGCCCGTTGGCTGTAAGCGACAACCTGGCCGCCCCTCGTGGGCGGCCTTTTTTGTACCTGGAG<br>GAAATATGAACACCTTGACTTACTCTCCA | intrinsic | -18.7 |
| | PAK_P3 T004 | 10211 | + | CTGGGACATGGAGGTCTACCGTAAAGTGGTGGGAATCCGGGGTCAGGTGACCCGAGTTCGAAACCTCAACC<br>AATCTGCACGTATGCGAGTAGAGCTTCTT | - | - |
| | PAK_P3 T005 | 11269 | + | ATTACAAAGCTCGTCGGGACGATCCAGTGGAGGGCGGATAACCGCCCCCTGCTGACGTTCAAGAAGAACCT<br>GGAACAGGTCAGAGACCGCCTCAACGAAG | - | - |
| | PAK_P3 T006 | 13706 | + | TACCGACCATGTGATTGCTCAGAATCAGAAGATGACCCTTTCGGGGGTCGTCTCTGACGCTGACTTCAACC<br>TGGACAGGCCAGTTATTACGGCTGCTGAC | - | - |
| | PAK_P3 T007 | 13778 | + | GGACAGGCCAGTTATTACGGCTGCTGACGCACAGGCTTTCGGGATCACCCCTAAGCCTTTTGTCAATAGTA<br>CGCCAGTGACATCGGGCCCCATCATCACC | - | - |
| | PAK_P3 T008 | 18374 | + | GCAGTAGGGGCTCAATTGGACGTTCTGGGGGAGATTGTAGGTCTTCCCCGGAGCCTTGTCACTGCCGAGAT<br>ATTCAAATTCTTCGGCTACAAAGATCGTC | - | - |

| φ | ID | TTS position | + /- | -60 to +40 sequence region | predicted type | ARNold hairpin energy (kcal/mol) |
|---|---|---|---|---|---|---|
| | PAK_P3 T009 | 23748 | + | CGGATAGCGTTGGTAGTCCTCGTTCTGGGGCTCTCCTGGCTTCTCCGGGAGAGTCATCAGAAAATCGACAG<br>GCTGGAGCAGAGCCTGGCTTCTCTGGAGG | - | - |
| | PAK_P3 T010 | 24869 | + | CAGGACAACCTCAAGCGGAAAAGATGAGCCCCACTGGCCACAAGCTGGTGGGGCTTTTTCACGCCCGGAGA<br>AAAGTAGTTGACATCCCCCACAACACCCT | intrinsic | -14.4 |
| | PAK_P3 T011 | 26024 | + | GAAGTAGCTGTTATGATTCCAGACCTCAAGGAGGCTTACCAGAAAGCCTCCAAGAGGACTAAGCTGTTGAT<br>ACTGGCTGCCCTGGTTAGCCAAGCAGCGA | - | - |
| | PAK_P3 T012 | 27936 | + | CGGCTTCCAGAACCGCGTCTACAAGTGACCAACAGGGGCCAGCCAGGCTGGCCCCATTTCTTTTGAAGGAG<br>GATGCATGGAACATAAAGCGTTGGAAGGT | intrinsic | -17.7 |
| | PAK_P3 T013 | 31308 | + | TCCCGAGCCACCACCTCTCCTGCTGGCGTAACCCTAAGGGGCCCTGATTGGGCCCTTTTTATTGCAGAAGG<br>AGTAGGAAGTGCTGATTGGGAAGTTCGCC | intrinsic | NA |
| | PAK_P3 T014 | 31491 | - | TCTTTCTCCTCTTTCCTCCTCTGGGCATGGGCACGTCGATCACCTTTGATGATCACTGCTCCCAACTCTTC<br>CCCCAGGGGAGACCTTGCCAGGGTGTTGT | - | - |
| | PAK_P3 T015 | 33396 | + | ATCGAGGCTACCGGGCTGATCGGGGAGGGGACGGTGGATTACACCGCCTCACCCTGGCGCCTGAAAGAGGA<br>TTTCAAGCTACACTGTGCTGTAGCTCGGG | - | - |
| | PAK_P3 T016 | 36540 | - | CTTTTGGGACGCAGTGCCATGTTTTACTTCTCCTATCAATTACCCTATCGGGCGGTAGTTGCCCCTAATCG<br>GGGGGTTTGTCGCTTGTTAGCGAACTTCT | - | - |
| | PAK_P3 T017 | 37510 | + | TTCGACGACGATATTCCGTTCTAAGACGGAACCCTCAGGGCCCTTCGGGGCCCTTTCTTTTACTCCAAAGG<br>AGGAGATGAGAATGGAAGATTACGTGGTG | | - |
| | PAK_P3 T018 | 41420 | + | GACCGGGAACACCTGGTAGTGGGCATCTGACACAAAGCCCACCTTCGGGTGGGCTCTTTCATATCTACAAG<br>GAGAATTCACGCATGATCATGAGCGACCT | intrinsic | -16.2 |
| | PAK_P3 T019 | 45611 | + | TGTGGCCGCATGAAATCTGGACTGGGCAGTAAAACAGAGGGGCCTTCGGGCCCCTTCTTTGAAAGGAGAGA<br>TTGTGAGCGTATTTGTTGAAAGCACCAGT | intrinsic | NA |
| | PAK_P3 T020 | 47943 | + | GATAAGTACGGCCAAAGCTGTTGACAGGTTGTATGGGGCCGGGTAAACTGGCCCTATTTCAACCGGGAGGT<br>AAAGATGAACGATCAAAACTACGATTTTC | - | - |
| | PAK_P3 T021 | 50279 | + | TATTTGACTGGCCCCCTATTTCGGGCTAGATATTTGAAGGGGCCTCCCAGCCCCGCCCTTACATTTTAAGC<br>CGGCCCCTGACTCACGGCCTGGGTTAGGG | - | - |
| | PAK_P3 T022 | 50625 | + | CCTGACAGATTTTCACAGCCGGGCGTAAGGTACTCGACAGATTTTCACCGAGCGCCCAAGGTACTCGACAG<br>TTTTTTACAGAGACAGGTATCAAAAGAAC | - | - |
| | PAK_P3 T023 | 52047 | + | ATACGTCAAGTAACAGCAGAGTGCTTTTAGGGTTGCCCCTGGACGCCGGGGGCAGCTTGCCTAAATCCACT<br>CCGTAGGAGATAACACGATGAACACTGTT | - | - |
| | PAK_P3 T024 | 53146 | + | AAGAAGAGGAAGAATAAGGGTTGACAAGGGGCCAGGAAACTCATAAACTGGCCCCAACACCAACGCACTGA<br>GGGTAACACAACATGGTCAAGTTCGAAGA | - | - |
| | PAK_P3 T025 | 53749 | + | AGGACTAACCTGGCGCCTGGGGCGCCTTCCCTAAACCCTGGACATTCTCCAGGGTTTACGAAAGATGCTCT<br>AGATGTTACCCTGCAACCCCACATTTGAG | - | - |

| ϕ | ID | TTS position | + /- | -60 to +40 sequence region | predicted type | ARNold hairpin energy (kcal/mol) |
|---|---|---|---|---|---|---|
| | PAK_P3 T026 | 54353 | + | GTGGCTGTTCAAATAAAAGATCAAAAAGGGCTTGACGTGCGCAACGGATGGCCCTAGTATCTCTCTCAGAA ACACAACCCCTTACTTACAAGGTAGATCG | - | - |
| | PAK_P3 T027 | 55204 | + | GTCGATGACTCCCTGGACTACTACAGTCGATAACCCAAGCCCCGCTCCTGCGGGGCTTTTGTTTGCCTGAA ATAAAGTGTTGACACTATAGGCCCTTCCT | intrinsic | NA |
| | PAK_P3 T028 | 58608 | + | ACCATCCGAAGTCTGGTAGAGCATTACTGAACACAAGCCCCCATTACGGGGGCTTTCTTTTTCCTACAGGG CTTGACCTGGCCCCCTCTTCCTGTAGAAT | intrinsic | NA |
| | PAK_P3 T029 | 59378 | + | TCTGAAAAACATCTTGACGGGGGTCCCAGGCTTCGGCATAATGACCCCCGTCAAGCAAACAAACCCCGGAG ATTCCCATGAACACCTTCGCATACGCCCA | - | - |
| | PAK_P3 T030 | 59945 | + | AAAAATACGCTGAGATGGGGATTTAATGCTTTACGGGGCCCGGCAAGGGCCCCATAATGCACCCATGCCAA ACAACAGCCCTGGAGGGCAATACAATGAA | - | - |
| | PAK_P3 T031 | 61972 | + | ATCTACGTCCGCAACCATCACAGCTAACCCTTACCAGCCCGGCACAGCCGGGCTTTCTCTCATAGGAGGAA CCATGAAAGTTCAGATCGGTCAGCATACC | - | - |
| | PAK_P3 T032 | 68654 | + | GCAGGCCAGACGTTAACGTGTGCTGGAGGGTTGACACTAGGTAGCCCCTCCGGTACCTTGTCTTTTCTTGA ATTTTAACCGAGGAGATTTTAATGAAAAC | - | - |
| | PAK_P3 T033 | 72389 | + | GGGATTACTTGAAAATCTGACTACATCCAGCCCCAGGACTTGACAGTCCTGGGGCTTTGTCGTATCTTCAT CTCTCTCAACAAATAGGAGGGTACATGAA | - | - |
| | PAK_P3 T034 | 73240 | + | CATCGCATAGCCTCGATGCTATCGAAGTGACCCCAAAGGCAGCTATTTAGCTGCCTTTTTTATTGCCCTTT TTCAGCCCTGCTTTGATCCTCGTCCCACA | intrinsic | NA |
| | PAK_P3 T035 | 73423 | - | CTAGGGTCGAATACCCGGGTTGAACGAAACCCCGCTGACGGAAGTTGGCGGGGTTTTCTTTTGCATGAAAG AAAGATGAAGGGCAGGACCGGTTTGAAAA | intrinsic | -13.6 |
| | PAK_P3 T036 | 78328 | - | GACAAAGAACGCAAGGGGAACAAGAAGGGGCGCCAGTTGGAAAACCAACTGATGCCCTTGTTCTCGTTCAT CGCGTTTGAGAGGGATGATCCCTGGCCGT | - | - |
| | PAK_P3 T037 | 83230 | - | GTCTCCGAAGAAGCGGTGGAAAGCTGATAACCTAAGCCCGCCCAGTGTGGCGGGCTTTTTTGCGCCTGAAA TAAGTTGACAGAGCAGGAGAGTCTGTGTA | intrinsic | NA |
| | PAK_P3 T038 | 83691 | - | AAACCGGCCCGTAAAAAGCGGCAGCCAAAGCCGAAACCTAAGCAGCAGGATGATGCTGAGCAAGAACAAAT TGAAAAAGGAGAGAATGAATGACTGAGAA | - | - |
| | PAK_P3 T039 | 85458 | + | GCTCTAAAGTCCTGGAGAAGTGTCAGCCGGACCTTTGAGGATGGGAGTCCACCGGCACTCTTGACACGGCG TACTGAACACGGTTTAATACCCTCCACGA | - | - |
| | PAK_P3 T040 | 86017 | + | GCTGATGTGCAGAGGAGGTGATCCAGCATCTCCCGGACGGGTCCGGGTTACCAACCCGCTAATTTCATAGC CCGACAGCAAGGGTAGTTCCTGCGGGGCC | - | - |
| | PAK_P3 T041 | 86426 | + | GTGGGGATAGAGGGCTCCCACCGCTGAGCAACAGGAGGTTCGATTCCTCAACACAGCCCCAAAACTGCTCC AGGCAGTTCAGCTAGGCTAGGCTTGAGAG | - | - |
| | PAK_P3 T042 | 86498 | + | GGCAGTTCAGCTAGGCTAGGCTTGAGAGGTAGTAGCCCGTTTTTCCAAAACCTCTCACTTACAATCAGGCG TGTGGCGAAAAGGTTTAACGCACTGGACT | - | - |

| ϕ | ID | TTS position | + /- | -60 to +40 sequence region | predicted type | ARNold hairpin energy (kcal/mol) |
|---|---|---|---|---|---|---|
| | PAK_P3 T043 | 86669 | + | GCTCCGGGTCAGCGACTCAGAAGTGACCCTTAGCCCCGGCGATCTTCGCTGGGGCTTTTTACTTTCTCAAC<br>CCGTAATGGAGGTTGCAAATGACGTTCCA | intrinsic | -16.7 |
| phiKZ | phiKZ T001 | 3756 | + | TAACTAAATAAATTAGTAACACTTATTGGCGACATACACCCTCCCTTCGGGGAGGGCTTTATTTTCAATAT<br>AGACATATTTTTTTAATACATTAGTTGGA | - | - |
| | phiKZ T002 | 5540 | + | AAAATAGTGGTAGGTTTTCTTGCAGTTCTAATAGAACTCACTACTGCTGGTAGTGAGTTCTATTATGAACT<br>GACCAAACCCGAAGGTAAGGTTTCGAAGG | - | - |
| | phiKZ T003 | 12011 | - | ACAGTGTAGTCGCGAATGCACTGTTTGGAACCCCATTGGTTTAGCGGCCAATGGGGAATTCCTTTTATGTT<br>GTCTCGACCTTCTTTTTAGTAGGCTTTTT | - | - |
| | phiKZ T004 | 12168 | - | GAAAATGTCAAGCTTAAACTTGACGACATATAGCCCTCCTGTTGCACCAGGAGGGCTAATCTTTTTCAGAA<br>CTTACGGTATTGTTATAATACTGCTAGCG | - | - |
| | phiKZ T005 | 12586 | - | GCTCGTGCAGAGCGTACACGTAGAGTGCGTATTAACAGATATTCTAGAAATAGTTGGTATTAAACAAAAAA<br>AATAATAATACGTTACCCTCTTCGCAATG | - | - |
| | phiKZ T006 | 14840 | - | GCATTACCTTGTTTACAACGTTAAATATAAAGCCCCCTAGGTGGTTCCTAGGGGGCTTTATAACGTCTTTT<br>AACACATTATCCATATGAGACTATACTTT | - | - |
| | phiKZ T007 | 19111 | - | GACCGACAGTTTGTTTATGAACTGAAACAGCTTACCCCACGCTACGGCGTGAGGGTAAACTGGGTGGTTGA<br>TTACACCCGGTATAAGCTACCAGTCCAAA | - | - |
| | phiKZ T008 | 20033 | + | GACAATTTGAGTAAATTTCTTAATATCCTCAAAATCATCAACTGGGGTAATCTGTCCACTATGCTCTAGCG<br>TTTCACCATTACGTGCATCAACAGGTGGA | - | - |
| | phiKZ T009 | 20489 | - | TACTACGTGGATCAAGCGATGCTTGACCATAGATGGATGCAAAACCATCATCAGAATCACCGACATAAGTG<br>GTTTCAGTCGGTCCTGTCTCAGTGAAGAG | - | - |
| | phiKZ T010 | 22435 | + | TTCAATCTGGAAACTGTTCGTATTGAAACCCCACCTGCTCAGCAATGAGTGGGTGGCTTTCAGAATGAGGA<br>AAAGTTAAATGGCTCGCTATAATGATCCG | - | - |
| | phiKZ T011 | 23552 | + | CTGTCGGTACTAACCAAAAAAGCTAAAGCATATCGCCTCCCTTCGGGGAGGCTTTATGTTGTTAAAAACTA<br>TGCTAAAAAATAATTAAGATATATATTAC | intrinsic | NA |
| | phiKZ T012 | 25208 | - | AAGAATTACTATACAAAAACAAAATAAACATAATGCCCTCCCGCAATGGGAGGGCTTATGCTGTTAAGTAA<br>CATTAGCTAGATTGATCATTACATAAATT | intrinsic | -18.1 |
| | phiKZ T013 | 25242 | + | TCAATCTAGCTAATGTTACTTAACAGCATAAGCCCTCCCATTGCGGGAGGGCATTATGTTTATTTTGTTTT<br>TGTATAGTAATTCTTAGTCATAGCACGTA | intrinsic | -18.4 |
| | phiKZ T014 | 27684 | + | TTTAATGCGTGTTTTAACACTATGTAAATATAATGCCCTCTCCTTATGGAGAGGGCTTATGATGTATTTTC<br>TCTTTTATCATCTGTAGTACAATAACTCA | intrinsic | -15.8 |
| | phiKZ T015 | 29374 | + | TGATCCAAACGCTGAAATAGATAAATGAACATATCGCCTCCCCTAGGGGAGGCTTTATGCTGAGTTAAAAA<br>AAGTTTAAGATCTATATTACTAATTTGCA | intrinsic | NA |
| | phiKZ T016 | 32943 | + | GCAACTAGTAGTGGTCTTGTATTCGATTAACTTATACTGGAGCCCTTCGGGGCTCCTTTTATTTTTTTTGT<br>AACGAGGTGTAATATGAAACCAGTAAATA | intrinsic | NA |

| ϕ | ID | TTS position | + /- | -60 to +40 sequence region | predicted type | ARNold hairpin energy (kcal/mol) |
|---|---|---|---|---|---|---|
| | phiKZ T017 | 33561 | + | ACTGCTAAAAAGCTCGACGAGCAAATGGCAGCTGAACTTGATCTGACGTTATCTCCGTTTACTATGGTTCT<br>GAAACACGTAGTTCGTCTCCCCAACATAG | - | - |
| | phiKZ T018 | 34911 | - | CGCTCAATAAAAAAAATAAAAAAAAAGACATAATGCCCTCCCATTGCGGGAGGGCTTATGCCATTAAGATA<br>ATATCTAAATAACACTATATTTAAAAACA | intrinsic | -18.1 |
| | phiKZ T019 | 34942 | + | ATAGTGTTATTTAGATATTATCTTAATGGCATAAGCCCTCCCGCAATGGGAGGGCATTATGTCTTTTTTTT<br>TATTTTTTTTATTGAGCGGCTGTGATTGT | intrinsic | -18.4 |
| | phiKZ T020 | 39545 | + | ATTGACGCAAATAGGAAAGCCTATCGGTTCACATAGGCTTCCTACTAGGAACCGATATGGAAAAACGAGTA<br>TTTATCGTATTTAAATCTAAAGAGTCAGA | - | - |
| | phiKZ T021 | 40406 | + | GCACCACCAACTTCAGTGACATATGAATAACTATAAAGGAGCCATTATGGCTCCTTTTATGTTGTTTTATT<br>TTTACTTACTTACATCATCTAATGTGAAT | intrinsic | NA |
| | phiKZ T022 | 40502 | + | GAATGTAAGTTGATGTGCTACCTCTCAATTTAGGGTACCCGTATAGGGTACCCATTCTTTCCTTTTCTTTT<br>TTAGTTAAAAAGAGAGAAATCATTCACTA | intrinsic | -17 |
| | phiKZ T023 | 42806 | + | GATCGACTCATCTCCGACTTCTATTAAGAAATATAGCCTCCCCTAGTGGGAGGCTTATATACTAAAAAATA<br>ATTAAGATTTATATAGTCTGATTGCCAAT | intrinsic | NA |
| | phiKZ T024 | 45844 | + | TCGCTCTGGTCGACTAGATGGCTCTGATGCTCGTTCAGCAGATGCCCTACAAGTATTCACTAAAGTAAAAG<br>AAGAAGCAATCCGCCATCAGGACCTACCT | - | - |
| | phiKZ T025 | 47888 | + | TCCGTTGGTGGTCCATTGTAAAAACTGATATAGCGGCTCCTTCGGGGGCCGCTTATATCTCCTTTTTTTGT<br>TAATCGATAAACAGGAGTTTGGTATGGGT | intrinsic | -16.7 |
| | phiKZ T026 | 50199 | + | GAAAAGAATGGGGTGCTTGATAAATACCTTATTAAGTTAATCAAATCTAGTTCCCCCAATGAGTAATCGTT<br>GGGTTGAGTTCTCTCTAATTGCTGGGATA | - | - |
| | phiKZ T027 | 50539 | + | AGAGCTAGAGCTGCACGTAATGGTGCGGGTATCAATTAACGACTGATACTGAACAATACGGCTGATTTTGA<br>CGGCGATCAATTGAATCTTACGTTAATGC | - | - |
| | phiKZ T028 | 52650 | + | TCAGTCGTAGGTAATCTTCTAGGTTAAATAAATGAGAGTCCCTACATGGGGCTCTCTTCTATCGTTTAGGA<br>GAGATCTATGTCAATTGATCTACTCGTAC | - | - |
| | phiKZ T029 | 53514 | + | ATGTAAGTATTTAACTGTATGTATCAAGCAATATAAAGGGAGCCATTTGGCTCCCATCTTAACTTAATTTT<br>TTTCTTTAGTTTTGGAGTTCAATATGCCA | - | - |
| | phiKZ T030 | 55165 | + | CAAAGAAACCTGGCATATTTCGATATTTAACAGTAGGCGATGTCCATATGGGGCATCGCCAAACCCCTACT<br>GAATTAATCATCAATAACTTTTGGAAAAC | - | - |
| | phiKZ T031 | 57939 | + | AGGATATGCCTTCCTTCTAAAACATATTTCGGTGCTTACTCTCGAAGTAGGCACCGATATGCCCTCAGTAC<br>TAAATGCCGTTATACTAAAAAAAATTTGA | - | - |
| | phiKZ T032 | 65671 | + | ATGCAGCTGCCTGCAATACGAGAGGCGGGGTAAGCGTAGCGAACTTACCTGAACATAACGAAGGGCGTCAT<br>CTGTAAGGTAATGGAAGATGAAGATATGC | - | - |
| | phiKZ T033 | 68709 | + | ACTGCGATAGACGTGGAGGACGTGAGGTTTGGGGAGCACTGCTCCCCTTACCGATCGTTGATTTTTTTATG<br>AAACCGATTCTTACAGCAGAACGATACAC | - | - |

| ϕ | ID | TTS position | + /- | -60 to +40 sequence region | predicted type | ARNold hairpin energy (kcal/mol) |
|---|---|---|---|---|---|---|
| | phiKZ T034 | 70298 | + | CGCATGATGCGTTTATAGTAAATGGCATAAATGATCCCTCCCTTAGTTGGGAGGGATTATGCTTTTTTTTT<br>ATTTTTTGTATTTAAAGCATTTTAAACAG | intrinsic | -16.9 |
| | phiKZ T035 | 71188 | + | CCATTAGGATATCTGATGGTTTAATAAATATTACCCCGCCTTTTATTGGTGGGGTAATATGCCATCTTTAT<br>ATCAAAAGACTTTTGAATATCATATCCCA | - | - |
| | phiKZ T036 | 71395 | - | CGTTAACTAAATAAAAAAAAATAAAGACAGGCCCCTCTCCACATGGAGAGGGGTTTATGTCGTCTTACTTA<br>CGTGCAAAGCTAGCTACAAGACCTTTGAT | intrinsic | -15.6 |
| | phiKZ T037 | 71448 | - | TACTCATAATGAGGCTGTTAATAACAATAACCAGTTAAATGAATTAAACAAGCCGTTAACTAAATAAAAAA<br>AAATAAAGACAGGCCCCTCTCCACATGGA | - | - |
| | phiKZ T038 | 76410 | + | TCTTATCGCGATCGCATCAGTTAAAATAGATGGAGAAGTTACTCAGCTTCTCCATTATTTAGATTATAACC<br>CGAAAGGAACTAGTAATGTCTAAAAAGAA | - | - |
| | phiKZ T039 | 77890 | + | CTTCTTCCGGCAGAAAGCTGGTATGTAAACATATTGCCTCCCCTAGGGGAGGCTTTATGCTGATTATTACT<br>TTTACTAGGTACTAATATAGTACGTAGAT | intrinsic | NA |
| | phiKZ T040 | 80313 | - | CAGAATTTAAAATACCAAATTAGTTAACGAAATAAACCCTCCCGTAATGGGAGGGTTTTATGATGTGTCGA<br>AGTTTATATTAGCAGTAATGATCAATTGA | intrinsic | -15.9 |
| | phiKZ T041 | 82568 | - | GACTTTACGAATATTTCCTTCAGTCCTATCATTAGCAAACTTCTCATAATCGGCTGGATCAACGTTTTTCT<br>TTAACCATGTTTCATCCAATAATTGAATC | - | - |
| | phiKZ T042 | 87899 | + | GATTGCATCAGTACTGGATTTCGTCTCAGGTTTAATAACTGATTTTTAAATTCACTTGCCATTAATTAACC<br>CTCTTTCTGAGGTGGTGGGGGATTGCGTA | - | - |
| | phiKZ T043 | 91723 | + | TCATTTAGTCATTCGGTCTTACTTGAATCAATCGACGGGGACAGCCCCGTCGATATGATCGCTACATGCGA<br>TGGATATCTAAAAGAATTATTGCCATCAA | - | - |
| | phiKZ T044 | 97650 | + | TTCCAAGCAGAATAATGATAAGTGACGACATATCGACGGGGCTCGGAAGAGTCCCGTCTTATGTTAGGAGT<br>TAGCAATGGAATTAATTGATATTTATAAA | - | - |
| | phiKZ T045 | 98802 | - | AGAGTTTATGAGTTATGCTATTCGTCTTAGTCATAACTCCAAGCGACGAACGTTAACATTCAAAATATTGC<br>GCTGGAATCTTACTTTAGATTTTCATCGC | - | - |
| | phiKZ T046 | 101370 | + | TGTGAATCGTGCCATTAAGTAACAAACATAACGGGGAGCCTTCATGGCTCCCTTTTATGCTGTTAGACCAT<br>GATCTTATGATCTTAAACACATGGAGTTA | intrinsic | -15.2 |
| | phiKZ T047 | 102031 | + | TCCCAGAATATTTGAATAAATATTTTACGTTCCCAACTACAGAATTAGCTGATCTTAGTCAAAAAAATAAA<br>TCACTGGTTATATACGAAAGACAGGTCGA | - | - |
| | phiKZ T048 | 107495 | + | AGCCTGGAAATGCACCCAATCTAATGAGAAGGTCTATGTTTTGATCTTCTCATTAGTACTCTTGTAGAAGG<br>AGCCATTATCATGGCTGAATTTATTATAG | - | - |
| | phiKZ T049 | 108979 | + | AAGTATTCGATACAGTTCGTAAGATGTAAAAATGGATAGGGCCTCACTGAGGCCCCATTCAAATATAATTA<br>CGATAATAAAATTATATTTGAATGAGTTT | - | - |
| | phiKZ T050 | 112512 | + | TATACACCAGCACGTATATTGGTCAATAGAAAAAAACCACCCTATATAAGGGTGGTTTATAAAAGAGGTGA<br>AATATGCATAAACTTAATCCTGGTTGTTA | intrinsic | -13.5 |

| ϕ | ID | TTS position | +/- | -60 to +40 sequence region | predicted type | ARNold hairpin energy (kcal/mol) |
|---|---|---|---|---|---|---|
| | phiKZ T051 | 114207 | - | TCTTTTATTCTTAAAAGTTTAACTGAATAAATGCCCTCTCCCGAAGGAGAGGGATTATGTCGTTAATTTAT TTAACCAGCGATGTTTCCCTAACACGAGA | intrinsic | -15.6 |
| | phiKZ T052 | 114239 | + | GAAACATCGCTGGTTAAATAAATTAACGACATAATCCCTCTCCTTCGGGAGAGGGCATTTATTCAGTTAAA CTTTTAAGAATAAAAGAATTAGCCCTACT | - | - |
| | phiKZ T053 | 114725 | - | TTTCTCTTTAGCGGGAGACTTAACTCCACTTGGGAGCCGCTTTAGCGGGCGGCTCTCACTTAAACAAAAAA AGAATTCAGATAAGTCCCTCTCCTTTAGG | - | - |
| | phiKZ T054 | 115943 | - | CCACTGGTGGAGATGACTGGGGATATTAACAACATAAAGCTCTCCCTTCGGGGAGAGCACTATAATTTTAA TAGGTAATCTAAATGAGTGCAATTGATAA | - | - |
| | phiKZ T055 | 117934 | - | ATCGTACACTAAATTACGGTTACCTGCATCCCTAAGTTCATTAGTCGCTAGGCGGCTCTCTAAACGATTGT GTAAATCCTTCATCTTAAATTGTTCAGGA | - | - |
| | phiKZ T056 | 118183 | - | TTTTGGTTTAAAAATTTGGGTCTACTAAGCATCTTTTCCGGAATCAACCTAGCAGCTCTCCAAATAATAGG TAAAAATTTGTGTTGACGCCACGGATTCA | - | - |
| | phiKZ T057 | 120629 | + | ATGCCCGGAAGTTAGAAGAGTAACAGACATAGGCCCTCCCTTCGGGGAGGGCTTTATGTTGTATTACATTA AAAAAAGTTTAAGATCTATATTACAGTAA | intrinsic | -18.4 |
| | phiKZ T058 | 126821 | - | TACTTTAATAATTTGTCATGGATACCAACAATACCAGTAACTCTTGAATTTGAAAGCAGTTAAATAAAAAA AAATAAAGACATAGCCCCTCTCCATGCGG | - | - |
| | phiKZ T059 | 133067 | - | CTTAGAATATTAGGTGTAGATGGTTAACGTCATAAGCCCTCCCGTAATGGGAGGGCATTATGTTGTTTAGA AATTATCCAATCTCATCTGATTCAGTTGA | intrinsic | -18.4 |
| | phiKZ T060 | 133099 | + | GATGAGATTGGATAATTTCTAAACAACATAATGCCCTCCCATTACGGGAGGGCTTATGACGTTAACCATCT ACACCTAATATTCTAAGATATGCGGTGTC | intrinsic | -18.1 |
| | phiKZ T061 | 135111 | - | ACCACGTGGGTTCGTAAATAGATCATTACGATTAGCATCTAAATGCTGTTGATCACTTGCCATTGCATAGT TTGGAACTAGTGACAAGCCAACTTGTGCT | - | - |
| | phiKZ T062 | 137190 | + | GACCGTCTATAAACATCTGACCTGGCATAAGAGACACAGCCAAAAGCTGTGTCTCTTTTTGCTGTTTCAGT TAAAAGGTATTTCTTATGTATAATCCTAT | intrinsic | -10.6 |
| | phiKZ T063 | 142947 | + | GTGGTGCTCCTGAAAAATTTGCTTAACAGCATAAATCCCCTCCATATGGAGGGGAACTCTTACAAATTTAC TCAATAGTACAGTGTACTAAATTTTAACC | - | - |
| | phiKZ T064 | 145649 | + | ATAAAGATGCTTATGCAATAAAAACTATAATAGCCTTCCTCTTCAAGAGGAAGGCTTTATAATGCTATGTG GATTAAACATTTTACCACTATAGAGGTTT | intrinsic | -12.7 |
| | phiKZ T065 | 147179 | + | TGATAAACCACATCTTGTGCCAATGTAACTTTTTGAGAGTCCTTCGGGACTCTCTTATGTTGTTAAAATAT TACTCATTAGCAATAGTACATTACTGTAT | - | - |
| | phiKZ T066 | 149761 | + | ATTGATGAACCCACTTCTGTAGTTAATGGACACATCCCGGTTTCTACCGGGGTGGGTCTTAATACCACTTT CGTTGATCCTGTTACTATACCTGGATTAA | - | - |
| | phiKZ T067 | 152963 | + | TTTCCAAAGGTGCTGTTATTTAAAACGACATATTGCCCTCCCTTCGGGGAGGGCTTTATTTCGTCGTTTAA GACACTTTTTATATTATCTATACCAATGT | intrinsic | -18.6 |

| ϕ | ID | TTS position | + /- | -60 to +40 sequence region | predicted type | ARNold hairpin energy (kcal/mol) |
|---|---|---|---|---|---|---|
| | phiKZ T068 | 153024 | - | AATACTTTTCCATGTTTAAGCCATGATACCCAATAGAAAGCCCTAATAGGGCTTTCTATGCTGTTTTAAGC TATAATTACTACATTGGTATAGATAATAT | - | - |
| | phiKZ T069 | 155687 | - | AGTATAGATGTTATCGCCAGTTTCAAAATAACTAGGGGTCAATAAGATCCCTAGTTATGTCAGGTTTGGTA TATGAATGTTTACACTTATCAGATAGCTA | - | - |
| | phiKZ T070 | 158210 | - | GAGAAGTGTATTCCAAATTGATTGGTACGACATAATGCCCTCTCCTTTTGGGAGAGGGCTTATGCTGTTAT CCAATGATTTCACGAAGAATATCTTCAGT | intrinsic | -15.2 |
| | phiKZ T071 | 158242 | + | CTTCGTGAAATCATTGGATAACAGCATAAGCCCTCTCCCAAAAGGAGAGGGCATTATGTCGTACCAATCAA TTTGGAATACACTTCTCATGTAAACTTAA | intrinsic | -15.5 |
| | phiKZ T072 | 160356 | - | TGCCTACACCTAAAAAATATAGTAAGGATACTAGTAAGAAGTGTATGGTTTTCGAACTACCAAGAAAGAAA CCGGTACCGCCAACACCGATTATTACAGA | - | - |
| | phiKZ T073 | 161591 | - | TATCTAGAAGTTTCTTAACTTCTAAAGGATCCTTAAACCAATCGTCATAGTTACCTTGTGCATTCTTACCA ATAGGGTGAACCACCTCAATTTTAAATTC | - | - |
| | phiKZ T074 | 162450 | + | GAGTGTAGAACAATTTGGTTAATGCATAAATGAGAGAGGGCAAAGGCCCTCTCTTATATTAATCTAAAGGT GGCGCAATGCGTAGATTCACTCTTAATCT | intrinsic | -15.8 |
| | phiKZ T075 | 166843 | + | AGCGATGTCATTTGCTTTAGAAAGTATCGATCCCGCCTTCTCCATAAAAGAAGGCGGTATAATTACACTAC GTGCTTTAAAAGATGCATTAGTTCGAGCT | - | - |
| | phiKZ T076 | 167868 | + | ATTATGAGAATGAAGGTATTTAACAGCATAAGCCCTCCCATAGTGGGAGGGCATTATGTTCTTTAACTACC ACAAGTAAGATTAGGTCTACCAGTAGCTG | intrinsic | -18.4 |
| | phiKZ T077 | 171597 | - | CTTAATAGCACTTATAAAGATAGTTACTAATATAATGCCCTCCCATTGCGGGAGGGCTTATGTTGCTTATA ATTATTTTAATACTAGATTTTTATTAAAT | intrinsic | -18.1 |
| | phiKZ T078 | 172327 | + | GTATTAACTTCGATGGAATACTGATATACATATTACCCTCTCCTTCGGGAGGGGGTTTATATCGCATTAAT TTTTTTATTACACTAAAAAAAGTTTAAGA | - | - |
| | phiKZ T079 | 173039 | + | CCGCTAAGCGCATGCCGCTGGAAACAATCACATCAGTTGCTGTTATTGGCGGGTTTATTTTGTACGTTGGT ACACGGTTTTATAAAGCATGTAAAAAATT | - | - |
| | phiKZ T080 | 173150 | + | GTCATGCAGTATACTTTAGGTACAATAACCCTAGAGGAAACTGCAATGGCCGTATTAGACCGTCAATTTAA TATTCGTTTAAATGGTGCTGATTTAGATA | - | - |
| | phiKZ T081 | 173797 | - | AAGAATTAGAAAACGATTGATATTAACGACATATAGCCCTCCCGTAATGGGAGGGCGTTATGCACTTAATA CAAATCGATATCTAATACATCACCAACTG | intrinsic | -18.3 |
| | phiKZ T082 | 177206 | + | AAAATAGTAGTTTTTAATATTGGGGCTTGATATTCCTTTGGGGTATCTGGCCCCAATCTTATAGATTATTT TTTTTGTTGGAGACTACCATGGTTGATTT | - | - |
| | phiKZ T083 | 178352 | - | GCGAAAAGTCTTGGCTGGGATATTTCACCAGTTAAACATAGGCGCCCTGGCTTCATGCGTTAAAAATAAAA AAAAGACATAATGCCCTCCCGTAATGGGA | - | - |
| | phiKZ T084 | 179971 | - | AATCATTAAAACATGCAATCGATGATAAACTCCGGTATTTGGAAGGATACCGGAGTAATACAGCGAATATT TAAAGGTGCATGATTTTGATTAAGTATTT | - | - |

| ϕ | ID | TTS position | +/- | -60 to +40 sequence region | predicted type | ARNold hairpin energy (kcal/mol) |
|---|---|---|---|---|---|---|
| | phiKZ T085 | 181455 | - | CCCTTGCGGTCTTTGCAGATGGTTTCAGTGTAGATGGCAACTCTAATAAAGTTGCCCTCTCTGAAGAGCTG<br>GAAGCATCTGTAGAAGAGTTCTTCGAGGA | - | - |
| | phiKZ T086 | 186346 | - | GAGAGCTAGAGCTGCACGTAATGGTGCGGGTATCAATTAACGACTGATACTGAACATAACGTTCGACGGGG<br>ACACTGTATCATTTAACGCAGTTTATTCA | - | - |
| | phiKZ T087 | 196087 | + | TGCCAATTAGACTACGTTAATCTATTAAAGGAGCCTCCCTATTGGGAGGTTCCTTATTTTGTCAAGGAGAT<br>TCCTATGGCGACAGATAGTAAACCAATTT | intrinsic | -13.2 |
| | phiKZ T088 | 199193 | + | TACTGAAGGTACCTATAGTGGTAATTGACATAAATGCCTCCCTTCGGGGAGGCTTTATTTCGTATTAAAAA<br>AAATTTGAGATTTATATTACTAGTGTGTA | intrinsic | -15.3 |
| | phiKZ T089 | 200561 | + | AATATATACTAGCTATTTCAATATAATAAGATCAGTGGGTGCATTGCCCACTGGTCATTTTTTATATTAAA<br>AAAGTTTAATGGCTAAATAGTACATTTTA | intrinsic | -10.2 |
| | phiKZ T090 | 202344 | + | TAATCGCGATAAAAAATAAAATACGATATAATAGAGTCCCCATTACGGGGACTCTTCTTTATAATTTTTTT<br>GTTAAAAGGTATATCAAGTGTTCAATAGA | intrinsic | -15.8 |
| | phiKZ T091 | 207011 | + | ATTCGTGGACTAAAAGTCTAATACTTTTATAGGAGAGAACCATGATGGTTCTCTCTTTTTGACGTAAGTGC<br>ATTATTGTTATGATAAGACTTTCGTACAA | - | - |
| | phiKZ T092 | 208177 | - | AATGATAAACTTGTAAAAGTTTAACGGCATAAGCCCTCCCGCAATGGGAGGGCATTATGTTGTTTTAACCA<br>CGCGTTATATTAAATAACCGTCTTTTCTG | intrinsic | -18.4 |
| | phiKZ T093 | 208211 | + | ATTTAATATAACGCGTGGTTAAAACAACATAATGCCCTCCCATTGCGGGAGGGCTTATGCCGTTAAACTTT<br>TACAAGTTTATCATTTTCTATTTTCCAAT | intrinsic | -18.1 |
| | phiKZ T094 | 210031 | - | TCAGATATTCAGAAGGTGTATTATACAGTGAGCCCGGAGTTACCTCCGGTGCTCTCTGAATACATTGCTTA<br>TGATGGTGACCCTGCAACAGGTGGTAGTC | - | - |
| | phiKZ T095 | 213001 | + | GACACGAACCCCAACTACGGTGCCTTGCTACTCACCGACCTCTAGATAGGGGTCGGACCTATTACATTAAA<br>GGAGCCCACCAGGGGCTCCTTTATGCCAT | - | - |
| | phiKZ T096 | 214091 | - | AGTCCTTATTCTATTAAACATATAAGTAAATAATCCCTCCCCAATAAAGGGGAGGGTATATGTTATCTTAA<br>TTCATGCTTTTGAAAGATCATATTTAATA | - | - |
| | phiKZ T097 | 214123 | + | TCTTTCAAAAGCATGAATTAAGATAACATATACCCTCCCCTTTATTGGGGAGGGATTATTTACTTATATGT<br>TTAATAGAATAAGGACTCTACCTTGGAAT | intrinsic | -18.8 |
| | phiKZ T098 | 218207 | + | TGCAGTTATAGGTGGATCATTATGGGAAATAATTAGACAGTGGTTTTAGCCACTGTCTTTATAATGTATTT<br>TGAGTTATAAAAATGTCCATAGAAAAAGC | intrinsic | -13.1 |
| | phiKZ T099 | 221066 | + | CTTCCGTAAGAACAAACAAAAAGATGTTCGTCATTGCAGTGTATGTGATAATATTAAACTAAAAGAAAATG<br>GTATCGAAGAAGGGCGGGCCAAACCAATT | - | - |
| | phiKZ T100 | 221335 | + | CAGTCTTCCGCCGCGCTATTACTGCCTAACGATATAGGGGACCTTCTGGTCCCCTTAGTCATTATTATTTT<br>TTTTTGGATTTTAGCTATGGATGATATTG | - | - |
| | phiKZ T101 | 221825 | + | GTTGCGATAATGAGTAAGCGGTAATGTTGTCATAAAGCCCTCCCTTCGGGGAGGGCTATTGCACTAAATTT<br>TATTTGAGATCTACATTATAGATTAGCAT | intrinsic | -16 |

| ϕ | ID | TTS position | + /- | -60 to +40 sequence region | predicted type | ARNold hairpin energy (kcal/mol) |
|---|---|---|---|---|---|---|
| | phiKZ T102 | 223931 | + | TTCTATTACGTACCTTTCGAGGCATATACTCCATGACTTATGTTGTGGGGTATATGTGTTATAATGTTTAA<br>GAGGTTAAAATGCAAAAGCAACTAAATTC | - | - |
| | phiKZ T103 | 224732 | + | ATTAGTACTGGAGTTATTATTCGTTTAATATTAAGGGACTCTTAATGAGAGTCCCTATTTATTTTGCGTAA<br>GTGTATATATGAAAAAGAAAGCATTTATT | intrinsic | -13.3 |
| | phiKZ T104 | 225793 | + | GACATTTAAAAGACTGGGCAAAAGACATTCTAAAAGAGTGTTGGCTGCCCAGTCATCTTTTCTGCCGTTTG<br>GTGAATTAACGTTGAGGAAGCAAAGAATG | - | - |
| | phiKZ T105 | 227922 | + | GATAACAACTACGGTAAAGCTGTGGTTGTAAAATAAATACCAGCCCTTCGGGGCTGGTTTATTTCATCTAG<br>ATGAGAATCTGTGGGTTGCAAATGTATAA | intrinsic | NA |
| | phiKZ T106 | 231265 | + | TAGGTAAGATACTAGCCGATTGGTTAGTTGCGCTTAGGGAGTCCTTCGGGACTCCCGTATATCGTCTAGGA<br>GGTTTTATGTATACAGTTACACTTGAACA | - | - |
| | phiKZ T107 | 233301 | + | CACACATTGCTGGTAAAGTTGCTGTCTAACATAAATGCCTCCCTTTGGGGAGGCTTTATTTCCCTATGGTT<br>CATTTGGGATATTCAGCGAATACTACTAT | intrinsic | NA |
| | phiKZ T108 | 235828 | + | ACAAAAAGAGTAAAACTATTTAGTTGCATAAAGAGAGAGGATAAATTCCTCTCTCTTTTTATGTTGTTTTA<br>TAATTTGTATACTACAGGTATATTTCAAT | intrinsic | -13.3 |
| | phiKZ T109 | 239132 | + | TGGACAAGCTTCACTATTGAAAAGCTACTTGTATAATAAGAGCCCTTCGGGGCTCTTTCTATTTCGCAATT<br>AGGTATCTGTATTAAATGGATAAGTATCC | intrinsic | NA |
| | phiKZ T110 | 240325 | + | GAAATTGTATGTTAAAATAATTACTTAATATACACCCTCCCTTCGGGGAGGGTTTATTTATCTTTTTTTG<br>TATTTAAACTATTTAAAAAGCTATATTAC | intrinsic | -16.3 |
| | phiKZ T111 | 243801 | + | CAAAATGGGATGCCCTGAGAGTTATAAGAAATAATTAGAGGACCTTCGGGTCCTCTTTTTAACCGACTGGA<br>CGGTTAAAACTAGGTGGGATAAATTAAAC | intrinsic | NA |
| | phiKZ T112 | 243999 | + | ACACTCATAAGAAAAAATATAAATAAAGTCATAAACCCTCTCCCAAAGGAGAGGGCTTTTACATTAAATTT<br>TTTTTGAAATCTATATTACACTAATACAA | intrinsic | -15.4 |
| | phiKZ T113 | 248230 | + | CGAACCATCAGCGCGAATTATCTCGCTGATTAAAGATGCGGTCGCATGACCGCTTTCTTTTAAGAAGGTAA<br>CTAATATGTCTATCTCTGATGTAGTATAT | - | - |
| | phiKZ T114 | 248532 | + | TTCAATTTGCTAAAGAAGCAGTACGCAATGCTGAAGGCATTGTAATAAAATAATCAATCTGGAAAAAAAGG<br>AAATATAAATGTATATTGATGAAAAGTAC | - | - |
| | phiKZ T115 | 249008 | + | AGTTGACCATTTCTGCATTCTAATTAATTATGAAAATGGGGGTCTAAAGACCCCCTATCTATTTAATAGAA<br>TGAAACTAATCTTAATAATGGTGAATACT | - | - |
| | phiKZ T116 | 250278 | + | GTGTGTCAATTCGTTTCACAATGTAACGTCATAATGGCCTCCCTTCGGGGAGGCTTTTATATCCCTTTATA<br>TAAACAGGTATATTCAAATGAAAATGAAT | intrinsic | NA |
| | phiKZ T117 | 254771 | + | AGTATATAGACTCGTAATTTAATAATGGTATAATTGCCCTCCTTCGGGAGGGCTTCTATTTCGTAAAAAGG<br>AAAATACAAATGAAAGCATTCATTCCAAA | intrinsic | NA |
| | phiKZ T118 | 262760 | + | GTTTCAGTTTTGCTGAAGAAGTACCAATGAGATACCCTCCCCTGATCGGGGAGGGTTTTTGACATCTAAGA<br>GGAAAATATAATGTGCGAACAACATCGTG | intrinsic | -17.8 |

| ϕ | ID | TTS position | + /- | -60 to +40 sequence region | predicted type | ARNold hairpin energy (kcal/mol) |
|---|---|---|---|---|---|---|
| | phiKZ T119 | 263292 | + | GAAGGTGCCGCCGAGATCCCACCTATTGATAATGCCCTCCCTTCGGGGAGGGCTTTATTTCGCTTAAGCCTATTTATTTTATTATAAGGAGTGAAAATGA | intrinsic | -18.6 |
| | phiKZ T120 | 265438 | - | TAAGTGAGTTTAAATAAAGAAAATAAAGACATAGCCCCTCTCCATGCGGAGAGGGGTTTATGCCGTAATTATACACTCACTTTGATAAGATTTTTAATAT | intrinsic | -15.6 |
| | phiKZ T121 | 265484 | - | AACTACTTAATGATTGGCTAGATGATATTCCAGTAGAAGATTATTCTAAGTGAGTTTAAATAAAGAAAATAAAGACATAGCCCCTCTCCATGCGGAGAGG | - | - |
| | phiKZ T122 | 266616 | - | GGGATTGAACGTGAAAGACTCAGTAAATCATATTGCCCCTCCATATGGAGGGGCTTATGTTTGTCATTAATCAGGAAAATATAAAATGAGTCATCTATTA | intrinsic | -18.2 |
| | phiKZ T123 | 267119 | + | GATGTAAAAGATGTAGCAGAGTAAGATAACATAAAGCTAGCCCATTAGGGGCTAGCTCTTATATTTATTTTTTATTTGTTTTAACTTAGAGTACTATAAT | intrinsic | -12.9 |
| | phiKZ T124 | 267142 | - | AATATAATCAAAGAGCAACAAGATGAATTAACTAAATTTATTTATTATAGTACTCTAAGTTAAAACAAATAAAAAATAAATATAAGAGCTAGCCCCTAAT | - | - |
| | phiKZ T125 | 268768 | - | AGATTTAGTTATATTTCACTACTATGTAAACTATAAAGGCAGCTAATGCTGCCTTTATGTCGTCTTGTAAAAGTACCCAGTAGTTGAATCTTATAGCTAA | intrinsic | NA |
| | phiKZ T126 | 270485 | - | GTGGTAGAACTTATTATAGAGTGTGTCTAAATGCCAGGGGTTTGCCACCCCTGGTTATATTCATTGTTACTATTATAAATTCATTTATAGATGAGAAAAG | - | - |
| | phiKZ T127 | 271740 | - | CGGTAGAGCAAGTGACTGTTAATCACTGGGTCCCTGGTTCGAGTCCAGGTCACGGAGCCATATTCTAAAGAGTAGCTTCGGCTACTCTTTTATGTTGCCA | intrinsic | NA |
| | phiKZ T128 | 271822 | - | GGTTAGAATACCCGCCTGTCACGCGGGTGGTCAGGGGTTCGAGTCCCCTTGGGCGCGCCATTTAATTCCGTGATAGCTCAGTCGGTAGAGCAAGTGACTG | - | - |
| | phiKZ T129 | 272869 | - | TTAGAGCAGGCGACTCATAATCGCTTGGTCGCAGGTTCAAGTCCTGCTGGGCCCACCATATACTAGCCTCCCACTTGGGGGAGGTTTTATACTGTCTCAT | intrinsic | -11.5 |
| | phiKZ T130 | 273267 | - | TAGCGCGCCTGCTTTGGGAGCAGGATGTCGGGAGTTCGAGTCTCCCCACTCCGACCATTTTAATAATAGGTAAATAGGATGGATAATAAATGGATATCAT | - | - |
| | phiKZ T131 | 274007 | - | TAAAACTCTTTGCTGGGCATTAATGACGATATATCGCCTCCCTTCGGGGAGGCTTTATATTTTGTTTTTACGTATGTATATTAAAATATGTATAAACAAC | intrinsic | NA |
| | phiKZ T132 | 276157 | - | AATGATGGTACACGAGATATAATGTAGACATAATGACCCTCCCTTAATTGGGAGGGTTTATGCTAACAATTCTATAGCACTCTTATTAACAGTCATCAAC | intrinsic | -16.8 |
| | phiKZ T133 | 277829 | - | TGCAGGCGGTTCGTGCACTCTTTAAGTGTATACACCCTCCCTTAATTGGGAGGGTGTTATTCCCAATTTATACTAACCTCTATTATTTATTGTAAGAAAT | intrinsic | -15.3 |
| YuA | YuA T001 | 60 | - | CGGGTCGCAGCCGCAACGGCGATGTCTTCGACTTGCTCCGGGGTCATGGGCATATCGGGGTCCTCGCGCCAAGATGATCTTGGGCCGTATTTTACGGAAC | - | - |
| YuA | YuA T002 | 395 | + | TCGCCCTCCTAGGCATCTTCTTCTCGGGGCTGGTCAGCTTGCTCCTCCTCGGGCTCAAGGAGTGGTTCCAAAAATGAGAATGCGCTTGGGATACTTACCT | - | - |

| φ | ID | TTS position | +/- | -60 to +40 sequence region | predicted type | ARNold hairpin energy (kcal/mol) |
|---|---|---|---|---|---|---|
| | YuA T003 | 411 | - | CGATGGCGCTTGGGGTCTATACTTCTTCAGGTGTGAAGGTAAGTATCCCAAGCGCATTCTCATTTTTGGAA<br>CCACTCCTTGAGCCCGAGGAGGAGCAAGC | - | - |
| | YuA T004 | 471 | + | AGAATGCGCTTGGGATACTTACCTTCACACCTGAAGAAGTATAGACCCCAAGCGCCATCGAAAGCCCAGCG<br>CCGTAACAGCTCCGACCCTCCTGCGCTTA | - | - |
| | YuA T005 | 1004 | + | AAGTCCAAGCTGACCATCGACACCGCCTGCTGGCTCTGGCTCCGCGGGCTCATCGACGGCGTGCTGGTGGT<br>CGCCCCGAACGGCGTACACCGCAACTGGG | - | - |
| | YuA T006 | 2477 | + | CTGTACATCGACCTCGTGGCTGAGGACTCCGTGGATGAGAAGGTGGTTGAAGCGTTGAGGAACAAATTCAA<br>CGTGGCGAGCCAGATCACCGGCGACCGCC | - | - |
| | YuA T007 | 2607 | + | CGTGTATTTGTTGTACAGAACCAACACCGCTGGGACCGCGACAAGCAGCGGTTCGAGCCCAAATTCAACCT<br>CGCCCCGGCGGAGGAGTTCGGTGAGCTGG | - | - |
| | YuA T008 | 3188 | + | GTGCCTGAACTCATGGACCAGATCGGCATTGGCGAGTTCAAGACCACCACCGGCCTGAAGATCAAGATTGA<br>CGAGACGATCCGCGCCAGCATCCCGAAGG | - | - |
| | YuA T009 | 3504 | + | TCCCCCTGGACCTGTTCGGGGTCCATCGTCAGCGCGTGTCCAAGATCGAAGTGTGACCAGAATGCCGGAGG<br>ACGGCCCCGGGCCTTCCTCCCGACCGAAC | - | - |
| | YuA T010 | 4557 | + | ACCCGGACGGCCCGCCCCCTCGGGCCGTACCTTCAGGGGGCCTTCGTGGCCCCCTTCTTTTCATCAGAAAC<br>GGAGAACGATCATCATGTGGAGTCCGCAA | intrinsic | -17.3 |
| | YuA T011 | 14259 | + | CCATGATCTTCTCGCCGTTCTGGTGGTACTGGAGCTGGTGGTTGGTCTGGCCCCCTCGCCCCCGCGTCCTT<br>CGCCGGACCGGCAACGTCATCTTCGTGGA | - | - |
| | YuA T012 | 22800 | + | CCATCATGAACTACAAGAAGCCTTCCGCCGCATACACGGGCACCCAATCCCGCGCGGCGCAAAACCACAAG<br>AAGTCGCACGAGGACTTGCTGCGCGAAGC | - | - |
| | YuA T013 | 29082 | + | AATTGCCGTGAATCGAAAAAAGTTGTATGAGGGGGCTTTACTTCTTCGGAGGCCCCCACTATACTTTCTTC<br>ACGTTCAACAAGAACGGCAACCACAGAAA | - | - |
| | YuA T014 | 29212 | + | TCTACACCGACCGCGCCACCGCCCGCTTCGAGGCCGGTTTCACCTCCAAGGCCGCTCAACAAGAAGCCCTC<br>CGCGACCTCAACAGCGCCTTCGAGCTGCT | - | - |
| | YuA T015 | 29343 | + | ATCGAGCACGACGCCCGCACCGCGGATCATGATGACGTGTACTGGAACCTGGCCGACTACCCCCACAACTG<br>GAAGGCCAAGCACTCCGCCCTCGCCCTCC | - | - |
| | YuA T016 | 31609 | + | AAATCATCGCCGCCATCGACTGACCAACCCCTCGGGGCGGAACCGATCCGCCCCGAACTTCCCAGAAAAGG<br>AGCAACACCATGACCAAGACCAAGACCAC | - | - |
| | YuA T017 | 33511 | + | GCCGCGTGATGCGCGATTGACGAACAGTACACTGAGAGGGGAGCGTGATGCGCCCGATTGAAAAACTTCAG<br>CGGAAGCCGATGGTGCCCTTCACCTCGGA | - | - |
| | YuA T018 | 33553 | + | GCGTGATGCGCCCGATTGAAAAACTTCAGCGGAAGCCGATGGTGCCCTTCACCTCGGACCGCAAGCAGCAG<br>TTCCTGGACCTGTTCCGGTCGCACCCGGA | - | - |
| | YuA T019 | 33639 | + | CGGTCGCACCCGGACTTGAAGGGGTGCCGGGGCCTTTGCGCGGAGGCGGTGGGCGTCTCCATAACCACTTT<br>GTATGATCATCTGAAGCGTGACCCGGAGT | - | - |
| | YuA T020 | 39269 | - | TCCGTCACGCCCGACCGCGGCCCGCCTTGAACTGACCCTTGGCGAGTCCCTGGGCGATCTTCTCGTTGGCC<br>GTCAGGACCTTGCCCTGCTGGCGCGGAGG | - | - |
| | YuA T021 | 39519 | + | CCGGGTGATCCGGCACGGCTGAAACCGCTTGATTCATCAACTTCTGAAGGAGGGCCAATCATGGCTTCTGT<br>TACCCTTGCCGAAAGCGCCAAGCTGGCCC | - | - |
| | YuA T022 | 39736 | + | TGGGCGACGTCATCATGGCCGGTGTCGGCACGACCTTCTCCGGCGCTGGTGCCGGTAAGGGCGCTGCGACC<br>TTCACCAAGGTCAACTCCAACCTCACCAC | - | - |

| ϕ | ID | TTS position | +/- | -60 to +40 sequence region | predicted type | ARNold hairpin energy (kcal/mol) |
|---|---|---|---|---|---|---|
| | YuA T023 | 40494 | + | GACGGCATCCTCAACTAAGGCAGTCCCGCCCTCCGGGGAGGGCTCTGCCCCGTAGTACCCGAAGGGCGGGC CGGTCTGAGACTGGTCCGCCCTTTTGTTT | - | - |
| | YuA T024 | 40533 | + | GGGCTCTGCCCCGTAGTACCCGAAGGGCGGGCCGGTCTGAGACTGGTCCGCCCTTTTGTTTGAATTCACAC TGGAGAACTCATCATGCCCGCTTACCTTG | intrinsic | -14.9 |
| | YuA T025 | 45006 | + | TTACAACCATGGCTAACAAGATTGACTCGAACGTCACCGGCCTCCGCTATGCGGAGGAGGACACGATCAAG AACCTGCCCGTGTCGCCGGTGTGGTATCC | - | - |
| | YuA T026 | 45248 | + | AACAACCTGACCCGCCTGCTTCAGGGCTTCTTCTTCGCGGACATCCGCGAGAAGGCCACCAACATCCCGAC CAACGGAACCGCCGTGCCCTTCACCGGCG | - | - |
| | YuA T027 | 45302 | + | GCCACCAACATCCCGACCAACGGAACCGCCGTGCCCTTCACCGGCGTGACGGGCACCAGCAAAACCTACAC CCTGGGCTCCGGCACCGTGGGCTCTCAGT | - | - |
| | YuA T028 | 45591 | + | TGTCGGGCAGCTATCCGCGCCTGCTCCGCGGCTCGGGCACGAAGGACCTGACGACCCTGGGCCTGATCCCG GGCGAGTGGGTGTTCATCGGCGGGGACGC | - | - |
| | YuA T029 | 47070 | + | CAGAAGATCATTGAGCAGTGTATGCGGTTCGGGATGCCCTTGCCCGACCGCATACAGAACGCCCCGGAGTT GACCCTTGGTTCGGAGCTGTACTACATCG | - | - |
| | YuA T030 | 47350 | + | CTAAGTGAGTTCAGCCGACGCATTACCCTCCGGGGCCGAAAGGTCGCGGAGGGCGCTGACGCCCTCACTCG CAAGGTCGCCCTCGCCGCGGATCAAGCCG | - | - |
| | YuA T031 | 47737 | + | GTCCAGGTTGTACAGTTCGGAAGGGTTGTTGACGGGGACCCGGGGAGCTGACGGTGGCCGAAGAACGCATT GACATAGTCATCACTGAACGAGGCTCGCG | - | - |
| | YuA T032 | 47883 | + | GCCGCGGGCGGGGTGGACTTCCTCAAGAACGCCCTGGCCTCCCTGGGCGCGTACCTGTCTGCGCGCGAACT CGTGCGCCTGCTGGACACCTACACCAACC | - | - |
| | YuA T033 | 53312 | + | GACAACCTCCCCATCAGTGAGCCTGAGTACAACTTCCCGTCCTGGCGGGGCATTAGCTTCCTGGATGACTA CTACAACCGCATCCACATCCGCCCCTCGA | - | - |
| | YuA T034 | 57947 | + | GTGGTCCGTCCGATCATCGACCACTCCGCGGCGAACATCGGCCCGGAGCCGGGGGTGACCTACCGCGTGGA CATCTACGCGGCCAACGGCAGCACGCTCT | - | - |

**Supplementary Table S5: Overview of phage transcription units (TUs) of 14-1, LUZ19, LUZ24, PAK_P3, phiKZ and YuA.** TUs are delineated by TSSs and TTSs defined by ONT-cappable-seq. The most distant 3' end of each annotated TSS is also provided (indicated in blue).

| ϕ | TU | TSS | TSS_ID | TTS | TTS_ID | TU length (bp) | +/- | Genes | #Genes | most distant 3' end | Genes in longest read with TSS | note |
|---|---|---|---|---|---|---|---|---|---|---|---|---|
| 14-1 | 14-1 TU001 | 707 | 14-1 P001 | 178 | 14-1 T001 | 529 | - | gp01 \| gp02 | 2 | 63751 | gp01 \| gp02 | |
| | 14-1 TU002 | 822 | 14-1 P002 | 2359 | - | 1537 | + | gp03 | 1 | 2359 | gp03 | |
| | 14-1 TU003 | 3708 | 14-1 P003 | 3581 | 14-1 T002 | 127 | - | - | 0 | 2229 | gp04 \| gp05 \| gp06 \| gp07 | |
| | 14-1 TU004 | 3974 | 14-1 P004 | 3581 | 14-1 T002 | 393 | - | - | 0 | 3216 | gp07 | |
| | 14-1 TU005 | 4070 | 14-1 P005 | 3581 | 14-1 T002 | 489 | - | - | 0 | 3245 | gp07 | |
| | 14-1 TU006 | 4479 | 14-1 P006 | 3581 | 14-1 T002 | 898 | - | gp08 | 1 | 3027 | gp07 \| gp08 | |
| | 14-1 TU007 | 5200 | 14-1 P007 | 4617 | - | 583 | - | - | 0 | 4617 | - | |
| | 14-1 TU008 | 5686 | 14-1 P008 | 5619 | 14-1 T003 | 67 | - | - | 0 | 5141 | gp11 | |
| | 14-1 TU009 | 6637 | 14-1 P009 | 5619 | 14-1 T003 | 1018 | - | gp12 | 1 | 4612 | gp10 \| gp11 \| gp12 | |
| | 14-1 TU010 | 8684 | 14-1 P010 | 6661 | 14-1 T004 | 2023 | - | gp13 \| gp14 \| gp15 \| gp16 | 4 | 6654 | gp13 \| gp14 \| gp15 \| gp16 | |
| | 14-1 TU011 | 8970 | 14-1 P011 | 6661 | 14-1 T004 | 2309 | - | gp13 \| gp14 \| gp15 \| gp16 | 4 | 6603 | gp13 \| gp14 \| gp15 \| gp16 | |
| | 14-1 TU012 | 9116 | 14-1 P012 | 6661 | 14-1 T004 | 2455 | - | gp13 \| gp14 \| gp15 \| gp16 \| gp17 | 5 | 6305 | gp13 \| gp14 \| gp15 \| gp16 \| gp17 | |

| ϕ | TU | TSS | TSS_ID | TTS | TTS_ID | TU length (bp) | +/- | Genes | #Genes | most distant 3' end | Genes in longest read with TSS | note |
|---|---|---|---|---|---|---|---|---|---|---|---|---|
| | 14-1 TU013 | 9247 | 14-1 P013 | 10828 | - | 1581 | + | - | 0 | 10828 | - | |
| | 14-1 TU014 | 13257 | 14-1 P014 | 14848 | - | 1591 | + | gp22 | 1 | 14848 | gp22 | |
| | 14-1 TU015 | 18699 | 14-1 P015 | 21518 | - | 2819 | + | gp30 \| gp31 \| gp32 | 3 | 21518 | gp30 \| gp31 \| gp32 | |
| | 14-1 TU016 | 23797 | 14-1 P016 | 24561 | - | 764 | + | gp38 | 1 | 24561 | gp38 | |
| | 14-1 TU017 | 23847 | 14-1 P017 | 24854 | - | 1007 | + | gp38 | 1 | 24854 | gp38 | |
| | 14-1 TU018 | 28318 | 14-1 P018 | 30402 | - | 2084 | + | gp42 \| gp43 | 2 | 30402 | gp42 \| gp43 | |
| | 14-1 TU019 | 31714 | 14-1 P019 | 34849 | - | 3135 | + | gp45 | 1 | 34849 | gp45 | |
| | 14-1 TU020 | 36158 | 14-1 P020 | 35525 | 14-1 T005 | 633 | - | gp48 | 1 | 34821 | gp48 | |
| | 14-1 TU021 | 36158 | 14-1 P020 | 35820 | 14-1 T006 | 338 | - | gp48 | 1 | 34821 | gp48 | |
| | 14-1 TU022 | 36804 | 14-1 P021 | 35820 | 14-1 T006 | 984 | - | gp48 | 1 | 35822 | gp48 | |
| | 14-1 TU023 | 38929 | 14-1 P022 | 38499 | 14-1 T008 | 430 | - | - | 0 | 38382 | - | |
| | 14-1 TU024 | 39095 | 14-1 P023 | 38499 | 14-1 T008 | 596 | - | - | 0 | 38383 | - | |
| | 14-1 TU025 | 39481 | 14-1 P024 | 38499 | 14-1 T008 | 982 | - | gp52 | 1 | 38384 | gp52 | |

| ϕ | TU | TSS | TSS_ID | TTS | TTS_ID | TU length (bp) | +/- | Genes | #Genes | most distant 3' end | Genes in longest read with TSS | note |
|---|---|---|---|---|---|---|---|---|---|---|---|---|
| | 14-1 TU026 | 40195 | 14-1 P025 | 38499 | 14-1 T008 | 1696 | - | gp52 \| gp53 | 2 | 38424 | gp52 \| gp53 | |
| | 14-1 TU027 | 43097 | 14-1 P026 | 40725 | - | 2372 | - | gp55 | 1 | 40725 | gp55 | |
| | 14-1 TU028 | 48622 | 14-1 P027 | 46442 | - | 2180 | - | gp59 \| gp60 \| gp61 \| gp62 | 4 | 46442 | gp59 \| gp60 \| gp61 \| gp62 | |
| | 14-1 TU029 | 49234 | 14-1 P028 | 46405 | - | 2829 | - | gp59 \| gp60 \| gp61 \| gp62 \| gp63 \| gp64 \| gp65 | 7 | 46405 | gp59 \| gp60 \| gp61 \| gp62 \| gp63 \| gp64 \| gp65 | |
| | 14-1 TU030 | 49599 | 14-1 P029 | 46817 | - | 2782 | - | gp59 \| gp60 \| gp61 \| gp62 \| gp63 \| gp64 \| gp65 | 7 | 46817 | gp59 \| gp60 \| gp61 \| gp62 \| gp63 \| gp64 \| gp65 | |
| | 14-1 TU031 | 50236 | 14-1 P030 | 48787 | - | 1449 | - | gp65 \| gp66 | 2 | 48787 | gp65 \| gp66 | |
| | 14-1 TU032 | 50440 | 14-1 P031 | 50323 | 14-1 T009 | 117 | - | - | 0 | 48915 | gp65 \| gp66 | |
| | 14-1 TU033 | 50518 | 14-1 P032 | 50323 | 14-1 T009 | 195 | - | - | 0 | 48431 | gp63 \| gp64 \| gp65 \| gp66 | |
| | 14-1 TU034 | 51005 | 14-1 P033 | 50323 | 14-1 T009 | 682 | - | - | 0 | 50320 | - | |
| | 14-1 TU035 | 51317 | 14-1 P034 | 50323 | 14-1 T009 | 994 | - | gp67 | 1 | 48852 | gp65 \| gp66 \| gp67 | |
| | 14-1 TU036 | 51317 | 14-1 P034 | 51089 | 14-1 T010 | 228 | - | - | 0 | 48852 | gp65 \| gp66 \| gp67 | |
| | 14-1 TU037 | 51666 | 14-1 P035 | 51089 | 14-1 T010 | 577 | - | - | 0 | 50541 | - | |
| | 14-1 TU038 | 53073 | 14-1 P036 | 51354 | 14-1 T011 | 1719 | - | gp68 \| gp69 | 2 | 49601 | gp67 \| gp68 \| gp69 | |

| ϕ | TU | TSS | TSS_ID | TTS | TTS_ID | TU length (bp) | +/- | Genes | #Genes | most distant 3' end | Genes in longest read with TSS | note |
|---|---|---|---|---|---|---|---|---|---|---|---|---|
| | 14-1 TU039 | 53073 | 14-1 P036 | 51089 | 14-1 T010 | 1984 | - | gp68 \| gp69 | 2 | 49601 | gp67 \| gp68 \| gp69 | |
| | 14-1 TU040 | 53073 | 14-1 P036 | 50323 | 14-1 T009 | 2750 | - | gp67 \| gp68 \| gp69 | 3 | 49601 | gp67 \| gp68 \| gp69 | |
| | 14-1 TU041 | 53097 | 14-1 P037 | 53335 | 14-1 T012 | 238 | + | - | 0 | 54776 | gp70 | |
| | 14-1 TU042 | 53097 | 14-1 P037 | 54728 | - | 1631 | + | gp70 | 1 | 54776 | gp70 | |
| | 14-1 TU043 | 53163 | 14-1 P038 | 53335 | 14-1 T012 | 172 | + | - | 1 | 54802 | gp70 | |
| | 14-1 TU044 | 54251 | 14-1 P039 | 54831 | - | 580 | + | - | 0 | 54692 | - | |
| | 14-1 TU045 | 59399 | 14-1 P040 | 60626 | - | 1227 | + | gp76 | 1 | 60626 | gp76 | |
| | 14-1 TU046 | 59807 | 14-1 P041 | 60727 | - | 920 | + | gp76 | 1 | 60637 | gp76 | |
| | 14-1 TU047 | 60949 | 14-1 P042 | 60583 | 14-1 T013 | 366 | - | - | 0 | 60578 | - | |
| | 14-1 TU048 | 61300 | 14-1 P043 | 60583 | 14-1 T013 | 717 | - | gp77 | 1 | 59828 | gp77 | |
| | 14-1 TU049 | 62417 | 14-1 P044 | 61649 | - | 768 | - | - | 0 | 61649 | - | |
| | 14-1 TU050 | 62463 | 14-1 P045 | 61593 | - | 870 | - | gp79 | 1 | 61593 | gp79 | |
| | 14-1 TU051 | 62562 | 14-1 P046 | 62469 | 14-1 T014 | 93 | - | - | 0 | 61613 | gp79 | |

| ϕ | TU | TSS | TSS_ID | TTS | TTS_ID | TU length (bp) | +/- | Genes | #Genes | most distant 3' end | Genes in longest read with TSS | note |
|---|---|---|---|---|---|---|---|---|---|---|---|---|
| | 14-1 TU052 | 63024 | 14-1 P047 | 62469 | 14-1 T014 | 555 | - | gp80 \| gp81 | 2 | 61396 | gp78 \| gp79 \| gp80 \| gp81 | |
| | 14-1 TU053 | 63303 | 14-1 P048 | 62469 | 14-1 T014 | 834 | - | gp80 \| gp81 | 2 | 61396 | gp78 \| gp79 \| gp80 \| gp81 \| gp82 | |
| | 14-1 TU054 | 65240 | 14-1 P049 | 63431 | - | 1809 | - | gp84 \| gp85 \| gp86 \| gp87 \| gp88 | 5 | 63431 | gp84 \| gp85 \| gp86 \| gp87 \| gp88 | |
| | 14-1 TU055 | 65678 | 14-1 P050 | 63778 | - | 1900 | - | gp85 \| gp86 \| gp87 \| gp88 \| gp89 | 5 | 63778 | gp85 \| gp86 \| gp87 \| gp88 \| gp89 | |
| | LUZ19 TU001 | 731 | LUZ19 P001 | 141 | LUZ19 T001 | 590 | - | - | 0 | 42269 (other side) | - | antisense |
| | LUZ19 TU002 | 731 | LUZ19 P001 | 457 | LUZ19 T002 | 274 | - | - | 0 | 42269 (other side) | - | antisense |
| | LUZ19 TU003 | 950 | LUZ19 P002 | 1704 | LUZ19 T003 | 754 | + | - | 0 | 3586 | gp0.1 \| gp1 \| gp2 \| gp3 \| gp4 | |
| | LUZ19 TU004 | 950 | LUZ19 P002 | 1768 | LUZ19 T004 | 818 | + | gp0.1 | 1 | 3586 | gp0.1 \| gp1 \| gp2 \| gp3 \| gp4 | |
| LUZ19 | LUZ19 TU005 | 950 | LUZ19 P002 | 2851 | LUZ19 T005 | 1901 | + | gp0.1 \| gp1 \| gp2 \| gp3 | 4 | 3586 | gp0.1 \| gp1 \| gp2 \| gp3 \| gp4 | |
| | LUZ19 TU006 | 1029 | LUZ19 P003 | 1704 | LUZ19 T003 | 675 | + | - | 0 | 1877 | gp0.1 | |
| | LUZ19 TU007 | 1029 | LUZ19 P003 | 1768 | LUZ19 T004 | 739 | + | gp0.1 | 1 | 1877 | gp0.1 | |
| | LUZ19 TU008 | 1182 | LUZ19 P004 | 1704 | LUZ19 T003 | 522 | + | - | 0 | 2513 | gp0.1 \| gp1 \| gp2 | |
| | LUZ19 TU009 | 1182 | LUZ19 P004 | 1768 | LUZ19 T004 | 586 | + | gp0.1 | 1 | 2513 | gp0.1 \| gp1 \| gp2 | |

| ϕ | TU | TSS | TSS_ID | TTS | TTS_ID | TU length (bp) | +/- | Genes | #Genes | most distant 3' end | Genes in longest read with TSS | note |
|---|---|---|---|---|---|---|---|---|---|---|---|---|
| | LUZ19 TU010 | 1284 | LUZ19 P005 | 1704 | LUZ19 T003 | 420 | + | - | 0 | 1815 | gp0.1 | |
| | LUZ19 TU011 | 1284 | LUZ19 P005 | 1768 | LUZ19 T004 | 484 | + | gp0.1 | 1 | 1815 | gp0.1 | |
| | LUZ19 TU012 | 1561 | LUZ19 P006 | 1704 | LUZ19 T003 | 143 | + | - | 0 | 2852 | gp0.1 \| gp1 \| gp2 \| gp3 | |
| | LUZ19 TU013 | 1561 | LUZ19 P006 | 1768 | LUZ19 T004 | 207 | + | gp0.1 | 1 | 2852 | gp0.1 \| gp1 \| gp2 \| gp3 | |
| | LUZ19 TU014 | 1561 | LUZ19 P006 | 2851 | LUZ19 T005 | 1290 | + | gp0.1 \| gp1 \| gp2 \| gp3 | 4 | 2852 | gp0.1 \| gp1 \| gp2 \| gp3 | |
| | LUZ19 TU015 | 21134 | LUZ19 P007 | 21207 | LUZ19 T009 | 73 | + | - | 0 | 23931 | gp27 \| gp28 \| gp29 \| gp30 | |
| | LUZ19 TU016 | 21134 | LUZ19 P007 | 21415 | LUZ19 T010 | 281 | + | - | 0 | 23931 | gp27 \| gp28 \| gp29 \| gp30 | |
| | LUZ19 TU017 | 21134 | LUZ19 P007 | 22355 | LUZ19 T011 | 1221 | + | gp27 \| gp28 \| gp29 | 3 | 23931 | gp27 \| gp28 \| gp29 \| gp30 | |
| | LUZ19 TU018 | 21134 | LUZ19 P007 | 22491 | LUZ19 T012 | 1357 | + | gp27 \| gp28 \| gp29 | 3 | 23931 | gp27 \| gp28 \| gp29 \| gp30 | |
| | LUZ19 TU019 | 21134 | LUZ19 P007 | 22582 | LUZ19 T013 | 1448 | + | gp27 \| gp28 \| gp29 | 3 | 23931 | gp27 \| gp28 \| gp29 \| gp30 | |
| | LUZ19 TU020 | 21331 | LUZ19 P008 | 21415 | LUZ19 T010 | 84 | + | - | 0 | 23940 | gp28 \| gp29 \| gp30 | |
| | LUZ19 TU021 | 21331 | LUZ19 P008 | 22355 | LUZ19 T011 | 1024 | + | gp28 \| gp29 | 2 | 23940 | gp28 \| gp29 \| gp30 | |
| | LUZ19 TU022 | 21331 | LUZ19 P008 | 22491 | LUZ19 T012 | 1160 | + | gp28 \| gp29 | 2 | 23940 | gp28 \| gp29 \| gp30 | |

| ϕ | TU | TSS | TSS_ID | TTS | TTS_ID | TU length (bp) | +/- | Genes | #Genes | most distant 3' end | Genes in longest read with TSS | note |
|---|---|---|---|---|---|---|---|---|---|---|---|---|
| | LUZ19 TU023 | 21331 | LUZ19 P008 | 22582 | LUZ19 T013 | 1251 | + | gp28 \| gp29 | 2 | 23940 | gp28 \| gp29 \| gp30 | |
| | LUZ19 TU024 | 35609 | LUZ19 P009 | 36296 | LUZ19 T016 | 687 | + | - | 0 | 40006 | gp38 \| gp39 \| gp40 \| gp41 \| gp42 | |
| | LUZ19 TU025 | 35609 | LUZ19 P009 | 38768 | LUZ19 T017 | 3159 | + | gp38 \| gp39 \| gp40 | 3 | 40006 | gp38 \| gp39 \| gp40 \| gp41 \| gp42 | |
| LUZ24 | LUZ24 TU001 | 238 | LUZ24 P001 | 998 | LUZ24 T001 | 760 | + | - | 0 | 1743 | gp01 \| gp02 | |
| | LUZ24 TU002 | 331 | LUZ24 P002 | 998 | LUZ24 T001 | 667 | + | - | 0 | 1794 | gp01 \| gp02 | |
| | LUZ24 TU003 | 410 | LUZ24 P003 | 998 | LUZ24 T001 | 588 | + | - | 0 | 1282 | gp01 | |
| | LUZ24 TU004 | 554 | LUZ24 P004 | 998 | LUZ24 T001 | 444 | + | - | 0 | 2158 | gp01 \| gp02 | |
| | LUZ24 TU005 | 554 | LUZ24 P004 | 1872 | LUZ24 T002 | 1318 | + | gp01 \| gp02 | 2 | 2158 | gp01 \| gp02 | |
| | LUZ24 TU006 | 554 | LUZ24 P004 | 1941 | LUZ24 T003 | 1387 | + | gp01 \| gp02 | 2 | 2158 | gp01 \| gp02 | |
| | LUZ24 TU007 | 657 | LUZ24 P005 | 998 | LUZ24 T001 | 341 | + | - | 0 | 1789 | gp01 \| gp02 | |
| | LUZ24 TU008 | 4869 | LUZ24 P006 | 5415 | LUZ24 T006 | 546 | + | gp13 | 1 | 5588 | gp13 | |
| | LUZ24 TU009 | 5143 | LUZ24 P007 | 5415 | LUZ24 T006 | 272 | + | - | 0 | 5585 | - | |
| | LUZ24 TU010 | 5249 | LUZ24 P008 | 5415 | LUZ24 T006 | 166 | + | - | 0 | 5721 | - | |

| ϕ | TU | TSS | TSS_ID | TTS | TTS_ID | TU length (bp) | +/- | Genes | #Genes | most distant 3' end | Genes in longest read with TSS | note |
|---|---|---|---|---|---|---|---|---|---|---|---|---|
| | LUZ24 TU011 | 6630 | LUZ24 P009 | 7605 | LUZ24 T007 | 975 | + | - | 0 | 7604 | - | |
| | LUZ24 TU012 | 17248 | LUZ24 P010 | 17476 | LUZ24 T013 | 228 | + | - | 0 | 19070 | gp32 \| gp33 \| gp34 | |
| | LUZ24 TU013 | 17248 | LUZ24 P010 | 18243 | LUZ24 T014 | 995 | + | gp32 \| gp33 | 2 | 19070 | gp32 \| gp33 \| gp34 | |
| | LUZ24 TU014 | 19143 | LUZ24 P011 | 19811 | LUZ24 T016 | 668 | + | gp35 | 1 | 20987 | gp35 \| gp36 \| gp37 | associated with group I intron |
| | LUZ24 TU015 | 19143 | LUZ24 P011 | 20338 | LUZ24 T017 | 1195 | + | gp35 \| gp36 | 2 | 20987 | gp35 \| gp36 \| gp37 | associated with group I intron |
| | LUZ24 TU016 | 19143 | LUZ24 P011 | 20978 | LUZ24 T018 | 1835 | + | gp35 \| gp36 \| gp37 | 3 | 20987 | gp35 \| gp36 \| gp37 | associated with group I intron |
| | LUZ24 TU017 | 19353 | LUZ24 P012 | 19270 | LUZ24 T015 | 83 | - | - | 0 | 19267 | - | antisense |
| | LUZ24 TU018 | 25961 | LUZ24 P013 | 25726 | LUZ24 T020 | 235 | - | - | 0 | 25723 | - | |
| | LUZ24 TU019 | 28915 | LUZ24 P014 | 29129 | LUZ24 T021 | 214 | + | - | 0 | 30020 | - | antisense |
| | LUZ24 TU020 | 39133 | LUZ24 P015 | 37885 | LUZ24 T023 | 1248 | - | gp62 | 1 | 37883 | gp62 | |
| | LUZ24 TU021 | 39133 | LUZ24 P015 | 38968 | LUZ24 T024 | 165 | - | - | 0 | 37883 | - | |
| | LUZ24 TU022 | 44883 | LUZ24 P016 | 43329 | LUZ24 T025 | 1554 | - | gp67 \| gp68 | 2 | 43204 | gp67 \| gp68 | |

| ϕ | TU | TSS | TSS_ID | TTS | TTS_ID | TU length (bp) | +/- | Genes | #Genes | most distant 3' end | Genes in longest read with TSS | note |
|---|---|---|---|---|---|---|---|---|---|---|---|---|
| PAK_P3 | PAK_P3 TU001 | 100 | PAK_P3 P001 | 6419 | - | 6319 | + | gp1 \| gp2 \| gp3 \| gp4 \| gp5 \| gp6 \| gp7 | 7 | 6419 | gp1 \| gp2 \| gp3 \| gp4 \| gp5 \| gp6 \| gp7 | |
| | PAK_P3 TU002 | 300 | PAK_P3 P002 | 4381 | - | 4081 | + | gp2 \| gp3 \| gp34 | 3 | 4381 | gp2 \| gp3 \| gp34 | |
| | PAK_P3 TU003 | 768 | PAK_P3 P003 | 87643 | - | 1222 | - | PAK_P3as01 | 1 | 87641 (other side) | PAK_P3as01 | antisense |
| | PAK_P3 TU004 | 1638 | PAK_P3 P004 | 447 | - | 1191 | - | PAK_P3as02 | 1 | 447 | PAK_P3as02 | antisense |
| | PAK_P3 TU005 | 2898 | PAK_P3 P005 | 2627 | PAK_P3 T001 | 271 | - | - | 0 | 1227 | PAK_P3as03 | antisense |
| | PAK_P3 TU006 | 5470 | PAK_P3 P006 | 4406 | - | 1064 | - | - | 0 | 3613 | - | antisense |
| | PAK_P3 TU007 | 5814 | PAK_P3 P007 | 5673 | PAK_P3 T002 | 141 | - | - | 0 | 4406 | PAK_P3as04 | antisense |
| | PAK_P3 TU008 | 6167 | PAK_P3 P008 | 9540 | PAK_P3 T003 | 3373 | + | gp8 \| gp9 \| gp10 \| gp11 \| gp12 | 5 | 9541 | gp8 \| gp9 \| gp10 \| gp11 \| gp12 | |
| | PAK_P3 TU009 | 9603 | PAK_P3 P009 | 10211 | PAK_P3 T004 | 608 | + | gp13 | 1 | 15153 | gp13 \| gp14 \| gp15 \| gp16 \| gp17 \| gp18 \| gp19 | |
| | PAK_P3 TU010 | 9603 | PAK_P3 P009 | 11269 | PAK_P3 T005 | 1666 | + | gp13 \| gp14 \| gp15 \| gp16 | 4 | 15153 | gp13 \| gp14 \| gp15 \| gp16 \| gp17 \| gp18 \| gp19 | |
| | PAK_P3 TU011 | 9603 | PAK_P3 P009 | 13706 | PAK_P3 T006 | 4103 | + | gp13 \| gp14 \| gp15 \| gp16 \| gp17 | 5 | 15153 | gp13 \| gp14 \| gp15 \| gp16 \| gp17 \| gp18 \| gp19 | |
| | PAK_P3 TU012 | 9603 | PAK_P3 P009 | 13778 | PAK_P3 T007 | 4175 | + | gp13 \| gp14 \| gp15 \| gp16 \| gp17 | 5 | 15153 | gp13 \| gp14 \| gp15 \| gp16 \| gp17 \| gp18 \| gp19 | |
| | PAK_P3 TU013 | 12231 | PAK_P3 P010 | 13706 | PAK_P3 T006 | 1475 | + | - | 0 | 16193 | gp18 \| gp19 \| gp20 | |

| ϕ | TU | TSS | TSS_ID | TTS | TTS_ID | TU length (bp) | +/- | Genes | #Genes | most distant 3' end | Genes in longest read with TSS | note |
|---|---|---|---|---|---|---|---|---|---|---|---|---|
| | PAK_P3 TU014 | 12231 | PAK_P3 P010 | 13778 | PAK_P3 T007 | 1547 | + | - | 0 | 16193 | gp18 \| gp19 \| gp20 | |
| | PAK_P3 TU015 | 17986 | PAK_P3 P011 | 18374 | PAK_P3 T008 | 388 | + | - | 0 | 23221 | gp24 \| gp25 \| gp26 \| gp27 | |
| | PAK_P3 TU016 | 18530 | PAK_P3 P012 | 19591 | - | 1061 | + | - | 0 | 19591 | - | |
| | PAK_P3 TU017 | 19745 | PAK_P3 P013 | 18516 | - | 1229 | - | PAK_P3as11 | 1 | 18516 | PAK_P3as11 | antisense |
| | PAK_P3 TU018 | 21185 | PAK_P3 P014 | 19829 | - | 1356 | - | PAK_P3as13 | 1 | 19829 | PAK_P3as13 | antisense |
| | PAK_P3 TU019 | 23122 | PAK_P3 P015 | 23748 | PAK_P3 T009 | 626 | + | gp29 | 1 | 26024 | gp29 \| gp30 \| gp31 \| gp32 \| gp33 \| gp34 \| gp35 \| gp36 | |
| | PAK_P3 TU020 | 23122 | PAK_P3 P015 | 24869 | PAK_P3 T010 | 1747 | + | gp29 \| gp30 \| gp31 \| gp32 \| gp33 | 5 | 26024 | gp29 \| gp30 \| gp31 \| gp32 \| gp33 \| gp34 \| gp35 \| gp36 | |
| | PAK_P3 TU021 | 23122 | PAK_P3 P015 | 26024 | PAK_P3 T011 | 2902 | + | gp29 \| gp30 \| gp31 \| gp32 \| gp33 \| gp34 \| gp35 \| gp36 | 8 | 26024 | gp29 \| gp30 \| gp31 \| gp32 \| gp33 \| gp34 \| gp35 \| gp36 | |
| | PAK_P3 TU022 | 24923 | PAK_P3 P016 | 26024 | PAK_P3 T011 | 1101 | + | gp34 \| gp35 \| gp36 | 3 | 33391 | gp34 \| gp35 \| gp36 \| gp37 \| gp38 \| gp39 \| gp40 \| gp41 \| gp42 \| gp43 \| gp44 \| gp45 \| gp46 \| gp47 \| gp48 \| gp49 | |
| | PAK_P3 TU023 | 24923 | PAK_P3 P016 | 27936 | PAK_P3 T012 | 3013 | + | gp34 \| gp35 \| gp36 \| gp37 \| gp38 \| gp39 \| gp40 \| gp41 | 8 | 33391 | gp34 \| gp35 \| gp36 \| gp37 \| gp38 \| gp39 \| gp40 \| gp41 \| gp42 \| gp43 \| gp44 \| gp45 \| gp46 \| gp47 \| gp48 \| gp49 | |
| | PAK_P3 TU024 | 24923 | PAK_P3 P016 | 31308 | PAK_P3 T013 | 6385 | + | gp34 \| gp35 \| gp36 \| gp37 \| gp38 \| gp39 \| gp40 \| gp41 \| gp42 \| gp43 \| gp44 \| gp45 \| gp46 \| gp47 \| gp48 | 15 | 33391 | gp34 \| gp35 \| gp36 \| gp37 \| gp38 \| gp39 \| gp40 \| gp41 \| gp42 \| gp43 \| gp44 \| gp45 \| gp46 \| gp47 \| gp48 \| gp49 | |

| ϕ | TU | TSS | TSS_ID | TTS | TTS_ID | TU length (bp) | +/- | Genes | #Genes | most distant 3' end | Genes in longest read with TSS | note |
|---|---|---|---|---|---|---|---|---|---|---|---|---|
| | PAK_P3 TU025 | 24923 | PAK_P3 P016 | 33396 | PAK_P3 T015 | 8473 | + | gp34 \| gp35 \| gp36 \| gp37 \| gp38 \| gp39 \| gp40 \| gp41 \| gp42 \| gp43 \| gp44 \| gp45 \| gp46 \| gp47 \| gp48 \| gp49 | 16 | 33391 | gp34 \| gp35 \| gp36 \| gp37 \| gp38 \| gp39 \| gp40 \| gp41 \| gp42 \| gp43 \| gp44 \| gp45 \| gp46 \| gp47 \| gp48 \| gp49 | |
| | PAK_P3 TU026 | 25397 | PAK_P3 P017 | 26024 | PAK_P3 T011 | 627 | + | gp35 \| gp36 | 2 | 27937 | gp35 \| gp36 \| gp37 \| gp38 \| gp39 \| gp40 \| gp41 | |
| | PAK_P3 TU027 | 25397 | PAK_P3 P017 | 27936 | PAK_P3 T012 | 2539 | + | gp35 \| gp36 \| gp37 \| gp38 \| gp39 \| gp40 \| gp41 | 7 | 27937 | gp35 \| gp36 \| gp37 \| gp38 \| gp39 \| gp40 \| gp41 | |
| | PAK_P3 TU028 | 26977 | PAK_P3 P018 | 27936 | PAK_P3 T012 | 959 | + | gp41 | 1 | 34814 | gp41 \| gp42 \| gp43 \| gp44 \| gp45 \| gp46 \| gp47 \| gp48 \| gp49 | |
| | PAK_P3 TU029 | 26977 | PAK_P3 P018 | 31308 | PAK_P3 T013 | 4331 | + | gp41 \| gp42 \| gp43 \| gp44 \| gp45 \| gp46 \| gp47 \| gp48 | 8 | 34814 | gp41 \| gp42 \| gp43 \| gp44 \| gp45 \| gp46 \| gp47 \| gp48 \| gp49 | |
| | PAK_P3 TU030 | 26977 | PAK_P3 P018 | 33396 | PAK_P3 T015 | 6419 | + | gp41 \| gp42 \| gp43 \| gp44 \| gp45 \| gp46 \| gp47 \| gp48 \| gp49 | 9 | 34814 | gp41 \| gp42 \| gp43 \| gp44 \| gp45 \| gp46 \| gp47 \| gp48 \| gp49 | |
| | PAK_P3 TU031 | 29755 | PAK_P3 P019 | 31308 | PAK_P3 T013 | 1553 | + | gp45 \| gp46 \| gp47 \| gp48 | 4 | 32068 | gp45 \| gp46 \| gp47 \| gp48 | |
| | PAK_P3 TU032 | 29962 | PAK_P3 P020 | 31308 | PAK_P3 T013 | 1346 | + | gp46 \| gp47 \| gp48 | 3 | 36009 | gp46 \| gp47 \| gp48 \| gp49 \| gp50 | |
| | PAK_P3 TU033 | 29962 | PAK_P3 P020 | 33396 | PAK_P3 T015 | 3434 | + | gp46 \| gp47 \| gp48 \| gp49 | 4 | 36009 | gp46 \| gp47 \| gp48 \| gp49 \| gp50 | |
| | PAK_P3 TU034 | 30016 | PAK_P3 P021 | 31308 | PAK_P3 T013 | 1292 | + | gp46 \| gp47 \| gp48 | 3 | 31955 | gp46 \| gp47 \| gp48 | |
| | PAK_P3 TU035 | 30038 | PAK_P3 P022 | 31308 | PAK_P3 T013 | 1270 | + | gp46 \| gp47 \| gp48 | 3 | 33807 | gp46 \| gp47 \| gp48 \| gp49 | |
| | PAK_P3 TU036 | 30038 | PAK_P3 P022 | 33396 | PAK_P3 T015 | 3358 | + | gp46 \| gp47 \| gp48 \| gp49 | 4 | 33807 | gp46 \| gp47 \| gp48 \| gp49 | |

| ϕ | TU | TSS | TSS_ID | TTS | TTS_ID | TU length (bp) | +/- | Genes | #Genes | most distant 3' end | Genes in longest read with TSS | note |
|---|---|---|---|---|---|---|---|---|---|---|---|---|
| | PAK_P3 TU037 | 30858 | PAK_P3 P023 | 31308 | PAK_P3 T013 | 450 | + | gp48 | 1 | 34105 | gp48 \| gp49 | |
| | PAK_P3 TU038 | 31149 | PAK_P3 P024 | 31308 | PAK_P3 T013 | 159 | + | - | 0 | 37977 | gp49 \| gp50 \| gp51 \| gp52 \| gp53 \| gp54 \| gp55 | |
| | PAK_P3 TU039 | 31149 | PAK_P3 P024 | 33396 | PAK_P3 T015 | 2247 | + | gp49 | 1 | 37977 | gp49 \| gp50 \| gp51 \| gp52 \| gp53 \| gp54 \| gp55 | |
| | PAK_P3 TU040 | 31149 | PAK_P3 P024 | 37510 | PAK_P3 T017 | 6361 | + | gp49 \| gp50 \| gp51 \| gp52 \| gp53 | 5 | 37977 | gp49 \| gp50 \| gp51 \| gp52 \| gp53 \| gp54 \| gp55 | |
| | PAK_P3 TU041 | 31313 | PAK_P3 P025 | 33396 | PAK_P3 T015 | 2083 | + | gp49 | 1 | 37511 | gp49 \| gp50 \| gp51 \| gp52 \| gp53 | |
| | PAK_P3 TU042 | 31313 | PAK_P3 P025 | 37510 | PAK_P3 T017 | 6197 | + | gp49 \| gp50 \| gp51 \| gp52 \| gp53 | 5 | 37511 | gp49 \| gp50 \| gp51 \| gp52 \| gp53 | |
| | PAK_P3 TU043 | 31341 | PAK_P3 P026 | 33396 | PAK_P3 T015 | 2055 | + | gp49 | 1 | 35918 | gp49 \| gp50 | |
| | PAK_P3 TU044 | 36310 | PAK_P3 P027 | 37510 | PAK_P3 T017 | 1200 | + | gp52 \| gp53 | 2 | 40988 | gp52 \| gp53 \| gp54 \| gp55 \| gp56 \| gp57 \| gp58 \| gp59 \| gp60 \| gp61 | |
| | PAK_P3 TU045 | 41196 | PAK_P3 P028 | 41420 | PAK_P3 T018 | 224 | + | - | 0 | 42925 | gp63 \| gp64 \| gp65 | |
| | PAK_P3 TU046 | 42029 | PAK_P3 P029 | 45611 | PAK_P3 T019 | 3582 | + | gp65 \| gp66 \| gp67 \| gp68 | 4 | 46686 | gp65 \| gp66 \| gp67 \| gp68 \| gp69 | |
| | PAK_P3 TU047 | 44517 | PAK_P3 P030 | 45611 | PAK_P3 T019 | 1094 | + | gp68 | 1 | 46689 | gp68 \| gp69 | |
| | PAK_P3 TU048 | 46660 | PAK_P3 P031 | 47943 | PAK_P3 T020 | 1283 | + | gp70 \| gp71 \| gp72 \| gp73 | 4 | 50630 | gp70 \| gp71 \| gp72 \| gp73 \| gp74 \| gp75 \| gp76 \| gp77 \| gp78 \| gp79 \| gp80 | |

| ϕ | TU | TSS | TSS_ID | TTS | TTS_ID | TU length (bp) | +/- | Genes | #Genes | most distant 3' end | Genes in longest read with TSS | note |
|---|---|---|---|---|---|---|---|---|---|---|---|---|
| | PAK_P3 TU049 | 46660 | PAK_P3 P031 | 50279 | PAK_P3 T021 | 3619 | + | gp70 \| gp71 \| gp72 \| gp73 \| gp74 \| gp75 \| gp76 \| gp77 \| gp78 \| gp79 | 10 | 50630 | gp70 \| gp71 \| gp72 \| gp73 \| gp74 \| gp75 \| gp76 \| gp77 \| gp78 \| gp79 \| gp80 | |
| | PAK_P3 TU050 | 46660 | PAK_P3 P031 | 50625 | PAK_P3 T022 | 3965 | + | gp70 \| gp71 \| gp72 \| gp73 \| gp74 \| gp75 \| gp76 \| gp77 \| gp78 \| gp79 \| gp80 | 11 | 50630 | gp70 \| gp71 \| gp72 \| gp73 \| gp74 \| gp75 \| gp76 \| gp77 \| gp78 \| gp79 \| gp80 | |
| | PAK_P3 TU051 | 47278 | PAK_P3 P032 | 47943 | PAK_P3 T020 | 665 | + | gp72 \| gp73 | 2 | 50872 | gp72 \| gp73 \| gp74 \| gp75 \| gp76 \| gp77 \| gp78 \| gp79 \| gp80 | |
| | PAK_P3 TU052 | 47278 | PAK_P3 P032 | 50279 | PAK_P3 T021 | 3001 | + | gp72 \| gp73 \| gp74 \| gp75 \| gp76 \| gp77 \| gp78 \| gp79 | 8 | 50872 | gp72 \| gp73 \| gp74 \| gp75 \| gp76 \| gp77 \| gp78 \| gp79 \| gp80 | |
| | PAK_P3 TU053 | 47278 | PAK_P3 P032 | 50625 | PAK_P3 T022 | 3347 | + | gp72 \| gp73 \| gp74 \| gp75 \| gp76 \| gp77 \| gp78 \| gp79 \| gp80 | 9 | 50872 | gp72 \| gp73 \| gp74 \| gp75 \| gp76 \| gp77 \| gp78 \| gp79 \| gp80 | |
| | PAK_P3 TU054 | 47939 | PAK_P3 P033 | 50279 | PAK_P3 T021 | 2340 | + | gp74 \| gp75 \| gp76 \| gp77 \| gp78 \| gp79 | 6 | 50787 | gp74 \| gp75 \| gp76 \| gp77 \| gp78 \| gp79 \| gp80 | |
| | PAK_P3 TU055 | 47939 | PAK_P3 P033 | 50625 | PAK_P3 T022 | 2686 | + | gp74 \| gp75 \| gp76 \| gp77 \| gp78 \| gp79 \| gp80 | 7 | 50787 | gp74 \| gp75 \| gp76 \| gp77 \| gp78 \| gp79 \| gp80 | |
| | PAK_P3 TU056 | 48439 | PAK_P3 P034 | 50279 | PAK_P3 T021 | 1840 | + | gp75 \| gp76 \| gp77 \| gp78 \| gp79 | 5 | 50897 | gp75 \| gp76 \| gp77 \| gp78 \| gp79 \| gp80 | |
| | PAK_P3 TU057 | 48439 | PAK_P3 P034 | 50625 | PAK_P3 T022 | 2186 | + | gp75 \| gp76 \| gp77 \| gp78 \| gp79 \| gp80 | 6 | 50897 | gp75 \| gp76 \| gp77 \| gp78 \| gp79 \| gp80 | |
| | PAK_P3 TU058 | 51049 | PAK_P3 P035 | 52047 | PAK_P3 T023 | 998 | + | gp81 \| gp82 \| gp83 \| gp84 | 4 | 54283 | gp81 \| gp82 \| gp83 \| gp84 \| gp85 \| gp86 \| gp87 \| gp88 \| gp89 \| gp90 | |
| | PAK_P3 TU059 | 51049 | PAK_P3 P035 | 53146 | PAK_P3 T024 | 2097 | + | gp81 \| gp82 \| gp83 \| gp84 \| gp85 \| gp86 | 6 | 54283 | gp81 \| gp82 \| gp83 \| gp84 \| gp85 \| gp86 \| gp87 \| gp88 \| gp89 \| gp90 | |

| ϕ | TU | TSS | TSS_ID | TTS | TTS_ID | TU length (bp) | +/- | Genes | #Genes | most distant 3' end | Genes in longest read with TSS | note |
|---|---|---|---|---|---|---|---|---|---|---|---|---|
| | PAK_P3 TU060 | 51049 | PAK_P3 P035 | 53749 | PAK_P3 T025 | 2700 | + | gp81 \| gp82 \| gp83 \| gp84 \| gp85 \| gp86 \| gp87 \| gp88 | 8 | 54283 | gp81 \| gp82 \| gp83 \| gp84 \| gp85 \| gp86 \| gp87 \| gp88 \| gp89 \| gp90 | |
| | PAK_P3 TU061 | 51771 | PAK_P3 P036 | 52047 | PAK_P3 T023 | 276 | + | gp84 | 1 | 54356 | gp84 \| gp85 \| gp86 \| gp87 \| gp88 \| gp89 \| gp90 | |
| | PAK_P3 TU062 | 51771 | PAK_P3 P036 | 53146 | PAK_P3 T024 | 1375 | + | gp84 \| gp85 \| g86 | 3 | 54356 | gp84 \| gp85 \| gp86 \| gp87 \| gp88 \| gp89 \| gp90 | |
| | PAK_P3 TU063 | 51771 | PAK_P3 P036 | 53749 | PAK_P3 T025 | 1978 | + | gp84 \| gp85 \| gp86 \| gp87 \| gp88 | 5 | 54356 | gp84 \| gp85 \| gp86 \| gp87 \| gp88 \| gp89 \| gp90 | |
| | PAK_P3 TU064 | 51771 | PAK_P3 P036 | 54353 | PAK_P3 T026 | 2582 | + | gp84 \| gp85 \| gp86 \| gp87 \| gp88 \| gp89 \| gp90 | 7 | 54356 | gp84 \| gp85 \| gp86 \| gp87 \| gp88 \| gp89 \| gp90 | |
| | PAK_P3 TU065 | 51944 | PAK_P3 P037 | 52047 | PAK_P3 T023 | 103 | + | - | 0 | 54437 | gp85 \| gp86 \| gp87 \| gp88 \| gp89 \| gp90 | |
| | PAK_P3 TU066 | 51944 | PAK_P3 P037 | 53146 | PAK_P3 T024 | 1202 | + | gp85 \| gp86 | 2 | 54437 | gp85 \| gp86 \| gp87 \| gp88 \| gp89 \| gp90 | |
| | PAK_P3 TU067 | 51944 | PAK_P3 P037 | 53749 | PAK_P3 T025 | 1805 | + | gp85 \| gp86 \| gp87 \| gp88 | 4 | 54437 | gp85 \| gp86 \| gp87 \| gp88 \| gp89 \| gp90 | |
| | PAK_P3 TU068 | 51944 | PAK_P3 P037 | 54353 | PAK_P3 T026 | 2409 | + | gp85 \| gp86 \| gp87 \| gp88 \| gp89 \| gp90 | 6 | 54437 | gp85 \| gp86 \| gp87 \| gp88 \| gp89 \| gp90 | |
| | PAK_P3 TU069 | 52664 | PAK_P3 P038 | 53146 | PAK_P3 T024 | 482 | + | gp86 | 1 | 55204 | gp86 \| gp87 \| gp88 \| gp89 \| gp90 \| gp91 \| gp92 \| gp93 | |
| | PAK_P3 TU070 | 52664 | PAK_P3 P038 | 53749 | PAK_P3 T025 | 1085 | + | gp86 \| gp87 \| gp88 | 3 | 55204 | gp86 \| gp87 \| gp88 \| gp89 \| gp90 \| gp91 \| gp92 \| gp93 | |
| | PAK_P3 TU071 | 52664 | PAK_P3 P038 | 54353 | PAK_P3 T026 | 1689 | + | gp86 \| gp87 \| gp88 \| gp89 \| gp90 | 5 | 55204 | gp86 \| gp87 \| gp88 \| gp89 \| gp90 \| gp91 \| gp92 \| gp93 | |
| | PAK_P3 TU072 | 52664 | PAK_P3 P038 | 55204 | PAK_P3 T027 | 2540 | + | gp86 \| gp87 \| gp88 \| gp89 \| gp90 \| gp91 \| gp92 \| gp93 | 8 | 55204 | gp86 \| gp87 \| gp88 \| gp89 \| gp90 \| gp91 \| gp92 \| gp93 | |

| ϕ | TU | TSS | TSS_ID | TTS | TTS_ID | TU length (bp) | +/- | Genes | #Genes | most distant 3' end | Genes in longest read with TSS | note |
|---|---|---|---|---|---|---|---|---|---|---|---|---|
| | PAK_P3 TU073 | 53142 | PAK_P3 P039 | 54353 | PAK_P3 T026 | 1211 | + | gp87 \| gp88 \| gp89 \| gp90 | 4 | 54491 | gp87 \| gp88 \| gp89 \| gp90 | |
| | PAK_P3 TU074 | 54361 | PAK_P3 P040 | 55204 | PAK_P3 T027 | 843 | + | gp91 \| gp92 \| gp93 | 3 | 56874 | gp91 \| gp92 \| gp93 \| gp94 \| gp95 \| gp96 | |
| | PAK_P3 TU075 | 55259 | PAK_P3 P041 | 58608 | PAK_P3 T028 | 3349 | + | gp94 \| gp95 \| gp96 \| gp97 \| gp98 \| gp99 \| gp100 | 7 | 58677 | gp94 \| gp95 \| gp96 \| gp97 \| gp98 \| gp99 \| gp100 | |
| | PAK_P3 TU076 | 55652 | PAK_P3 P042 | 58608 | PAK_P3 T028 | 2956 | + | gp95 \| gp96 \| gp97 \| gp98 \| gp99 \| gp100 | 6 | 58608 | gp95 \| gp96 \| gp97 \| gp98 \| gp99 \| gp100 | |
| | PAK_P3 TU077 | 57078 | PAK_P3 P043 | 58608 | PAK_P3 T028 | 1530 | + | gp98 \| gp99 \| gp100 | 3 | 60630 | gp98 \| gp99 \| gp100 \| gp101 \| gp102 \| gp103 \| gp104 \| gp105 | |
| | PAK_P3 TU078 | 57078 | PAK_P3 P043 | 59378 | PAK_P3 T029 | 2300 | + | gp98 \| gp99 \| gp100 \| gp101 \| gp102 | 5 | 60630 | gp98 \| gp99 \| gp100 \| gp101 \| gp102 \| gp103 \| gp104 \| gp105 | |
| | PAK_P3 TU079 | 57078 | PAK_P3 P043 | 59945 | PAK_P3 T030 | 2867 | + | gp98 \| gp99 \| gp100 \| gp101 \| gp102 \| gp103 | 6 | 60630 | gp98 \| gp99 \| gp100 \| gp101 \| gp102 \| gp103 \| gp104 \| gp105 | |
| | PAK_P3 TU080 | 57408 | PAK_P3 P044 | 58608 | PAK_P3 T028 | 1200 | + | gp99 \| gp100 | 2 | 60270 | gp99 \| gp100 \| gp101 \| gp102 \| gp103 \| gp104 | |
| | PAK_P3 TU081 | 58020 | PAK_P3 P045 | 58608 | PAK_P3 T028 | 588 | + | gp100 | 1 | 60634 | gp100 \| gp101 \| gp102 \| gp103 \| gp104 \| gp105 | |
| | PAK_P3 TU082 | 58656 | PAK_P3 P046 | 59378 | PAK_P3 T029 | 722 | + | gp101 \| gp102 | 2 | 61660 | gp101 \| gp102 \| gp103 \| gp104 \| gp105 \| gp106 \| gp107 \| gp108 | |
| | PAK_P3 TU083 | 58656 | PAK_P3 P046 | 59945 | PAK_P3 T030 | 1289 | + | gp101 \| gp102 \| gp103 | 3 | 61660 | gp101 \| gp102 \| gp103 \| gp104 \| gp105 \| gp106 \| gp107 \| gp108 | |
| | PAK_P3 TU084 | 58850 | PAK_P3 P047 | 59378 | PAK_P3 T029 | 528 | + | gp102 | 1 | 62440 | gp102 \| gp103 \| gp104 \| gp105 \| gp106 \| gp107 \| | |

| ϕ | TU | TSS | TSS_ID | TTS | TTS_ID | TU length (bp) | +/- | Genes | #Genes | most distant 3' end | Genes in longest read with TSS | note |
|---|---|---|---|---|---|---|---|---|---|---|---|---|
| | | | | | | | | | | | gp108 \| gp109 \| gp110 \| gp111 | |
| | PAK_P3 TU085 | 58850 | PAK_P3 P047 | 59945 | PAK_P3 T030 | 1095 | + | gp102 \| gp103 | 2 | 62440 | gp102 \| gp103 \| gp104 \| gp105 \| gp106 \| gp107 \| gp108 \| gp109 \| gp110 \| gp111 | |
| | PAK_P3 TU086 | 58850 | PAK_P3 P047 | 61972 | PAK_P3 T031 | 3122 | + | gp102 \| gp103 \| gp104 \| gp105 \| gp106 \| gp107 \| gp108 \| gp109 | 8 | 62440 | gp102 \| gp103 \| gp104 \| gp105 \| gp106 \| gp107 \| gp108 \| gp109 \| gp110 \| gp111 | |
| | PAK_P3 TU087 | 58993 | PAK_P3 P048 | 59378 | PAK_P3 T029 | 385 | + | - | 0 | 62661 | gp103 \| gp104 \| gp105 \| gp106 \| gp107 \| gp108 \| gp109 \| gp110 \| gp111 | |
| | PAK_P3 TU088 | 58993 | PAK_P3 P048 | 59945 | PAK_P3 T030 | 952 | + | gp103 | 1 | 62661 | gp103 \| gp104 \| gp105 \| gp106 \| gp107 \| gp108 \| gp109 \| gp110 \| gp111 | |
| | PAK_P3 TU089 | 58993 | PAK_P3 P048 | 61972 | PAK_P3 T031 | 2979 | + | gp103 \| gp104 \| gp105 \| gp106 \| gp107 \| gp108 \| gp109 | 7 | 62661 | gp103 \| gp104 \| gp105 \| gp106 \| gp107 \| gp108 \| gp109 \| gp110 \| gp111 | |
| | PAK_P3 TU090 | 59369 | PAK_P3 P049 | 59945 | PAK_P3 T030 | 576 | + | gp103 | 1 | 62885 | gp103 \| gp104 \| gp105 \| gp106 \| gp107 \| gp108 \| gp109 \| gp110 \| gp111 \| gp112 | |
| | PAK_P3 TU091 | 59369 | PAK_P3 P049 | 61972 | PAK_P3 T031 | 2603 | + | gp103 \| gp104 \| gp105 \| gp106 \| gp107 \| gp108 \| gp109 | 7 | 62885 | gp103 \| gp104 \| gp105 \| gp106 \| gp107 \| gp108 \| gp109 \| gp110 \| gp111 \| gp112 | |
| | PAK_P3 TU092 | 59950 | PAK_P3 P050 | 61972 | PAK_P3 T031 | 2022 | + | gp104 \| gp105 \| gp106 \| gp107 \| gp108 \| gp109 | 6 | 62661 | gp104 \| gp105 \| gp106 \| gp107 \| gp108 \| gp109 \| gp110 \| gp111 | |

| ϕ | TU | TSS | TSS_ID | TTS | TTS_ID | TU length (bp) | +/- | Genes | #Genes | most distant 3' end | Genes in longest read with TSS | note |
|---|---|---|---|---|---|---|---|---|---|---|---|---|
| | PAK_P3 TU093 | 60536 | PAK_P3 P051 | 61972 | PAK_P3 T031 | 1436 | + | gp106 \| gp107 \| gp108 \| gp109 | 4 | 62729 | gp106 \| gp107 \| gp108 \| gp109 \| gp110 \| gp111 | |
| | PAK_P3 TU094 | 61068 | PAK_P3 P052 | 61972 | PAK_P3 T031 | 904 | + | gp107 \| gp108 \| gp109 | 3 | 62684 | gp107 \| gp108 \| gp109 \| gp110 \| gp111 | |
| | PAK_P3 TU095 | 63712 | PAK_P3 P053 | 67467 | - | 3755 | + | gp114 \| gp115 \| gp116 \| gp117 \| gp118 \| gp119 \| gp120 \| gp121 | 8 | 67467 | gp114 \| gp115 \| gp116 \| gp117 \| gp118 \| gp119 \| gp120 \| gp121 | |
| | PAK_P3 TU096 | 63855 | PAK_P3 P054 | 66228 | - | 2373 | + | gp115 \| gp116 \| gp117 \| gp118 | 4 | 65865 | gp115 \| gp116 \| gp117 \| gp118 | |
| | PAK_P3 TU097 | 64001 | PAK_P3 P055 | 66453 | - | 2452 | + | gp115 \| gp116 \| gp117 \| gp118 \| gp119 | 5 | 66453 | gp115 \| gp116 \| gp117 \| gp118 \| gp119 | |
| | PAK_P3 TU098 | 64117 | PAK_P3 P056 | 68322 | - | 4205 | + | gp115 \| gp116 \| gp117 \| gp118 \| gp119 \| gp120 \| gp121 \| gp122 \| gp123 | 9 | 68322 | gp115 \| gp116 \| gp117 \| gp118 \| gp119 \| gp120 \| gp121 \| gp122 \| gp123 | |
| | PAK_P3 TU099 | 65172 | PAK_P3 P057 | 68654 | PAK_P3 T032 | 3482 | + | gp118 \| gp119 \| gp120 \| gp121 \| gp122 \| gp123 \| gp124 \| gp125 | 8 | 69604 | gp118 \| gp119 \| gp120 \| gp121 \| gp122 \| gp123 \| gp124 \| gp125 \| gp126 | |
| | PAK_P3 TU100 | 66422 | PAK_P3 P058 | 68654 | PAK_P3 T032 | 2232 | + | gp120 \| gp121 \| gp122 \| gp123 \| gp124 \| gp125 | 6 | 69642 | gp120 \| gp121 \| gp122 \| gp123 \| gp124 \| gp125 \| gp126 | |
| | PAK_P3 TU101 | 67450 | PAK_P3 P059 | 68654 | PAK_P3 T032 | 1204 | + | gp122 \| gp123 \| gp124 \| gp125 | 4 | 69628 | gp122 \| gp123 \| gp124 \| gp125 \| gp126 | |
| | PAK_P3 TU102 | 70777 | PAK_P3 P060 | 72389 | PAK_P3 T033 | 1612 | + | gp130 \| gp131 \| gp132 \| gp133 \| gp166 | 5 | 73240 | gp130 \| gp131 \| gp132 \| gp133 \| gp166 \| gp134 \| gp135 \| gp136 | |
| | PAK_P3 TU103 | 70777 | PAK_P3 P060 | 73240 | PAK_P3 T034 | 2463 | + | gp130 \| gp131 \| gp132 \| gp133 \| gp166 \| gp134 \| gp135 \| gp136 | 8 | 73240 | gp130 \| gp131 \| gp132 \| gp133 \| gp166 \| gp134 \| gp135 \| gp136 | |

| ϕ | TU | TSS | TSS_ID | TTS | TTS_ID | TU length (bp) | +/- | Genes | #Genes | most distant 3' end | Genes in longest read with TSS | note |
|---|---|---|---|---|---|---|---|---|---|---|---|---|
| | PAK_P3 TU104 | 72404 | PAK_P3 P061 | 73240 | PAK_P3 T034 | 836 | + | gp134 \| gp135 \| gp136 | 3 | 73412 | gp134 \| gp135 \| gp136 | |
| | PAK_P3 TU105 | 72475 | PAK_P3 P062 | 73240 | PAK_P3 T034 | 765 | + | gp135 \| gp136 | 2 | 73243 | gp135 \| gp136 | |
| | PAK_P3 TU106 | 78100 | PAK_P3 P063 | 73423 | PAK_P3 T035 | 4677 | - | gp137 \| gp138 \| gp139 \| gp140 \| gp141 \| gp142 \| gp143 \| gp144 \| gp145 \| gp146 \| gp147 | 11 | 73633 | gp137 \| gp138 \| gp139 \| gp140 \| gp141 \| gp142 \| gp143 \| gp144 \| gp145 \| gp146 \| gp147 | |
| | PAK_P3 TU107 | 78951 | PAK_P3 P064 | 78328 | PAK_P3 T036 | 623 | - | - | 0 | 75238 | gp143 \| gp144 \| gp145 \| gp146 \| gp147 \| gp148 | |
| | PAK_P3 TU108 | 79497 | PAK_P3 P065 | 78328 | PAK_P3 T036 | 1169 | - | gp149 \| gp150 | 2 | 74800 | gp141 \| gp142 \| gp143 \| gp144 \| gp145 \| gp146 \| gp147 \| gp148 \| gp149 \| gp150 | |
| | PAK_P3 TU109 | 80430 | PAK_P3 P066 | 78328 | PAK_P3 T036 | 2102 | - | gp149 \| gp150 \| gp151 \| gp152 | 4 | 77628 | gp148 \| gp149 \| gp150 \| gp151 \| gp152 | |
| | PAK_P3 TU110 | 82342 | PAK_P3 P067 | 78872 | - | 3470 | - | gp150 \| gp151 \| gp152 \| gp153 \| gp154 | 5 | 78872 | gp150 \| gp151 \| gp152 \| gp153 \| gp154 | |
| | PAK_P3 TU111 | 83179 | PAK_P3 P068 | 78328 | PAK_P3 T036 | 4851 | - | gp149 \| gp150 \| gp151 \| gp152 \| gp153 \| gp154 \| gp155 \| gp156 \| gp157 | 9 | 74811 | gp141 \| gp142 \| gp143 \| gp144 \| gp145 \| gp146 \| gp147 \| gp148 \| gp149 \| gp150 \| gp151 \| gp152 \| gp153 \| gp154 \| gp155 \| gp156 \| gp157 | |
| | PAK_P3 TU112 | 84247 | PAK_P3 P069 | 83230 | PAK_P3 T037 | 1017 | - | gp158 \| gp159 | 2 | 83017 | gp158 \| gp159 | |
| | PAK_P3 TU113 | 84247 | PAK_P3 P069 | 83691 | PAK_P3 T038 | 556 | - | gp159 | 1 | 83017 | gp158 \| gp159 | |
| | PAK_P3 TU114 | 84321 | PAK_P3 P070 | 85458 | PAK_P3 T039 | 1137 | + | gp160 \| gp161 \| gp162 \| PAK_P3sRNA1 | 4 | 86928 | gp160 \| gp161 \| gp162 \| PAK_P3sRNA1 \| gp163 \| | |

| ϕ | TU | TSS | TSS_ID | TTS | TTS_ID | TU length (bp) | +/- | Genes | #Genes | most distant 3' end | Genes in longest read with TSS | note |
|---|---|---|---|---|---|---|---|---|---|---|---|---|
| | | | | | | | | | | | PAK_P3tRNA01 \| PAK_P3sRNA2 \| PAK_P3tRNA02 | |
| | PAK_P3 TU115 | 84321 | PAK_P3 P070 | 86017 | PAK_P3 T040 | 1696 | + | gp160 \| gp161 \| gp162 \| PAK_P3sRNA1 \| gp163 | 5 | 86928 | gp160 \| gp161 \| gp162 \| PAK_P3sRNA1 \| gp163 \| PAK_P3tRNA01 \| PAK_P3sRNA2 \| PAK_P3tRNA02 | |
| | PAK_P3 TU116 | 84633 | PAK_P3 P071 | 85458 | PAK_P3 T039 | 825 | + | gp161 \| gp162 \| PAK_P3sRNA1 | 3 | 88097 | gp161 \| gp162 \| PAK_P3sRNA1 \| gp163 \| PAK_P3tRNA01 \| PAK_P3sRNA2 \| PAK_P3tRNA02 \| gp164 \| gp165 | |
| | PAK_P3 TU117 | 84633 | PAK_P3 P071 | 86017 | PAK_P3 T040 | 1384 | + | gp161 \| gp162 \| PAK_P3sRNA1 \| gp163 | 4 | 88097 | gp161 \| gp162 \| PAK_P3sRNA1 \| gp163 \| PAK_P3tRNA01 \| PAK_P3sRNA2 \| PAK_P3tRNA02 \| gp164 \| gp165 | |
| | PAK_P3 TU118 | 84633 | PAK_P3 P071 | 86426 | PAK_P3 T041 | 1793 | + | gp161 \| gp162 \| PAK_P3sRNA1 \| gp163 \| PAK_P3tRNA01 | 5 | 88097 | gp161 \| gp162 \| PAK_P3sRNA1 \| gp163 \| PAK_P3tRNA01 \| PAK_P3sRNA2 \| PAK_P3tRNA02 \| gp164 \| gp165 | |
| | PAK_P3 TU119 | 84633 | PAK_P3 P071 | 86498 | PAK_P3 T042 | 1865 | + | gp161 \| gp162 \| PAK_P3sRNA1 \| gp163 \| PAK_P3tRNA01 \| PAK_P3sRNA2 | 6 | 88097 | gp161 \| gp162 \| PAK_P3sRNA1 \| gp163 \| PAK_P3tRNA01 \| PAK_P3sRNA2 \| PAK_P3tRNA02 \| gp164 \| gp165 | |
| | PAK_P3 TU120 | 84633 | PAK_P3 P071 | 86669 | PAK_P3 T043 | 2036 | + | gp161 \| gp162 \| PAK_P3sRNA1 \| gp163 \| PAK_P3tRNA01 \| PAK_P3sRNA2 \| PAK_P3tRNA02 | 7 | 88097 | gp161 \| gp162 \| PAK_P3sRNA1 \| gp163 \| PAK_P3tRNA01 \| PAK_P3sRNA2 \| PAK_P3tRNA02 \| gp164 \| gp165 | |

| ϕ | TU | TSS | TSS_ID | TTS | TTS_ID | TU length (bp) | +/- | Genes | #Genes | most distant 3' end | Genes in longest read with TSS | note |
|---|---|---|---|---|---|---|---|---|---|---|---|---|
| | PAK_P3 TU121 | 85495 | PAK_P3 P072 | 86017 | PAK_P3 T040 | 522 | + | gp163 | 1 | 87273 | gp163 \| PAK_P3tRNA01 \| PAK_P3sRNA2 \| PAK_P3tRNA02 | |
| | PAK_P3 TU122 | 85495 | PAK_P3 P072 | 86426 | PAK_P3 T041 | 931 | + | gp163 \| PAK_P3tRNA01 | 2 | 87273 | gp163 \| PAK_P3tRNA01 \| PAK_P3sRNA2 \| PAK_P3tRNA02 | |
| | PAK_P3 TU123 | 85495 | PAK_P3 P072 | 86498 | PAK_P3 T042 | 1003 | + | gp163 \| PAK_P3tRNA01 \| PAK_P3sRNA2 | 3 | 87273 | gp163 \| PAK_P3tRNA01 \| PAK_P3sRNA2 \| PAK_P3tRNA02 | |
| | PAK_P3 TU124 | 85495 | PAK_P3 P072 | 86669 | PAK_P3 T043 | 1174 | + | gp163 \| PAK_P3tRNA01 \| PAK_P3sRNA2 \| PAK_P3tRNA02 | 4 | 87273 | gp163 \| PAK_P3tRNA01 \| PAK_P3sRNA2 \| PAK_P3tRNA02 | |
| | PAK_P3 TU125 | 85931 | PAK_P3 P073 | 86017 | PAK_P3 T040 | 86 | + | - | 0 | 88097 | PAK_P3tRNA01 \| PAK_P3sRNA2 \| PAK_P3tRNA02 \| gp164 \| gp165 | |
| | PAK_P3 TU126 | 85931 | PAK_P3 P073 | 86426 | PAK_P3 T041 | 495 | + | PAK_P3tRNA01 | 1 | 88097 | PAK_P3tRNA01 \| PAK_P3sRNA2 \| PAK_P3tRNA02 \| gp164 \| gp165 | |
| | PAK_P3 TU127 | 85931 | PAK_P3 P073 | 86498 | PAK_P3 T042 | 567 | + | PAK_P3tRNA01 \| PAK_P3sRNA2 | 2 | 88097 | PAK_P3tRNA01 \| PAK_P3sRNA2 \| PAK_P3tRNA02 \| gp164 \| gp165 | |
| | PAK_P3 TU128 | 85931 | PAK_P3 P073 | 86669 | PAK_P3 T043 | 738 | + | PAK_P3tRNA01 \| PAK_P3sRNA2 \| PAK_P3tRNA02 | 3 | 88097 | PAK_P3tRNA01 \| PAK_P3sRNA2 \| PAK_P3tRNA02 \| gp164 \| gp165 | |
| | PAK_P3 TU129 | 86865 | PAK_P3 P074 | 86468 | - | 397 | - | PAK_P3as20 | 1 | 86468 | PAK_P3as20 | antisense |

| ϕ | TU | TSS | TSS_ID | TTS | TTS_ID | TU length (bp) | +/- | Genes | #Genes | most distant 3' end | Genes in longest read with TSS | note |
|---|---|---|---|---|---|---|---|---|---|---|---|---|
| | PAK_P3 TU130 | 88067 | PAK_P3 P075 | 5014 | - | 5044 | + | gp1 \| gp2 \| gp3 \| gp4 \| gp5 | 5 | 5014 (other side) | gp1 \| gp2 \| gp3 \| gp4 \| gp5 | |
| phiKZ | phiKZ TU001 | 2176 | phiKZ P001 | 3756 | phiKZ T001 | 1580 | + | PHIKZ005 \| PHIKZ006 \| PHIKZ007 \| PHIKZ008 | 4 | 5773 | PHIKZ005 \| PHIKZ006 \| PHIKZ007 \| PHIKZ008 \| PHIKZ_p01 \| PHIKZ010 \| phiKZ_p02 \| PHIKZ011 \| PHIKZ012 | |
| | phiKZ TU002 | 2176 | phiKZ P001 | 5540 | phiKZ T002 | 3364 | + | PHIKZ005 \| PHIKZ006 \| PHIKZ007 \| PHIKZ008 \| PHIKZ_p01 \| PHIKZ010 \| phiKZ_p02 \| PHIKZ011 \| PHIKZ012 | 9 | 5773 | PHIKZ005 \| PHIKZ006 \| PHIKZ007 \| PHIKZ008 \| PHIKZ_p01 \| PHIKZ010 \| phiKZ_p02 \| PHIKZ011 \| PHIKZ012 | |
| | phiKZ TU003 | 4180 | phiKZ P002 | 5540 | phiKZ T002 | 1360 | + | PHIKZ010 \| phiKZ_p02 \| PHIKZ011 \| PHIKZ012 | 4 | 7354 | PHIKZ010 \| phiKZ_p02 \| PHIKZ011 \| PHIKZ012 \| PHIKZ013 \| PHIKZ014 | |
| | phiKZ TU004 | 10695 | phiKZ P003 | 12204 | - | 1509 | + | PHIKZ022 \| PHIKZ023 | 2 | 12204 | PHIKZ022 \| PHIKZ023 | |
| | phiKZ TU005 | 12388 | phiKZ P004 | 12011 | phiKZ T003 | 377 | - | PHIKZ_p04 | 1 | 8970 | PHIKZ_p04 | |
| | phiKZ TU006 | 12388 | phiKZ P004 | 12168 | phiKZ T004 | 220 | - | PHIKZ_p04 | 1 | 8970 | PHIKZ_p04 | |
| | phiKZ TU007 | 12482 | phiKZ P005 | 12011 | phiKZ T003 | 471 | - | PHIKZ_p04 | 1 | 7381 | PHIKZ_p04 | |
| | phiKZ TU008 | 12482 | phiKZ P005 | 12168 | phiKZ T004 | 314 | - | PHIKZ_p04 | 1 | 7381 | PHIKZ_p04 | |
| | phiKZ TU009 | 14812 | phiKZ P006 | 12011 | phiKZ T003 | 2801 | - | PHIKZ_p04 \| PHIKZ_p05 \| PHIKZ025 | 3 | 12005 | PHIKZ_p04 \| PHIKZ_p05 \| PHIKZ025 | |
| | phiKZ TU010 | 14812 | phiKZ P006 | 12586 | phiKZ T005 | 2226 | - | PHIKZ025 | 1 | 12005 | PHIKZ_p04 \| PHIKZ_p05 \| PHIKZ025 | |

| ϕ | TU | TSS | TSS_ID | TTS | TTS_ID | TU length (bp) | +/- | Genes | #Genes | most distant 3' end | Genes in longest read with TSS | note |
|---|---|---|---|---|---|---|---|---|---|---|---|---|
| | phiKZ TU011 | 19891 | phiKZ P007 | 19111 | phiKZ T007 | 780 | - | - | 0 | 17548 | - | |
| | phiKZ TU012 | 19894 | phiKZ P008 | 20033 | phiKZ T008 | 139 | + | - | 0 | 22110 | - | |
| | phiKZ TU013 | 20123 | phiKZ P009 | 19111 | phiKZ T007 | 1012 | - | PHIKZ028 | 1 | 17332 | PHIKZ028 | |
| | phiKZ TU014 | 20273 | phiKZ P010 | 22435 | phiKZ T010 | 2162 | + | PHIKZ029 | 1 | 23560 | PHIKZ029 \| PHIKZ030 \| PHIKZ_p06 | |
| | phiKZ TU015 | 20273 | phiKZ P010 | 23552 | phiKZ T011 | 3279 | + | PHIKZ029 \| PHIKZ030 \| PHIKZ_p06 | 3 | 23560 | PHIKZ029 \| PHIKZ030 \| PHIKZ_p06 | |
| | phiKZ TU016 | 21311 | phiKZ P011 | 19111 | phiKZ T007 | 2200 | - | PHIKZ028 | 1 | 19109 | PHIKZ028 | |
| | phiKZ TU017 | 21311 | phiKZ P011 | 20489 | phiKZ T009 | 822 | - | - | 0 | 19109 | PHIKZ028 | antisense |
| | phiKZ TU018 | 23599 | phiKZ P012 | 25242 | phiKZ T013 | 1643 | + | PHIKZ031 \| PHIKZ_p07 | 2 | 25686 | PHIKZ031 \| PHIKZ_p07 | |
| | phiKZ TU019 | 26501 | phiKZ P013 | 25208 | phiKZ T012 | 1293 | - | PHIKZ_p08 | 1 | 25206 | PHIKZ_p08 | |
| | phiKZ TU020 | 26543 | phiKZ P014 | 27684 | phiKZ T014 | 1141 | + | PHIKZ033 \| PHIKZ_p09 | 2 | 27687 | PHIKZ033 \| PHIKZ_p09 | |
| | phiKZ TU021 | 27554 | phiKZ P015 | 27684 | phiKZ T014 | 130 | + | - | 0 | 29370 | PHIKZ034 \| PHIKZ035 | |
| | phiKZ TU022 | 27554 | phiKZ P015 | 29374 | phiKZ T015 | 1820 | + | PHIKZ034 \| PHIKZ035 | 2 | 29370 | PHIKZ034 \| PHIKZ035 | |
| | phiKZ TU023 | 27710 | phiKZ P016 | 29374 | phiKZ T015 | 1664 | + | PHIKZ034 \| PHIKZ035 | 2 | 30803 | PHIKZ034 \| PHIKZ035 \| PHIKZ036 | |

| ϕ | TU | TSS | TSS_ID | TTS | TTS_ID | TU length (bp) | +/- | Genes | #Genes | most distant 3' end | Genes in longest read with TSS | note |
|---|---|---|---|---|---|---|---|---|---|---|---|---|
| | phiKZ TU024 | 29413 | phiKZ P017 | 32943 | phiKZ T016 | 3530 | + | PHIKZ036 \| PHIKZ037 \| PHIKZ038 \| PHIKZ039 | 4 | 32942 | PHIKZ036 \| PHIKZ037 \| PHIKZ038 \| PHIKZ039 | |
| | phiKZ TU025 | 30150 | phiKZ P018 | 32484 | - | 2334 | + | PHIKZ037 \| PHIKZ038 | 2 | 32484 | PHIKZ037 \| PHIKZ038 | |
| | phiKZ TU026 | 33472 | phiKZ P019 | 33561 | phiKZ T017 | 89 | + | - | 0 | 34945 | PHIKZ041 \| PHIKZ_p10 \| PHIKZ_p10 \| PHIKZ_p11 \| PHIKZ_p12 | |
| | phiKZ TU027 | 33472 | phiKZ P019 | 34942 | phiKZ T019 | 1470 | + | PHIKZ041 \| PHIKZ_p10 \| PHIKZ_p10 \| PHIKZ_p11 \| PHIKZ_p12 | 5 | 34945 | PHIKZ041 \| PHIKZ_p10 \| PHIKZ_p10 \| PHIKZ_p11 \| PHIKZ_p12 | |
| | phiKZ TU028 | 35112 | phiKZ P020 | 34911 | phiKZ T018 | 201 | - | - | 0 | 34909 | - | |
| | phiKZ TU029 | 39463 | phiKZ P021 | 39545 | phiKZ T020 | 82 | + | - | 0 | 40503 | PHIKZ048 \| PHIKZ049 | |
| | phiKZ TU030 | 39463 | phiKZ P021 | 40406 | phiKZ T021 | 943 | + | PHIKZ048 \| PHIKZ049 | 2 | 40503 | PHIKZ048 \| PHIKZ049 | |
| | phiKZ TU031 | 39463 | phiKZ P021 | 40502 | phiKZ T022 | 1039 | + | PHIKZ048 \| PHIKZ049 | 2 | 40503 | PHIKZ048 \| PHIKZ049 | |
| | phiKZ TU032 | 39935 | phiKZ P022 | 40406 | phiKZ T021 | 471 | + | PHIKZ049 | 1 | 41108 | PHIKZ049 | |
| | phiKZ TU033 | 39935 | phiKZ P022 | 40502 | phiKZ T022 | 567 | + | PHIKZ049 | 1 | 41108 | PHIKZ049 | |
| | phiKZ TU034 | 40441 | phiKZ P023 | 40502 | phiKZ T022 | 61 | + | - | 0 | 43796 | PHIKZ050 \| PHIKZ_p15 \| PHIKZ051 | |
| | phiKZ TU035 | 40441 | phiKZ P023 | 42806 | phiKZ T023 | 2365 | + | PHIKZ050 | 1 | 43796 | PHIKZ050 \| PHIKZ_p15 \| PHIKZ051 | |
| | phiKZ TU036 | 42692 | phiKZ P024 | 42806 | phiKZ T023 | 114 | + | - | 0 | 43799 | PHIKZ_p15 \| PHIKZ051 | |

| ɸ | TU | TSS | TSS_ID | TTS | TTS_ID | TU length (bp) | +/- | Genes | #Genes | most distant 3' end | Genes in longest read with TSS | note |
|---|---|---|---|---|---|---|---|---|---|---|---|---|
| | phiKZ TU037 | 44937 | phiKZ P025 | 45844 | phiKZ T024 | 907 | + | PHIKZ053 | 1 | 48522 | PHIKZ053 \| PHIKZ054 | |
| | phiKZ TU038 | 44937 | phiKZ P025 | 47888 | phiKZ T025 | 2951 | + | PHIKZ053 \| PHIKZ054 | 2 | 48522 | PHIKZ053 \| PHIKZ054 | |
| | phiKZ TU039 | 45377 | phiKZ P026 | 45844 | phiKZ T024 | 467 | + | - | 0 | 52647 | PHIKZ054 \| PHIKZ056 \| PHIKZ_p17 \| PHIKZ057 \| PHIKZ058 \| PHIKZ_p18 \| PHIKZ059 \| PHIKZ055 | |
| | phiKZ TU040 | 45377 | phiKZ P026 | 47888 | phiKZ T025 | 2511 | + | PHIKZ054 | 1 | 52647 | PHIKZ054 \| PHIKZ056 \| PHIKZ_p17 \| PHIKZ057 \| PHIKZ058 \| PHIKZ_p18 \| PHIKZ059 \| PHIKZ055 | |
| | phiKZ TU041 | 45377 | phiKZ P026 | 52650 | phiKZ T028 | 7273 | + | PHIKZ054 \| PHIKZ056 \| PHIKZ_p17 \| PHIKZ057 \| PHIKZ058 \| PHIKZ_p18 \| PHIKZ059 \| PHIKZ055 | 8 | 52647 | PHIKZ054 \| PHIKZ056 \| PHIKZ_p17 \| PHIKZ057 \| PHIKZ058 \| PHIKZ_p18 \| PHIKZ059 \| PHIKZ055 | |
| | phiKZ TU042 | 45490 | phiKZ P027 | 45844 | phiKZ T024 | 354 | + | - | 0 | 50794 | PHIKZ054 \| PHIKZ056 \| PHIKZ_p17 \| PHIKZ055 | |
| | phiKZ TU043 | 45490 | phiKZ P027 | 47888 | phiKZ T025 | 2398 | + | PHIKZ054 | 1 | 50794 | PHIKZ054 \| PHIKZ056 \| PHIKZ_p17 \| PHIKZ055 | |
| | phiKZ TU044 | 45583 | phiKZ P028 | 45844 | phiKZ T024 | 261 | + | - | 0 | 52650 | PHIKZ054 \| PHIKZ056 \| PHIKZ_p17 \| PHIKZ057 \| PHIKZ058 \| PHIKZ_p18 \| PHIKZ059 \| PHIKZ055 | |
| | phiKZ TU045 | 45583 | phiKZ P028 | 47888 | phiKZ T025 | 2305 | + | PHIKZ054 | 1 | 52650 | PHIKZ054 \| PHIKZ056 \| PHIKZ_p17 \| PHIKZ057 \| PHIKZ058 \| PHIKZ_p18 \| PHIKZ059 \| PHIKZ055 | |

| ϕ | TU | TSS | TSS_ID | TTS | TTS_ID | TU length (bp) | +/- | Genes | #Genes | most distant 3' end | Genes in longest read with TSS | note |
|---|---|---|---|---|---|---|---|---|---|---|---|---|
| | phiKZ TU046 | 45583 | phiKZ P028 | 52650 | phiKZ T028 | 7067 | + | PHIKZ054 \| PHIKZ056 \| PHIKZ_p17 \| PHIKZ057 \| PHIKZ058 \| PHIKZ_p18 \| PHIKZ059 \| PHIKZ055 | 8 | 52650 | PHIKZ054 \| PHIKZ056 \| PHIKZ_p17 \| PHIKZ057 \| PHIKZ058 \| PHIKZ_p18 \| PHIKZ059 \| PHIKZ055 | |
| | phiKZ TU047 | 49168 | phiKZ P029 | 50199 | phiKZ T026 | 1031 | + | PHIKZ056 | 1 | 53514 | PHIKZ056 \| PHIKZ_p17 \| PHIKZ057 \| PHIKZ058 \| PHIKZ_p18 \| PHIKZ059 \| PHIKZ060 \| PHIKZ061 | |
| | phiKZ TU048 | 49168 | phiKZ P029 | 50539 | phiKZ T027 | 1371 | + | PHIKZ056 | 1 | 53514 | PHIKZ056 \| PHIKZ_p17 \| PHIKZ057 \| PHIKZ058 \| PHIKZ_p18 \| PHIKZ059 \| PHIKZ060 \| PHIKZ061 | |
| | phiKZ TU049 | 49168 | phiKZ P029 | 53514 | phiKZ T029 | 4346 | + | PHIKZ056 \| PHIKZ_p17 \| PHIKZ057 \| PHIKZ058 \| PHIKZ_p18 \| PHIKZ059 \| PHIKZ060 \| PHIKZ061 | 8 | 53514 | PHIKZ056 \| PHIKZ_p17 \| PHIKZ057 \| PHIKZ058 \| PHIKZ_p18 \| PHIKZ059 \| PHIKZ060 \| PHIKZ061 | |
| | phiKZ TU050 | 51729 | phiKZ P030 | 52650 | phiKZ T028 | 921 | + | PHIKZ_p18 \| PHIKZ059 | 2 | 53528 | PHIKZ_p18 \| PHIKZ059 \| PHIKZ060 \| PHIKZ061 | |
| | phiKZ TU051 | 51729 | phiKZ P030 | 53514 | phiKZ T029 | 1785 | + | PHIKZ_p18 \| PHIKZ059 \| PHIKZ060 \| PHIKZ061 | 4 | 53528 | PHIKZ_p18 \| PHIKZ059 \| PHIKZ060 \| PHIKZ061 | |
| | phiKZ TU052 | 52559 | phiKZ P031 | 52650 | phiKZ T028 | 91 | + | - | 0 | 55165 | PHIKZ060 \| PHIKZ061 \| PHIKZ062 \| PHIKZ063 \| PHIKZ064 | |
| | phiKZ TU053 | 52559 | phiKZ P031 | 53514 | phiKZ T029 | 955 | + | PHIKZ060 \| PHIKZ061 | 2 | 55165 | PHIKZ060 \| PHIKZ061 \| PHIKZ062 \| PHIKZ063 \| PHIKZ064 | |
| | phiKZ TU054 | 52559 | phiKZ P031 | 55165 | phiKZ T030 | 2606 | + | PHIKZ060 \| PHIKZ061 \| PHIKZ062 \| PHIKZ063 \| PHIKZ064 | 5 | 55165 | PHIKZ060 \| PHIKZ061 \| PHIKZ062 \| PHIKZ063 \| PHIKZ064 | |

| ϕ | TU | TSS | TSS_ID | TTS | TTS_ID | TU length (bp) | +/- | Genes | #Genes | most distant 3' end | Genes in longest read with TSS | note |
|---|---|---|---|---|---|---|---|---|---|---|---|---|
| | phiKZ TU055 | 53699 | phiKZ P032 | 55165 | phiKZ T030 | 1466 | + | PHIKZ063 \| PHIKZ064 | 2 | 57874 | PHIKZ063 \| PHIKZ064 \| PHIKZ_p19 \| PHIKZ066 \| PHIKZ067 | |
| | phiKZ TU056 | 57100 | phiKZ P033 | 57939 | phiKZ T031 | 839 | + | PHIKZ067 | 1 | 59512 | PHIKZ067 \| PHIKZ068 | |
| | phiKZ TU057 | 57998 | phiKZ P034 | 62039 | - | 4041 | + | PHIKZ068 \| PHIKZ069 | 2 | 62039 | PHIKZ068 \| PHIKZ069 | |
| | phiKZ TU058 | 64454 | phiKZ P035 | 65671 | phiKZ T032 | 1217 | + | PHIKZ072 \| PHIKZ_p21 | 2 | 67857 | PHIKZ072 \| PHIKZ_p21 \| PHIKZ_p22 | |
| | phiKZ TU059 | 68441 | phiKZ P036 | 68709 | phiKZ T033 | 268 | + | - | 0 | 70374 | PHIKZ075 | |
| | phiKZ TU060 | 68441 | phiKZ P036 | 70298 | phiKZ T034 | 1857 | + | PHIKZ075 | 1 | 70374 | PHIKZ075 | |
| | phiKZ TU061 | 70913 | phiKZ P037 | 71188 | phiKZ T035 | 275 | + | PHIKZ_p24 | 1 | 73363 | PHIKZ_p24 \| PHIKZ_p25 | |
| | phiKZ TU062 | 74086 | phiKZ P038 | 71395 | phiKZ T036 | 2691 | - | PHIKZ_p26 \| PHIKZ077 \| PHIKZ078 | 3 | 71394 | PHIKZ_p26 \| PHIKZ077 \| PHIKZ078 | |
| | phiKZ TU063 | 74086 | phiKZ P038 | 71448 | phiKZ T037 | 2638 | - | PHIKZ_p26 \| PHIKZ077 \| PHIKZ078 | 3 | 71394 | PHIKZ_p26 \| PHIKZ077 \| PHIKZ078 | |
| | phiKZ TU064 | 74150 | phiKZ P039 | 76410 | phiKZ T038 | 2260 | + | PHIKZ_p27 \| PHIKZ080 | 2 | 77892 | PHIKZ_p27 \| PHIKZ080 \| PHIKZ081 | |
| | phiKZ TU065 | 74150 | phiKZ P039 | 77890 | phiKZ T039 | 3740 | + | PHIKZ_p27 \| PHIKZ080 \| PHIKZ081 | 3 | 77892 | PHIKZ_p27 \| PHIKZ080 \| PHIKZ081 | |
| | phiKZ TU066 | 77926 | phiKZ P040 | 80345 | - | 2419 | + | PHIKZ082 \| PHIKZ_p28 | 2 | 80345 | PHIKZ082 \| PHIKZ_p28 | |
| | phiKZ TU067 | 81776 | phiKZ P041 | 80313 | phiKZ T040 | 1463 | - | PHIKZ083 | 1 | 79804 | PHIKZ083 | |

| ϕ | TU | TSS | TSS_ID | TTS | TTS_ID | TU length (bp) | +/- | Genes | #Genes | most distant 3' end | Genes in longest read with TSS | note |
|---|---|---|---|---|---|---|---|---|---|---|---|---|
| | phiKZ TU068 | 81893 | phiKZ P042 | 80313 | phiKZ T040 | 1580 | - | PHIKZ083 | 1 | 80311 | PHIKZ083 | |
| | phiKZ TU069 | 81932 | phiKZ P043 | 84982 | - | 3050 | + | PHIKZ084 \| PHIKZ085 \| PHIKZ_p29 | 3 | 84982 | PHIKZ084 \| PHIKZ085 \| PHIKZ_p29 | |
| | phiKZ TU070 | 81969 | phiKZ P044 | 80313 | phiKZ T040 | 1656 | - | PHIKZ083 | 1 | 79622 | PHIKZ083 | |
| | phiKZ TU071 | 82828 | phiKZ P045 | 84648 | - | 1820 | + | PHIKZ085 | 1 | 84648 | PHIKZ085 | |
| | phiKZ TU072 | 88779 | phiKZ P046 | 91723 | phiKZ T043 | 2944 | + | PHIKZ089 \| PHIKZ090 \| PHIKZ_p30 | 3 | 92773 | PHIKZ089 \| PHIKZ090 \| PHIKZ_p30 | |
| | phiKZ TU073 | 89053 | phiKZ P047 | 91723 | phiKZ T043 | 2670 | + | PHIKZ089 \| PHIKZ090 \| PHIKZ_p30 | 3 | 92538 | PHIKZ089 \| PHIKZ090 \| PHIKZ_p30 | |
| | phiKZ TU074 | 90115 | phiKZ P048 | 91723 | phiKZ T043 | 1608 | + | PHIKZ090 \| PHIKZ_p30 | 2 | 92491 | PHIKZ090 \| PHIKZ_p30 | |
| | phiKZ TU075 | 94330 | phiKZ P049 | 97650 | phiKZ T044 | 3320 | + | PHIKZ094 \| PHIKZ095 | 2 | 98186 | PHIKZ094 \| PHIKZ095 | |
| | phiKZ TU076 | 95917 | phiKZ P050 | 97650 | phiKZ T044 | 1733 | + | PHIKZ095 | 1 | 101366 | PHIKZ095 \| PHIKZ096 \| PHIKZ097 | |
| | phiKZ TU077 | 95917 | phiKZ P050 | 101370 | phiKZ T046 | 5453 | + | PHIKZ095 \| PHIKZ096 \| PHIKZ097 | 3 | 101366 | PHIKZ095 \| PHIKZ096 \| PHIKZ097 | |
| | phiKZ TU078 | 97086 | phiKZ P051 | 97650 | phiKZ T044 | 564 | + | - | 0 | 98432 | - | |
| | phiKZ TU079 | 98764 | phiKZ P052 | 101370 | phiKZ T046 | 2606 | + | PHIKZ097 | 1 | 101371 | PHIKZ097 | |
| | phiKZ TU080 | 98803 | phiKZ P053 | 101370 | phiKZ T046 | 2567 | + | PHIKZ097 | 1 | 102138 | PHIKZ097 | |

| ϕ | TU | TSS | TSS_ID | TTS | TTS_ID | TU length (bp) | +/- | Genes | #Genes | most distant 3' end | Genes in longest read with TSS | note |
|---|---|---|---|---|---|---|---|---|---|---|---|---|
| | phiKZ TU081 | 98803 | phiKZ P053 | 102031 | phiKZ T047 | 3228 | + | PHIKZ097 | 1 | 102138 | PHIKZ097 | |
| | phiKZ TU082 | 101390 | phiKZ P054 | 102031 | phiKZ T047 | 641 | + | - | 0 | 106437 | PHIKZ098 \| PHIKZ099 \| PHIKZ100 \| PHIKZ101 | |
| | phiKZ TU083 | 106304 | phiKZ P055 | 107495 | phiKZ T048 | 1191 | + | PHIKZ_p31 \| PHIKZ_p32 \| PHIKZ103 | 3 | 108038 | PHIKZ_p31 \| PHIKZ_p32 \| PHIKZ103 \| PHIKZ104 | |
| | phiKZ TU084 | 106889 | phiKZ P056 | 107495 | phiKZ T048 | 606 | + | PHIKZ_p32 \| PHIKZ103 | 2 | 108157 | PHIKZ_p32 \| PHIKZ103 \| PHIKZ104 | |
| | phiKZ TU085 | 108843 | phiKZ P057 | 108979 | phiKZ T049 | 136 | + | - | 0 | 109949 | PHIKZ_p33 \| PHIKZ107 | |
| | phiKZ TU086 | 114163 | phiKZ P058 | 114239 | phiKZ T052 | 76 | + | - | 0 | 114302 | - | |
| | phiKZ TU087 | 114456 | phiKZ P059 | 114207 | phiKZ T051 | 249 | - | - | 0 | 114205 | - | |
| | phiKZ TU088 | 114922 | phiKZ P060 | 114207 | phiKZ T051 | 715 | - | PHIKZ_p39 \| PHIKZ_p40 | 2 | 113009 | PHIKZ_p39 \| PHIKZ_p40 | |
| | phiKZ TU089 | 114922 | phiKZ P060 | 114725 | phiKZ T053 | 197 | - | - | 0 | 113009 | PHIKZ_p39 \| PHIKZ_p40 | |
| | phiKZ TU090 | 116062 | phiKZ P061 | 115943 | phiKZ T054 | 119 | - | - | 0 | 115327 | PHIKZ_p41 \| PHIKZ_p42 | |
| | phiKZ TU091 | 117303 | phiKZ P062 | 119656 | - | 2353 | + | PHIKZ119 | 1 | 119656 | PHIKZ119 | |
| | phiKZ TU092 | 117616 | phiKZ P063 | 120629 | phiKZ T057 | 3013 | + | PHIKZ119 \| PHIKZ120 | 2 | 120631 | PHIKZ119 \| PHIKZ120 | |
| | phiKZ TU093 | 117754 | phiKZ P064 | 120629 | phiKZ T057 | 2875 | + | PHIKZ120 | 1 | 120630 | PHIKZ120 | |

| ϕ | TU | TSS | TSS_ID | TTS | TTS_ID | TU length (bp) | +/- | Genes | #Genes | most distant 3' end | Genes in longest read with TSS | note |
|---|---|---|---|---|---|---|---|---|---|---|---|---|
| | phiKZ TU094 | 117934 | phiKZ P065 | 120629 | phiKZ T057 | 2695 | + | PHIKZ120 | 1 | 120629 | PHIKZ120 | |
| | phiKZ TU095 | 118012 | phiKZ P066 | 120629 | phiKZ T057 | 2617 | + | PHIKZ120 | 1 | 120630 | PHIKZ120 | |
| | phiKZ TU096 | 118114 | phiKZ P067 | 120629 | phiKZ T057 | 2515 | + | PHIKZ120 | 1 | 120631 | PHIKZ120 | |
| | phiKZ TU097 | 118184 | phiKZ P068 | 120629 | phiKZ T057 | 2445 | + | PHIKZ120 | 1 | 120992 | PHIKZ120 | |
| | phiKZ TU098 | 119075 | phiKZ P069 | 117934 | phiKZ T055 | 1141 | - | - | 0 | 117511 | - | antisense |
| | phiKZ TU099 | 119075 | phiKZ P069 | 118183 | phiKZ T056 | 892 | - | - | 0 | 117511 | - | antisense |
| | phiKZ TU100 | 130258 | phiKZ P070 | 133099 | phiKZ T060 | 2841 | + | PHIKZ129 | 1 | 133099 | PHIKZ129 | |
| | phiKZ TU101 | 132129 | phiKZ P071 | 133099 | phiKZ T060 | 970 | + | - | 0 | 133099 | - | |
| | phiKZ TU102 | 134420 | phiKZ P072 | 137190 | phiKZ T062 | 2770 | + | PHIKZ131 \| PHIKZ132 | 2 | 140278 | PHIKZ131 \| PHIKZ132 \| PHIKZ133 \| PHIKZ134 | |
| | phiKZ TU103 | 134428 | phiKZ P073 | 133067 | phiKZ T059 | 1361 | - | PHIKZ130 | 1 | 132946 | PHIKZ130 | |
| | phiKZ TU104 | 142840 | phiKZ P074 | 142947 | phiKZ T063 | 107 | + | - | 0 | 145625 | PHIKZ139 \| PHIKZ140 \| PHIKZ141 \| PHIKZ_p44 \| PHIKZ142 | |
| | phiKZ TU105 | 142840 | phiKZ P074 | 145649 | phiKZ T064 | 2809 | + | PHIKZ139 \| PHIKZ140 \| PHIKZ141 \| PHIKZ_p44 \| PHIKZ142 | 5 | 145625 | PHIKZ139 \| PHIKZ140 \| PHIKZ141 \| PHIKZ_p44 \| PHIKZ142 | |
| | phiKZ TU106 | 145659 | phiKZ P075 | 147179 | phiKZ T065 | 1520 | + | PHIKZ143 \| PHIKZ144 \| PHIKZ_p45 | 3 | 149761 | PHIKZ143 \| PHIKZ144 \| PHIKZ_p45 \| PHIKZ145 | |

| ϕ | TU | TSS | TSS_ID | TTS | TTS_ID | TU length (bp) | +/- | Genes | #Genes | most distant 3' end | Genes in longest read with TSS | note |
|---|---|---|---|---|---|---|---|---|---|---|---|---|
| | phiKZ TU107 | 145659 | phiKZ P075 | 149761 | phiKZ T066 | 4102 | + | PHIKZ143 \| PHIKZ144 \| PHIKZ_p45 \| PHIKZ145 | 4 | 149761 | PHIKZ143 \| PHIKZ144 \| PHIKZ_p45 \| PHIKZ145 | |
| | phiKZ TU108 | 146355 | phiKZ P076 | 147179 | phiKZ T065 | 824 | + | PHIKZ144 \| PHIKZ_p45 | 2 | 148437 | PHIKZ144 \| PHIKZ_p45 | |
| | phiKZ TU109 | 146766 | phiKZ P077 | 147179 | phiKZ T065 | 413 | + | PHIKZ_p45 | 1 | 147460 | PHIKZ_p45 | |
| | phiKZ TU110 | 147221 | phiKZ P078 | 149761 | phiKZ T066 | 2540 | + | PHIKZ145 | 1 | 152964 | PHIKZ145 \| PHIKZ146 | |
| | phiKZ TU111 | 147221 | phiKZ P078 | 152963 | phiKZ T067 | 5742 | + | PHIKZ145 \| PHIKZ146 | 2 | 152964 | PHIKZ145 \| PHIKZ146 | |
| | phiKZ TU112 | 148383 | phiKZ P079 | 149761 | phiKZ T066 | 1378 | + | - | 0 | 152965 | PHIKZ146 | |
| | phiKZ TU113 | 148383 | phiKZ P079 | 152963 | phiKZ T067 | 4580 | + | PHIKZ146 | 1 | 152965 | PHIKZ146 | |
| | phiKZ TU114 | 154974 | phiKZ P080 | 153024 | phiKZ T068 | 1950 | - | PHIKZ147 \| PHIKZ148 \| PHIKZ149 | 3 | 152931 | PHIKZ147 \| PHIKZ148 \| PHIKZ149 | |
| | phiKZ TU115 | 155550 | phiKZ P081 | 153024 | phiKZ T068 | 2526 | - | PHIKZ147 \| PHIKZ148 \| PHIKZ149 \| PHIKZ150 | 4 | 152991 | PHIKZ147 \| PHIKZ148 \| PHIKZ149 \| PHIKZ150 | |
| | phiKZ TU116 | 157220 | phiKZ P082 | 153024 | phiKZ T068 | 4196 | - | PHIKZ147 \| PHIKZ148 \| PHIKZ149 \| PHIKZ150 \| PHIKZ151 \| PHIKZ152 | 6 | 152937 | PHIKZ147 \| PHIKZ148 \| PHIKZ149 \| PHIKZ150 \| PHIKZ151 \| PHIKZ152 | |
| | phiKZ TU117 | 157220 | phiKZ P082 | 155687 | phiKZ T069 | 1533 | - | PHIKZ152 | 1 | 152937 | PHIKZ147 \| PHIKZ148 \| PHIKZ149 \| PHIKZ150 \| PHIKZ151 \| PHIKZ152 | |
| | phiKZ TU118 | 157249 | phiKZ P083 | 158242 | phiKZ T071 | 993 | + | PHIKZ153 | 1 | 158989 | PHIKZ153 | |
| | phiKZ TU119 | 159280 | phiKZ P084 | 158210 | phiKZ T070 | 1070 | - | PHIKZ_p46 | 1 | 158151 | PHIKZ_p46 | |

| ϕ | TU | TSS | TSS_ID | TTS | TTS_ID | TU length (bp) | +/- | Genes | #Genes | most distant 3' end | Genes in longest read with TSS | note |
|---|---|---|---|---|---|---|---|---|---|---|---|---|
| | phiKZ TU120 | 161035 | phiKZ P085 | 162450 | phiKZ T074 | 1415 | + | PHIKZ157 | 1 | 164325 | PHIKZ157 \| PHIKZ158 \| PHIKZ_p48 \| PHIKZ160 | |
| | phiKZ TU121 | 161236 | phiKZ P086 | 158210 | phiKZ T070 | 3026 | - | PHIKZ_p46 \| PHIKZ155 \| PHIKZ156 \| PHIKZ_p47 | 4 | 158204 | PHIKZ_p46 \| PHIKZ155 \| PHIKZ156 \| PHIKZ_p47 | |
| | phiKZ TU122 | 161236 | phiKZ P086 | 160356 | phiKZ T072 | 880 | - | PHIKZ_p47 | 1 | 158204 | PHIKZ_p46 \| PHIKZ155 \| PHIKZ156 \| PHIKZ_p47 | |
| | phiKZ TU123 | 164370 | phiKZ P087 | 166843 | phiKZ T075 | 2473 | + | PHIKZ_p49 \| PHIKZ162 | 2 | 167868 | PHIKZ_p49 \| PHIKZ162 \| PHIKZ163 | |
| | phiKZ TU124 | 164370 | phiKZ P087 | 167868 | phiKZ T076 | 3498 | + | PHIKZ_p49 \| PHIKZ162 \| PHIKZ163 | 3 | 167868 | PHIKZ_p49 \| PHIKZ162 \| PHIKZ163 | |
| | phiKZ TU125 | 165041 | phiKZ P088 | 166843 | phiKZ T075 | 1802 | + | PHIKZ162 | 1 | 167868 | PHIKZ162 \| PHIKZ163 | |
| | phiKZ TU126 | 165041 | phiKZ P088 | 167868 | phiKZ T076 | 2827 | + | PHIKZ162 \| PHIKZ163 | 2 | 167868 | PHIKZ162 \| PHIKZ163 | |
| | phiKZ TU127 | 171868 | phiKZ P089 | 172327 | phiKZ T078 | 459 | + | PHIKZ166 | 1 | 173137 | PHIKZ166 \| PHIKZ_p53 \| PHIKZ167 | |
| | phiKZ TU128 | 171868 | phiKZ P089 | 173150 | phiKZ T080 | 1282 | + | PHIKZ166 \| PHIKZ_p53 \| PHIKZ167 | 3 | 173137 | PHIKZ166 \| PHIKZ_p53 \| PHIKZ167 | |
| | phiKZ TU129 | 172165 | phiKZ P090 | 171597 | phiKZ T077 | 568 | - | PHIKZ_p52 | 1 | 170046 | PHIKZ_p52 | |
| | phiKZ TU130 | 172385 | phiKZ P091 | 173039 | phiKZ T079 | 654 | + | PHIKZ_p53 | 1 | 173830 | PHIKZ_p53 \| PHIKZ167 \| PHIKZ168 | |
| | phiKZ TU131 | 172385 | phiKZ P091 | 173150 | phiKZ T080 | 765 | + | PHIKZ_p53 \| PHIKZ167 | 2 | 173830 | PHIKZ_p53 \| PHIKZ167 \| PHIKZ168 | |
| | phiKZ TU132 | 174427 | phiKZ P092 | 173797 | phiKZ T081 | 630 | - | PHIKZ169 | 1 | 172695 | PHIKZ169 | |

| ϕ | TU | TSS | TSS_ID | TTS | TTS_ID | TU length (bp) | +/- | Genes | #Genes | most distant 3' end | Genes in longest read with TSS | note |
|---|---|---|---|---|---|---|---|---|---|---|---|---|
| | phiKZ TU133 | 179469 | phiKZ P093 | 178352 | phiKZ T083 | 1117 | - | PHIKZ175 | 1 | 178301 | PHIKZ175 | |
| | phiKZ TU134 | 181662 | phiKZ P094 | 178352 | phiKZ T083 | 3310 | - | PHIKZ175 \| PHIKZ176 \| PHIKZ177 | 3 | 178304 | PHIKZ175 \| PHIKZ176 \| PHIKZ177 | |
| | phiKZ TU135 | 181662 | phiKZ P094 | 179971 | phiKZ T084 | 1691 | - | PHIKZ177 | 1 | 178304 | PHIKZ175 \| PHIKZ176 \| PHIKZ177 | |
| | phiKZ TU136 | 181662 | phiKZ P094 | 181455 | phiKZ T085 | 207 | - | - | 0 | 178304 | PHIKZ175 \| PHIKZ176 \| PHIKZ177 | |
| | phiKZ TU137 | 182372 | phiKZ P095 | 178352 | phiKZ T083 | 4020 | - | PHIKZ175 \| PHIKZ176 \| PHIKZ177 \| PHIKZ_p55 | 4 | 178301 | PHIKZ175 \| PHIKZ176 \| PHIKZ177 \| PHIKZ_p55 | |
| | phiKZ TU138 | 182372 | phiKZ P095 | 179971 | phiKZ T084 | 2401 | - | PHIKZ177 \| PHIKZ_p55 | 2 | 178301 | PHIKZ175 \| PHIKZ176 \| PHIKZ177 \| PHIKZ_p55 | |
| | phiKZ TU139 | 182372 | phiKZ P095 | 181455 | phiKZ T085 | 917 | - | PHIKZ_p55 | 1 | 178301 | PHIKZ175 \| PHIKZ176 \| PHIKZ177 \| PHIKZ_p55 | |
| | phiKZ TU140 | 187788 | phiKZ P096 | 186346 | phiKZ T086 | 1442 | - | PHIKZ179 | 1 | 184769 | PHIKZ179 | |
| | phiKZ TU141 | 195984 | phiKZ P097 | 196087 | phiKZ T087 | 103 | + | - | 0 | 199194 | PHIKZ182 \| PHIKZ183 \| PHIKZ184 | |
| | phiKZ TU142 | 195984 | phiKZ P097 | 199193 | phiKZ T088 | 3209 | + | PHIKZ182 \| PHIKZ183 \| PHIKZ184 | 3 | 199194 | PHIKZ182 \| PHIKZ183 \| PHIKZ184 | |
| | phiKZ TU143 | 198121 | phiKZ P098 | 199193 | phiKZ T088 | 1072 | + | PHIKZ183 \| PHIKZ184 | 2 | 200147 | PHIKZ183 \| PHIKZ184 \| PHIKZ185 \| PHIKZ_p56 | |
| | phiKZ TU144 | 199232 | phiKZ P099 | 200561 | phiKZ T089 | 1329 | + | PHIKZ185 \| PHIKZ_p56 \| PHIKZ186 | 3 | 203438 | PHIKZ185 \| PHIKZ_p56 \| PHIKZ186 \| PHIKZ187 \| PHIKZ188 \| PHIKZ189 \| PHIKZ190 | |
| | phiKZ TU145 | 199232 | phiKZ P099 | 202344 | phiKZ T090 | 3112 | + | PHIKZ185 \| PHIKZ_p56 \| PHIKZ186 \| PHIKZ187 \| PHIKZ188 \| PHIKZ189 | 6 | 203438 | PHIKZ185 \| PHIKZ_p56 \| PHIKZ186 \| PHIKZ187 \| | |

| ϕ | TU | TSS | TSS_ID | TTS | TTS_ID | TU length (bp) | +/- | Genes | #Genes | most distant 3' end | Genes in longest read with TSS | note |
|---|---|---|---|---|---|---|---|---|---|---|---|---|
| | | | | | | | | | | | PHIKZ188 \| PHIKZ189 \| PHIKZ190 | |
| | phiKZ TU146 | 199463 | phiKZ P100 | 200561 | phiKZ T089 | 1098 | + | PHIKZ_p56 \| PHIKZ186 | 2 | 201187 | PHIKZ_p56 \| PHIKZ186 \| PHIKZ187 | |
| | phiKZ TU147 | 200175 | phiKZ P101 | 200561 | phiKZ T089 | 386 | + | - | 0 | 201142 | PHIKZ187 | |
| | phiKZ TU148 | 203327 | phiKZ P102 | 202268 | - | 1059 | - | - | 0 | 202268 | - | antisense |
| | phiKZ TU149 | 206786 | phiKZ P103 | 207011 | phiKZ T091 | 225 | + | - | 0 | 208212 | PHIKZ199 \| PHIKZ200 | |
| | phiKZ TU150 | 206786 | phiKZ P103 | 208211 | phiKZ T093 | 1425 | + | PHIKZ199 \| PHIKZ200 | 2 | 208212 | PHIKZ199 \| PHIKZ200 | |
| | phiKZ TU151 | 207035 | phiKZ P104 | 208211 | phiKZ T093 | 1176 | + | PHIKZ199 \| PHIKZ200 | 2 | 208212 | PHIKZ199 \| PHIKZ200 | |
| | phiKZ TU152 | 210184 | phiKZ P105 | 210031 | phiKZ T094 | 153 | - | - | 0 | 209529 | - | |
| | phiKZ TU153 | 211034 | phiKZ P106 | 208177 | phiKZ T092 | 2857 | - | PHIKZ201 \| PHIKZ202 | 2 | 208176 | PHIKZ201 \| PHIKZ202 | |
| | phiKZ TU154 | 211034 | phiKZ P106 | 210031 | phiKZ T094 | 1003 | - | PHIKZ202 | 1 | 208176 | PHIKZ201 \| PHIKZ202 | |
| | phiKZ TU155 | 212808 | phiKZ P107 | 210251 | - | 2557 | - | PHIKZ202 \| PHIKZ203 | 2 | 210251 | PHIKZ202 \| PHIKZ203 | |
| | phiKZ TU156 | 217730 | phiKZ P108 | 218207 | phiKZ T098 | 477 | + | PHIKZ210 \| PHIKZ_p59 | 2 | 219955 | PHIKZ210 \| PHIKZ_p59 \| PHIKZ_p60 \| PHIKZ211 \| PHIKZ112 \| PHIKZ_p61 | |
| | phiKZ TU157 | 220796 | phiKZ P109 | 221066 | phiKZ T099 | 270 | + | - | 0 | 222222 | PHIKZ216 \| PHIKZ_p62 \| PHIKZ218 \| PHIKZ_p63 | |

| ϕ | TU | TSS | TSS_ID | TTS | TTS_ID | TU length (bp) | +/- | Genes | #Genes | most distant 3' end | Genes in longest read with TSS | note |
|---|---|---|---|---|---|---|---|---|---|---|---|---|
| | phiKZ TU158 | 220796 | phiKZ P109 | 221335 | phiKZ T100 | 539 | + | PHIKZ216 | 1 | 222222 | PHIKZ216 \| PHIKZ_p62 \| PHIKZ218 \| PHIKZ_p63 | |
| | phiKZ TU159 | 220796 | phiKZ P109 | 221825 | phiKZ T101 | 1029 | + | PHIKZ216 \| PHIKZ_p62 \| PHIKZ218 | 3 | 222222 | PHIKZ216 \| PHIKZ_p62 \| PHIKZ218 \| PHIKZ_p63 | |
| | phiKZ TU160 | 220879 | phiKZ P110 | 221066 | phiKZ T099 | 187 | + | - | 0 | 223924 | PHIKZ_p62 \| PHIKZ218 \| PHIKZ_p63 \| PHIKZ219 \| PHIKZ220 | |
| | phiKZ TU161 | 220879 | phiKZ P110 | 221335 | phiKZ T100 | 456 | + | - | 0 | 223924 | PHIKZ_p62 \| PHIKZ218 \| PHIKZ_p63 \| PHIKZ219 \| PHIKZ220 | |
| | phiKZ TU162 | 220879 | phiKZ P110 | 221825 | phiKZ T101 | 946 | + | PHIKZ_p62 \| PHIKZ218 | 2 | 223924 | PHIKZ_p62 \| PHIKZ218 \| PHIKZ_p63 \| PHIKZ219 \| PHIKZ220 | |
| | phiKZ TU163 | 220879 | phiKZ P110 | 223931 | phiKZ T102 | 3052 | + | PHIKZ_p62 \| PHIKZ218 \| PHIKZ_p63 \| PHIKZ219 \| PHIKZ220 | 5 | 223924 | PHIKZ_p62 \| PHIKZ218 \| PHIKZ_p63 \| PHIKZ219 \| PHIKZ220 | |
| | phiKZ TU164 | 221507 | phiKZ P111 | 221825 | phiKZ T101 | 318 | + | PHIKZ218 | 1 | 223046 | PHIKZ218 \| PHIKZ_p63 | |
| | phiKZ TU165 | 223862 | phiKZ P112 | 223931 | phiKZ T102 | 69 | + | - | 0 | 225791 | PHIKZ221 \| PHIKZ222 \| PHIKZ223 \| PHIKZ224 | |
| | phiKZ TU166 | 223862 | phiKZ P110 | 224732 | phiKZ T103 | 870 | + | PHIKZ221 \| PHIKZ222 | 2 | 225791 | PHIKZ221 \| PHIKZ222 \| PHIKZ223 \| PHIKZ224 | |
| | phiKZ TU167 | 223862 | phiKZ P110 | 225793 | phiKZ T104 | 1931 | + | PHIKZ221 \| PHIKZ222 \| PHIKZ223 \| PHIKZ224 | 4 | 225791 | PHIKZ221 \| PHIKZ222 \| PHIKZ223 \| PHIKZ224 | |
| | phiKZ TU168 | 224867 | phiKZ P113 | 225793 | phiKZ T104 | 926 | + | PHIKZ223 \| PHIKZ224 | 2 | 225986 | PHIKZ223 \| PHIKZ224 \| PHIKZ_p64 | |
| | phiKZ TU169 | 226820 | phiKZ P114 | 227922 | phiKZ T105 | 1102 | + | PHIKZ228 \| PHIKZ229 | 2 | 228569 | PHIKZ228 \| PHIKZ229 \| PHIKZ_p65 \| PHIKZ_p66 | |

| ϕ | TU | TSS | TSS_ID | TTS | TTS_ID | TU length (bp) | +/- | Genes | #Genes | most distant 3' end | Genes in longest read with TSS | note |
|---|---|---|---|---|---|---|---|---|---|---|---|---|
| | phiKZ TU170 | 227508 | phiKZ P115 | 227922 | phiKZ T105 | 414 | + | PHIKZ229 | 1 | 229091 | PHIKZ229 \| PHIKZ_p65 \| PHIKZ_p66 \| PHIKZ_p67 | |
| | phiKZ TU171 | 231207 | phiKZ P116 | 231265 | phiKZ T106 | 58 | + | - | 0 | 233300 | PHIKZ234 \| PHIKZ_p70 \| PHIKZ235 | |
| | phiKZ TU172 | 231207 | phiKZ P116 | 233301 | phiKZ T107 | 2094 | + | PHIKZ234 \| PHIKZ_p70 \| PHIKZ235 | 3 | 233300 | PHIKZ234 \| PHIKZ_p70 \| PHIKZ235 | |
| | phiKZ TU173 | 231384 | phiKZ P117 | 233301 | phiKZ T107 | 1917 | + | PHIKZ_p70 \| PHIKZ235 | 2 | 233681 | PHIKZ_p70 \| PHIKZ235 | |
| | phiKZ TU174 | 232161 | phiKZ P118 | 233301 | phiKZ T107 | 1140 | + | - | 0 | 235274 | PHIKZ236 \| PHIKZ237 \| PHIKZ238 | |
| | phiKZ TU175 | 235361 | phiKZ P119 | 235828 | phiKZ T108 | 467 | + | PHIKZ_p72 | 1 | 236822 | PHIKZ_p72 \| PHIKZ239 \| PHIKZ240 | |
| | phiKZ TU176 | 235469 | phiKZ P120 | 235828 | phiKZ T108 | 359 | + | PHIKZ_p72 | 1 | 236822 | PHIKZ_p72 \| PHIKZ239 \| PHIKZ240 | |
| | phiKZ TU177 | 238722 | phiKZ P121 | 239132 | phiKZ T109 | 410 | + | PHIKZ244 | 1 | 240325 | PHIKZ244 \| PHIKZ245 \| PHIKZ246 | |
| | phiKZ TU178 | 238722 | phiKZ P121 | 240325 | phiKZ T110 | 1603 | + | PHIKZ244 \| PHIKZ245 \| PHIKZ246 | 3 | 240325 | PHIKZ244 \| PHIKZ245 \| PHIKZ246 | |
| | phiKZ TU179 | 240373 | phiKZ P122 | 243801 | phiKZ T111 | 3428 | + | PHIKZ_p73 \| PHIKZ247 \| PHIKZ248 \| PHIKZ249 \| PHIKZ250 \| PHIKZ251 \| PHIKZ252 \| PHIKZ253 | 8 | 243999 | PHIKZ_p73 \| PHIKZ247 \| PHIKZ248 \| PHIKZ249 \| PHIKZ250 \| PHIKZ251 \| PHIKZ252 \| PHIKZ253 \| PHIKZ_p74 | |
| | phiKZ TU180 | 240373 | phiKZ P122 | 243999 | phiKZ T112 | 3626 | + | PHIKZ_p73 \| PHIKZ247 \| PHIKZ248 \| PHIKZ249 \| PHIKZ250 \| PHIKZ251 \| PHIKZ252 \| PHIKZ253 \| PHIKZ_p74 | 9 | 243999 | PHIKZ_p73 \| PHIKZ247 \| PHIKZ248 \| PHIKZ249 \| PHIKZ250 \| PHIKZ251 \| PHIKZ252 \| PHIKZ253 \| PHIKZ_p74 | |

| ϕ | TU | TSS | TSS_ID | TTS | TTS_ID | TU length (bp) | +/- | Genes | #Genes | most distant 3' end | Genes in longest read with TSS | note |
|---|---|---|---|---|---|---|---|---|---|---|---|---|
| | phiKZ TU181 | 243202 | phiKZ P123 | 243801 | phiKZ T111 | 599 | + | PHIKZ252 \| PHIKZ253 | 2 | 244142 | PHIKZ252 \| PHIKZ253 \| PHIKZ_p74 | |
| | phiKZ TU182 | 243202 | phiKZ P123 | 243999 | phiKZ T112 | 797 | + | PHIKZ252 \| PHIKZ253 \| PHIKZ_p74 | 3 | 244142 | PHIKZ252 \| PHIKZ253 \| PHIKZ_p74 | |
| | phiKZ TU183 | 243620 | phiKZ P124 | 243801 | phiKZ T111 | 181 | + | - | 0 | 245240 | PHIKZ_p74 \| PHIKZ254 \| PHIKZ_p75 \| PHIKZ256 | |
| | phiKZ TU184 | 243620 | phiKZ P124 | 243999 | phiKZ T112 | 379 | + | PHIKZ_p74 | 1 | 245240 | PHIKZ_p74 \| PHIKZ254 \| PHIKZ_p75 \| PHIKZ256 | |
| | phiKZ TU185 | 246721 | phiKZ P125 | 248230 | phiKZ T113 | 1509 | + | PHIKZ260 \| PHIKZ_p76 \| PHIKZ261 \| PHIKZ262 | 4 | 250238 | PHIKZ260 \| PHIKZ_p76 \| PHIKZ261 \| PHIKZ262 \| PHIKZ263 \| PHIKZ264 \| PHIKZ265 \| PHIKZ266 | |
| | phiKZ TU186 | 246721 | phiKZ P125 | 248532 | phiKZ T114 | 1811 | + | PHIKZ260 \| PHIKZ_p76 \| PHIKZ261 \| PHIKZ262 \| PHIKZ263 | 5 | 250238 | PHIKZ260 \| PHIKZ_p76 \| PHIKZ261 \| PHIKZ262 \| PHIKZ263 \| PHIKZ264 \| PHIKZ265 \| PHIKZ266 | |
| | phiKZ TU187 | 246721 | phiKZ P125 | 249008 | phiKZ T115 | 2287 | + | PHIKZ260 \| PHIKZ_p76 \| PHIKZ261 \| PHIKZ262 \| PHIKZ263 \| PHIKZ264 | 6 | 250238 | PHIKZ260 \| PHIKZ_p76 \| PHIKZ261 \| PHIKZ262 \| PHIKZ263 \| PHIKZ264 \| PHIKZ265 \| PHIKZ266 | |
| | phiKZ TU188 | 246721 | phiKZ P125 | 250278 | phiKZ T116 | 3557 | + | PHIKZ260 \| PHIKZ_p76 \| PHIKZ261 \| PHIKZ262 \| PHIKZ263 \| PHIKZ264 \| PHIKZ265 \| PHIKZ266 | 8 | 250238 | PHIKZ260 \| PHIKZ_p76 \| PHIKZ261 \| PHIKZ262 \| PHIKZ263 \| PHIKZ264 \| PHIKZ265 \| PHIKZ266 | |
| | phiKZ TU189 | 248091 | phiKZ P126 | 248230 | phiKZ T113 | 139 | + | - | 0 | 250280 | PHIKZ263 \| PHIKZ264 \| PHIKZ265 \| PHIKZ266 | |
| | phiKZ TU190 | 248091 | phiKZ P126 | 248532 | phiKZ T114 | 441 | + | PHIKZ263 | 1 | 250280 | PHIKZ263 \| PHIKZ264 \| PHIKZ265 \| PHIKZ266 | |
| | phiKZ TU191 | 248091 | phiKZ P126 | 249008 | phiKZ T115 | 917 | + | PHIKZ263 \| PHIKZ264 | 2 | 250280 | PHIKZ263 \| PHIKZ264 \| PHIKZ265 \| PHIKZ266 | |

| ϕ | TU | TSS | TSS_ID | TTS | TTS_ID | TU length (bp) | +/- | Genes | #Genes | most distant 3' end | Genes in longest read with TSS | note |
|---|---|---|---|---|---|---|---|---|---|---|---|---|
| | phiKZ TU192 | 248091 | phiKZ P126 | 250278 | phiKZ T116 | 2187 | + | PHIKZ263 \| PHIKZ264 \| PHIKZ265 \| PHIKZ266 | 4 | 250280 | PHIKZ263 \| PHIKZ264 \| PHIKZ265 \| PHIKZ266 | |
| | phiKZ TU193 | 249790 | phiKZ P127 | 250278 | phiKZ T116 | 488 | + | PHIKZ266 | 1 | 250280 | PHIKZ266 | |
| | phiKZ TU194 | 261514 | phiKZ P128 | 262760 | phiKZ T118 | 1246 | + | PHIKZ_p81 \| PHIKZ287 | 2 | 263293 | PHIKZ_p81 \| PHIKZ287 \| PHIKZ288 | |
| | phiKZ TU195 | 261514 | phiKZ P128 | 263292 | phiKZ T119 | 1778 | + | PHIKZ_p81 \| PHIKZ287 \| PHIKZ288 | 3 | 263293 | PHIKZ_p81 \| PHIKZ287 \| PHIKZ288 | |
| | phiKZ TU196 | 262671 | phiKZ P129 | 262760 | phiKZ T118 | 89 | + | - | 0 | 263293 | PHIKZ288 | |
| | phiKZ TU197 | 262671 | phiKZ P129 | 263292 | phiKZ T119 | 621 | + | PHIKZ288 | 1 | 263293 | PHIKZ288 | |
| | phiKZ TU198 | 263172 | phiKZ P130 | 263292 | phiKZ T119 | 120 | + | - | 0 | 264292 | PHIKZ289 | |
| | phiKZ TU199 | 264056 | phiKZ P131 | 265474 | - | 1418 | + | PHIKZ290 \| PHIKZ_p82 \| PHIKZ219 | 3 | 265474 | PHIKZ290 \| PHIKZ_p82 \| PHIKZ219 | |
| | phiKZ TU200 | 265938 | phiKZ P132 | 265438 | phiKZ T120 | 500 | - | PHIKZ_p83 | 1 | 265433 | PHIKZ_p83 | |
| | phiKZ TU201 | 265938 | phiKZ P132 | 265484 | phiKZ T121 | 454 | - | PHIKZ_p83 | 1 | 265433 | PHIKZ_p83 | |
| | phiKZ TU202 | 266394 | phiKZ P133 | 265438 | phiKZ T120 | 956 | - | PHIKZ_p83 \| PHIKZ293 | 2 | 265434 | PHIKZ_p83 \| PHIKZ293 | |
| | phiKZ TU203 | 266394 | phiKZ P133 | 265484 | phiKZ T121 | 910 | - | PHIKZ_p83 \| PHIKZ293 | 2 | 265434 | PHIKZ_p83 \| PHIKZ293 | |
| | phiKZ TU204 | 266808 | phiKZ P134 | 265438 | phiKZ T120 | 1370 | - | PHIKZ_p83 \| PHIKZ293 \| PHIKZ_p84 \| PHIKZ_t01 | 4 | 265434 | PHIKZ_p83 \| PHIKZ293 \| PHIKZ_p84 \| PHIKZ_t01 | |

| ϕ | TU | TSS | TSS_ID | TTS | TTS_ID | TU length (bp) | +/- | Genes | #Genes | most distant 3' end | Genes in longest read with TSS | note |
|---|---|---|---|---|---|---|---|---|---|---|---|---|
| | phiKZ TU205 | 266808 | phiKZ P134 | 265484 | phiKZ T121 | 1324 | - | PHIKZ_p83 \| PHIKZ293 \| PHIKZ_p84 \| PHIKZ_t01 | 4 | 265434 | PHIKZ_p83 \| PHIKZ293 \| PHIKZ_p84 \| PHIKZ_t01 | |
| | phiKZ TU206 | 266808 | phiKZ P134 | 266616 | phiKZ T122 | 192 | - | PHIKZ_t01 | 1 | 265434 | PHIKZ_p83 \| PHIKZ293 \| PHIKZ_p84 \| PHIKZ_t01 | |
| | phiKZ TU207 | 266897 | phiKZ P135 | 267119 | phiKZ T123 | 222 | + | PHIKZ_p91 | 1 | 267398 | PHIKZ_p91 | |
| | phiKZ TU208 | 267851 | phiKZ P136 | 266616 | phiKZ T122 | 1235 | - | PHIKZ_t01 \| PHIKZ294 | 2 | 266614 | PHIKZ_t01 \| PHIKZ294 | |
| | phiKZ TU209 | 267851 | phiKZ P136 | 267142 | phiKZ T124 | 709 | - | PHIKZ294 | 1 | 266614 | PHIKZ_t01 \| PHIKZ294 | |
| | phiKZ TU210 | 269756 | phiKZ P137 | 267142 | phiKZ T124 | 2614 | - | PHIKZ294 \| PHIKZ_p90 \| PHIKZ_p89 \| PHIKZ295 \| PHIKZ_p88 \| PHIKZ296 \| PHIKZ_t02 | 7 | 267089 | PHIKZ294 \| PHIKZ_p90 \| PHIKZ_p89 \| PHIKZ295 \| PHIKZ_p88 \| PHIKZ296 \| PHIKZ_t02 | |
| | phiKZ TU211 | 269756 | phiKZ P137 | 268768 | phiKZ T125 | 988 | - | PHIKZ296 \| PHIKZ_t02 | 2 | 267089 | PHIKZ294 \| PHIKZ_p90 \| PHIKZ_p89 \| PHIKZ295 \| PHIKZ_p88 \| PHIKZ296 \| PHIKZ_t02 | |
| | phiKZ TU212 | 270595 | phiKZ P138 | 267142 | phiKZ T124 | 3453 | - | PHIKZ294 \| PHIKZ_p90 \| PHIKZ_p89 \| PHIKZ295 \| PHIKZ_p88 \| PHIKZ296 \| PHIKZ_t02 \| PHIKZ297 | 8 | 267089 | PHIKZ294 \| PHIKZ_p90 \| PHIKZ_p89 \| PHIKZ295 \| PHIKZ_p88 \| PHIKZ296 \| PHIKZ_t02 \| PHIKZ297 | |
| | phiKZ TU213 | 270595 | phiKZ P138 | 268768 | phiKZ T125 | 1827 | - | PHIKZ296 \| PHIKZ_t02 \| PHIKZ297 | 3 | 267089 | PHIKZ294 \| PHIKZ_p90 \| PHIKZ_p89 \| PHIKZ295 \| PHIKZ_p88 \| PHIKZ296 \| PHIKZ_t02 \| PHIKZ297 | |
| | phiKZ TU214 | 270595 | phiKZ P138 | 270485 | phiKZ T126 | 110 | - | - | 0 | 267089 | PHIKZ294 \| PHIKZ_p90 \| PHIKZ_p89 \| PHIKZ295 \| PHIKZ_p88 \| PHIKZ296 \| PHIKZ_t02 \| PHIKZ297 | |

| ϕ | TU | TSS | TSS_ID | TTS | TTS_ID | TU length (bp) | +/- | Genes | #Genes | most distant 3' end | Genes in longest read with TSS | note |
|---|---|---|---|---|---|---|---|---|---|---|---|---|
| | phiKZ TU215 | 271686 | phiKZ P139 | 268768 | phiKZ T125 | 2918 | - | PHIKZ296 \| PHIKZ_t02 \| PHIKZ297 \| PHIKZ298 | 4 | 268767 | PHIKZ296 \| PHIKZ_t02 \| PHIKZ297 \| PHIKZ298 | |
| | phiKZ TU216 | 271686 | phiKZ P139 | 270485 | phiKZ T126 | 1201 | - | PHIKZ298 | 1 | 268767 | PHIKZ296 \| PHIKZ_t02 \| PHIKZ297 \| PHIKZ298 | |
| | phiKZ TU217 | 271922 | phiKZ P140 | 268768 | phiKZ T125 | 3154 | - | PHIKZ296 \| PHIKZ_t02 \| PHIKZ297 \| PHIKZ298 \| PHIKZ_t03 \| PHIKZ_t04 | 6 | 268766 | PHIKZ296 \| PHIKZ_t02 \| PHIKZ297 \| PHIKZ298 \| PHIKZ_t03 \| PHIKZ_t04 | |
| | phiKZ TU218 | 271922 | phiKZ P140 | 270485 | phiKZ T126 | 1437 | - | PHIKZ298 \| PHIKZ_t03 \| PHIKZ_t04 | 3 | 268766 | PHIKZ296 \| PHIKZ_t02 \| PHIKZ297 \| PHIKZ298 \| PHIKZ_t03 \| PHIKZ_t04 | |
| | phiKZ TU219 | 271922 | phiKZ P140 | 271740 | phiKZ T127 | 182 | - | PHIKZ_t03 \| PHIKZ_t04 | 2 | 268766 | PHIKZ296 \| PHIKZ_t02 \| PHIKZ297 \| PHIKZ298 \| PHIKZ_t03 \| PHIKZ_t04 | |
| | phiKZ TU220 | 271922 | phiKZ P140 | 271822 | phiKZ T128 | 100 | - | PHIKZ_t04 | 1 | 268766 | PHIKZ296 \| PHIKZ_t02 \| PHIKZ297 \| PHIKZ298 \| PHIKZ_t03 \| PHIKZ_t04 | |
| | phiKZ TU221 | 273454 | phiKZ P141 | 271740 | phiKZ T127 | 1714 | - | PHIKZ_t03 \| PHIKZ_t04 \| PHIKZ299 \| PHIKZ_p87 \| PHIKZ_p86 \| PHIKZ_t05 \| PHIKZ_301 \| PHIKZ_t06 \| PHIKZ_t07 | 9 | 271704 | PHIKZ_t03 \| PHIKZ_t04 \| PHIKZ299 \| PHIKZ_p87 \| PHIKZ_p86 \| PHIKZ_t05 \| PHIKZ_301 \| PHIKZ_t06 \| PHIKZ_t07 | |
| | phiKZ TU222 | 273454 | phiKZ P141 | 271822 | phiKZ T128 | 1632 | - | PHIKZ_t04 \| PHIKZ299 \| PHIKZ_p87 \| PHIKZ_p86 \| PHIKZ_t05 \| PHIKZ_301 \| PHIKZ_t06 \| PHIKZ_t07 | 8 | 271704 | PHIKZ_t03 \| PHIKZ_t04 \| PHIKZ299 \| PHIKZ_p87 \| PHIKZ_p86 \| PHIKZ_t05 \| PHIKZ_301 \| PHIKZ_t06 \| PHIKZ_t07 | |
| | phiKZ TU223 | 273454 | phiKZ P141 | 272869 | phiKZ T129 | 585 | - | PHIKZ_t05 \| PHIKZ_301 \| PHIKZ_t06 \| PHIKZ_t07 | 4 | 271704 | PHIKZ_t03 \| PHIKZ_t04 \| PHIKZ299 \| PHIKZ_p87 \| PHIKZ_p86 \| PHIKZ_t05 \| PHIKZ_301 \| PHIKZ_t06 \| PHIKZ_t07 | |

| ϕ | TU | TSS | TSS_ID | TTS | TTS_ID | TU length (bp) | +/- | Genes | #Genes | most distant 3' end | Genes in longest read with TSS | note |
|---|---|---|---|---|---|---|---|---|---|---|---|---|
| | phiKZ TU224 | 273454 | phiKZ P141 | 273267 | phiKZ T130 | 187 | - | PHIKZ_t06 \| PHIKZ_t07 | 2 | 271704 | PHIKZ_t03 \| PHIKZ_t04 \| PHIKZ299 \| PHIKZ_p87 \| PHIKZ_p86 \| PHIKZ_t05 \| PHIKZ_301 \| PHIKZ_t06 \| PHIKZ_t07 | |
| | phiKZ TU225 | 273633 | phiKZ P142 | 271740 | phiKZ T127 | 1893 | - | PHIKZ_t03 \| PHIKZ_t04 \| PHIKZ299 \| PHIKZ_p87 \| PHIKZ_p86 \| PHIKZ_t05 \| PHIKZ_301 \| PHIKZ_t06 \| PHIKZ_t07 | 9 | 271704 | PHIKZ_t03 \| PHIKZ_t04 \| PHIKZ299 \| PHIKZ_p87 \| PHIKZ_p86 \| PHIKZ_t05 \| PHIKZ_301 \| PHIKZ_t06 \| PHIKZ_t07 | |
| | phiKZ TU226 | 273633 | phiKZ P142 | 271822 | phiKZ T128 | 1811 | - | PHIKZ_t04 \| PHIKZ299 \| PHIKZ_p87 \| PHIKZ_p86 \| PHIKZ_t05 \| PHIKZ_301 \| PHIKZ_t06 \| PHIKZ_t07 | 8 | 271704 | PHIKZ_t03 \| PHIKZ_t04 \| PHIKZ299 \| PHIKZ_p87 \| PHIKZ_p86 \| PHIKZ_t05 \| PHIKZ_301 \| PHIKZ_t06 \| PHIKZ_t07 | |
| | phiKZ TU227 | 273633 | phiKZ P142 | 272869 | phiKZ T129 | 764 | - | PHIKZ_t05 \| PHIKZ_301 \| PHIKZ_t06 \| PHIKZ_t07 | 4 | 271704 | PHIKZ_t03 \| PHIKZ_t04 \| PHIKZ299 \| PHIKZ_p87 \| PHIKZ_p86 \| PHIKZ_t05 \| PHIKZ_301 \| PHIKZ_t06 \| PHIKZ_t07 | |
| | phiKZ TU228 | 273633 | phiKZ P142 | 273267 | phiKZ T130 | 366 | - | PHIKZ_t06 \| PHIKZ_t07 | 2 | 271704 | PHIKZ_t03 \| PHIKZ_t04 \| PHIKZ299 \| PHIKZ_p87 \| PHIKZ_p86 \| PHIKZ_t05 \| PHIKZ_301 \| PHIKZ_t06 \| PHIKZ_t07 | |
| | phiKZ TU229 | 273977 | phiKZ P143 | 271740 | phiKZ T127 | 2237 | - | PHIKZ_t03 \| PHIKZ_t04 \| PHIKZ299 \| PHIKZ_p87 \| PHIKZ_p86 \| PHIKZ_t05 \| PHIKZ_301 \| PHIKZ_t06 \| PHIKZ_t07 \| PHIKZ302 | 10 | 270741 | PHIKZ_t03 \| PHIKZ_t04 \| PHIKZ299 \| PHIKZ_p87 \| PHIKZ_p86 \| PHIKZ_t05 \| PHIKZ_301 \| PHIKZ_t06 \| PHIKZ_t07 \| PHIKZ302 | |
| | phiKZ TU230 | 273977 | phiKZ P143 | 271822 | phiKZ T128 | 2155 | - | PHIKZ_t04 \| PHIKZ299 \| PHIKZ_p87 \| PHIKZ_p86 \| PHIKZ_t05 \| PHIKZ_301 \| PHIKZ_t06 \| PHIKZ_t07 \| PHIKZ302 | 9 | 270741 | PHIKZ_t03 \| PHIKZ_t04 \| PHIKZ299 \| PHIKZ_p87 \| PHIKZ_p86 \| PHIKZ_t05 \| | |

| ϕ | TU | TSS | TSS_ID | TTS | TTS_ID | TU length (bp) | +/- | Genes | #Genes | most distant 3' end | Genes in longest read with TSS | note |
|---|---|---|---|---|---|---|---|---|---|---|---|---|
| | | | | | | | | | | | PHIKZ_301 \| PHIKZ_t06 \| PHIKZ_t07 \| PHIKZ302 | |
| | phiKZ TU231 | 273977 | phiKZ P143 | 272869 | phiKZ T129 | 1108 | - | PHIKZ_t05 \| PHIKZ_301 \| PHIKZ_t06 \| PHIKZ_t07 \| PHIKZ302 | 5 | 270741 | PHIKZ_t03 \| PHIKZ_t04 \| PHIKZ299 \| PHIKZ_p87 \| PHIKZ_p86 \| PHIKZ_t05 \| PHIKZ_301 \| PHIKZ_t06 \| PHIKZ_t07 \| PHIKZ302 | |
| | phiKZ TU232 | 273977 | phiKZ P143 | 273267 | phiKZ T130 | 710 | - | PHIKZ_t06 \| PHIKZ_t07 \| PHIKZ302 | 3 | 270741 | PHIKZ_t03 \| PHIKZ_t04 \| PHIKZ299 \| PHIKZ_p87 \| PHIKZ_p86 \| PHIKZ_t05 \| PHIKZ_301 \| PHIKZ_t06 \| PHIKZ_t07 \| PHIKZ302 | |
| | phiKZ TU233 | 274148 | phiKZ P144 | 271740 | phiKZ T127 | 2408 | - | PHIKZ_t03 \| PHIKZ_t04 \| PHIKZ299 \| PHIKZ_p87 \| PHIKZ_p86 \| PHIKZ_t05 \| PHIKZ_301 \| PHIKZ_t06 \| PHIKZ_t07 \| PHIKZ302 | 10 | 271704 | PHIKZ_t03 \| PHIKZ_t04 \| PHIKZ299 \| PHIKZ_p87 \| PHIKZ_p86 \| PHIKZ_t05 \| PHIKZ_301 \| PHIKZ_t06 \| PHIKZ_t07 \| PHIKZ302 | |
| | phiKZ TU234 | 274148 | phiKZ P144 | 271822 | phiKZ T128 | 2326 | - | PHIKZ_t04 \| PHIKZ299 \| PHIKZ_p87 \| PHIKZ_p86 \| PHIKZ_t05 \| PHIKZ_301 \| PHIKZ_t06 \| PHIKZ_t07 \| PHIKZ302 | 9 | 271704 | PHIKZ_t03 \| PHIKZ_t04 \| PHIKZ299 \| PHIKZ_p87 \| PHIKZ_p86 \| PHIKZ_t05 \| PHIKZ_301 \| PHIKZ_t06 \| PHIKZ_t07 \| PHIKZ302 | |
| | phiKZ TU235 | 274148 | phiKZ P144 | 272869 | phiKZ T129 | 1279 | - | PHIKZ_t05 \| PHIKZ_301 \| PHIKZ_t06 \| PHIKZ_t07 \| PHIKZ302 | 5 | 271704 | PHIKZ_t03 \| PHIKZ_t04 \| PHIKZ299 \| PHIKZ_p87 \| PHIKZ_p86 \| PHIKZ_t05 \| PHIKZ_301 \| PHIKZ_t06 \| PHIKZ_t07 \| PHIKZ302 | |
| | phiKZ TU236 | 274148 | phiKZ P144 | 273267 | phiKZ T130 | 881 | - | PHIKZ_t06 \| PHIKZ_t07 \| PHIKZ302 | 3 | 271704 | PHIKZ_t03 \| PHIKZ_t04 \| PHIKZ299 \| PHIKZ_p87 \| PHIKZ_p86 \| PHIKZ_t05 \| PHIKZ_301 \| PHIKZ_t06 \| PHIKZ_t07 \| PHIKZ302 | |

| ϕ | TU | TSS | TSS_ID | TTS | TTS_ID | TU length (bp) | +/- | Genes | #Genes | most distant 3' end | Genes in longest read with TSS | note |
|---|---|---|---|---|---|---|---|---|---|---|---|---|
| | phiKZ TU237 | 274148 | phiKZ P144 | 274007 | phiKZ T131 | 141 | - | - | 0 | 271704 | PHIKZ_t03 \| PHIKZ_t04 \| PHIKZ299 \| PHIKZ_p87 \| PHIKZ_p86 \| PHIKZ_t05 \| PHIKZ_301 \| PHIKZ_t06 \| PHIKZ_t07 \| PHIKZ302 | |
| | phiKZ TU238 | 276905 | phiKZ P145 | 274007 | phiKZ T131 | 2898 | - | PHIKZ303 \| PHIKZ304 | 2 | 274006 | PHIKZ303 \| PHIKZ304 | |
| | phiKZ TU239 | 277797 | phiKZ P146 | 274007 | phiKZ T131 | 3790 | - | PHIKZ303 \| PHIKZ304 \| PHIKZ305 | 3 | 274005 | PHIKZ303 \| PHIKZ304 \| PHIKZ305 | |
| | phiKZ TU240 | 277797 | phiKZ P146 | 276157 | phiKZ T132 | 1640 | - | PHIKZ303 \| PHIKZ304 | 2 | 274005 | PHIKZ303 \| PHIKZ304 \| PHIKZ305 | |
| | phiKZ TU241 | 278754 | phiKZ P147 | 276157 | phiKZ T132 | 2597 | - | PHIKZ303 \| PHIKZ304 | 2 | 276156 | PHIKZ303 \| PHIKZ304 | |
| | phiKZ TU242 | 278754 | phiKZ P147 | 277829 | phiKZ T133 | 925 | - | - | 0 | 276156 | PHIKZ303 \| PHIKZ304 | |
| | phiKZ TU243 | 280201 | phiKZ P148 | 274007 | phiKZ T131 | 6194 | - | PHIKZ303 \| PHIKZ304 \| PHIKZ305 \| PHIKZ_p85 | 4 | 274006 | PHIKZ303 \| PHIKZ304 \| PHIKZ305 \| PHIKZ_p85 | |
| | phiKZ TU244 | 280201 | phiKZ P148 | 276157 | phiKZ T132 | 4044 | - | PHIKZ304 \| PHIKZ305 \| PHIKZ_p85 | 3 | 274006 | PHIKZ303 \| PHIKZ304 \| PHIKZ305 \| PHIKZ_p85 | |
| | phiKZ TU245 | 280201 | phiKZ P148 | 277829 | phiKZ T133 | 2372 | - | PHIKZ_p85 | 1 | 274006 | PHIKZ303 \| PHIKZ304 \| PHIKZ305 \| PHIKZ_p85 | |
| | phiKZ TU246 | 280271 | phiKZ P149 | 276157 | phiKZ T132 | 4114 | - | PHIKZ304 \| PHIKZ305 \| PHIKZ_p85 | 3 | 276156 | PHIKZ304 \| PHIKZ305 \| PHIKZ_p85 | |
| | phiKZ TU247 | 280271 | phiKZ P149 | 277829 | phiKZ T133 | 2442 | - | PHIKZ_p85 | 1 | 276156 | PHIKZ304 \| PHIKZ305 \| PHIKZ_p85 | |
| YuA | YuA T001 | 40 | YuA P001 | 395 | YuA T002 | 355 | + | gp01 | 1 | 1068 | gp01 | |

| ϕ | TU | TSS | TSS_ID | TTS | TTS_ID | TU length (bp) | +/- | Genes | #Genes | most distant 3' end | Genes in longest read with TSS | note |
|---|---|---|---|---|---|---|---|---|---|---|---|---|
| | YuA T002 | 40 | YuA P001 | 471 | YuA T004 | 431 | + | gp01 | 1 | 1068 | gp01 | |
| | YuA T003 | 40 | YuA P001 | 1004 | YuA T005 | 964 | + | gp01 | 1 | 1068 | gp01 | |
| | YuA T004 | 612 | YuA P002 | 60 | YuA T001 | 552 | - | - | 0 | 0 | - | antisense |
| | YuA T005 | 612 | YuA P002 | 411 | YuA T003 | 201 | - | - | 0 | 0 | - | antisense |
| | YuA T006 | 814 | YuA P003 | 1004 | YuA T005 | 190 | + | - | 0 | 1796 | - | |
| | YuA T007 | 1715 | YuA P004 | 967 | - | 748 | - | - | 0 | 967 | - | antisense |
| | YuA T008 | 2349 | YuA P005 | 2477 | YuA T006 | 128 | + | - | 0 | 3589 | gp03 \| gp04 | |
| | YuA T009 | 2349 | YuA P005 | 2607 | YuA T007 | 258 | + | - | 0 | 3589 | gp03 \| gp04 | |
| | YuA T010 | 2349 | YuA P005 | 3188 | YuA T008 | 839 | + | gp3 | 1 | 3589 | gp03 \| gp04 | |
| | YuA T011 | 2349 | YuA P005 | 3504 | YuA T009 | 1155 | + | gp3 \| gp4 | 2 | 3589 | gp03 \| gp04 | |
| | YuA T012 | 2914 | YuA P006 | 3188 | YuA T008 | 274 | + | - | 0 | 4320 | gp04 | |
| | YuA T013 | 2914 | YuA P006 | 3504 | YuA T009 | 590 | + | gp4 | 1 | 4320 | gp04 | |
| | YuA T014 | 5809 | YuA P007 | 6420 | - | 611 | + | - | 0 | 6420 | - | |

| ϕ | TU | TSS | TSS_ID | TTS | TTS_ID | TU length (bp) | +/- | Genes | #Genes | most distant 3' end | Genes in longest read with TSS | note |
|---|---|---|---|---|---|---|---|---|---|---|---|---|
| | YuA T015 | 7758 | YuA P008 | 8128 | - | 370 | + | - | 0 | 8128 | - | |
| | YuA T016 | 14010 | YuA P009 | 14259 | YuA T011 | 249 | + | - | 0 | 14471 | gp16 | |
| | YuA T017 | 22676 | YuA P010 | 22800 | YuA T012 | 124 | + | - | 0 | 22957 | - | |
| | YuA T018 | 23521 | YuA P011 | 22940 | - | 581 | - | - | 0 | 22940 | - | antisense |
| | YuA T019 | 29094 | YuA P012 | 29212 | YuA T014 | 118 | + | - | 0 | 29551 | - | |
| | YuA T020 | 29094 | YuA P012 | 29343 | YuA T015 | 249 | + | - | 0 | 29551 | - | |
| | YuA T021 | 31134 | YuA P013 | 31609 | YuA T016 | 475 | + | gp45 | 1 | 32161 | gp45 \| gp46 | |
| | YuA T022 | 33417 | YuA P014 | 33511 | YuA T017 | 94 | + | - | 0 | 34028 | gp50 | |
| | YuA T023 | 33417 | YuA P014 | 33553 | YuA T018 | 136 | + | - | 0 | 34028 | gp50 | |
| | YuA T024 | 33417 | YuA P014 | 33639 | YuA T019 | 222 | + | - | 0 | 34028 | gp50 | |
| | YuA T025 | 38492 | YuA P015 | 39519 | YuA T021 | 1027 | + | gp55 | 1 | 39863 | gp55 | |
| | YuA T026 | 38492 | YuA P015 | 39736 | YuA T022 | 1244 | + | gp55 | 1 | 39863 | gp55 | |
| | YuA T027 | 39396 | YuA P016 | 39519 | YuA T021 | 123 | + | - | 0 | 40534 | gp56 | |

| ϕ | TU | TSS | TSS_ID | TTS | TTS_ID | TU length (bp) | +/- | Genes | #Genes | most distant 3' end | Genes in longest read with TSS | note |
|---|---|---|---|---|---|---|---|---|---|---|---|---|
| | YuA T028 | 39396 | YuA P016 | 39736 | YuA T022 | 340 | + | - | 0 | 40534 | gp56 | |
| | YuA T029 | 39396 | YuA P016 | 40494 | YuA T023 | 1098 | + | gp56 | 1 | 40534 | gp56 | |
| | YuA T030 | 39396 | YuA P016 | 40533 | YuA T024 | 1137 | + | gp56 | 1 | 40534 | gp56 | |
| | YuA T031 | 44790 | YuA P017 | 45006 | YuA T025 | 216 | + | - | 0 | 45674 | - | |
| | YuA T032 | 44790 | YuA P017 | 45248 | YuA T026 | 458 | + | - | 0 | 45674 | - | |
| | YuA T033 | 44790 | YuA P017 | 45302 | YuA T027 | 512 | + | - | 0 | 45674 | - | |
| | YuA T034 | 44790 | YuA P017 | 45591 | YuA T028 | 801 | + | - | 0 | 45674 | - | |
| | YuA T035 | 46497 | YuA P018 | 47070 | YuA T029 | 573 | + | gp67 | 1 | 47429 | gp67 | |
| | YuA T036 | 46497 | YuA P018 | 47350 | YuA T030 | 853 | + | gp67 | 1 | 47429 | gp67 | |
| | YuA T037 | 47574 | YuA P019 | 47737 | YuA T031 | 163 | + | - | 0 | 48203 | - | |
| | YuA T038 | 47574 | YuA P019 | 47883 | YuA T032 | 309 | + | - | 0 | 48203 | - | |
| | YuA T039 | 52854 | YuA P020 | 53312 | YuA T033 | 458 | + | - | 0 | 55207 | gp72 | |
| | YuA T040 | 57837 | YuA P021 | 57947 | YuA T034 | 110 | + | - | 0 | 58098 | - | |

**Supplementary Table S6: Identification of phage-specific promoters across *Phikmvvirus* members, based on the promoters found in LUZ19 .** Nucleotides indicated in red deviate from the consensus sequence.

| *Phikmvvirus* species | Representative | Accession number | Location of LUZ19-like promoter motif | Sequence |
|---|---|---|---|---|
| LUZ19 | LUZ19 | NC_010326 | 732-751 [-] | GGATATGTCACACACAATGG |
| | | | 21115-21134 [+] | GATTATGTCACCCACATAGG |
| | | | 21312-21331 [+] | GGTTATGTCACCCACAGTGG |
| | | | 35590-35609 [+] | GGTTATGTCACATACAGAGG |
| 15pyo | PaeP_PAO1_1-15pyo | NC_047967.1 | 702-721 [-] | GGTTATGTCACACACAATGG |
| | | | 21055-21074 [+] | GATTATGTCACCCACATAGG |
| | | | 21241-21260 [+] | GGTTATGTCACCCACAGTGG |
| | | | 35519-35538 [+] | GGTTATGTCACATACAGAGG |
| Ab05 | PaeP_PAO1_ Ab05 | NC_026602.1 | 718-737 [-] | GGTTATGTCACACACAATGG |
| | | | 21448-21467 [+] | GATTATGTCACCCACATAGG |
| | | | 21634-21563 [+] | GGTTATGTCACCCACAGTGG |
| | | | 35912-35931 [+] | GGTTATGTCACATACAGAGG |
| ABTNL | PaeP_PPA-ABTNL | NC_027375.1 | 715-734 [-] | GGTTATGTCACACACAATGG |
| | | | 21122-21141 [+] | GATTATGTCACCCACATAGG |
| | | | 21336-21355 [+] | GGTTATGTCACCCACAGTGG |
| | | | 35614-35633 [+] | GGTTATGTCACATACAGAGG |
| DL62 | DL62 | NC_028836.1 | 722-741 [-] | GGTTATGTCACACACAATGG |
| | | | 20680-20699 [+] | GATTATGTCACCCACATAGG |
| | | | 20878-20897 [+] | GGTTATGTCACCCACAGTGG |
| | | | 35156-35175 [+] | GGTTATGTCACATACAGAGG |
| kF77 | phikF77 | NC_012418 | 720-729 [-] | GGTTATGTCACACACAATGG |
| | | | 21013-21031 [+] | GATTATGTCACCCACATAGG |
| | | | 21194-21213 [+] | GATTATGTCACCCACAGTGG |
| | | | 35478-35497 [+] | GGTTATGTCACATACAGAGG |
| LKD16 | LKD16 | NC_009935 | 610-629 [-] | GGTTATGTCACACACAATGG |
| | | | 21073-21092 [+] | GGTTATGTCACCCACATAGG |
| | | | 21253-21272 [+] | GATTATGTCACCCACAGTGG |
| | | | 35547-35566 [+] | GGTTATGTCACATACAGAGG |
| MPK6 | MPK6 | NC_022746 | 740-759 [-] | GGTTATGTCACACACAATGG |
| | | | 21238-21257 [+] | GATTATGTCACCCACATAGG |
| | | | 21451-21470 [+] | GGTTATGTCACCCACAGTGG |

| | | | | |
|---|---|---|---|---|
| | | | 35729-35748 [+] | GGTTATGTCACATACAGAGG |
| MPK7 | MPK7 | NC_022091 | 708-727 [-] | GGTTATGTCACACACAATGG |
| | | | 21225-21244 [+] | GATTATGTCACCCACATAGG |
| | | | 21436-21455 [+] | GGTTATGTCACCCACAGTGG |
| | | | 35715-35734 [+] | GGTTATGTCACATACAGAGG |
| NFS | phiNFS | NC_047852 | 378-397 [-] | GGTTATGTCACACACAATGG |
| | | | 20238-20258 [+] | GATTATGTCACCCACATAGG |
| | | | 20425-20444 [+] | GGTTATGTCACCCACAGTGG |
| | | | 34703-34722 [+] | GGTTATGTCACATACAGAGG |
| PAXYB1 | PAXYB1 | NC_047952 | 743-763 [-] | GGTTATGTCACACACAATGG |
| | | | 21166-21185 [+] | GATTATGTCACCCACATAGG |
| | | | 21379-21398 [+] | GGTTATGTCACCCACAGTGG |
| | | | 35638-35657 [+] | GGTTATGTCACATACAGAGG |
| phiKMV | phiKMV | NC_005045 | 711-730 [-] | GGTTATGTCACACACAATGG |
| | | | 20489-20508 [+] | GATTATGTCACCCACATAGG |
| | | | 20675-20694 [+] | GGTTATGTCACCCACAGTGG |
| | | | 34953-34972 [+] | GGTTATGTCACATACAGAGG |
| PT2 | PT2 | NC_011107 | 702-721 [-] | GGTTATGTCACACACAATGG |
| | | | 20841-20860 [+] | GATTATGTCACCCACATAGG |
| | | | 21038-21057 [+] | GGTTATGTCACCCACAGTGG |
| | | | 35316-35335 [+] | GGTTATGTCACATACAGAGG |
| PT5 | PT5 | NC_011105 | 702-721 [-] | GGTTATGTCACACACAATGG |
| | | | 20854-20873 [+] | GATTATGTCACCCACATAGG |
| | | | 21040-21059 [+] | GGTTATGTCACCCACAGTGG |
| | | | 35318-35337 [+] | GGTTATGTCACATACAGAGG |
| pv130113 | PaeP_130_113 | NC_047953 | 755-774 [-] | GGTTATGTCACACACAATGG |
| | | | 21224-21243 [+] | GATTATGTCACCCACATAGG |
| | | | 21437-21456 [+] | GGTTATGTCACCCACAGTGG |
| | | | 35715-35734 [+] | GGTTATGTCACATACAGAGG |
| RLP | RLP | NC_048168 | 29300-29319 [-] | GGTTATGTCACACACAATGG |
| | | | 6692-6711 [+] | GATTATGTCACCCACATAGG |
| | | | 6914-6933 [+] | GGTTATGTCACCCACAGTGG |
| | | | 21352-21371 [+] | GGTTATGTCACATACAGAGG |

**Supplementary Table S7: Overview of putative phage-encoded ncRNA candidates identified by ONT-cappable-seq.**

| ϕ | TSS | TTS | +/- | TTS ID | Type | Associated gene | RNA fragment length (bp) (<250bp) | Validation | Sequence (5'-3') | RNAfold (kcal/mol) |
|---|---|---|---|---|---|---|---|---|---|---|
| 14-1 | 62561 | 62469 | - | 14-1 T014 | 5'UTR-derived | 141_gp79 | 92 | | gactggcgcggtaaatatcgcgcatatgacaaagtcgttctgttgaa gcgcgactaacaacaaccagcctctgcgactcggcagaggctttct | -37.40 |
| LUZ19 | 21134 | 21207 | + | LUZ19 T009 | 5'UTR-derived | LUZ19_gp27 | 73 | | gagcaagtgcatccgtccaaggccctcgtagagggagcgggggagag gagaggtcagggaagacctggtagagg | -22.80 |
| LUZ19 | 21331 | 21415 | + | LUZ19 T010 | 5'UTR-derived | LUZ19_gp28 | 84 | | gaaggatcgagaacaggcaagggtacggcccgtcctgtcacgttcac ctctcagcagatcgagtggttagaacagaccttccccg | -26.50 |
| LUZ24 | 5249 | 5415 | + | LUZ2 T006 | 5'UTR-derived | LUZ24_gp14 | 166 | | gagtgtcctgtacctgcaccctcgggaactccccgtgtatttcgtgg cgatacccgagggtgaaactctcaaggtgaaggtgtgatgtactatg acggaacagcagcgtcgcggccccttctaagagggcgttagtaatgc ccctaggaactccctggggggtattgc | -69.60 |
| LUZ24 | 39133 | 38968 | - | LUZ2 T024 | 5'UTR-derived | LUZ24_gp62 | 165 | | gatatctgatgcccgccaactggctgtactcgatgcagcacgtaagt gggctgaatctcaggtccgtcgcaaggctgccctagagaagaaggag gagactgaactacctgccgcagttaaacctgctgctcgtagggcgga agcctcagagggttctaagcgggtt | -57.90 |
| PAK_P3 | 31147 | 31308 | + | PAK_P3 T013 | 5'UTR-derived | PAK_P30049 | 161 | | tggagtcttccaagtgtcttaccctctgaacaagggggagcgtggaa gctacaagcacctctatgacaagcaggagaaggagcgaaaggcgaac aacccgcctcccgagccaccacctctcctgctggcgtaaccctaagg ggccctgattgggccctttttt | -63.80 |
| PAK_P3 | 41196 | 41420 | + | PAK_P3 T018 | 5'UTR-derived | PAK_P30063 | 224 | | tcgctctctgcgttgatccctcagaaggtcactcctctttgccctgt gtgctgttcggacgatcttcgtcccactggcaagctgtactacaccg cagtgtcggccttcgaggttctccgctgccatagctgtgggaagcac tcccgcacacgtcagaacgtcctggaccgggaacacctggtagtggg catctgacacaaagcccaccttcgggtgggctctttc | -91.80 |
| PAK_P3 | 51944 | 52047 | + | PAK_P3 T023 | 5'UTR-derived | PAK_P30085 | 103 | | tgcgaatacctcgcctttggcaaggctggcaatacggtgcctgtata cgtcaagtaacagcagagtgcttttagggttgcccctggacgccggg ggcagcttgc | -41.70 |

| ϕ | TSS | TTS | +/- | TTS ID | Type | Associated gene | RNA fragment length (bp) (<250bp) | Validation | Sequence (5'-3') | RNAfold (kcal/mol) |
|---|---|---|---|---|---|---|---|---|---|---|
| PAK_P3 | 85248 | 85458 | + | PAK_P3 T039 | 3'UTR-derived | PAK_P3_gp162 | 210 | Northern blots (JVO-21131), RNA-seq (sRNA1; Chevallereau et al. 2016) | atccatgagattgagggctatgtcctcggctcttgtggtgtaggggc ctcccagccccactaggcgttcccggagccgtatggcgggaggggtt gaaggttcaccacacgaacccccctacgccccggaggttgccggggta ggtcctccaggctctaaagtcctggagaagtgtcagccggacctttg aggatgggagtccaccggcactc | -94.00 |
| PAK_P3 | 85937 | 86017 | + | PAK_P3 T040 | 5'UTR-derived | PAK_P3tRNA01 | 80 | | ttcagatgtgggaccgacgaagctgatgtgcagaggaggtgatccag catctcccggacgggtccgggttaccaacccgct | -25.70 |
| PAK_P3 | 86307 | 86426 | + | PAK_P3 T041 | 3'UTR-derived | PAK_P3_gp163 / tRNA region | 119 | Northern blots (JVO-21132), RNA-seq (sRNA2; Chevallereau et al. 2016) | gctgtgtaagttggactgttgcagtgaagcgataggctccccggatc gtaagccggtaacgtggggatagagggctcccaccgctgagcaacag gaggttcgattcctcaacacagcccc | -42.30 |
| phiKZ | 19894 | 20033 | + | phiKZ T008 | 5'UTR-derived | PHIKZ029 | 139 | | ggaataccagtacgttgttcaatctttttatatagacgagcatcaac gactagcttaggattatcacgatcaccgatattgacaatttgagtaa atttcttaatatcctcaaaatcatcaactggggtaatctgtccact | -21.70 |
| phiKZ | 27554 | 27684 | + | phiKZ T014 | 5'UTR-derived | PHIKZ034 | 130 | | ttgatagtaatggcgatgtcatcatacaactagagcaatcattaatt tatactggttgctctttatctaaatttaatgcgtgttttaacactat gtaaatataatgccctctccttatggagagggcttat | -38.70 |
| phiKZ | 40441 | 40502 | + | phiKZ T022 | 5'UTR-derived | PHIKZ050 | 61 | | gtgaatgtaagttgatgtgctacctctcaatttagggtacccgtata gggtacccattcttt | -21.30 |
| phiKZ | 42692 | 42806 | + | phiKZ T023 | 5'UTR-derived | PHIKZ_p15 | 114 | | tgtattctatctaaccttagagtccctaggtattttccttattgata aaaataacgatcgactcatctccgacttctattaagaaatatagcct cccctagtgggaggcttatat | -23.40 |
| phiKZ | 52559 | 52650 | + | phiKZ T028 | 5'UTR-derived | PHIKZ060 | 91 | | ggcatctggacgcaggtgaagaagatccaacttcagtcgtaggtaat cttctaggttaaataaatgagagtccctacatggggctctcttct | -32.20 |

| ϕ | TSS | TTS | +/- | TTS ID | Type | Associated gene | RNA fragment length (bp) (<250bp) | Validation | Sequence (5’-3’) | RNAfold (kcal/mol) |
|---|---|---|---|---|---|---|---|---|---|---|
| phiKZ | 108843 | 108979 | + | phiKZ T049 | 5'UTR-derived | PHIKZ_p33 | 136 | | gagcaatattgaaaaacatcgactggatgtaattcgtaataagctgt cgatcatttatggcgatgactatacgaatcaagtattcgatacagtt cgtaagatgtaaaaatggatagggcctcactgaggccccattc | -38.10 |
| phiKZ | 114922 | 114725 | - | phiKZ T053 | 5'UTR-derived | PHIKZ_p40 | 197 | Northern blots (JVO-18831), Grad-Seq (5’UTR_gp40; Gerovac et al. 2021) | gtattgatcattctgattaagtcaaattagttaataaacaaatcaat tatcaacgagtgataacgggtattgattactagctaaagagattaat taggatatctacatgaggagtcgctaccccatgtattctggagtttc tctttagcgggagacttaactccacttgggagccgctttagcgggcg gctctcactt | -65.10 |
| phiKZ | 142840 | 142947 | + | phiKZ T063 | 5'UTR-derived | PHIKZ139 | 107 | | aagtaaaacgctcccataatggacaatcatggatctgtgtaatggta cgtggtgctcctgaaaaatttgcttaacagcataaatcccctccata tggaggggaactct | -29.60 |
| phiKZ | 195984 | 196087 | + | phiKZ T087 | 5'UTR-derived | PHIKZ182 | 103 | | agctccctctcaacgtcctggtcgggtagtggataaccgtccattgc caattagactacgttaatctattaaaggagcctccctattgggaggt tccttatttt | -44.00 |
| phiKZ | 206786 | 207011 | + | phiKZ T091 | 5'UTR-derived | PHIKZ199 | 225 | | ggaccacgatgtcgcaacctttgtcctaacacataataaaattagcg acgaattacttaaagaattgatgagtgttgagaatcctaaagctgta ctcaatcactataaaattcctgtaaagatttctagaaatagaataat tgcagcttgtcaaataatcttcaatattcgtggactaaaagtctaat acttttataggagagaaccatgatggttctctcttttt | -53.40 |
| phiKZ | 223862 | 223931 | + | phiKZ T102 | 5'UTR-derived | PHIKZ221 | 69 | Northern blots (JVO-21411) | tagtagtatttctattacgtacctttcgaggcatatactccatgact tatgttgtggggtatatgtgtt | -26.40 |
| phiKZ | 231207 | 231265 | + | phiKZ T106 | 5'UTR-derived | PHIKZ234 | 58 | Northern blots (JVO-21410) | aggtaagatactagccgattggttagttgcgcttagggagtccttcg ggactcccgtat | -26.80 |
| phiKZ | 248091 | 248230 | + | phiKZ T113 | 5'UTR-derived | PHIKZ263 | 139 | | aatcacgatgcagcaaagcaatcgtcagcttttatgcatcacgatag ccagttcaataatgtacgtattgaattttggcgcgaaccatcagcgc gaattatctcgctgattaaagatgcggtcgcatgaccgctttcttt | -49.20 |

| ϕ | TSS | TTS | +/- | TTS ID | Type | Associated gene | RNA fragment length (bp) (<250bp) | Validation | Sequence (5’-3’) | RNAfold (kcal/mol) |
|---|---|---|---|---|---|---|---|---|---|---|
| phiKZ | 262671 | 262760 | + | phiKZ T118 | 5'UTR-derived | PHIKZ288 | 89 | | tagttcctttacgattgctcttttatcttagtttcagttttgctgaa gaagtaccaatgagataccctcccctgatcggggagggttttt | -26.40 |
| phiKZ | 263172 | 263292 | + | phiKZ T119 | 5'UTR-derived | PHIKZ289 | 120 | | taaagaatacccaaatcgggatctcccggactatctcccttcagcgt tcccgtccgcaaatgaaggtgccgccgagatcccacctattgataat gccctcccttcggggagggctttattt | -46.60 |
| phiKZ | 270595 | 270485 | - | phiKZ T126 | 5'UTR-derived | PHIKZ297 | 110 | Northern blots (JVO-18830 & JVO-21375), Grad-Seq (3'UTR_PHIKZ298; Gerovac et al. 2021) | taagagtactataataaagtattcttaaatctatccactaaacacac tcggtggtagaacttattatagagtgtgtctaaatgccaggggtttg ccacccctggttatat | -40.30 |
| phiKZ | 274148 | 274007 | - | phiKZ T131 | 5'UTR-derived | PHIKZ302 | 141 | Northern blots (JVO-21409) | aatataaagcctccccgaagggaggcgatatatcgtcattaatgccc agcaaagagttttactctctgggagccagttggaaggggggactggct ccgatctattatgactaatgtagaaacaaggggggaaggattatttct ac | -48.70 |
| YuA | 2349 | 2477 | + | YuA T006 | 5'UTR-derived | YuA_gp03 | 128 | | agcttcaagctgatcgaccgccttcagtcggaggaccgcgcccaccg tatcgggcagaccaacaacgtgctgtacatcgacctcgtggctgagg actccgtggatgagaaggtggttgaagcgttgagg | -50.30 |
| YuA | 39396 | 39519 | + | YuA T021 | 5'UTR-derived | YuA_gp56 | 123 | | acttttggttgtatagccagaggcgaaggcccacaggagtgattccg cgggccgcgacggaacaccgggtgatccggcacggctgaaaccgctt gattcatcaacttctgaaggagggccaatc | -48.20 |
| YuA | 57837 | 57947 | + | YuA T034 | 5'UTR-derived | YuA_gp77 | 110 | | cctcggaggtctgagcctatcttgggcgcaccgcgaccggacccagc aggtggtccgtccgatcatcgaccactccgcggcgaacatcggcccg gagccgggggtgacc | -46.60 |
